# Linking B-factor and temperature-induced conformational transition

**DOI:** 10.1101/2023.03.12.532309

**Authors:** Fernando de Sá Ribeiro, Luís Maurício T. R. Lima

## Abstract

The crystallographic B-factor, also called temperature factor or Debye-Waller factor, has long been used as a surrogate for local protein flexibility. However, the use of the absolute B-factor as a probe for protein motion requires reproducibility and intervalidation against chemical and physical variables. Here we report the investigation of the thermal dependence of the crystallographic B-factor and its correlation with protein conformational changes. We solved the B-factor reproducibility issue at high resolution (1.5 Å) over a broad temperature range (100 K to 325 K) by protecting crystals with hydrocarbon grease during data collection. We found that the crystallographic protein conformation varies as a function of temperature. Further, the demonstrated that the thermal dependence of B-factor as a function of temperature were similar for all atoms (Cα, N-amide and side chains), without local variations, indicating lack of correlation between temperature-dependent conformational change and the B-factor. These data indicate a linear correlation of B-factor with temperature due to global rigid body motion.

## Introduction

Protein conformational polymorphism and dynamics are part of the proteostasis network involving synthesis and folding, conformational maintenance and degradation [1]. While high resolution structure of macromolecules provides conformational snapshots, molecular dynamics are also required to infer mechanisms required for the understanding of biological systems.

There is increasing prevalence of computational modeling approaches in the study of macromolecular structure and dynamics [2,3]. However, wet-bench techniques are still required [4]. Structure determination of macromolecules can be performed by a large set of state-of-the-art methods, such as in solid phase from crystals (nano to micro; powder, poly or single) by diffraction (electron, x-ray, neutron, XFEL), or in solution by NMR and vitreous by cryo-EM. Time-resolved techniques can also infer dynamics directly such as NMR and ultrafast serial femtosecond crystallography (SFX) techniques [5,6].

Conformational polymorphism and dynamics may also be inferred by solution NMR techniques, or from ensemble structure solved at varying conditions such as chemical, including pH, ligands, cosolutes, and hydration changes [7,8] or physical such as temperature and pressure [9–11].

Single-crystal X-ray diffraction (SCXRD) has been used for over 60 years in solving protein structure [12], and holds an outstanding position in structural biology for being a highly reproducible technique [13–15]. While the high-resolution crystal structure reflects the average coordinate position of the proteins in the crystal, the crystallographic B-factor would allow inferring conformational flexibility from this ensemble. However, It has been argued that the B-factor is highly inaccurate, requiring experimental and analytical improvements [16].

The B-factor – often called temperature factor [17–20] - relates to the mean-square atomic displacement (*X*^2^) by the Debye-Waller function

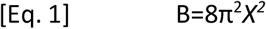

where *X*^2^ embodies additive crystal contributions from atomic fluctuation (*X*^2^_A_), conformational motions (*X*^2^_C_), rigid body vibration (*X*^2^ _RB_) and crystal lattice disorders (*X*^2^ _LT_) [21]. The crystallographic B-factor is also susceptible to variables from crystal (such as hydration, cosolutes, cryoprotection), XRD setup (such as beam, detector), gas stream for temperature control and diffuse scattering [14]. Diffraction shows contributions from both ordered lattice and diffuse scattering [22–25], while diffuse scattering can show contributions from varying sources including rigid-body motion [26] and background scattering [14]. For these reasons, direct correlations between B-factor with protein dynamics and consequent propensity for conformational changes are still controverse [27].

Attempts to solve the uncertainty problem in B-factor backs from decades ago. The broad distribution of B-factor can be empirically scaled down by its average B-factor (B_avg_) resulting in normalized B-factor (B_norm_) [17] in the form of

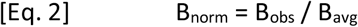

which differ from the *Z-score* as proposed elsewhere [27,28].

We have recently shown that B-factor reproducibility can be achieved by an empirical scaling [15]. The reproducibility was accessed using used 20 crystals of lysozyme collected in 2 different XRD setups, as well as reanalyzed data from human insulin from four dissimilar biotechnology processes collected at four dissimilar XRD setups [13]. The B_norm_ for each amino acid showed low standard deviation among independently collected a processed dataset. We further analyzed data from RCSB, and found that local variations in B_norm_ was related to space group. The correlation between B-factor and protein dynamic remained to be solved, requiring further investigation preferably by using variables to promoting change in conformation and B-factor, separating contributions from static (temperature independent) and dynamic (temperature-dependent) disorder [21,29].

The temperature influence over protein conformation has long been investigated by crystallography for ribonuclease from 98 to 325 K [29], high-pressure (200 MPa, 2 kbar) cryocooled thaumatin from 80 to 160 K [30], thaumatin from 25 to 300 K [10], myoglobin from 80 to 240 K [31], and hen egg white lysozyme (HEWL) solved at cryogenic and room temperatures and shown to vary in conformation and B-factor [32,33] resulting in expansion of about 0.9 % per 100 K.

Analysis of tetragonal lysozyme data currently deposited in the RCSB evidences a gap in the temperature dimension resulting in two subgroups of structures (**Fig S1**), one in the cryogenic range (about 100 K) and another close to room temperature (at about 300 K, with highest temperature at 308 K, i.e., 35 °C). We also observe a large dispersion in cell parameters (**Fig. S1A** and **Fig S1B**) and the lack of direct correlation between them (**Fig. S1D**). There is no apparent correlation between crystallization pH and data collection temperature (**Fig. S1E**), or with the average B-factor for the final structural model (**Fig. S1F**), although a clear correlation is found between the Wilson and average structural B-factor (**Fig. S1G**). From these data it became evident the lack of robust information on the effects of data collection temperature on crystallographic and structural parameters, requiring a systematic approach to use broad-range temperature as a physical variable in protein crystallography.

A major limitation in using physico-chemical variables in the study of B-factor is the lack of experimental repeatability and reproducibility in raw (non-normalized) B-factor, mostly due to dehydration. In small-molecule crystallography, non-dehydration agents are routinely used, such as oils and greases. In protein crystallography, non-dehydration techniques involves capillaries [34], polymer tubes (such as CrystalCap®, Hampton Research), uncured glue / epoxy cement [29,35], silicon nitride wafers (Silson®) [36], lipid cubic phase (LCP) [37], oil [10,38,39], agarose/gelatin [40], poly-acrylamide [41] mineral oil-based grease [42] shortening [43] and animal fat [44,45].

In this study we aimed to solve the experimental reproducibility of B-factor and elucidate the temperature dependence of B-factor. We solved the structure of tetragonal lysozyme over a broad temperature range (from 100 K to 325 K, 25 K intervals), from 30 independent crystals immersed into hydrocarbon grease, and discuss the potential correlation between observed conformational changes and B-factors.

## Results

### Reproducible crystal diffraction at high resolution from 100 K to 325 K

While glycerol is an effective cryoprotectant, higher temperatures requires further protection against dehydration. For this purpose, the crystals were embedded into hydrocarbon grease before data collection. We were able to diffract at 1.5 Å resolution from 100 K to 325 K (52 °C) in 25 K intervals, in triplicate for each temperature. Attempts to diffract at higher temperature resulted in progressive loss in order as indicated by loss of diffraction data (not shown). All dataset were obtained under similar data collection strategy, automatically chosen by the controller program, and the crystal structures were solved using the same workflow, avoiding individual interpretations and interventions.

We observed a positive dependence of the unit cell lengths on data collection temperature, for both a=b (**Fig. S2A**) and c (**Fig. S2B**), and a positive correlation between these dimensions (**Fig. S2C**). The increase in unit cell lengths corresponded to a volume expansion of the cell (**Fig. S3A**), as well as in the increase in volume of the protein in the asymmetric unit (**Fig. S3B**), with no significant changes in the protein accessible surface area (**Fig. S3C**). The volumetric changes in cell unit and protein showed positive correlation (**Fig. S3D**), which might be related to contribution from the protein to the expansion of the cell. A positive correlation between ASA and the unit cell volume (**Fig. S3E**) and with the protein volume (**Fig. S3F**) was also found, despite minor correlation between ASA and temperature (**Fig. S3C**).

### Loss of diffraction power due to protein unfolding over 325 K

We measured the thermal denaturation of lysozyme in sodium acetate buffer pH 4.6 by circular dichroism (CD). A progressive decrease in ellipticity was observed by increasing temperature from 15 °C to 90 °C in 5 °C steps (**Fig. S4A**). The thermal denaturation curves for each wavelength demonstrate a decrease in the steepness of the transitions as a function of the wavelength, converging to about 360 K for complete thermal denaturation transition (**Fig. S4B**). We adjusted the thermal denaturation curves for each wavelength with a sigmoidal function, obtaining both the half-transition temperature (T_50%_) and the slope of the transition (**Fig. S4C**). The T_50%_ for thermal denaturation of lysozyme was found to be about 310 K in the 212 nm to 225 nm wavelength range. These data indicate that the temperature range of 340 K to 360 K are compatible with denaturation of lysozyme in solution in buffer conditions close to the crystallization milieu.

We further probed the overall conformational distribution of lysozyme by ion mobility spectrometry coupled to mass spectrometry by electrospray ionization (ESI-IMS-MS). We evaluated the overall ion mobility distribution for lysozyme at varying charged state as a function of source temperature, from 50 °C to 250 °C in 50 °C intervals (**Fig. S5**). Attempts for measurement with source temperature below 50 °C resulted in spectra with insufficient signal. We have found a progressive drift and broadening in the mobility as a function of increasing temperature, indicative of changes in the protein conformational landscape. Collectively, these spectroscopic data (CD and IMS) indicate the thermal instability of lysozyme in solution at about 360 K and higher temperatures.

### Temperature induces local conformational changes in crystal phase

A pairwise global structural alignment of the lysozyme structures demonstrates an average root-mean square deviation (RMSD) up to 0.3 Å for Cα (**Fig. S6**). Superposition of the protein structures showed global folding similarity in lysozyme conformation, with local changes for both backbone (**Fig. 1A**) and side chains (**Fig. 1B**). Overall deviation in side chains (**Fig. 1C**), Cα (**Fig. 1D**) and amide nitrogen (N-amide; **Fig. 1E**) showed a positive linear correlation with temperature. No overall change in secondary structure content were observed in crystal structures as a function of data collection temperature in the range of 100 K to 325 K (**Fig. 1F**).

**Figure 1.**
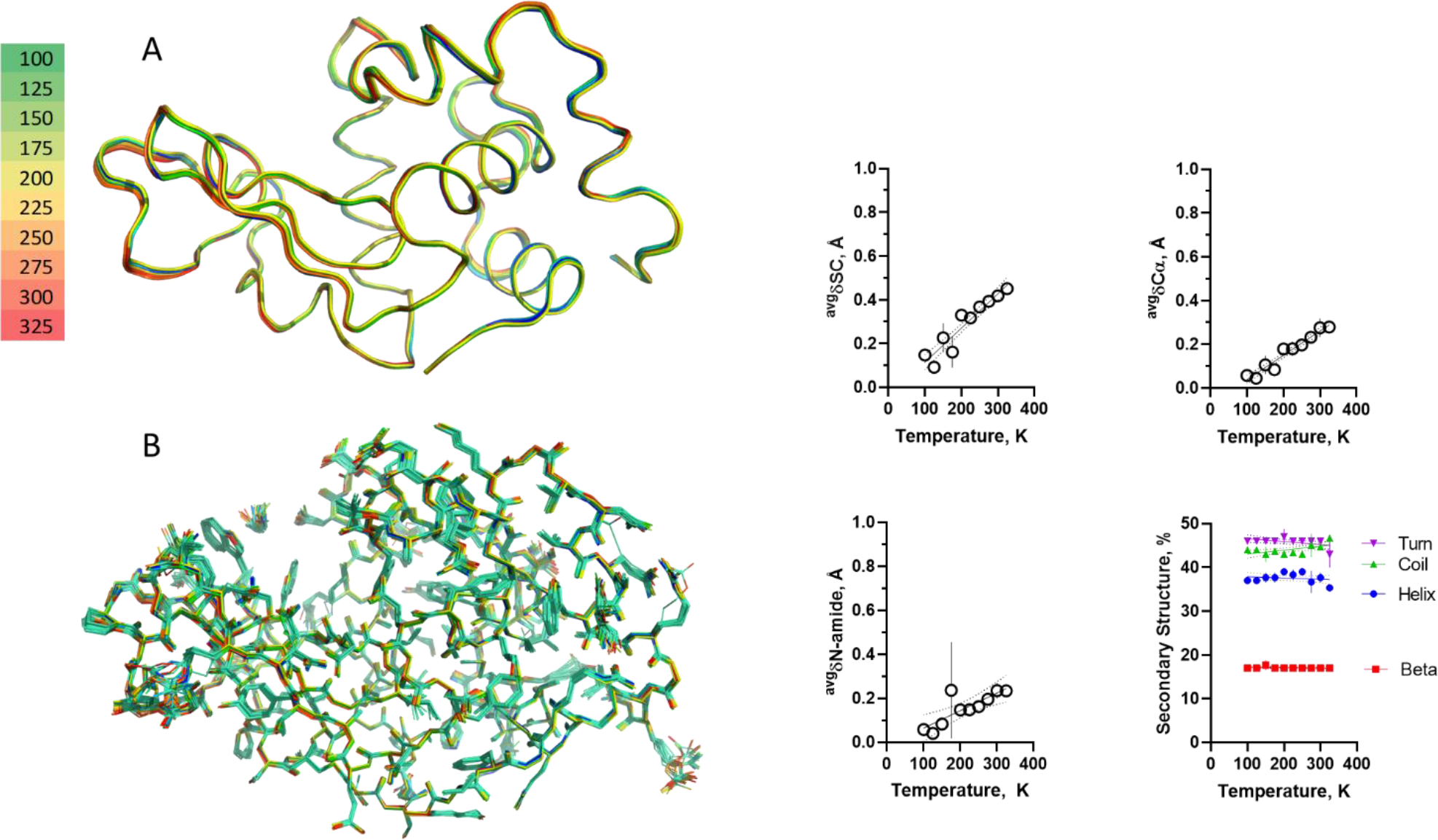
Global temperature-dependent conformational changes. Lysozyme structure models solved from data collected at varying temperatures from 100 K to 325 K were aligned and are showed for A) backbone and B) side chains. Colors represents data collection temperature as indicated. n=3 per temperature. Global (average values for residues 1-129) changes in conformation were inferred from superposition with a reference structure at 100 Kelvin. Continuous line are first order linear regression and dotted lines are 95 % confidence interval. We observed a small (< 1 Å) but progressive and significant increase in global conformational changes as inferred for distance changes in (C) SC (0.001566 Å.K-1; p≤ 0.0001), (D) Cα (0.001110 Å.K^-1^; p≤ 0.0001) and (E) amide nitrogen (0.0007971 Å.K^-1^; p= 0.0037) from reference structure. No significant temperature-dependent changes were inferred for total content in helix (p=0.4260), beta (p=0.3935), coil (p=0.0107) and turn (p=0.0458) as shown in (F). Symbol is average and bar is standard deviation (n=3).

A general overview of conformational changes as a function of data collection temperature could be further inferred from the change in Cα RMSD distribution over the polypeptide chain at varying temperatures using a reference structure at 100 K (**Fig. 2**). The positional drift induced by temperature was analyzed by first-order linear regression (**Fig. 2B; Fig. S7**), and by non-linear regression with a single exponential (**Fig. 2D; Fig. S8**). The thermal constant for conformational changes **k** was distributed as a function of polypeptide chain for first-order linear regression (**Fig. 2C**) and for non-linear exponential regression (**Fig. 2E**). The correlation coefficient for the regressions were higher at regions with higher conformation-temperature constants, and apparently higher for first-order polynomial function compared to exponential function (**Fig. S9**), indicating better adequacy of the linear function for this analysis. The thermal expansion of the lysozyme was discontinuous along the polypeptide chain (**Fig. 2C**), indicating that it occurs preferably in certain regions.

**Figure 2.**
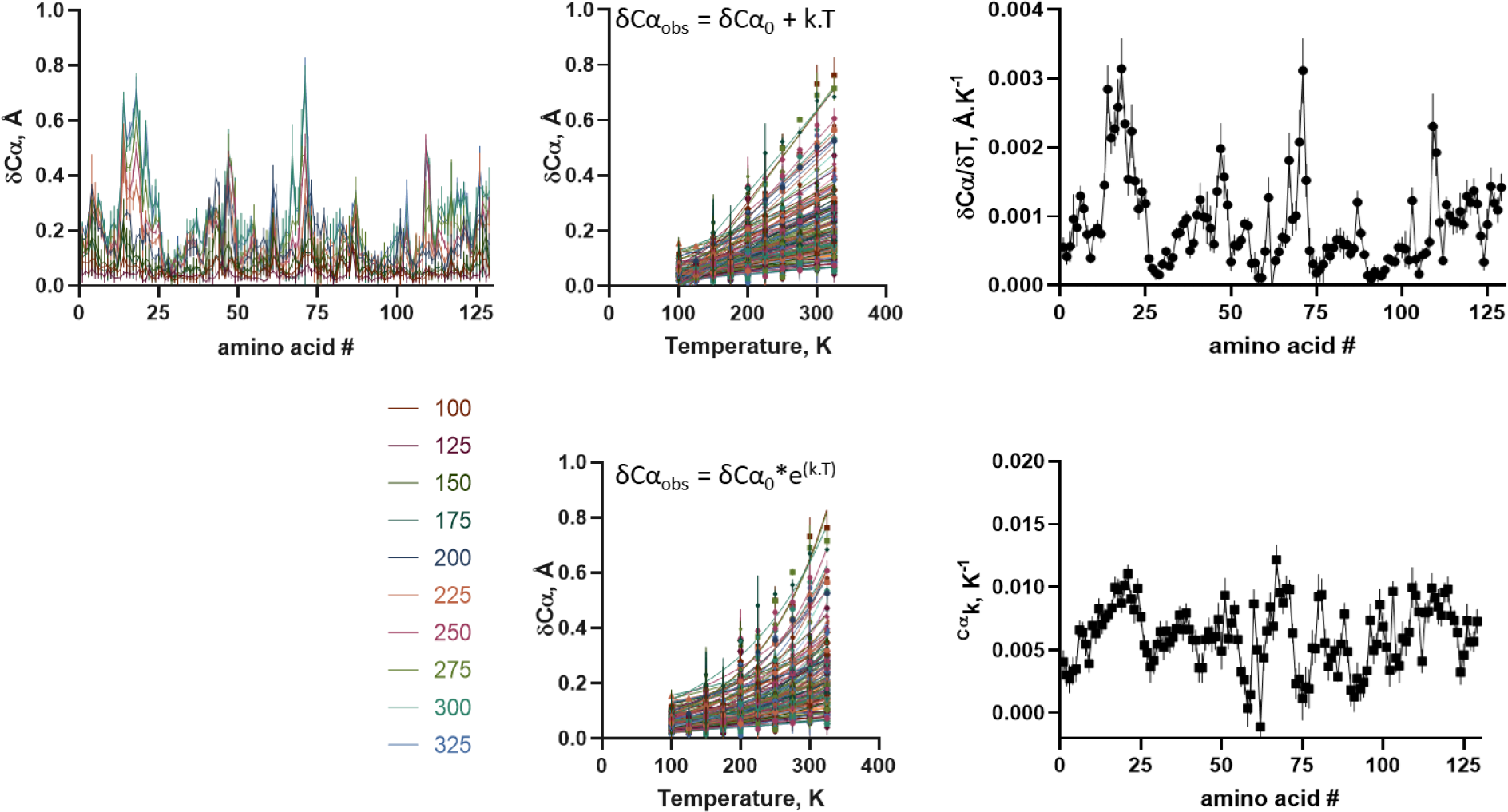
Temperature induced conformational changes in lysozyme. Structures were aligned with a reference structure at 100 K using *Superpose* (CCP4) and distances between Cα were obtained. A) Changes in Cα distances along protein sequence at varying temperatures (different colors). B) Changes in Cα distances as a function of temperature for each amino acid residue (different colors) protein sequence at varying temperatures. Lines are first order linear regression, from which were obtained their thermal-dependence of conformational change (δCα; angular coefficient of linear regression). C) Distribution of thermal-dependent conformational change (δCα) along protein sequence. D) Changes in Cα distances as a function of temperature for each amino acid residue (different colors) protein sequence at varying temperatures. Lines are exponential non-linear regression, from which were obtained their exponential thermal constant (^δCα^k). E) Distribution of exponential thermal constant (^δCα^k) of conformational change along protein sequence. Symbol is average and bar is standard deviation (n=3).

### Temperature affects average B-factor uniformly

We further analyzed the changes in B-factor as a function of temperature. **The average** B-factor from data as estimated from the Wilson plot (**Fig. 3A**), and from the final refined structure side-chains (**Fig. 3B**), Cα (**Fig. 3C**) or N-amide (**Fig. 3D**) showed similar exponential positive correlation with temperature, increasing about 2.5 times from 100 K to 325 K. The thermal dependence of average observed B-factors (^avg^B) was adjusted with a single exponential function according to **[Eq. 3]**

**Figure 3.**
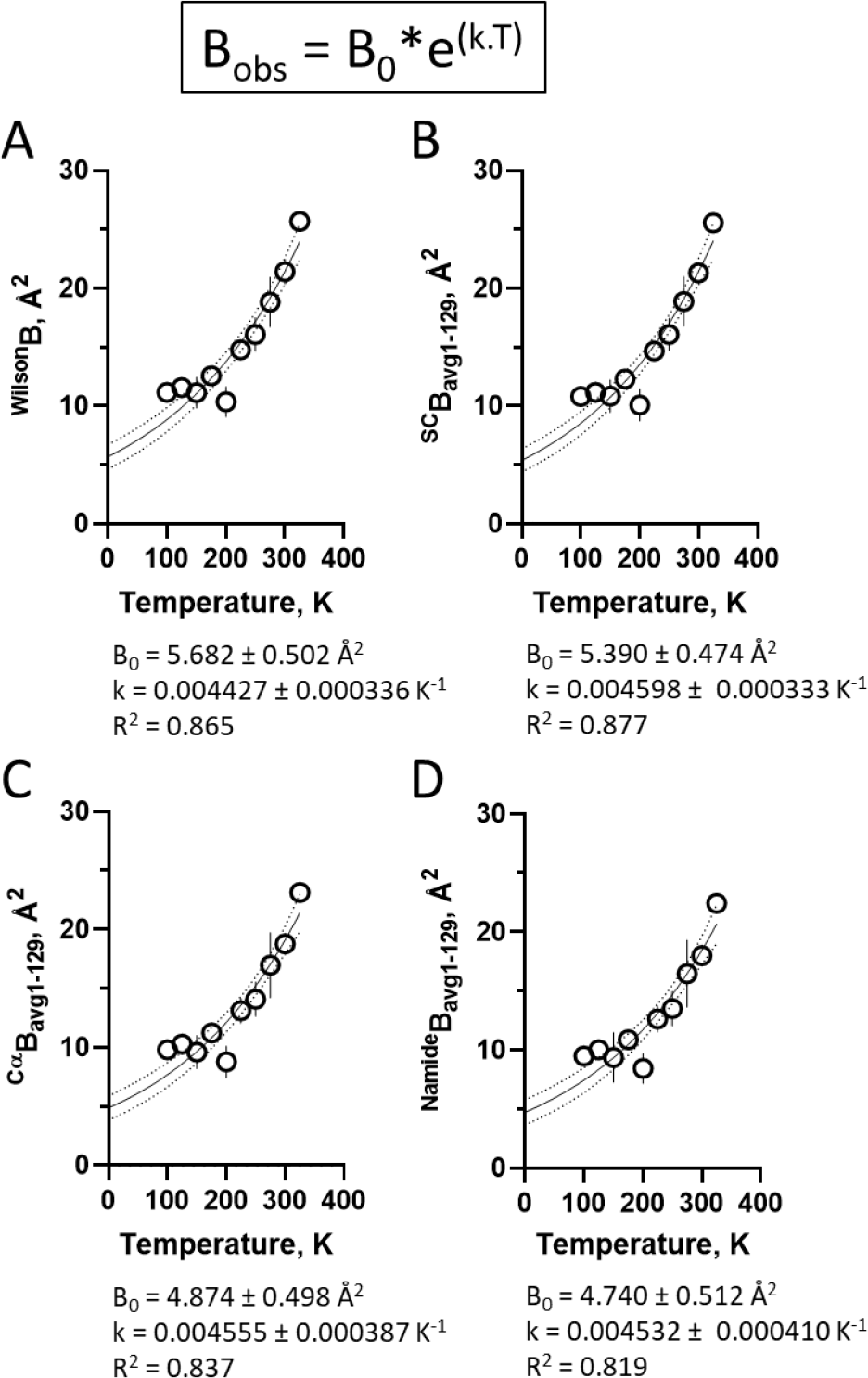
Temperature-dependent changes in B-factor (B). Overall changes in B-factor as a function of temperature for **A)** Wilson plot, **B)** side-chains, **C)** Cα and **D)** amide nitrogen. Values corresponds to the extrapolated B factor at zero Kelvin (B_0_) and the thermal coefficient k, as inferred from single exponential (equation above panels) non-linear regression of their respective panels (solid lines; dotted lines are 95% confidence interval). Symbol is average and bar is standard deviation (n=3).

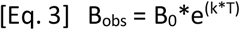

where **B**_**obs**_ is the refined B-factor, **B**_**0**_ is the B-factor interpolated to zero Kelvin from regression, **k** is the thermal constant, and T is the data collection temperature (in Kelvin), resulting in close average thermal factors of about ^avg^k = 0.0045 K^-1^ and in ^avg^B_0_ of about 5 Å^2^

### Temperature B-factor is a global constant

A detailed picture of B-factor dependence on data collection temperature could be inferred from the B-factor **distribution over the polypeptide chain** at varying temperatures for side-chains (**Fig. 4A; Fig. S10**), Cα (**Fig. 5A; Fig. S11**) and N-amide (**Fig. 6A; Fig. S12**). These data revealed similar B-factor distribution pattern along the polypeptide chain for all temperature tested.

**Figure 4.**
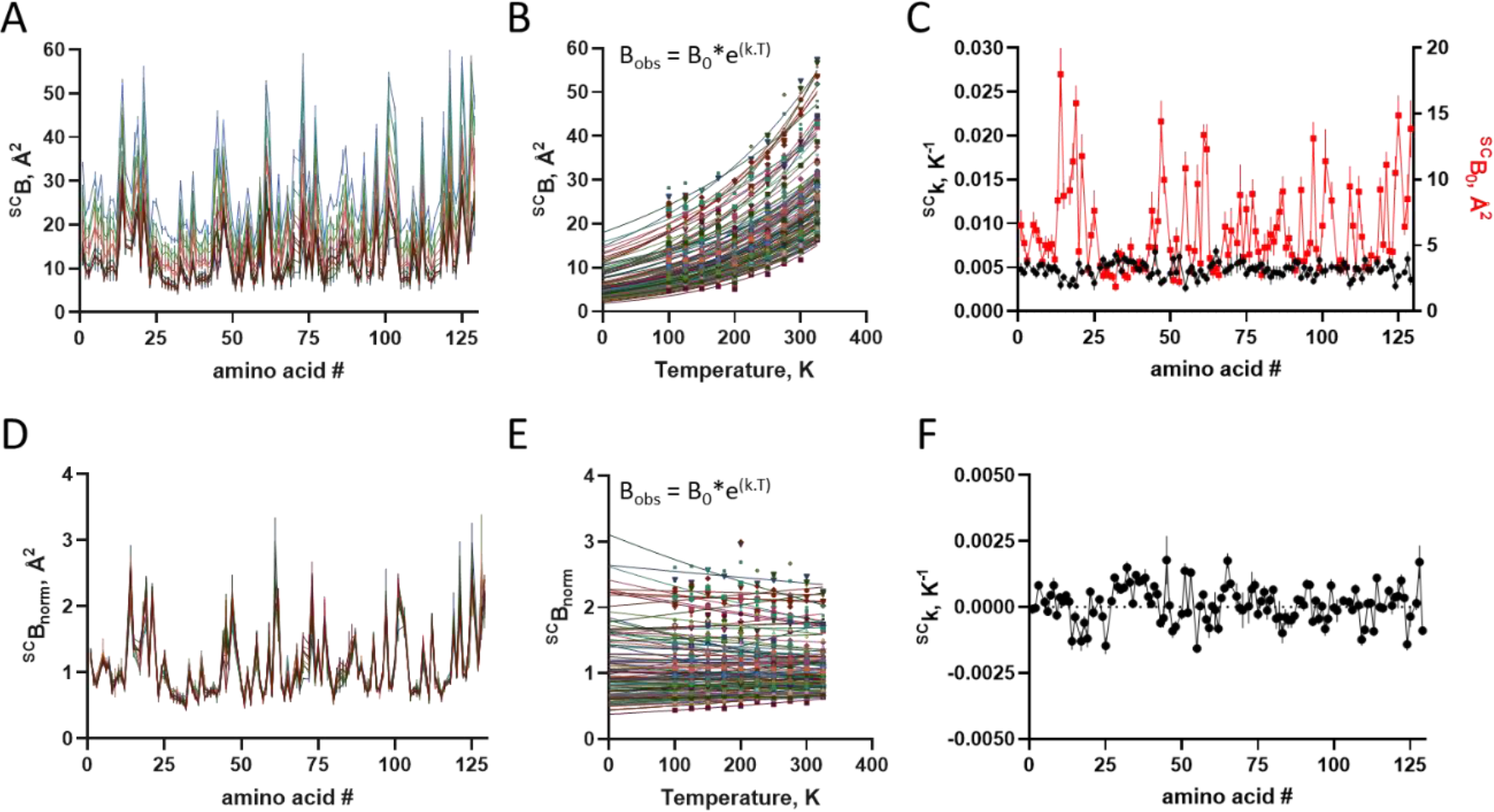
Temperature-dependent changes in side-chain B-factor. A) B-factor distribution for side chain at varying temperatures (different colors). B) Changes in raw B-factor as a function of temperature for each amino acid side chain (different colors). Lines are exponential non-linear regression, from which were obtained the raw B_0_ and the thermal constant k. C) Distribution of raw B_0_ and raw k along protein sequence. D) Normalized B-factor (B_norm_) distribution for side chain at varying temperatures (different colors). E) Changes in B_norm_ as a function of temperature for each amino acid side chain (different colors). Lines are exponential non-linear regression, from which was obtained the thermal constant ^norm^k. C) Distribution of ^norm^k along protein sequence. Symbol is average and bar is standard deviation (n=3).

**Figure 5.**
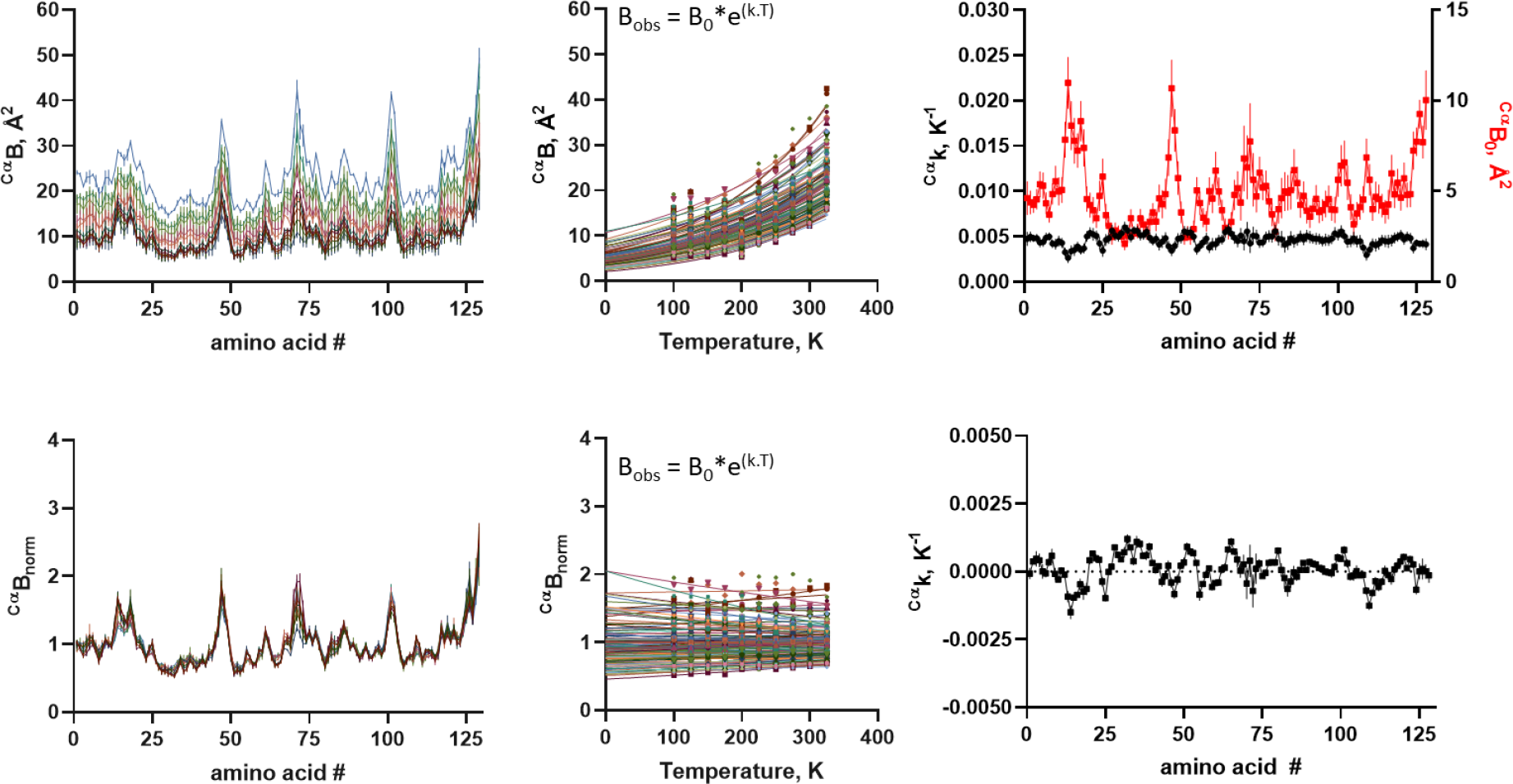
Temperature-dependent changes in Cα B-factor. A) B-factor distribution for Cα at varying temperatures (different colors). B) Changes in raw B-factor as a function of temperature for each amino acid Cα (different colors). Lines are exponential non-linear regression, from which were obtained the raw B_0_ and the thermal constant k. C) Distribution of raw B_0_ and raw k along protein sequence. D) Normalized B-factor (B_norm_) distribution for Cα at varying temperatures (different colors). E) Changes in B_norm_ as a function of temperature for each amino acid Cα (different colors). Lines are exponential non-linear regression, from which was obtained the thermal constant ^norm^k. F) Distribution of ^norm^k along protein sequence. Symbol is average and bar is standard deviation (n=3).

**Figure 6.**
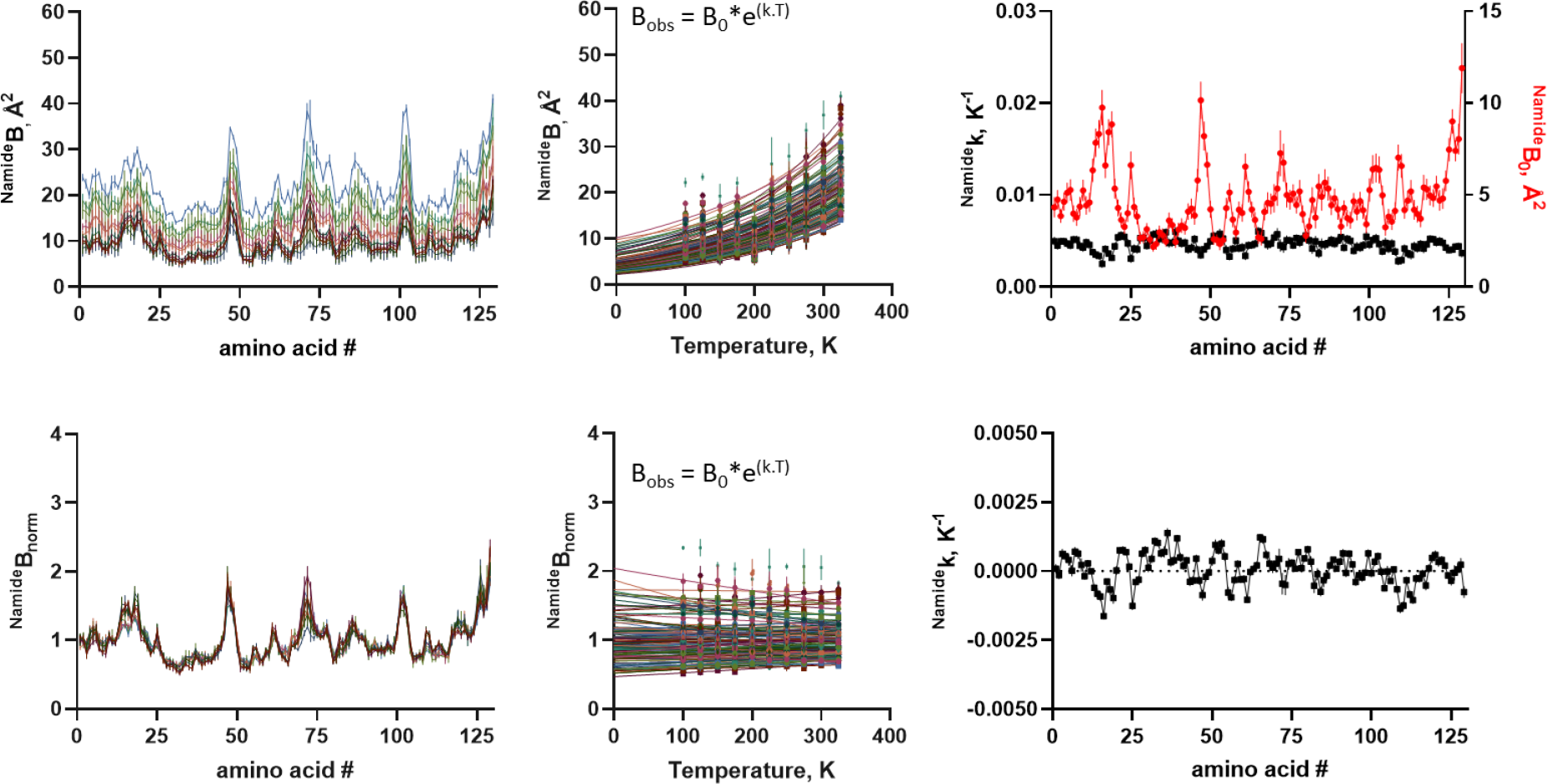
Temperature-dependent changes in amide nitrogen (N-amide) B-factor. A) B-factor distribution for N-amide α at varying temperatures (different colors). B) Changes in raw B-factor as a function of temperature for each amino acid Cα (different colors). Lines are exponential non-linear regression, from which were obtained the raw B_0_ and the thermal constant k. C) Distribution of raw B_0_ and raw k along protein sequence. D) Normalized B-factor (B_norm_) distribution for N-amide at varying temperatures (different colors). E) Changes in B_norm_ as a function of temperature for each amino acid N-amide (different colors). Lines are exponential non-linear regression, from which was obtained the thermal constant ^norm^k. F) Distribution of ^norm^k along protein sequence. Symbol is average and bar is standard deviation (n=3).

We further analyzed the changes in B-factor as a function of temperature for each amino acid residue, for their side-chain (**Fig. 4B**), Cα (**Fig. 5B**) and N-amide (**Fig. 6B**), and adjusted these temperature curves by a single exponential non-linear regression according to [Eq. 3], which described adequately the thermal-dependent changes in ^Wilson^B and average ^SC^B, ^Cα^B and ^N-amide^B (**Fig. 8**). The distributions of the calculated B_0_ along the polypeptide chains (**Fig. 4C, Fig. 5C** and **Fig. 6C**) showed similar patterns for the raw B-factor distributions (**Fig. 4A, Fig. 5A** and **Fig. 6A**). In opposite, the estimated thermal factor ^CαB^k was found to be roughly constant at about 0.005 K^-1^ along the polypeptide chain for side-chains (**Fig. 4C**), Cα (**Fig. 5C**) and N-amide (**Fig. 6C**), in close agreement with those found for ^avg^k (**Fig. 3**). These data suggests that the B-factor increases uniformly for the whole system (side chain, Cα and N-amide), and thus confirming it as a **temperature-dependent factor**. However, the thermal factor ^**B**^**k** shows no association with the observed B-factor (**Fig. 4C, Fig. 5C, Fig. 6C**), indicating that the absolute B-factor does not predict its temperature dependence. Moreover, the similarity of the thermal factor ^**B**^**k** between all amino acid residues (both side-chains, Cα and N-amide) indicates that this parameter may not correlate with the observed temperature-induced protein conformational changes (**Fig. 2**).

### Local temperature dependence of B-factor is null

Since the temperature-dependence of the raw B-factor (**B**_**obs**_) was found to be similar for all protein atoms (**Fig. 4-6**), as well as for the Wilson B-factor (**Fig. 8**), we hypothesize that a uniform thermal contribution may additively influence one component of the mean-square atomic displacement, resulting in the observed raw B-factor.

Converting [Eq. 3] to a log function results in

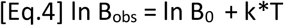

Incorporating the contribution of a thermal component B_other_ to the raw B_obs_ becomes [Eq. 5] ln B_norm_ = ln

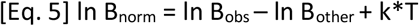

which rearranges back to

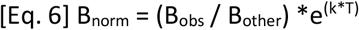

Since the thermal contribution is indeed uniform among all atoms, we may estimate B_other_ as the average of the B_obs_, i.e., B_avg_, resulting in

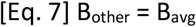

and thus

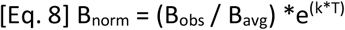

We normalized the **B**_**obs**_ into **B**_**norm**_ by scaling it down with the average B-factor (**B**_**avg**_) according to [**Eq. 2**]. The **B**_**norm**_ distribution profile becomes stable over a broad temperature range (100 K to 325 K), for side-chain (**Fig. 4D**), Cα (**Fig. 5D**) and N-amide (**Fig. 6D**), which were similar for all temperatures. The **B**_**norm**_ was roughly constant as a function of temperature for each amino acid side-chain (**Fig. 4E**), Cα (**Fig. 5E**) and N-amide (**Fig. 6E**). We adjusted each temperature curves for each amino acid residue side-chain (**Fig. 4E**), Cα (**Fig. 5E**) and N-amide (**Fig. 6E**) with single exponential function [**Eq. 3**]. The estimated thermal factor ^**Bnorm**^**k** was found to fluctuate around zero, for side-chain (**Fig. 4F**), Cα (**Fig. 5F**) and N-amide (**Fig. 6F**). These data suggest that the scaled **B**_**norm**_ embodies only the static, temperature-independent component of the B-factor.

We further explored the hypothesis for a link between conformational changes and B-factor, as depicted in **Fig. 7**. While ^avg^B and temperature were found to be tightly correlated (**Fig. 3**; R^2^>0.8), only a loose correlation was found between thermal expansion coefficient ^δ**Cα/**δ**T**^**k** and **B**_**0**_ (R^2^=0.371; (**Fig. 7A** and **Fig. 7C**), or between ^**CαB**^**k** (R^2^=0.175; **Fig. 7B** and **Fig. 7D**). While a progressive shift in conformation occurs for side-chain (**Fig. S10**), Cα (**Fig. S11**) and N-amide (**Fig. S12**), no major changes in distribution for **B**_**norm**_ seems to occur as a function of temperature.

**Figure 7.**
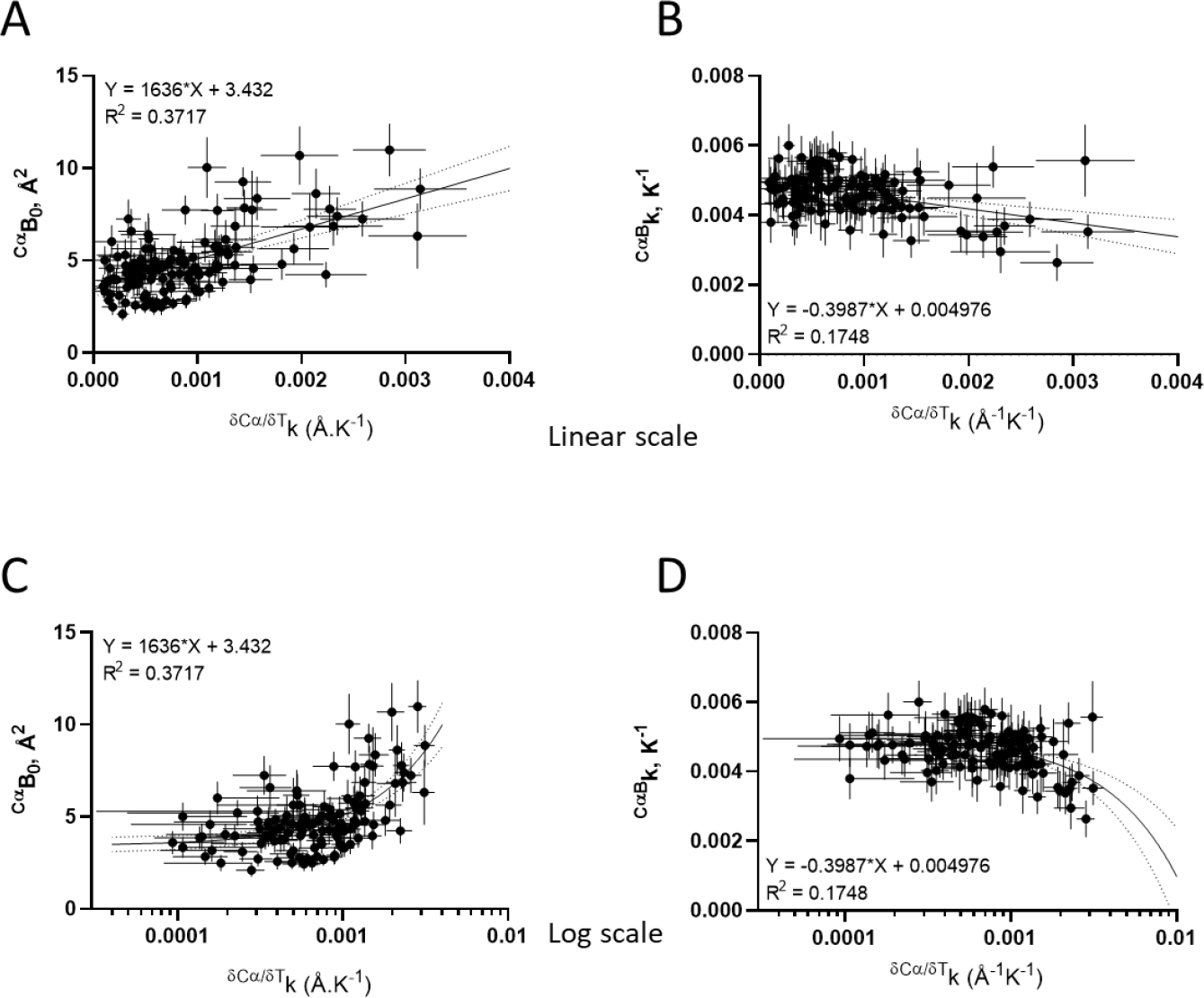
Correlation between changes in conformation B-factor parameters. Correlation between B-factor parameters (B_0;_ panels **A** and **C**) and (^CαB^k) and temperature-induced conformational changes constant (^δCα/δT^k; panels **B** and **D**). Data correspond to calculated ^δCα^B_0_ from **Fig. 5C** and ^δCα/δT^k from **Fig. 2C**. Data are plotted in linear (**A** and **B**) or log scale (**C** and D) for ease of observation of data span. Lines correspond to first-order linear regression (continuous) and 95 % confidence interval (dotted).

In order to test the validity to other systems, we analyzed the RCSB for datasets available from single-crystal diffraction under similar controlled conditions and temperature as variable, as inspected from their respective published articles. We have found data from thaumatin crystal structures solved at temperatures from 25 K to 300K. These data showed increasing B-factor as a function of temperature (**Fig. 8A**), which revealed similar pattern for B_norm_ distribution along the amino acid sequence (**Fig. 8B**). Thaumatin change conformation as a function of temperature (**Fig. 8C**), although the expansion coefficient ^δ**Cα/**δ**T**^**k** does not correlate with either the absolute B-factor (**Fig. 8D**) or the thermal factor ^**CαB**^**k**.

**Figure 8.**
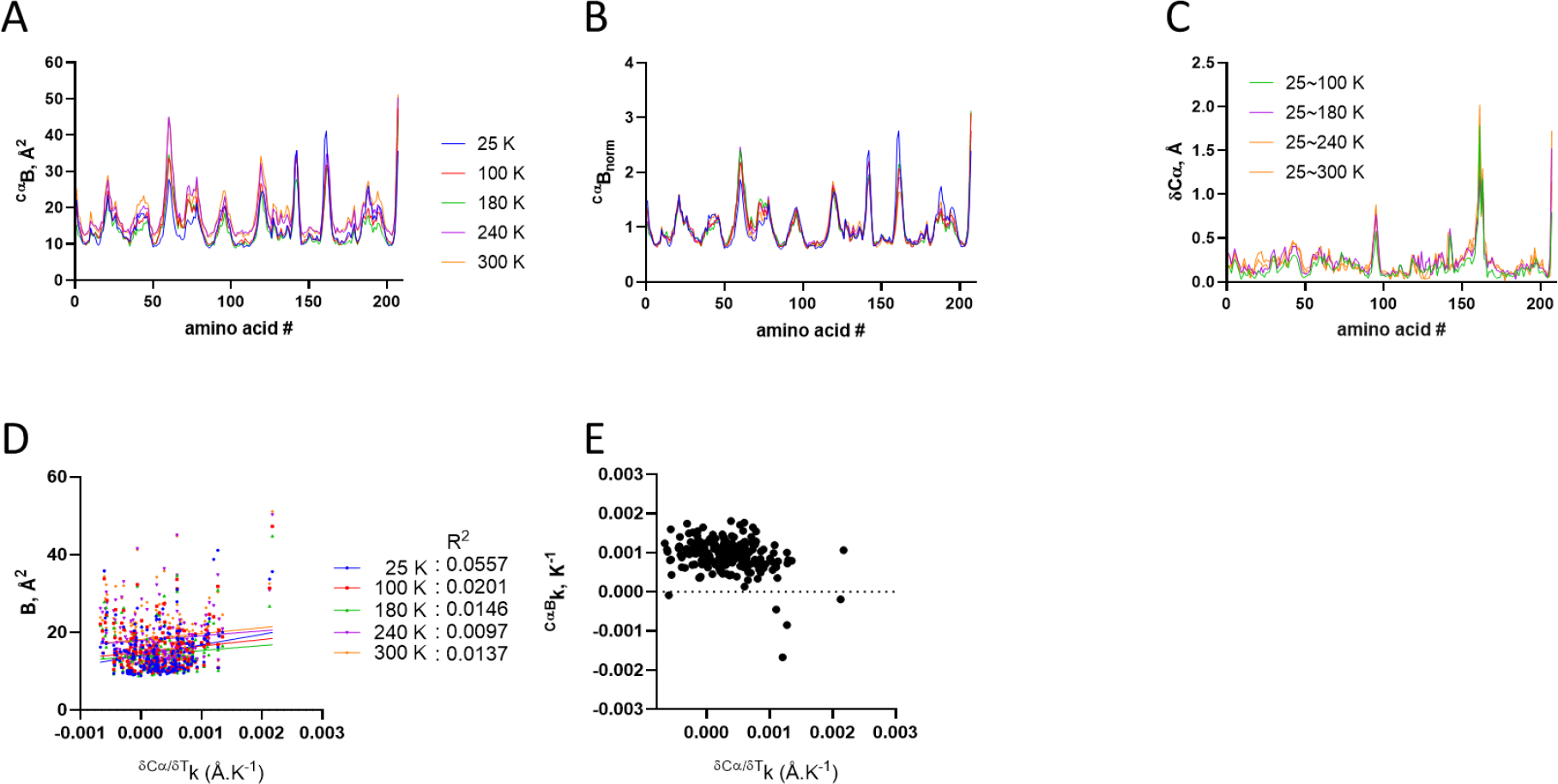
Temperature-dependence of B-factor and conformation of Thaumatin. X-ray structures solved at 25 K (PDB ID 4ek0), 100 K (PDPB ID 4ekb), 180 K (PDB ID 4eko), 240 K (PDB ID 4el2) and 300 K (PDB ID 4el7), as depicted in the Figure legends. A) B-factor distribution for Cα at varying temperatures (different colors). B) Normalized B-factor (B_norm_) distribution for Cα at varying temperatures (different colors). C) Changes in Cα distances along protein sequence at varying temperatures (different colors). D) Correlation between B-factor and backbone thermal expansion coefficient (^δCα/ δT^k) at varying temperatures (different colors). Each point corresponds to one amino acid. E) Dependence of thermal factor (^CαB^k) on the thermal expansion coefficient (^δCα/ δT^k). Data from [10].

## Discussion

Inferring conformational changes from a given protein require testing chemical (pH, ligands, drugs, metals, cofactors, cosolutes, point mutation, hydration, among others) and/or physical (temperature, pressure) variables which might induce dynamic conformational changes and accumulation of polymorphs. In the case of crystal structures, it may even result in dissimilar packing and space groups.

Previous work has shown that tetragonal lysozyme present changes in B-factor as a function of temperature in a non-uniform patter along the protein, with smaller changes at regions involved in symmetry-related intermolecular contacts [32,33]. Although the lysozyme (44 mg/mL) was crystallized in similar conditions reported here, at same space group (P4_3_2_1_2) from 0.7 M NaCl in sodium acetate pH 4.8, main-chain distances over 20 Å were found between structures solved at 298 and 100 K. However, while data collected at 100 K was obtained from crystals mounted in a glass fiber, the data obtained at 298 K was collected from crystal mounted into a capillary tube, making it difficult to infer the variable dictating crystallographic changes. Ribonuclease-A type III (RNAse) has been also studied at 1.5 Å resolution, over a broad range of temperature, from 98 K to 320 K [29,46]. However, each crystal was collected under dissimilar conditions of solvent composition (50 % MPD, for 98 K to 220 K, or 50 % methanol, for 240 K to 320 K), crystal mounting technique (capillary or glass fiber), diffractometer, instrument controllers and data collection procedures [29], limiting interpretation concerning the effective variable.

The reproducibility of the crystallographic model of lysozyme at high resolution (1.6 Å) has been shown to be robust when using tetragonal lysozyme obtained in sodium acetate pH 4.6 and NaCl by hanging drop vapor diffusion technique and using two dissimilar home-source diffractometers [15]. While the final structural models showed reproducible, the raw B-factor was found non-reproducible despite their similar distribution along the polypeptide chain. The normalization of the B-factor by the average B-factor allowed obtaining reproducible B-factor patterns, suggesting that the B-factor could be influenced by variables from crystal handling to data collection. It thus became evident the need for stabilization of crystal aiming a more stable and reproducible data collection.

In the present work, we used same cryoprotectant (glycerol), mounting device (nylon loop) and stabilization (synthetic hydrocarbon grease, Apiezon®) for all dataset, allowing high repeatability of B-factors at each temperature compared to equivalent data collected in the absence of grease [15]. We have reported here a broad data collection temperature investigation of tetragonal lysozyme crystals allowing the observation of dissociation between lysozyme conformational changes and B-factor. These data were possible to obtain thanks to the strategy of crystal stabilization with a hydrocarbon grease, allowing x-ray diffraction at high resolution in a broad temperature range, from 100 K up to 325 K, without loss in lattice order and diffraction power. In fact, since we obtained high resolution (1.5 Å) through the investigated temperature range, one can assume that the B-factor is not dominated by the well-organized lattice [21,29].

As previously envisioned by Wilson [47], the B-factor is susceptible to temperature, and indeed we have confidently observed this phenomena here once solved the experimental reproducibility of the B-factor by using hydrophobic grease. However, when the B_obs_ was normalized to the B_norm_ by scaling down with the B_avg_ (from all atoms of the polypeptide chain) we observed similar B-factor distribution pattern over the polypeptide chain for the temperature range assayed here (100 K to 325 K), both for Cα (^Cα^B), (^N-amide^B) and side chain (^SC^B). These data indicate that the inherent protein B_norm_ is not temperature-dependent, and that the total B-factor (including the B_Wilson_) is most likely dominated by dynamic, rigid body contributions [26,29]. The variations in the inherent protein B-factor along the polypeptide chain is likely to be influenced by the atom environment, by immediate local interactions, and by adjacent amino acids as previously reported [17]. While local conformational changes take place in lysozyme as a function of temperature, it did not correlate with B-factor parameters (**Fig. 7**) given the distribution profile and correlation coefficient [48], suggesting a dissociation between these two parameters, also observed for the unrelated protein model thaumatin (**Fig. 8**) [10].

We have shown the extension of our findings with lysozyme to thaumatin, and even being conducted with a rigorous triplicate for each temperature and a standard data collection and processing workflow, we used only one protein (lysozyme) at one crystallization condition and space group, requiring future independent studies to corroborate or not the present conclusion. The use of the grease during data collection may accelerate this field, providing further assistance in the thermal protection against dehydration.

We envision that the raw B_obs_ is composed by the mostly thermal-insensitive atomic vibration B_A_ and an additive contribution of a thermal-sensitive rigid body motion B_RB_ along with other contributions to baseline B-factor rising from sources other including crystal handling and data collection, which could be minimized by using a sealant such as hydrocarbon grease. The present study with our original data from thermal-induced changes in lysozyme (and thaumatin) demonstrate that changes in B-factor does not correlate with conformational changes. While the raw B-factor may indeed correlate with temperature, the normalized B-factor does not seem to be susceptible by this variable, lacking local changes that could be related to concomitant conformational transitions in the protein.

## Conclusions

We have found that embedding protein crystals within hydrocarbon grease allowed stable and repeatable x-ray diffraction data collection and structural solution in a broad temperature range. We observed a lack of correlation between local protein B-factor and temperature-induced conformational changes, demonstrating a dissociation of the mean-squared deviation with conformational plasticity. More importantly, since the raw B-factor is uniformly influenced by temperature at same constant, it suggests a global, dynamic correlation with rigid body motion and/or any other scattering and crystal property. Further development should consider separation the B-factor components – atomic and system (including rigid body, scattering, among others) - contributing for the composite raw B-factor.

## Materials and Methods

### Material

HEWL was obtained from Fluka (Cat #62971; Lot #BCG4805V) and kept at -20°C until use. All other reagents were of analytical grade.

## Methods

### Protein crystallography

Crystals of HEWL were obtained by vapor diffusion, sitting-drop technique, using Corning 3552 plates with 1 µL HEWL 50 mg/mL in water, and 1 µL precipitant, equilibrated against 80 µL precipitant solution in the well, comprising 1.2 M NaCl, 100 mM sodium acetate pH 4.6 (adjusted either with acetic acid or NaOH), at 22 °C ± 2 °C. This method was used throughout the study in order to avoid the large variability inherent to the vapor-diffusion crystallization step [49]. Diffraction-quality crystals grew within 1 day and were used for diffraction after 2 days. Crystals were handled with 20 µm nylon CryoLoop™ (Hampton Research). Each crystal was soaked for 5 to 10 sec in the same crystallization solution from the well, supplemented with glycerol (for 30% v/v final concentration, with proportional dilution of the well solution), following by immersion into Apiezon® N hydrocarbon grease, and mounted onto the goniometer under N_2g_ stream at desired temperature for data collection. Each dataset corresponds to a single crystal at a given temperature.

Since Apiezon® N has a recommended working range of 4 to 303 K (−269 °C to 30 °C), we also tested the Apiezon® T which has recommended use range of 283 to 393 K (10 °C to 120 °C), although it provided no additional benefits, since lysozyme lost diffraction power in temperatures over 55 °C (not shown).

### Diffractometer setup and data acquisition

The crystals were subjected to X-ray diffraction and data collection using CuKα radiation at constant exposure time using a 30 W air-cooled |µS microfocus source (Incoatec) mounted on D8-Venture diffractometer (Bruker; at a CENABIO-UFRJ facility) operating at 50 kV and 1.1 mA, and recorded on a Photon II detector (Bruker). Crystals were under N_2g_ stream at desired temperature with flow of 1.2 L/h, controlled by a CryoStream 800 (Oxford Cryogenics). All datasets were collected using 60 sec exposures with 0.5° oscillation/image and at leas 99% completeness assuming identical Friedel pairs, targeting 1.5 Å resolution. Data were collected, indexed, integrated and scaled using *Proteum3* (Bruker AXS Inc.), followed by analysis with *Truncate* [50].

A nonredundant set of information regarding the tetragonal (P4_3_2_1_2) hen lysozyme crystal structures determined by X-ray diffraction of single crystals was obtained from The Research Collaboratory for Structural Bioinformatics Protein Data Bank (RCSB PDB, rcsb.org), as of February 15, 2022 (**Supporting Information**). After filtering for entries with missing information, data was plotted according to the desired analysis (**Fig. S1**).

### Data processing

Data was further processed and analyzed using *C*.*C*.*P*.*4*. v7.0.071 [51]. The crystal structures were solved by isomorphous replacement with 20 cycles of rigid body search with RefMac v5.8.0238 [52] [53] using PDB entry 3A8Z [54] edited for single occupancy and no water as a search model, allowing only a single sodium ion, resulting in a clear solution for a monomer in the asymmetric unit. The solution was further subjected to 10 cycles of restrained refinement using *Refmac* [52]. Real space refinement was conducted by visual inspection of both the map and the model with *C*.*O*.*O*.*T*. v0.8.9.2 [55], adjusting solely misplaced side chains, followed by water addition using 1.2 σ as threshold. Additional restrained refinement with 10 cycles was performed using *Refmac*. All data processing was performed using default mode in order to avoid bias. Further data analysis was conducted with *Superpose* v1.05 from *C*.*C*.*P*.*4*. Global pairwise alignment of the HEWL structures was performed using ProSMART [56]. Analysis of B-factor was performed with *Baverage* (CCP4).

A summary of crystal parameters along with data collection and refinement statistics are presented in **Supporting Information**, in **Table S1** and as a function of the data collection temperature (**Fig. S13**). Structural validation of the model performed with PROCHECK. All structure figures were generated with *PyMOL* v2.0 [57]. Graphics were generated with GraphPad Prism v. 8.0.2 for Windows (GraphPad Software, San Diego, California USA, www.graphpad.com). The atomic coordinates have been deposited with the Protein Data Bank and have been assigned the codes depicted in **Supporting information**, along with the RCSB validation report.

### Circular Dichroism

Thermal denaturation of HEWL (20 mg/mL) was performed in 50 mM AcONa pH 4.6, in a circular dichroism spectropolarimeter (JASCO® J-810; Tokyo, Japan; LMProt-ICB-UFMG, Brazil), equipped with Peltier Jasco® temperature control system - PFD-425S (Tokyo, Japan), using a 100 µm pathlength quartz cuvette (Uvonic Instruments, Plainview, NY). All the spectra were recorded with three scans from 260 to 190 nm using a 0.2 nm spectral bandwidth, 100 nm/min scan speed, and 1 sec response time. Similar experiments with the respective blank solutions were also carried out in order to allow for proper background subtraction. The temperature was increased at a rate of 5 °C/min, and allowed equilibration for 60 sec before measurements. Temperature curve was analyzed using a sigmoidal 4 parameters function.

### Direct Infusion ESI-Traveling Wave – Ion Mobility Spectrometry – Mass Spectrometry (ESI-TWIM-MS)

The lysozyme was diluted to 50 µM in 0.1 % formic acid. The ESI-TWIM-MS measurements were performed in a Traveling Wave Ion Mobility Mass Spectrometer (TWIM-MS, Synapt G1 HDMS, Waters, UK), with perfusion at 20 µL/min in positive ESI with a capillary voltage of 4.0 kV and N_2_g used as mobility gas at 0.4 bar, and source temperature varying between 50 °C and 250 °C in 50 °C intervals. The data acquisition was conducted over the range of *m/z* 500 to 3,000 for 5 min. The mass spectrometer calibration was performed with phosphoric acid 0.1 % (v/v) in acetonitrile:H_2_O (1:1). Other typical instrumental settings are as described previously [58]. Data were analyzed using DriftScope [59] (Waters Corporation, UK). All measurements were performed at the *Laboratory of Macromolecules* – LAMAC / *Life Science Metrology Division* – DIMAV, INMETRO, Rio de Janeiro, Brazil.

### Structural analysis from RCSB-PDB

In order to further extend our temperature analysis, we searched the RCSB for crystallographic protein structures solved by x-ray diffraction at varying temperatures (**<400 K**) and resolution ≤ 2.0 Å, deposited between 01.01.2000 and 02.28.2022. Further search parameters included average B-factor and Wilson B-factor estimate ≤ 100 Å^2^ and refinement R-free ≤ 0.56. This search returned 24,755 entries. These entries were searched for works showing works reporting crystal structures solved with at least **five** equivalent entries (such as same protein, pH, crystallization condition, space group) under data collection temperatures spanning most regularly for at least 200 K.

The RCSB was also searched for hen egg white **lysozyme** crystal structures solved by x-ray diffraction in space group **P4**_**3**_**2**_**1**_**2** up to 02.15.2022. This search returned 516 entries which were analyzed for refinement and structural parameters as reported in **Supporting Information**.

### Data analysis

Graphics were generated with GraphPad Prism v. 8.0.2 for Windows (GraphPad Software, San Diego, California USA, www.graphpad.com). Significance was considered when p<0.005 [60].

## Supporting information

full crystallographic report

## Abbreviations

HEWL: hen egg white lysozyme
CD: circular dichroism
ESI: Electrospray ionization
MS: mass spectrometry
IMS: ion mobility spectrometry.

## Acknowledgments

We would like to thanks Prof. Mariana Quezado (UFMG) for valuable discussion along the development of this work. We thank Dr. Sandra Scapin (INMETRO) for helpful assistance during ESI-IMS-MS data collection.

## Author Contributions

**Fernando de Sá Ribeiro** - Data curation; Formal analysis; Investigation; Methodology; Validation; Visualization; Writing - review & editing.

**Luís Maurício T. R. Lima -** Conceptualization; Data curation; Formal analysis; Funding acquisition; Investigation; Methodology; Project administration; Supervision; Validation; Visualization; Roles/Writing - original draft; Writing - review & editing.

