## Supporting Information for "Linking B-factor and temperature-induced conformational transition"

**Running title:** Thermal-dependence of B-factor

*To whom correspondence should be addressed
**Authors Address and Contact**
- Luis Mauricio T. R. Lima – Pharmaceutical Biotechnology Laboratory (pbiotech), Faculty of Pharmacy, Federal University of Rio de Janeiro – UFRJ, CCS, Bss24, Ilha do Fundão, 21941-590, Rio de Janeiro, RJ, Brazil. Phone/Fax: (+55-21) 3938-6639 –. Social Media: @pbiotech

**AUTHOR LIST**
Fernando de Sá Ribeiro – – 0000-0002-8521-6745

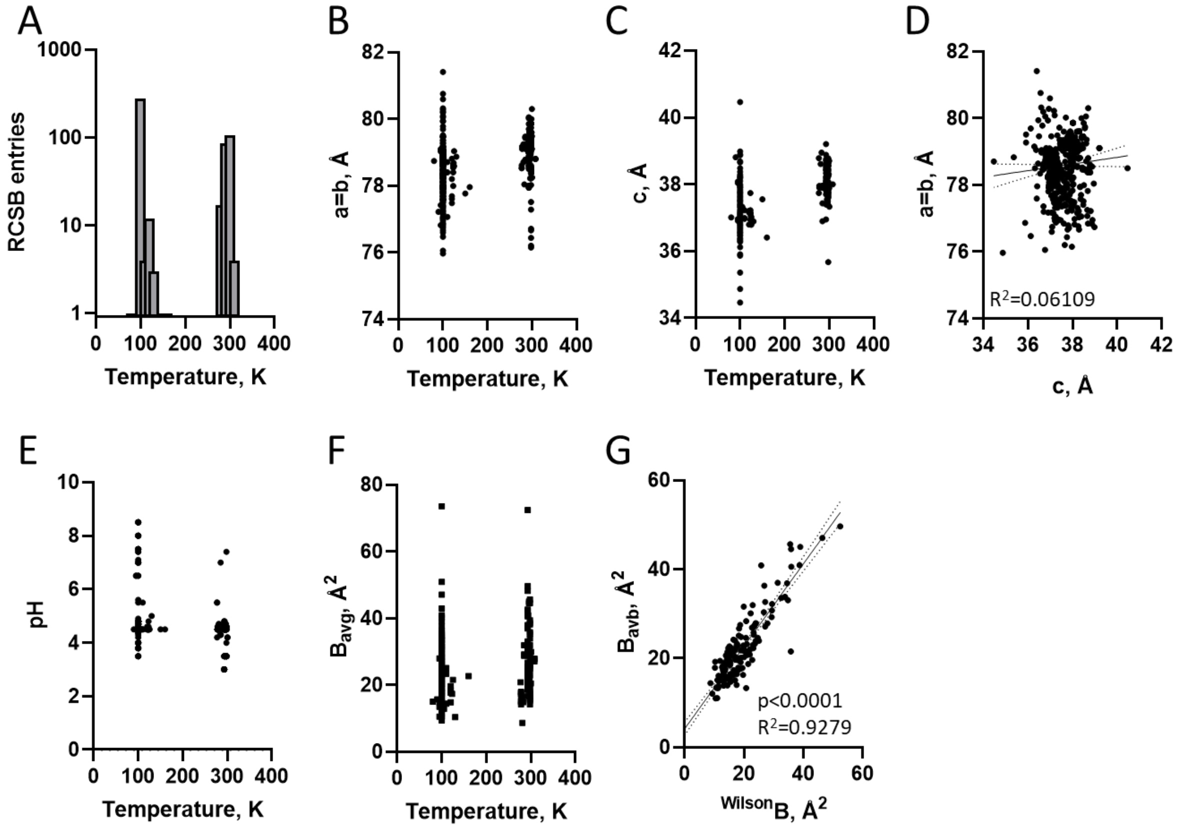


**Figure S1. Tetragonal lysozyme in the RCSB**
Data from the RCSB (access Feb 15, 2022) for lysozyme in $P4_32_12$.
**A)** distribution of entries according to data collection temperature, found between 80 k to 308 K.
**B)** Unit cell length in the a=b axis as a function of temperature.
**C)** Unit cell length in the c axis as a function of temperature.
Correlation between data collection temperature (from 80 k to 308 l) and **(D)** crystallographic unit cell parameters (516 entries), **(E)** crystallization pH (381 entries), and **(F)** average B-factor from final model (391 entries).
**G)** Correlation between B-factors from Wilson estimate and average from final structure model (171 entries), according to $B_{avg}$= = 0.9279*$^{Wilson}$B + 4.149. Continuous line are first order linear regression and dotted lines are 95 % confidence interval.

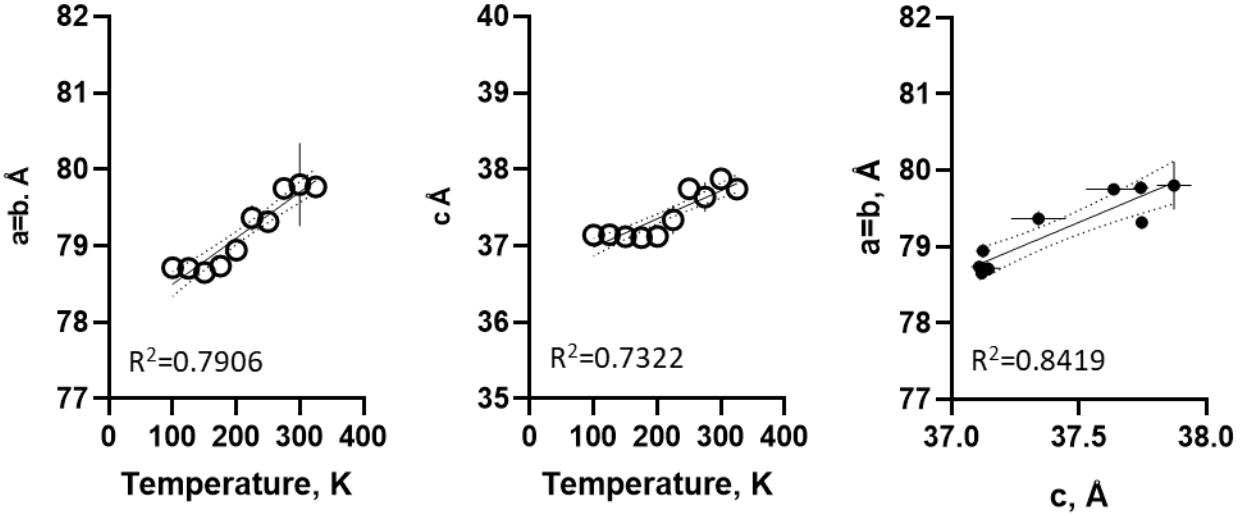


**Figure S2. Thermal-dependent changes in unit cell parameters.**
Correlation between cell unit parameters and data collection temperature for single crystal x-ray diffraction of lysozyme in $P4_32_12$. Continuous line are first order linear regression and dotted lines are 95 % confidence interval. The linear correlation between data collection temperature and changes in unit cell dimensions were found significant for a=b ($p<0.0001$) with 0.006019 Å.K$^{-1}$, and c ($p<0.0001$) with 0.003702 Å.K$^{-1}$. Symbol is average and bar is standard deviation (n=3).

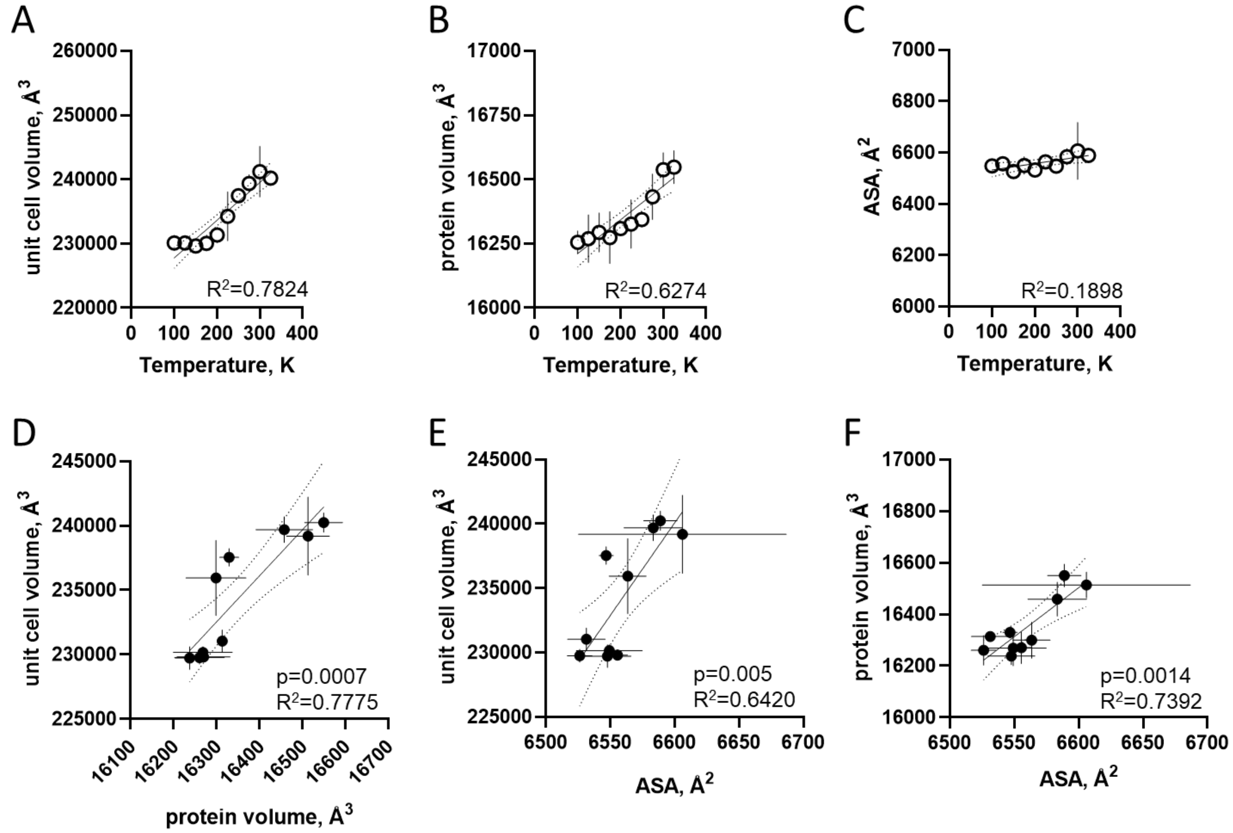


**Figure S3. Thermal-dependent changes in volumetric properties.**

Continuous line are first order linear regression and dotted lines are 95 % confidence interval. A linear dependence on data collection temperature was found significant ($p<0.0001$) for unit cell volume (58.72 $Å^3.K^{-1}$), protein volume ($p<0.0001$; 1.315 $Å^2.K^{-1}$), and non-significant for ASA ($p=0.0161$; 0.2505 $Å.K^{-1}$). Linear correlations between temperature-induced changes in unit cell volume, ASA and protein volume were found significant ($p \leq 0.005$) as indicated in their respective panels. Symbol is average and bar is standard deviation (n=3).

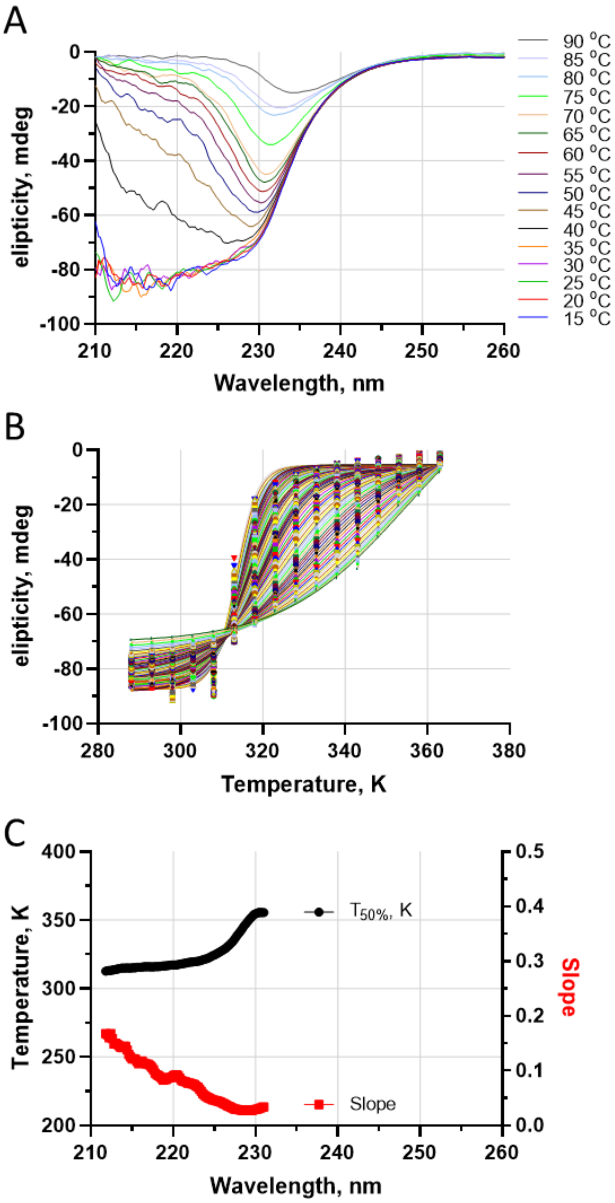


**Figure S4. Thermal-dependent conformational changes in lysozyme.**

Temperature effect on lysozyme was monitored by circular dichroism providing information regarding secondary structure. A) Circular dichroism spectra of lysozyme at temperatures from 15 °C to 90 °C in 5 °C intervals. B) Temperature effect on lysozyme monitored at 212 nm (triangles), 222 nm (squares) and 230 nm (circles). Measurements were performed with lysozyme at 20 mg/mL in 50 mM AcONa pH 4.6. T50% transition corresponds to 354.7 K at 230 nm, 319.2 K at 222 nm and 313.1 at 212 nm. Continuous line are non-linear regression using sigmoidal four-parameter function, and dotted lines are 95 % confidence interval.

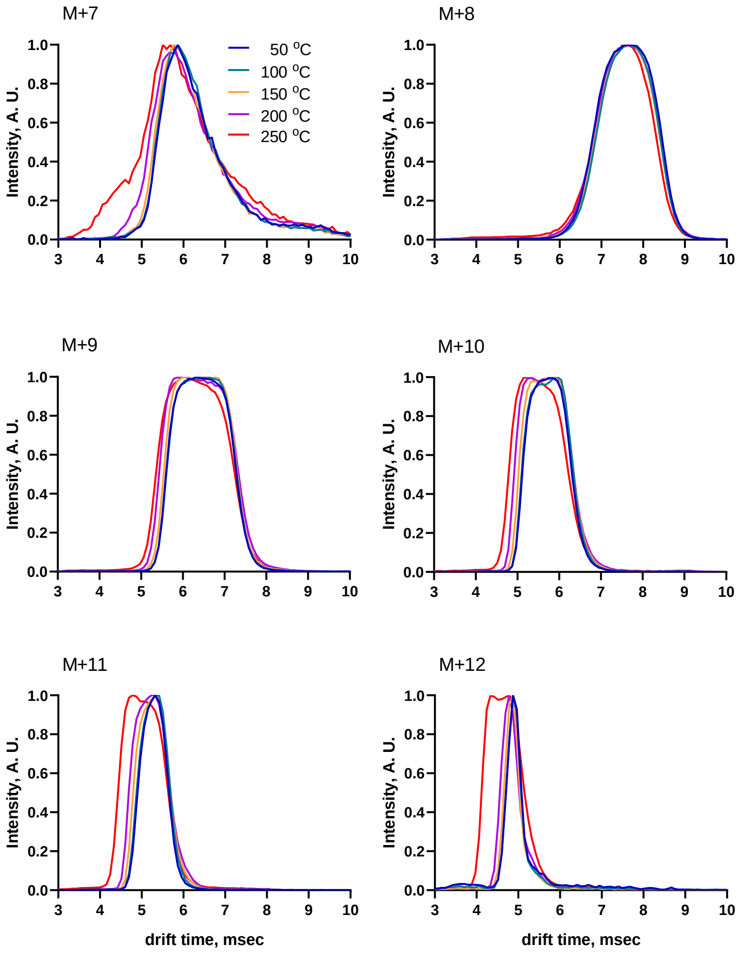


**Figure S5. Thermal effect on lysozyme conformation probed by ion mobility spectrometry.**
The thermal effect on lysozyme was accessed by variations in source temperature upon ionization by electrospray followed by ion mobility spectrometry and mass spectrometry. Each panel corresponds to a charged state of monomeric lysozyme and colors represent different temperatures from 323 K (50 $^oC$) to 523 K (250 $^oC$) in 50 degrees intervals.

| 100 K | | | 125 K | | | 150 K | | | 175 K | | | 200 K | | | 225 K | | | 250 K | | | 275 K | | | 300 K | | | 325 k | | | | |
|---|---|---|---|---|---|---|---|---|---|---|---|---|---|---|---|---|---|---|---|---|---|---|---|---|---|---|---|---|---|---|---|
| N1 | N2 | N3 | N1 | N2 | N3 | N1 | N2 | N3 | N1 | N2 | N3 | N1 | N2 | N3 | N1 | N2 | N3 | N1 | N2 | N3 | N1 | N2 | N3 | N1 | N2 | N3 | N1 | N2 | N3 | | |
| 0.000 | 0.082 | 0.057 | 0.041 | 0.043 | 0.050 | 0.090 | 0.075 | 0.152 | 0.067 | 0.110 | 0.075 | 0.197 | 0.157 | 0.181 | 0.216 | 0.158 | 0.163 | 0.195 | 0.202 | 0.194 | 0.235 | 0.234 | 0.227 | 0.324 | 0.253 | 0.249 | 0.276 | 0.289 | 0.271 | N1 | |
| | 0.000 | 0.048 | 0.087 | 0.087 | 0.072 | 0.046 | 0.052 | 0.105 | 0.118 | 0.085 | 0.075 | 0.157 | 0.114 | 0.137 | 0.189 | 0.149 | 0.154 | 0.194 | 0.202 | 0.189 | 0.241 | 0.234 | 0.227 | 0.340 | 0.256 | 0.254 | 0.281 | 0.297 | 0.279 | N2 | 100 K |
| | | 0.000 | 0.059 | 0.062 | 0.048 | 0.062 | 0.053 | 0.122 | 0.094 | 0.086 | 0.066 | 0.175 | 0.133 | 0.156 | 0.202 | 0.152 | 0.158 | 0.195 | 0.203 | 0.192 | 0.240 | 0.235 | 0.228 | 0.338 | 0.257 | 0.252 | 0.280 | 0.296 | 0.278 | N3 | |
| | | | 0.000 | 0.042 | 0.041 | 0.099 | 0.083 | 0.160 | 0.071 | 0.116 | 0.083 | 0.207 | 0.167 | 0.190 | 0.225 | 0.168 | 0.174 | 0.203 | 0.212 | 0.203 | 0.244 | 0.243 | 0.235 | 0.336 | 0.261 | 0.254 | 0.282 | 0.296 | 0.279 | N1 | |
| | | | | 0.000 | 0.043 | 0.102 | 0.080 | 0.164 | 0.058 | 0.120 | 0.090 | 0.214 | 0.174 | 0.197 | 0.234 | 0.178 | 0.184 | 0.211 | 0.221 | 0.210 | 0.256 | 0.254 | 0.246 | 0.348 | 0.271 | 0.265 | 0.293 | 0.308 | 0.291 | N2 | 125 K |
| | | | | | 0.000 | 0.088 | 0.073 | 0.149 | 0.078 | 0.108 | 0.082 | 0.199 | 0.159 | 0.182 | 0.222 | 0.169 | 0.175 | 0.206 | 0.215 | 0.204 | 0.251 | 0.248 | 0.240 | 0.348 | 0.267 | 0.260 | 0.289 | 0.304 | 0.286 | N3 | |
| | | | | | | 0.000 | 0.051 | 0.082 | 0.125 | 0.067 | 0.068 | 0.138 | 0.093 | 0.119 | 0.173 | 0.126 | 0.132 | 0.176 | 0.183 | 0.172 | 0.222 | 0.220 | 0.209 | 0.322 | 0.239 | 0.238 | 0.266 | 0.281 | 0.262 | N1 | |
| | | | | | | | 0.000 | 0.109 | 0.100 | 0.070 | 0.057 | 0.169 | 0.120 | 0.145 | 0.192 | 0.146 | 0.154 | 0.189 | 0.197 | 0.186 | 0.237 | 0.230 | 0.224 | 0.333 | 0.250 | 0.247 | 0.274 | 0.290 | 0.273 | N2 | 150 K |
| | | | | | | | | 0.000 | 0.184 | 0.074 | 0.109 | 0.096 | 0.055 | 0.076 | 0.130 | 0.112 | 0.123 | 0.168 | 0.172 | 0.163 | 0.209 | 0.203 | 0.194 | 0.310 | 0.224 | 0.226 | 0.252 | 0.268 | 0.250 | N3 | |
| | | | | | | | | | 0.000 | 0.135 | 0.100 | 0.234 | 0.191 | 0.215 | 0.251 | 0.190 | 0.196 | 0.223 | 0.231 | 0.224 | 0.261 | 0.260 | 0.254 | 0.345 | 0.276 | 0.270 | 0.296 | 0.309 | 0.293 | N1 | |
| | | | | | | | | | | 0.000 | 0.058 | 0.145 | 0.097 | 0.123 | 0.165 | 0.119 | 0.134 | 0.172 | 0.179 | 0.171 | 0.218 | 0.212 | 0.204 | 0.316 | 0.231 | 0.228 | 0.256 | 0.271 | 0.256 | N2 | 175 K |
| | | | | | | | | | | | 0.000 | 0.164 | 0.116 | 0.142 | 0.185 | 0.130 | 0.140 | 0.177 | 0.184 | 0.176 | 0.219 | 0.215 | 0.209 | 0.311 | 0.236 | 0.232 | 0.258 | 0.272 | 0.255 | N3 | |
| | | | | | | | | | | | | 0.000 | 0.073 | 0.053 | 0.121 | 0.134 | 0.134 | 0.182 | 0.181 | 0.174 | 0.206 | 0.209 | 0.193 | 0.300 | 0.225 | 0.234 | 0.258 | 0.271 | 0.248 | N1 | |
| | | | | | | | | | | | | | 0.000 | 0.049 | 0.122 | 0.115 | 0.121 | 0.171 | 0.172 | 0.164 | 0.202 | 0.197 | 0.189 | 0.298 | 0.220 | 0.224 | 0.249 | 0.263 | 0.243 | N2 | 200 K |
| | | | | | | | | | | | | | | 0.000 | 0.121 | 0.130 | 0.134 | 0.183 | 0.183 | 0.177 | 0.210 | 0.207 | 0.197 | 0.307 | 0.230 | 0.237 | 0.262 | 0.275 | 0.253 | N3 | |
| | | | | | | | | | | | | | | | 0.000 | 0.123 | 0.129 | 0.153 | 0.150 | 0.148 | 0.166 | 0.153 | 0.153 | 0.256 | 0.179 | 0.189 | 0.210 | 0.225 | 0.203 | N1 | |
| | | | | | | | | | | | | | | | | 0.000 | 0.047 | 0.088 | 0.088 | 0.096 | 0.126 | 0.144 | 0.111 | 0.232 | 0.148 | 0.148 | 0.184 | 0.196 | 0.174 | N2 | 225 K |
| | | | | | | | | | | | | | | | | | 0.000 | 0.083 | 0.079 | 0.087 | 0.124 | 0.149 | 0.108 | 0.225 | 0.144 | 0.149 | 0.184 | 0.197 | 0.173 | N3 | |
| | | | | | | | | | | | | | | | | | | 0.000 | 0.035 | 0.050 | 0.101 | 0.139 | 0.077 | 0.196 | 0.095 | 0.100 | 0.142 | 0.157 | 0.140 | N1 | |
| | | | | | | | | | | | | | | | | | | | 0.000 | 0.053 | 0.091 | 0.133 | 0.066 | 0.185 | 0.089 | 0.098 | 0.139 | 0.152 | 0.132 | N2 | 250 K |
| | | | | | | | | | | | | | | | | | | | | 0.000 | 0.113 | 0.145 | 0.088 | 0.201 | 0.100 | 0.109 | 0.146 | 0.163 | 0.146 | N3 | |
| | | | | | | | | | | | | | | | | | | | | | 0.000 | 0.116 | 0.064 | 0.164 | 0.099 | 0.106 | 0.138 | 0.144 | 0.107 | N1 | |
| | | | | | | | | | | | | | | | | | | | | | | 0.000 | 0.114 | 0.194 | 0.130 | 0.132 | 0.156 | 0.167 | 0.133 | N2 | 275 K |
| | | | | | | | | | | | | | | | | | | | | | | | 0.000 | 0.167 | 0.076 | 0.086 | 0.124 | 0.135 | 0.102 | N3 | |
| | | | | | | | | | | | | | | | | | | | | | | | | 0.000 | 0.139 | 0.149 | 0.139 | 0.122 | 0.131 | N1 | |
| | | | | | | | | | | | | | | | | | | | | | | | | | 0.000 | 0.055 | 0.085 | 0.099 | 0.083 | N2 | 300 K |
| | | | | | | | | | | | | | | | | | | | | | | | | | | 0.000 | 0.081 | 0.089 | 0.087 | N3 | |
| | | | | | | | | | | | | | | | | | | | | | | | | | | | 0.000 | 0.044 | 0.084 | N1 | |
| | | | | | | | | | | | | | | | | | | | | | | | | | | | | 0.000 | 0.091 | N2 | 325 k |
| | | | | | | | | | | | | | | | | | | | | | | | | | | | | | 0.000 | N3 | |

**Figure S6. Pairwise global alignment of crystal structures collected under varying temperatures.** A crystal structure collected at 100 K was used as reference for alignement. Colour indicate similarity, and numbers are overall Cα RMSD (Å). Pairwise alignment was performed with ProSmart (CCP4).

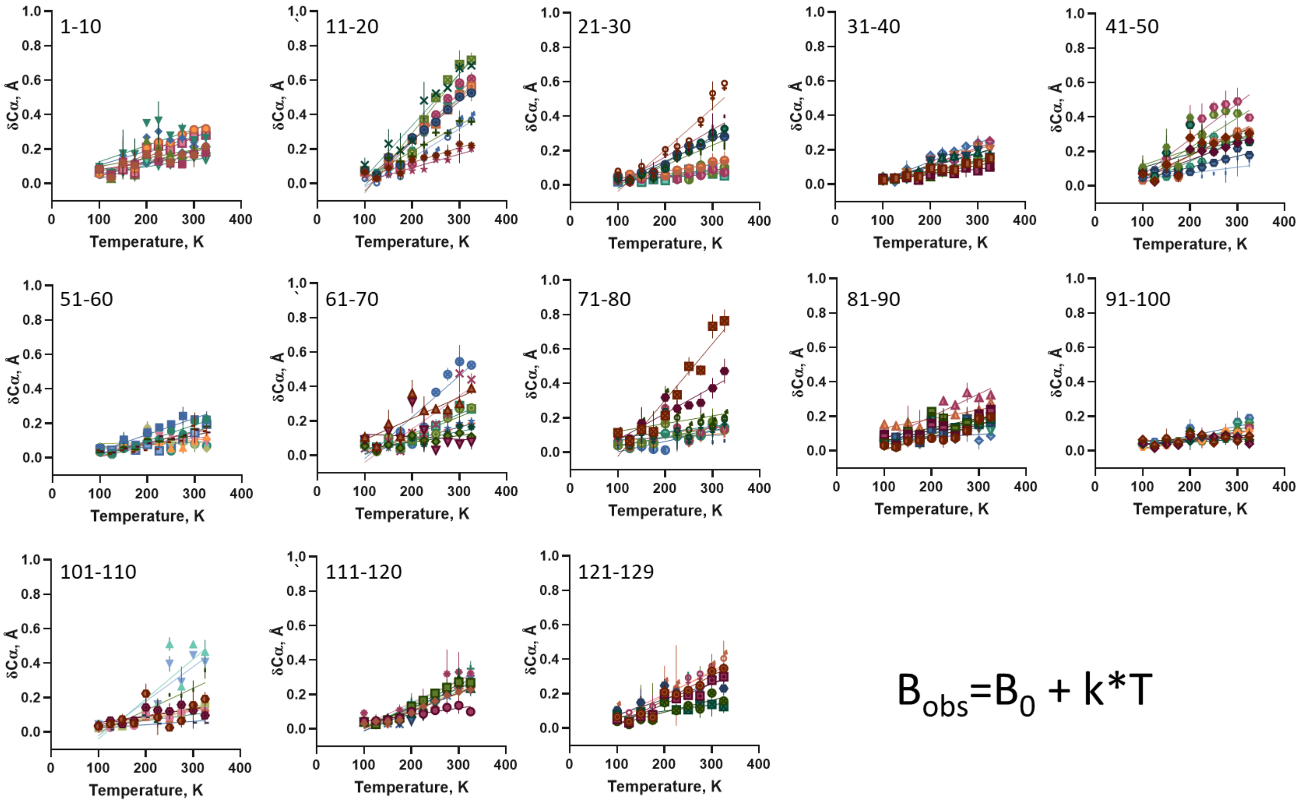


**Figure S7. Temperature-dependent changes in main-chain B-factor**

A) Changes in raw B-factor as a function of temperature for each amino acid main-chain (different colors). Lines are linear regression, from which were obtained the raw $B_0$ and the thermal constant k.

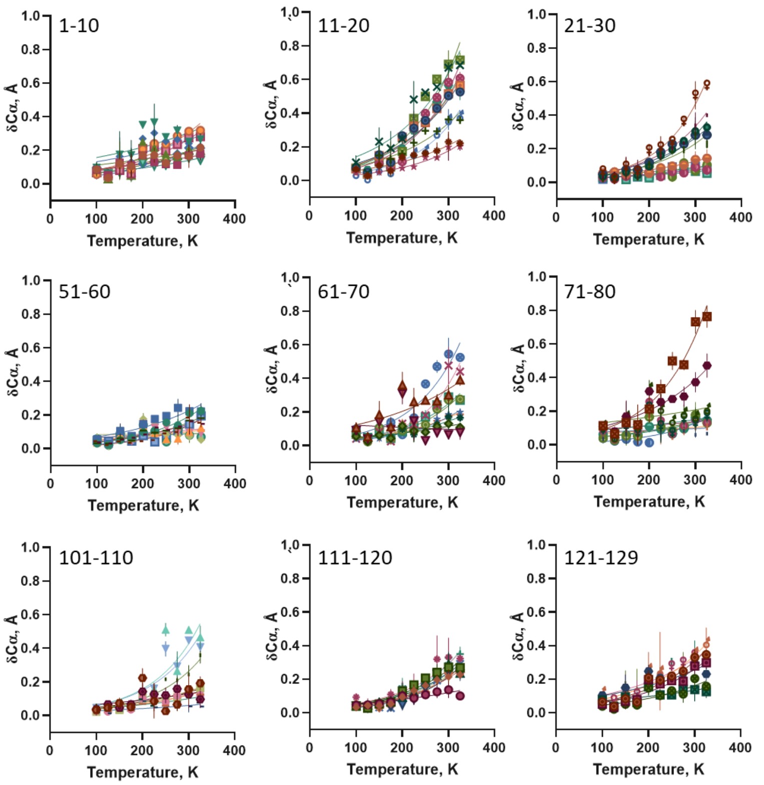


$$B_{obs}=B_0*e^{(k*T)}$$

**Figure S8. Temperature-dependent changes in main-chain B-factor**

A) Changes in raw B-factor as a function of temperature for each amino acid main-chain (different colors). Lines are exponential non-linear regression, from which were obtained the raw $B_0$ and the thermal constant k.

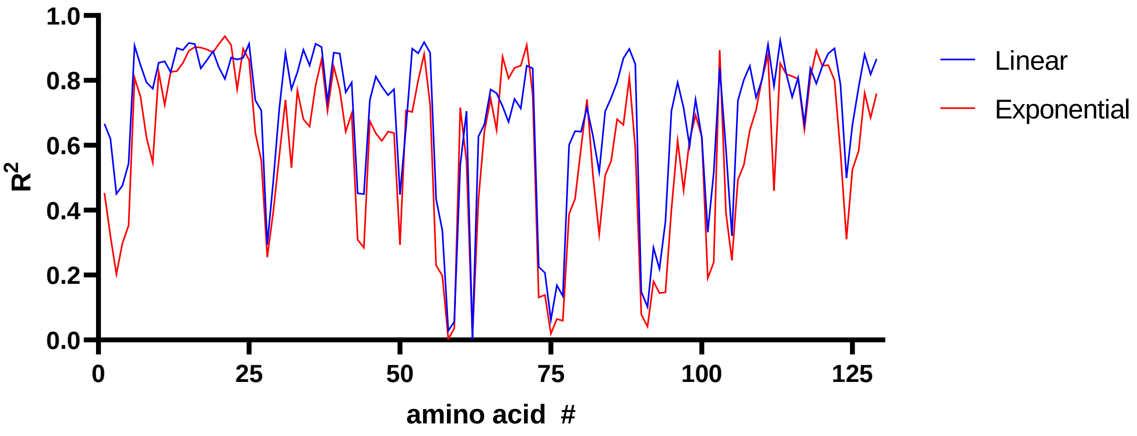


**Figure S9. Correlation coefficient for conformational change in lysozyme Cα as a function of temperature.** Linear (Fig. S2) and exponential (Fig. S3) functions.

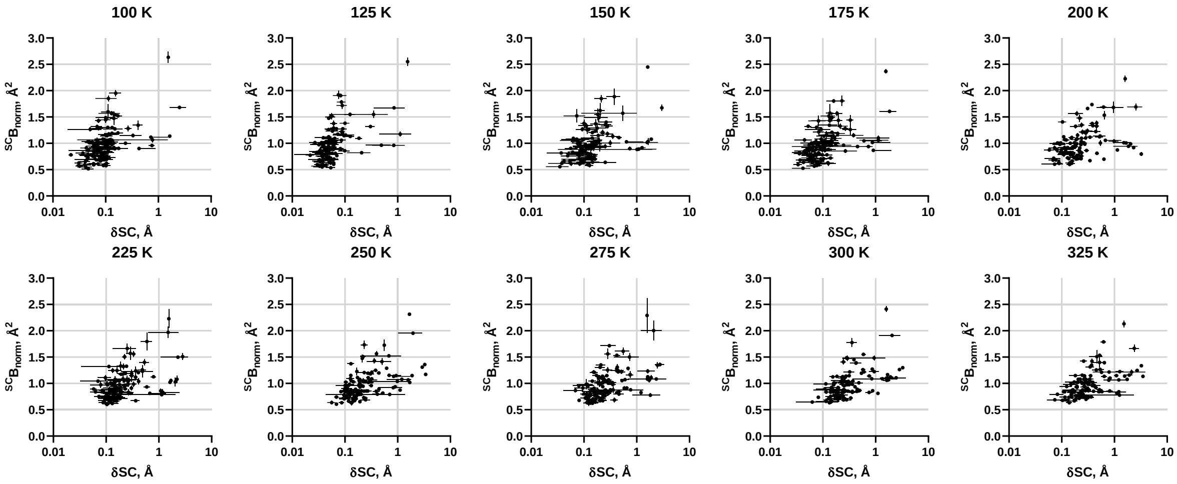


**Figure S10. Profiling side-chain B-factor with conformation.**

Two-dimensional distribution for average amino acid side-chain B-factor *vs* conformational change at varying temperature (as indicated) using a reference structure at 100 K. Symbol is average and bar is standard deviation (n=3).

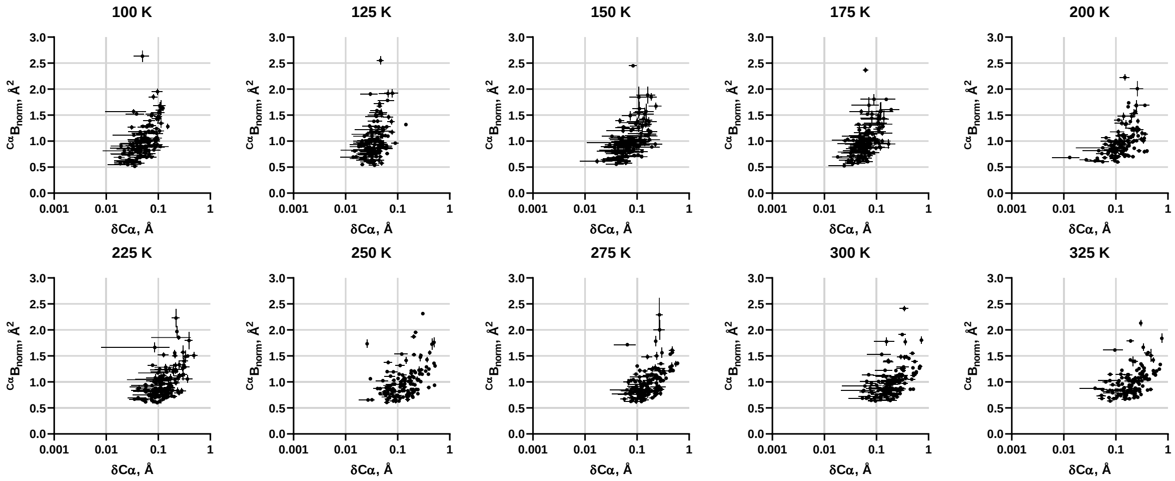


**Figure S11. Profiling Cα B-factor with conformation.**
Two-dimensional distribution for average amino acid Cα B-factor *vs* conformational change at varying temperature (as indicated) using a reference structure at 100 K. Symbol is average and bar is standard deviation (n=3).

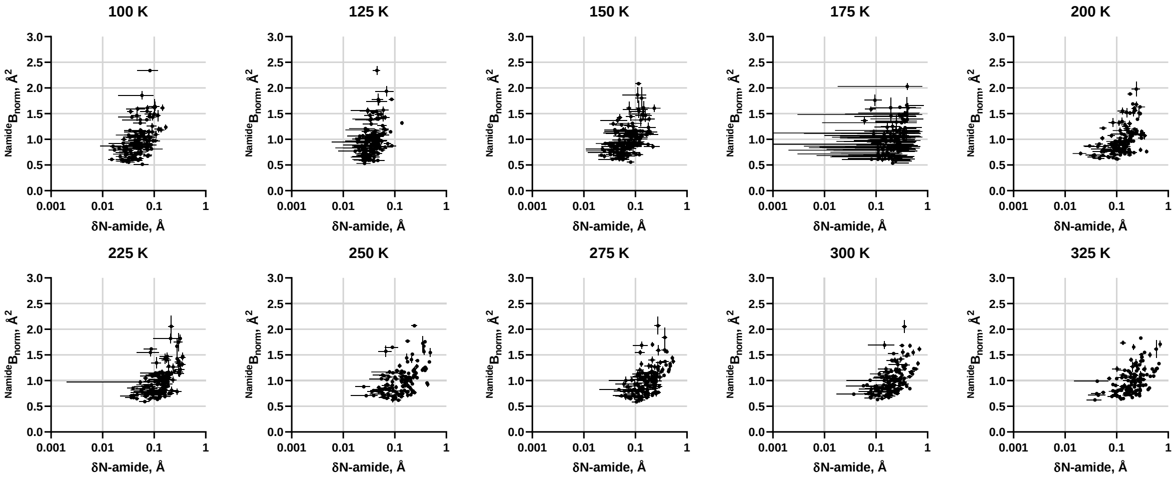


**Figure S12. Profiling amide nitrogen (N-amide) B-factor with conformation.**
Two-dimensional distribution for average amino acid N-amide B-factor *vs* conformational change at varying temperature (as indicated) using a reference structure at 100 K. Symbol is average and bar is standard deviation (n=3).

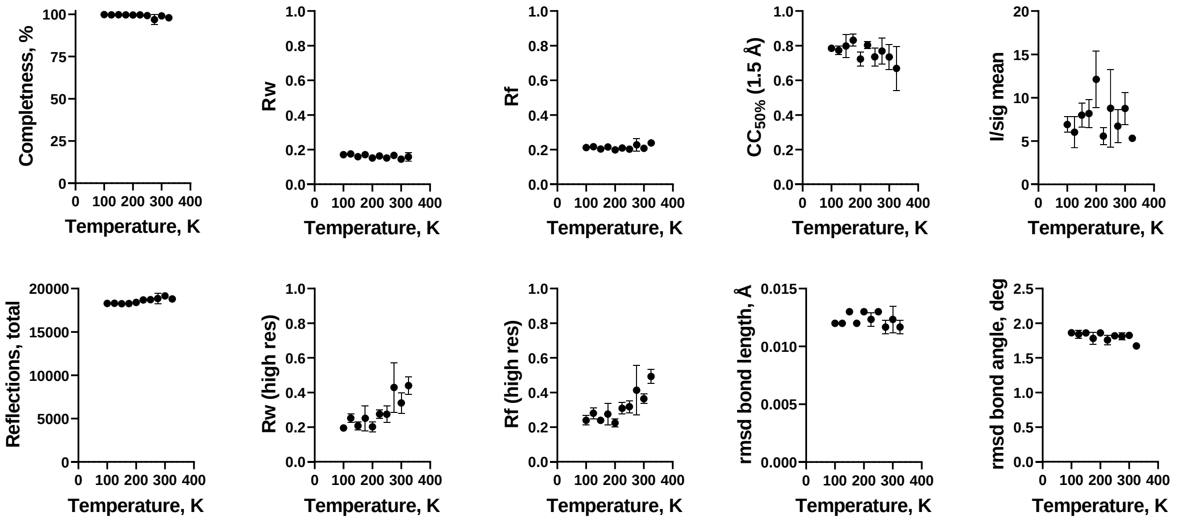


**Figure S13. Crystal quality parameters over temperature**

The data collection and refinement statistics are represented as a function of temperature as depicted in the crystallographic table, demonstrating their stability along the temperature variable. Symbol is average and bar is standard deviation (n=3).
