## Supplementary material for "Linking B-factor and temperature-induced conformational transition": full crystallographic report

### Full wwPDB X-ray Structure Validation Report (i)

Jun 7, 2022 – 11:39 AM EDT

| | | |
|---|---|---|
| PDB ID | : | 8D5S |
| Title | : | Crystal structure of hen egg white lysozyme at 100 Kelvin |
| Deposited on | : | 2022-06-06 |
| Resolution | : | 1.50 Å(reported) |

**This wwPDB validation report is for manuscript review**

For Manuscript Review

This is a Full wwPDB X-ray Structure Validation Report.

This report is produced by the wwPDB biocuration pipeline after annotation of the structure.

We welcome your comments at **
A user guide is available at
https://www.wwpdb.org/validation/2017/XrayValidationReportHelp
with specific help available everywhere you see the (i) symbol.

The following versions of software and data (see references (i)) were used in the production of this report:

| | | |
|---|---|---|
| MolProbity | : | 4.02b-467 |
| Mogul | : | 1.8.5 (274361), CSD as541be (2020) |
| Xtriage (Phenix) | : | 1.13 |
| EDS | : | 2.28 |
| Percentile statistics | : | 20191225.v01 (using entries in the PDB archive December 25th 2019) |
| Refmac | : | 5.8.0158 |
| CCP4 | : | 7.0.044 (Gargrove) |
| Ideal geometry (proteins) | : | Engh & Huber (2001) |
| Ideal geometry (DNA, RNA) | : | Parkinson et al. (1996) |
| Validation Pipeline (wwPDB-VP) | : | 2.28 |

### 1 Overall quality at a glance (i)

The following experimental techniques were used to determine the structure:
*X-RAY DIFFRACTION*

The reported resolution of this entry is 1.50 Å.

Percentile scores (ranging between 0-100) for global validation metrics of the entry are shown in the following graphic. The table shows the number of entries on which the scores are based.

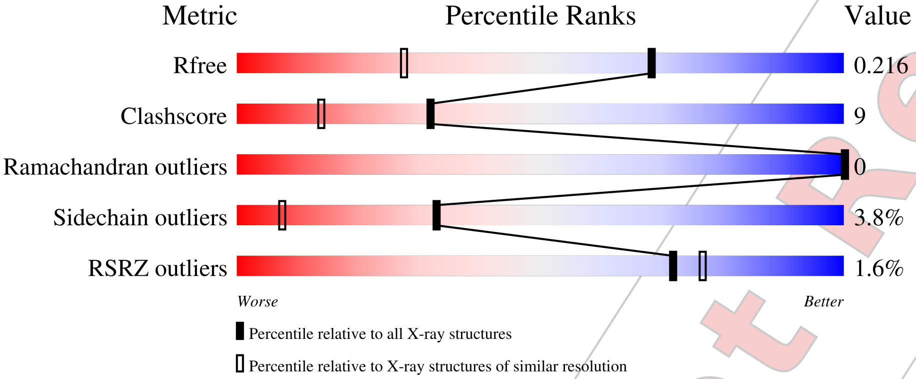


| Metric | Whole archive (#Entries) | Similar resolution (#Entries, resolution range(Å)) |
|---|---|---|
| $R_{free}$ | 130704 | 2936 (1.50-1.50) |
| Clashscore | 141614 | 3144 (1.50-1.50) |
| Ramachandran outliers | 138981 | 3066 (1.50-1.50) |
| Sidechain outliers | 138945 | 3064 (1.50-1.50) |
| RSRZ outliers | 127900 | 2884 (1.50-1.50) |

The table below summarises the geometric issues observed across the polymeric chains and their fit to the electron density. The red, orange, yellow and green segments of the lower bar indicate the fraction of residues that contain outliers for >=3, 2, 1 and 0 types of geometric quality criteria respectively. A grey segment represents the fraction of residues that are not modelled. The numeric value for each fraction is indicated below the corresponding segment, with a dot representing fractions <=5% The upper red bar (where present) indicates the fraction of residues that have poor fit to the electron density. The numeric value is given above the bar.

| Mol | Chain | Length | Quality of chain |
|---|---|---|---|
| 1 | A | 129 | 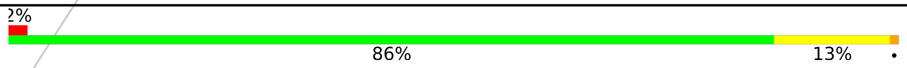  |

For Manuscript Review

### 2 Entry composition (i)

There are 5 unique types of molecules in this entry. The entry contains 1141 atoms, of which 0 are hydrogens and 0 are deuteriums.

In the tables below, the ZeroOcc column contains the number of atoms modelled with zero occupancy, the AltConf column contains the number of residues with at least one atom in alternate conformation and the Trace column contains the number of residues modelled with at most 2 atoms.

- Molecule 1 is a protein called Lysozyme C.

| Mol | Chain | Residues | Atoms | | | | | ZeroOcc | AltConf | Trace |
|---|---|---|---|---|---|---|---|---|---|---|
| | | | Total | C | N | O | S | | | |
| 1 | A | 129 | 1013 | 619 | 195 | 189 | 10 | 0 | 2 | 0 |

- Molecule 2 is SODIUM ION (three-letter code: NA) (formula: Na).

| Mol | Chain | Residues | Atoms | | ZeroOcc | AltConf |
|---|---|---|---|---|---|---|
| | | | Total | Na | | |
| 2 | A | 1 | 1 | 1 | 0 | 0 |

- Molecule 3 is CHLORIDE ION (three-letter code: CL) (formula: Cl).

| Mol | Chain | Residues | Atoms | | ZeroOcc | AltConf |
|---|---|---|---|---|---|---|
| | | | Total | Cl | | |
| 3 | A | 7 | 7 | 7 | 0 | 0 |

- Molecule 4 is GLYCEROL (three-letter code: GOL) (formula: $C_3H_8O_3$).

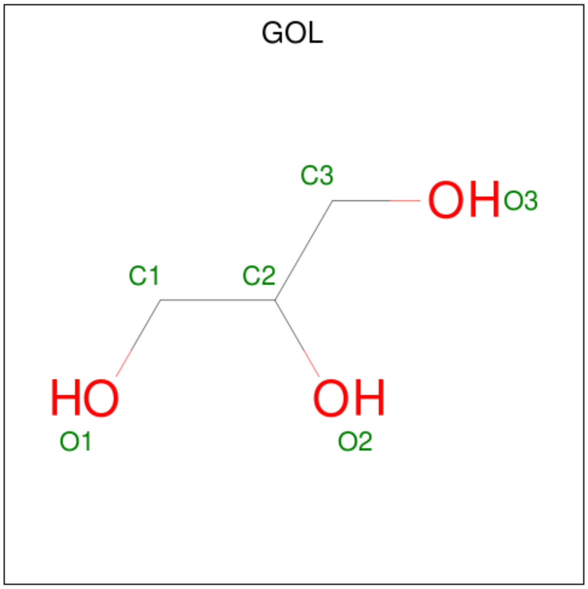


For Manuscript Review

| Mol | Chain | Residues | Atoms | ZeroOcc | AltConf |
|---|---|---|---|---|---|
| 4 | A | 1 | Total C O<br>6 3 3 | 0 | 0 |
| 4 | A | 1 | Total C O<br>6 3 3 | 0 | 0 |

- Molecule 5 is water.

| Mol | Chain | Residues | Atoms | ZeroOcc | AltConf |
|---|---|---|---|---|---|
| 5 | A | 108 | Total O<br>108 108 | 0 | 0 |

For Manuscript Review

#### 3 Residue-property plots (i)

These plots are drawn for all protein, RNA, DNA and oligosaccharide chains in the entry. The first graphic for a chain summarises the proportions of the various outlier classes displayed in the second graphic. The second graphic shows the sequence view annotated by issues in geometry and electron density. Residues are color-coded according to the number of geometric quality criteria for which they contain at least one outlier: green = 0, yellow = 1, orange = 2 and red = 3 or more. A red dot above a residue indicates a poor fit to the electron density (RSRZ > 2). Stretches of 2 or more consecutive residues without any outlier are shown as a green connector. Residues present in the sample, but not in the model, are shown in grey.

• Molecule 1: Lysozyme C

Chain A:

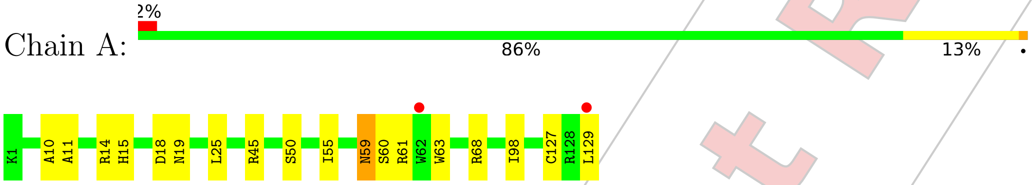


For Manuscript Review

### 4 Data and refinement statistics (i)

| Property | Value | Source |
|---|---|---|
| Space group | P 43 21 2 | Depositor |
| Cell constants<br>a, b, c, $\alpha$, $\beta$, $\gamma$ | 78.83Å 78.83Å 37.24Å<br>90.00° 90.00° 90.00° | Depositor |
| Resolution (Å) | 25.62 – 1.50<br>25.60 – 1.50 | Depositor<br>EDS |
| % Data completeness<br>(in resolution range) | 99.7 (25.62-1.50)<br>99.7 (25.60-1.50) | Depositor<br>EDS |
| $R_{merge}$ | (Not available) | Depositor |
| $R_{sym}$ | (Not available) | Depositor |
| $< I/\sigma(I) >$ [1] | 1.73 (at 1.49Å) | Xtriage |
| Refinement program | REFMAC 5.8.0267 | Depositor |
| R, $R_{free}$ | 0.180 , 0.213<br>0.189 , 0.216 | Depositor<br>DCC |
| $R_{free}$ test set | 994 reflections (5.11%) | wwPDB-VP |
| Wilson B-factor (Å$^2$) | 10.2 | Xtriage |
| Anisotropy | 0.035 | Xtriage |
| Bulk solvent $k_{sol}$(e/Å$^3$), $B_{sol}$(Å$^2$) | 0.35 , 38.4 | EDS |
| L-test for twinning[2] | $< \|L\| > = 0.48$, $< L^2 > = 0.31$ | Xtriage |
| Estimated twinning fraction | No twinning to report. | Xtriage |
| $F_o$,$F_c$ correlation | 0.95 | EDS |
| Total number of atoms | 1141 | wwPDB-VP |
| Average B, all atoms (Å$^2$) | 12.0 | wwPDB-VP |

Xtriage's analysis on translational NCS is as follows: *The largest off-origin peak in the Patterson function is 9.87% of the height of the origin peak. No significant pseudotranslation is detected.*

[1] Intensities estimated from amplitudes.
[2] Theoretical values of $< |L| >$, $< L^2 >$ for acentric reflections are 0.5, 0.333 respectively for untwinned datasets, and 0.375, 0.2 for perfectly twinned datasets.

For Manuscript Review

### 5 Model quality (i)

#### 5.1 Standard geometry (i)

Bond lengths and bond angles in the following residue types are not validated in this section: CL, NA, GOL

The Z score for a bond length (or angle) is the number of standard deviations the observed value is removed from the expected value. A bond length (or angle) with $|Z| > 5$ is considered an outlier worth inspection. RMSZ is the root-mean-square of all Z scores of the bond lengths (or angles).

| Mol | Chain | Bond lengths | | Bond angles | |
|---|---|---|---|---|---|
| | | RMSZ | #$\|Z\|$ >5 | RMSZ | #$\|Z\|$ >5 |
| 1 | A | 0.78 | 0/1033 | 0.92 | 0/1395 |

There are no bond length outliers.

There are no bond angle outliers.

There are no chirality outliers.

There are no planarity outliers.

#### 5.2 Too-close contacts (i)

In the following table, the Non-H and H(model) columns list the number of non-hydrogen atoms and hydrogen atoms in the chain respectively. The H(added) column lists the number of hydrogen atoms added and optimized by MolProbity. The Clashes column lists the number of clashes within the asymmetric unit, whereas Symm-Clashes lists symmetry-related clashes.

| Mol | Chain | Non-H | H(model) | H(added) | Clashes | Symm-Clashes |
|---|---|---|---|---|---|---|
| 1 | A | 1013 | 0 | 967 | 18 | 0 |
| 2 | A | 1 | 0 | 0 | 0 | 0 |
| 3 | A | 7 | 0 | 0 | 0 | 0 |
| 4 | A | 12 | 0 | 16 | 0 | 0 |
| 5 | A | 108 | 0 | 0 | 1 | 0 |
| All | All | 1141 | 0 | 983 | 18 | 0 |

The all-atom clashscore is defined as the number of clashes found per 1000 atoms (including hydrogen atoms). The all-atom clashscore for this structure is 9.

All (18) close contacts within the same asymmetric unit are listed below, sorted by their clash magnitude.

For Manuscript Review

| Atom-1 | Atom-2 | Interatomic distance (Å) | Clash overlap (Å) |
|---|---|---|---|
| 1:A:127:CYS:HB2 | 1:A:129:LEU:HD23 | 1.79 | 0.65 |
| 1:A:127:CYS:HB3 | 1:A:129:LEU:CD2 | 2.28 | 0.63 |
| 1:A:127:CYS:CB | 1:A:129:LEU:CD2 | 2.78 | 0.60 |
| 1:A:11:ALA:O | 1:A:15:HIS:HD2 | 1.85 | 0.59 |
| 1:A:15:HIS:HE1 | 5:A:355:HOH:O | 1.87 | 0.58 |
| 1:A:50:SER:HB2 | 1:A:59:ASN:HD21 | 1.68 | 0.57 |
| 1:A:127:CYS:CB | 1:A:129:LEU:HD23 | 2.37 | 0.55 |
| 1:A:127:CYS:HB3 | 1:A:129:LEU:HD21 | 1.88 | 0.54 |
| 1:A:59:ASN:HD22 | 1:A:60:SER:N | 2.06 | 0.53 |
| 1:A:50:SER:CB | 1:A:59:ASN:HD21 | 2.22 | 0.51 |
| 1:A:50:SER:HB2 | 1:A:59:ASN:ND2 | 2.30 | 0.47 |
| 1:A:45:ARG:NE | 1:A:68:ARG:NH2 | 2.63 | 0.47 |
| 1:A:59:ASN:ND2 | 1:A:61:ARG:H | 2.14 | 0.45 |
| 1:A:127:CYS:HB2 | 1:A:129:LEU:CD2 | 2.41 | 0.45 |
| 1:A:45:ARG:CZ | 1:A:68:ARG:NH2 | 2.80 | 0.44 |
| 1:A:10:ALA:HA | 1:A:129:LEU:CD1 | 2.50 | 0.42 |
| 1:A:18:ASP:HB3 | 1:A:25:LEU:HD13 | 2.01 | 0.42 |
| 1:A:63:TRP:CE2 | 1:A:98:ILE:HG12 | 2.55 | 0.41 |

There are no symmetry-related clashes.

#### 5.3 Torsion angles (i)

##### 5.3.1 Protein backbone (i)

In the following table, the Percentiles column shows the percent Ramachandran outliers of the chain as a percentile score with respect to all X-ray entries followed by that with respect to entries of similar resolution.

The Analysed column shows the number of residues for which the backbone conformation was analysed, and the total number of residues.

| Mol | Chain | Analysed | Favoured | Allowed | Outliers | Percentiles | |
|---|---|---|---|---|---|---|---|
| 1 | A | 129/129 (100%) | 127 (98%) | 2 (2%) | 0 | 100 | 100 |

There are no Ramachandran outliers to report.

##### 5.3.2 Protein sidechains (i)

In the following table, the Percentiles column shows the percent sidechain outliers of the chain as a percentile score with respect to all X-ray entries followed by that with respect to entries of similar resolution.

For Manuscript Review

The Analysed column shows the number of residues for which the sidechain conformation was analysed, and the total number of residues.

| Mol | Chain | Analysed | Rotameric | Outliers | Percentiles | |
|---|---|---|---|---|---|---|
| 1 | A | 107/105 (102%) | 103 (96%) | 4 (4%) | 34 | 8 |

All (4) residues with a non-rotameric sidechain are listed below:

| Mol | Chain | Res | Type |
|---|---|---|---|
| 1 | A | 14 | ARG |
| 1 | A | 19 | ASN |
| 1 | A | 55 | ILE |
| 1 | A | 59 | ASN |

Sometimes sidechains can be flipped to improve hydrogen bonding and reduce clashes. All (4) such sidechains are listed below:

| Mol | Chain | Res | Type |
|---|---|---|---|
| 1 | A | 15 | HIS |
| 1 | A | 19 | ASN |
| 1 | A | 59 | ASN |
| 1 | A | 65 | ASN |

###### 5.3.3 RNA (i)

There are no RNA molecules in this entry.

##### 5.4 Non-standard residues in protein, DNA, RNA chains (i)

There are no non-standard protein/DNA/RNA residues in this entry.

##### 5.5 Carbohydrates (i)

There are no monosaccharides in this entry.

##### 5.6 Ligand geometry (i)

Of 10 ligands modelled in this entry, 8 are monoatomic - leaving 2 for Mogul analysis.

In the following table, the Counts columns list the number of bonds (or angles) for which Mogul statistics could be retrieved, the number of bonds (or angles) that are observed in the model and the number of bonds (or angles) that are defined in the Chemical Component Dictionary. The

For Manuscript Review

Link column lists molecule types, if any, to which the group is linked. The Z score for a bond length (or angle) is the number of standard deviations the observed value is removed from the expected value. A bond length (or angle) with $|Z| > 2$ is considered an outlier worth inspection. RMSZ is the root-mean-square of all Z scores of the bond lengths (or angles).

| Mol | Type | Chain | Res | Link | Bond lengths | | | Bond angles | | |
|---|---|---|---|---|---|---|---|---|---|---|
| | | | | | Counts | RMSZ | $\#\|Z\| > 2$ | Counts | RMSZ | $\#\|Z\| > 2$ |
| 4 | GOL | A | 209 | - | 5,5,5 | 0.14 | 0 | 5,5,5 | 0.37 | 0 |
| 4 | GOL | A | 210 | - | 5,5,5 | 0.12 | 0 | 5,5,5 | 0.21 | 0 |

In the following table, the Chirals column lists the number of chiral outliers, the number of chiral centers analysed, the number of these observed in the model and the number defined in the Chemical Component Dictionary. Similar counts are reported in the Torsion and Rings columns. '-' means no outliers of that kind were identified.

| Mol | Type | Chain | Res | Link | Chirals | Torsions | Rings |
|---|---|---|---|---|---|---|---|
| 4 | GOL | A | 209 | - | - | 0/4/4/4 | - |
| 4 | GOL | A | 210 | - | - | 0/4/4/4 | - |

There are no bond length outliers.

There are no bond angle outliers.

There are no chirality outliers.

There are no torsion outliers.

There are no ring outliers.

No monomer is involved in short contacts.

##### 5.7 Other polymers (i)

There are no such residues in this entry.

##### 5.8 Polymer linkage issues (i)

There are no chain breaks in this entry.

For Manuscript Review

### 6 Fit of model and data (i)

#### 6.1 Protein, DNA and RNA chains (i)

In the following table, the column labelled '#RSRZ> 2' contains the number (and percentage) of RSRZ outliers, followed by percent RSRZ outliers for the chain as percentile scores relative to all X-ray entries and entries of similar resolution. The OWAB column contains the minimum, median, $95^{th}$ percentile and maximum values of the occupancy-weighted average B-factor per residue. The column labelled 'Q< 0.9' lists the number of (and percentage) of residues with an average occupancy less than 0.9.

| Mol | Chain | Analysed | <RSRZ> | #RSRZ>2 | | | OWAB(Å$^2$) | Q<0.9 |
|---|---|---|---|---|---|---|---|---|
| 1 | A | 129/129 (100%) | 0.14 | 2 (1%) | 72 | 77 | 5, 10, 20, 27 | 0 |

All (2) RSRZ outliers are listed below:

| Mol | Chain | Res | Type | RSRZ |
|---|---|---|---|---|
| 1 | A | 129 | LEU | 6.6 |
| 1 | A | 62 | TRP | 2.9 |

#### 6.2 Non-standard residues in protein, DNA, RNA chains (i)

There are no non-standard protein/DNA/RNA residues in this entry.

#### 6.3 Carbohydrates (i)

There are no monosaccharides in this entry.

#### 6.4 Ligands (i)

In the following table, the Atoms column lists the number of modelled atoms in the group and the number defined in the chemical component dictionary. The B-factors column lists the minimum, median, $95^{th}$ percentile and maximum values of B factors of atoms in the group. The column labelled 'Q< 0.9' lists the number of atoms with occupancy less than 0.9.

| Mol | Type | Chain | Res | Atoms | RSCC | RSR | B-factors(Å$^2$) | Q<0.9 |
|---|---|---|---|---|---|---|---|---|
| 4 | GOL | A | 210 | 6/6 | 0.83 | 0.16 | 48,49,50,52 | 0 |
| 4 | GOL | A | 209 | 6/6 | 0.85 | 0.20 | 23,34,37,42 | 0 |
| 3 | CL | A | 208 | 1/1 | 0.95 | 0.21 | 37,37,37,37 | 0 |
| 3 | CL | A | 205 | 1/1 | 0.97 | 0.05 | 23,23,23,23 | 0 |
| 3 | CL | A | 204 | 1/1 | 0.97 | 0.06 | 15,15,15,15 | 0 |

*Continued on next page...*

For Manuscript Review

*Continued from previous page...*

| Mol | Type | Chain | Res | Atoms | RSCC | RSR | B-factors(Å$^2$) | Q<0.9 |
|---|---|---|---|---|---|---|---|---|
| 3 | CL | A | 207 | 1/1 | 0.98 | 0.08 | 19,19,19,19 | 0 |
| 3 | CL | A | 206 | 1/1 | 0.98 | 0.09 | 22,22,22,22 | 0 |
| 3 | CL | A | 202 | 1/1 | 0.99 | 0.04 | 12,12,12,12 | 0 |
| 3 | CL | A | 203 | 1/1 | 0.99 | 0.03 | 14,14,14,14 | 0 |
| 2 | NA | A | 201 | 1/1 | 0.99 | 0.12 | 13,13,13,13 | 0 |

#### 6.5 Other polymers (i)

There are no such residues in this entry.

For Manuscript Review

### Full wwPDB X-ray Structure Validation Report ⓘ

Jun 7, 2022 – 12:14 PM EDT

| | | |
|---|---|---|
| PDB ID | : | 8D5T |
| Title | : | Crystal structure of hen egg white lysozyme at 100 Kelvin (Duplicate) |
| Deposited on | : | 2022-06-06 |
| Resolution | : | 1.50 Å(reported) |

**This wwPDB validation report is for manuscript review**

This is a Full wwPDB X-ray Structure Validation Report.

This report is produced by the wwPDB biocuration pipeline after annotation of the structure.

We welcome your comments at **
A user guide is available at
https://www.wwpdb.org/validation/2017/XrayValidationReportHelp
with specific help available everywhere you see the ⓘ symbol.

The following versions of software and data (see references ⓘ) were used in the production of this report:

| | | |
|---|---|---|
| MolProbity | : | 4.02b-467 |
| Mogul | : | 1.8.5 (274361), CSD as541be (2020) |
| Xtriage (Phenix) | : | 1.13 |
| EDS | : | 2.28 |
| Percentile statistics | : | 20191225.v01 (using entries in the PDB archive December 25th 2019) |
| Refmac | : | 5.8.0158 |
| CCP4 | : | 7.0.044 (Gargrove) |
| Ideal geometry (proteins) | : | Engh & Huber (2001) |
| Ideal geometry (DNA, RNA) | : | Parkinson et al. (1996) |
| Validation Pipeline (wwPDB-VP) | : | 2.28 |

For Manuscript Review

### 1 Overall quality at a glance (i)

The following experimental techniques were used to determine the structure:
*X-RAY DIFFRACTION*

The reported resolution of this entry is 1.50 Å.

Percentile scores (ranging between 0-100) for global validation metrics of the entry are shown in the following graphic. The table shows the number of entries on which the scores are based.

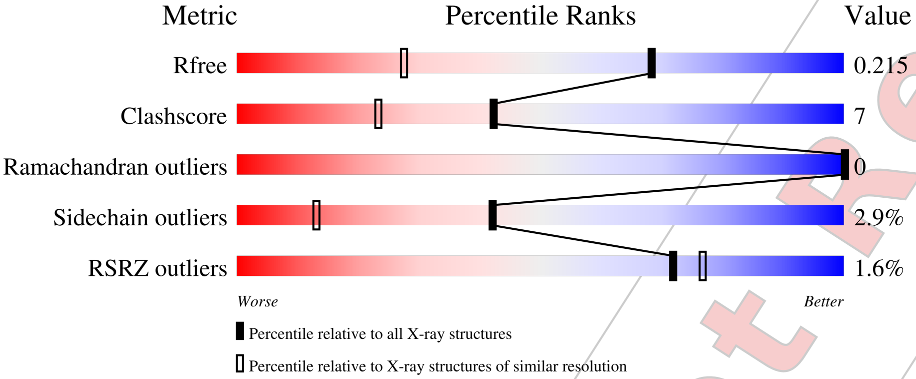


| Metric | Whole archive (#Entries) | Similar resolution (#Entries, resolution range(Å)) |
|---|---|---|
| $R_{free}$ | 130704 | 2936 (1.50-1.50) |
| Clashscore | 141614 | 3144 (1.50-1.50) |
| Ramachandran outliers | 138981 | 3066 (1.50-1.50) |
| Sidechain outliers | 138945 | 3064 (1.50-1.50) |
| RSRZ outliers | 127900 | 2884 (1.50-1.50) |

The table below summarises the geometric issues observed across the polymeric chains and their fit to the electron density. The red, orange, yellow and green segments of the lower bar indicate the fraction of residues that contain outliers for >=3, 2, 1 and 0 types of geometric quality criteria respectively. A grey segment represents the fraction of residues that are not modelled. The numeric value for each fraction is indicated below the corresponding segment, with a dot representing fractions <=5% The upper red bar (where present) indicates the fraction of residues that have poor fit to the electron density. The numeric value is given above the bar.

| Mol | Chain | Length | Quality of chain |
|---|---|---|---|
| 1 | A | 129 | 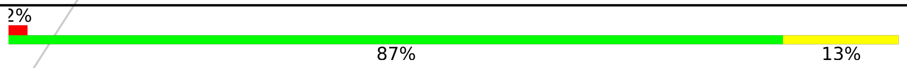 |

For Manuscript Review

### 2 Entry composition (i)

There are 5 unique types of molecules in this entry. The entry contains 1153 atoms, of which 0 are hydrogens and 0 are deuteriums.

In the tables below, the ZeroOcc column contains the number of atoms modelled with zero occupancy, the AltConf column contains the number of residues with at least one atom in alternate conformation and the Trace column contains the number of residues modelled with at most 2 atoms.

- Molecule 1 is a protein called Lysozyme C.

| Mol | Chain | Residues | Atoms | | | | | ZeroOcc | AltConf | Trace |
|---|---|---|---|---|---|---|---|---|---|---|
| | | | Total | C | N | O | S | | | |
| 1 | A | 129 | 1013 | 619 | 195 | 189 | 10 | 0 | 2 | 0 |

- Molecule 2 is SODIUM ION (three-letter code: NA) (formula: Na).

| Mol | Chain | Residues | Atoms | | ZeroOcc | AltConf |
|---|---|---|---|---|---|---|
| | | | Total | Na | | |
| 2 | A | 1 | 1 | 1 | 0 | 0 |

- Molecule 3 is CHLORIDE ION (three-letter code: CL) (formula: Cl).

| Mol | Chain | Residues | Atoms | | ZeroOcc | AltConf |
|---|---|---|---|---|---|---|
| | | | Total | Cl | | |
| 3 | A | 7 | 7 | 7 | 0 | 0 |

- Molecule 4 is GLYCEROL (three-letter code: GOL) (formula: $C_3H_8O_3$).

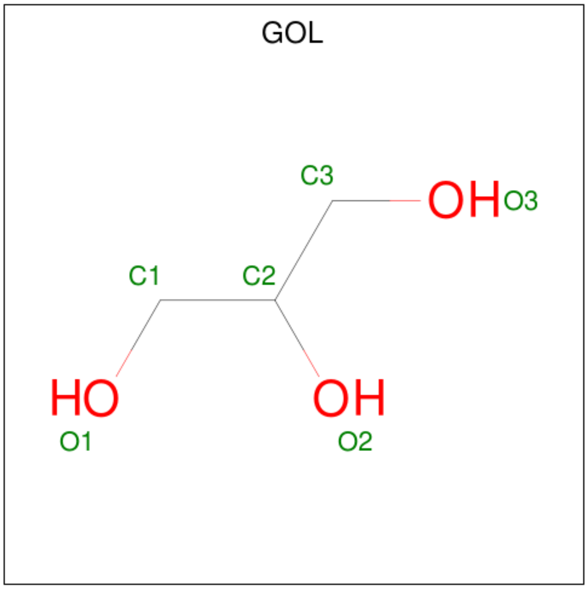


For Manuscript Review

| Mol | Chain | Residues | Atoms | ZeroOcc | AltConf |
|---|---|---|---|---|---|
| 4 | A | 1 | Total C O<br>6 3 3 | 0 | 0 |
| 4 | A | 1 | Total C O<br>6 3 3 | 0 | 0 |

- Molecule 5 is water.

| Mol | Chain | Residues | Atoms | ZeroOcc | AltConf |
|---|---|---|---|---|---|
| 5 | A | 120 | Total O<br>120 120 | 0 | 0 |

For Manuscript Review

### 3 Residue-property plots (i)

These plots are drawn for all protein, RNA, DNA and oligosaccharide chains in the entry. The first graphic for a chain summarises the proportions of the various outlier classes displayed in the second graphic. The second graphic shows the sequence view annotated by issues in geometry and electron density. Residues are color-coded according to the number of geometric quality criteria for which they contain at least one outlier: green = 0, yellow = 1, orange = 2 and red = 3 or more. A red dot above a residue indicates a poor fit to the electron density (RSRZ > 2). Stretches of 2 or more consecutive residues without any outlier are shown as a green connector. Residues present in the sample, but not in the model, are shown in grey.

- Molecule 1: Lysozyme C

Chain A:

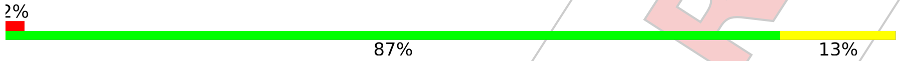


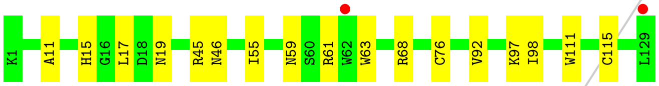


For Manuscript Review

### 4 Data and refinement statistics (i)

| Property | Value | Source |
|---|---|---|
| Space group | P 43 21 2 | Depositor |
| Cell constants<br>a, b, c, $\alpha$, $\beta$, $\gamma$ | 78.63Å 78.63Å 37.07Å<br>90.00° 90.00° 90.00° | Depositor |
| Resolution (Å) | 24.88 – 1.50<br>24.86 – 1.50 | Depositor<br>EDS |
| % Data completeness<br>(in resolution range) | 99.9 (24.88-1.50)<br>100.0 (24.86-1.50) | Depositor<br>EDS |
| $R_{merge}$ | (Not available) | Depositor |
| $R_{sym}$ | (Not available) | Depositor |
| $< I/\sigma(I) >$ [1] | 6.11 (at 1.50Å) | Xtriage |
| Refinement program | REFMAC 5.8.0267 | Depositor |
| R, $R_{free}$ | 0.175 , 0.212<br>0.184 , 0.215 | Depositor<br>DCC |
| $R_{free}$ test set | 983 reflections (5.13%) | wwPDB-VP |
| Wilson B-factor (Å$^2$) | 9.6 | Xtriage |
| Anisotropy | 0.048 | Xtriage |
| Bulk solvent $k_{sol}$(e/Å$^3$), $B_{sol}$(Å$^2$) | 0.36 , 39.4 | EDS |
| L-test for twinning[2] | $< \|L\| >$ = 0.49, $< L^2 >$ = 0.32 | Xtriage |
| Estimated twinning fraction | No twinning to report. | Xtriage |
| $F_o$,$F_c$ correlation | 0.95 | EDS |
| Total number of atoms | 1153 | wwPDB-VP |
| Average B, all atoms (Å$^2$) | 12.0 | wwPDB-VP |

Xtriage's analysis on translational NCS is as follows: *The largest off-origin peak in the Patterson function is 8.42% of the height of the origin peak. No significant pseudotranslation is detected.*

[1] Intensities estimated from amplitudes.

[2] Theoretical values of $< |L| >$, $< L^2 >$ for acentric reflections are 0.5, 0.333 respectively for untwinned datasets, and 0.375, 0.2 for perfectly twinned datasets.

For Manuscript Review

### 5 Model quality (i)

#### 5.1 Standard geometry (i)

Bond lengths and bond angles in the following residue types are not validated in this section: GOL, CL, NA

The Z score for a bond length (or angle) is the number of standard deviations the observed value is removed from the expected value. A bond length (or angle) with $|Z| > 5$ is considered an outlier worth inspection. RMSZ is the root-mean-square of all Z scores of the bond lengths (or angles).

| Mol | Chain | Bond lengths | | Bond angles | |
|---|---|---|---|---|---|
| | | RMSZ | #$\|Z\|$ >5 | RMSZ | #$\|Z\|$ >5 |
| 1 | A | 0.76 | 0/1033 | 0.92 | 0/1395 |

There are no bond length outliers.

There are no bond angle outliers.

There are no chirality outliers.

There are no planarity outliers.

#### 5.2 Too-close contacts (i)

In the following table, the Non-H and H(model) columns list the number of non-hydrogen atoms and hydrogen atoms in the chain respectively. The H(added) column lists the number of hydrogen atoms added and optimized by MolProbity. The Clashes column lists the number of clashes within the asymmetric unit, whereas Symm-Clashes lists symmetry-related clashes.

| Mol | Chain | Non-H | H(model) | H(added) | Clashes | Symm-Clashes |
|---|---|---|---|---|---|---|
| 1 | A | 1013 | 0 | 967 | 14 | 0 |
| 2 | A | 1 | 0 | 0 | 0 | 0 |
| 3 | A | 7 | 0 | 0 | 0 | 0 |
| 4 | A | 12 | 0 | 16 | 0 | 0 |
| 5 | A | 120 | 0 | 0 | 1 | 0 |
| All | All | 1153 | 0 | 983 | 14 | 0 |

The all-atom clashscore is defined as the number of clashes found per 1000 atoms (including hydrogen atoms). The all-atom clashscore for this structure is 7.

All (14) close contacts within the same asymmetric unit are listed below, sorted by their clash magnitude.

For Manuscript Review

| Atom-1 | Atom-2 | Interatomic distance (Å) | Clash overlap (Å) |
|---|---|---|---|
| 1:A:59:ASN:ND2 | 1:A:61:ARG:HB3 | 2.08 | 0.68 |
| 1:A:59:ASN:HD22 | 1:A:61:ARG:HB3 | 1.58 | 0.67 |
| 1:A:59:ASN:ND2 | 1:A:61:ARG:CB | 2.63 | 0.60 |
| 1:A:59:ASN:ND2 | 1:A:61:ARG:H | 2.01 | 0.59 |
| 1:A:11:ALA:O | 1:A:15:HIS:HD2 | 1.86 | 0.57 |
| 1:A:45:ARG:NE | 1:A:68:ARG:NH2 | 2.54 | 0.56 |
| 1:A:68:ARG:HD2 | 5:A:312:HOH:O | 2.08 | 0.53 |
| 1:A:46:ASN:ND2 | 1:A:59:ASN:OD1 | 2.42 | 0.52 |
| 1:A:17:LEU:HG | 1:A:92:VAL:HG13 | 1.92 | 0.51 |
| 1:A:59:ASN:HD21 | 1:A:61:ARG:HB2 | 1.82 | 0.44 |
| 1:A:63:TRP:CE2 | 1:A:98:ILE:HG12 | 2.55 | 0.42 |
| 1:A:45:ARG:CZ | 1:A:68:ARG:NH2 | 2.82 | 0.41 |
| 1:A:111:TRP:CD1 | 1:A:115:CYS:HB2 | 2.57 | 0.40 |
| 1:A:63:TRP:O | 1:A:76:CYS:HB2 | 2.21 | 0.40 |

There are no symmetry-related clashes.

#### 5.3 Torsion angles (i)

##### 5.3.1 Protein backbone (i)

In the following table, the Percentiles column shows the percent Ramachandran outliers of the chain as a percentile score with respect to all X-ray entries followed by that with respect to entries of similar resolution.

The Analysed column shows the number of residues for which the backbone conformation was analysed, and the total number of residues.

| Mol | Chain | Analysed | Favoured | Allowed | Outliers | Percentiles | |
|---|---|---|---|---|---|---|---|
| 1 | A | 129/129 (100%) | 128 (99%) | 1 (1%) | 0 | 100 | 100 |

There are no Ramachandran outliers to report.

##### 5.3.2 Protein sidechains (i)

In the following table, the Percentiles column shows the percent sidechain outliers of the chain as a percentile score with respect to all X-ray entries followed by that with respect to entries of similar resolution.

The Analysed column shows the number of residues for which the sidechain conformation was analysed, and the total number of residues.

For Manuscript Review

| Mol | Chain | Analysed | Rotameric | Outliers | Percentiles | |
|---|---|---|---|---|---|---|
| 1 | A | 107/105 (102%) | 104 (97%) | 3 (3%) | 43 | 14 |

All (3) residues with a non-rotameric sidechain are listed below:

| Mol | Chain | Res | Type |
|---|---|---|---|
| 1 | A | 19 | ASN |
| 1 | A | 55 | ILE |
| 1 | A | 97 | LYS |

Sometimes sidechains can be flipped to improve hydrogen bonding and reduce clashes. All (5) such sidechains are listed below:

| Mol | Chain | Res | Type |
|---|---|---|---|
| 1 | A | 15 | HIS |
| 1 | A | 19 | ASN |
| 1 | A | 46 | ASN |
| 1 | A | 59 | ASN |
| 1 | A | 65 | ASN |

##### 5.3.3 RNA (i)

There are no RNA molecules in this entry.

#### 5.4 Non-standard residues in protein, DNA, RNA chains (i)

There are no non-standard protein/DNA/RNA residues in this entry.

#### 5.5 Carbohydrates (i)

There are no monosaccharides in this entry.

#### 5.6 Ligand geometry (i)

Of 10 ligands modelled in this entry, 8 are monoatomic - leaving 2 for Mogul analysis.

In the following table, the Counts columns list the number of bonds (or angles) for which Mogul statistics could be retrieved, the number of bonds (or angles) that are observed in the model and the number of bonds (or angles) that are defined in the Chemical Component Dictionary. The Link column lists molecule types, if any, to which the group is linked. The Z score for a bond length (or angle) is the number of standard deviations the observed value is removed from the

For Manuscript Review

expected value. A bond length (or angle) with $|Z| > 2$ is considered an outlier worth inspection. RMSZ is the root-mean-square of all Z scores of the bond lengths (or angles).

| Mol | Type | Chain | Res | Link | Bond lengths | | | Bond angles | | |
|---|---|---|---|---|---|---|---|---|---|---|
| | | | | | Counts | RMSZ | $\#\|Z\| > 2$ | Counts | RMSZ | $\#\|Z\| > 2$ |
| 4 | GOL | A | 209 | - | 5,5,5 | 0.15 | 0 | 5,5,5 | 0.38 | 0 |
| 4 | GOL | A | 210 | - | 5,5,5 | 0.11 | 0 | 5,5,5 | 0.19 | 0 |

In the following table, the Chirals column lists the number of chiral outliers, the number of chiral centers analysed, the number of these observed in the model and the number defined in the Chemical Component Dictionary. Similar counts are reported in the Torsion and Rings columns. '-' means no outliers of that kind were identified.

| Mol | Type | Chain | Res | Link | Chirals | Torsions | Rings |
|---|---|---|---|---|---|---|---|
| 4 | GOL | A | 209 | - | - | 0/4/4/4 | - |
| 4 | GOL | A | 210 | - | - | 0/4/4/4 | - |

There are no bond length outliers.

There are no bond angle outliers.

There are no chirality outliers.

There are no torsion outliers.

There are no ring outliers.

No monomer is involved in short contacts.

#### 5.7 Other polymers (i)

There are no such residues in this entry.

#### 5.8 Polymer linkage issues (i)

There are no chain breaks in this entry.

For Manuscript Review

### 6 Fit of model and data (i)

#### 6.1 Protein, DNA and RNA chains (i)

In the following table, the column labelled '#RSRZ> 2' contains the number (and percentage) of RSRZ outliers, followed by percent RSRZ outliers for the chain as percentile scores relative to all X-ray entries and entries of similar resolution. The OWAB column contains the minimum, median, $95^{th}$ percentile and maximum values of the occupancy-weighted average B-factor per residue. The column labelled 'Q< 0.9' lists the number of (and percentage) of residues with an average occupancy less than 0.9.

| Mol | Chain | Analysed | <RSRZ> | #RSRZ>2 | | | OWAB(Å$^2$) | Q<0.9 |
|---|---|---|---|---|---|---|---|---|
| 1 | A | 129/129 (100%) | -0.04 | 2 (1%) | 72 | 77 | 5, 10, 19, 26 | 0 |

All (2) RSRZ outliers are listed below:

| Mol | Chain | Res | Type | RSRZ |
|---|---|---|---|---|
| 1 | A | 129 | LEU | 3.1 |
| 1 | A | 62 | TRP | 2.1 |

#### 6.2 Non-standard residues in protein, DNA, RNA chains (i)

There are no non-standard protein/DNA/RNA residues in this entry.

#### 6.3 Carbohydrates (i)

There are no monosaccharides in this entry.

#### 6.4 Ligands (i)

In the following table, the Atoms column lists the number of modelled atoms in the group and the number defined in the chemical component dictionary. The B-factors column lists the minimum, median, $95^{th}$ percentile and maximum values of B factors of atoms in the group. The column labelled 'Q< 0.9' lists the number of atoms with occupancy less than 0.9.

| Mol | Type | Chain | Res | Atoms | RSCC | RSR | B-factors(Å$^2$) | Q<0.9 |
|---|---|---|---|---|---|---|---|---|
| 4 | GOL | A | 210 | 6/6 | 0.74 | 0.16 | 40,41,42,45 | 0 |
| 4 | GOL | A | 209 | 6/6 | 0.84 | 0.19 | 23,31,33,39 | 0 |
| 3 | CL | A | 208 | 1/1 | 0.94 | 0.14 | 29,29,29,29 | 0 |
| 3 | CL | A | 207 | 1/1 | 0.97 | 0.06 | 19,19,19,19 | 0 |
| 3 | CL | A | 204 | 1/1 | 0.97 | 0.06 | 15,15,15,15 | 0 |

*Continued on next page...*

For Manuscript Review

*Continued from previous page...*

| Mol | Type | Chain | Res | Atoms | RSCC | RSR | B-factors($Å^2$) | Q<0.9 |
|---|---|---|---|---|---|---|---|---|
| 3 | CL | A | 205 | 1/1 | 0.98 | 0.06 | 22,22,22,22 | 0 |
| 3 | CL | A | 206 | 1/1 | 0.98 | 0.06 | 19,19,19,19 | 0 |
| 3 | CL | A | 202 | 1/1 | 0.99 | 0.04 | 12,12,12,12 | 0 |
| 3 | CL | A | 203 | 1/1 | 1.00 | 0.03 | 11,11,11,11 | 0 |
| 2 | NA | A | 201 | 1/1 | 1.00 | 0.04 | 9,9,9,9 | 0 |

#### 6.5 Other polymers (i)

There are no such residues in this entry.

For Manuscript Review

For Manuscript Review

### Full wwPDB X-ray Structure Validation Report (i)

Jun 7, 2022 – 12:28 PM EDT

| | | |
|---|---|---|
| PDB ID | : | 8D5U |
| Title | : | Crystal structure of hen egg white lysozyme at 100 Kelvin (Triplicate) |
| Deposited on | : | 2022-06-06 |
| Resolution | : | 1.50 Å(reported) |

**This wwPDB validation report is for manuscript review**

This is a Full wwPDB X-ray Structure Validation Report.

This report is produced by the wwPDB biocuration pipeline after annotation of the structure.

We welcome your comments at **
A user guide is available at
https://www.wwpdb.org/validation/2017/XrayValidationReportHelp
with specific help available everywhere you see the (i) symbol.

---

The following versions of software and data (see references (i)) were used in the production of this report:

| | | |
|---|---|---|
| MolProbity | : | 4.02b-467 |
| Mogul | : | 1.8.5 (274361), CSD as541be (2020) |
| Xtriage (Phenix) | : | 1.13 |
| EDS | : | 2.28 |
| Percentile statistics | : | 20191225.v01 (using entries in the PDB archive December 25th 2019) |
| Refmac | : | 5.8.0158 |
| CCP4 | : | 7.0.044 (Gargrove) |
| Ideal geometry (proteins) | : | Engh & Huber (2001) |
| Ideal geometry (DNA, RNA) | : | Parkinson et al. (1996) |
| Validation Pipeline (wwPDB-VP) | : | 2.28 |

### 1 Overall quality at a glance (i)

The following experimental techniques were used to determine the structure:
*X-RAY DIFFRACTION*

The reported resolution of this entry is 1.50 Å.

Percentile scores (ranging between 0-100) for global validation metrics of the entry are shown in the following graphic. The table shows the number of entries on which the scores are based.

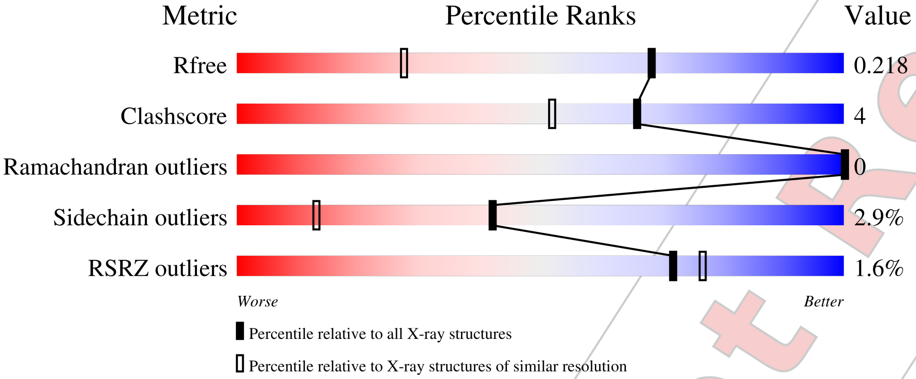


| Metric | Whole archive (#Entries) | Similar resolution (#Entries, resolution range(Å)) |
|---|---|---|
| $R_{free}$ | 130704 | 2936 (1.50-1.50) |
| Clashscore | 141614 | 3144 (1.50-1.50) |
| Ramachandran outliers | 138981 | 3066 (1.50-1.50) |
| Sidechain outliers | 138945 | 3064 (1.50-1.50) |
| RSRZ outliers | 127900 | 2884 (1.50-1.50) |

The table below summarises the geometric issues observed across the polymeric chains and their fit to the electron density. The red, orange, yellow and green segments of the lower bar indicate the fraction of residues that contain outliers for >=3, 2, 1 and 0 types of geometric quality criteria respectively. A grey segment represents the fraction of residues that are not modelled. The numeric value for each fraction is indicated below the corresponding segment, with a dot representing fractions <=5% The upper red bar (where present) indicates the fraction of residues that have poor fit to the electron density. The numeric value is given above the bar.

| Mol | Chain | Length | Quality of chain |
|---|---|---|---|
| 1 | A | 129 | 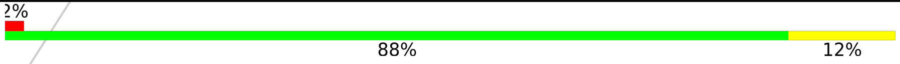 |

For Manuscript Review

### 2 Entry composition (i)

There are 5 unique types of molecules in this entry. The entry contains 1145 atoms, of which 0 are hydrogens and 0 are deuteriums.

In the tables below, the ZeroOcc column contains the number of atoms modelled with zero occupancy, the AltConf column contains the number of residues with at least one atom in alternate conformation and the Trace column contains the number of residues modelled with at most 2 atoms.

- Molecule 1 is a protein called Lysozyme C.

| Mol | Chain | Residues | Atoms | | | | | ZeroOcc | AltConf | Trace |
|---|---|---|---|---|---|---|---|---|---|---|
| | | | Total | C | N | O | S | | | |
| 1 | A | 129 | 1013 | 619 | 195 | 189 | 10 | 0 | 2 | 0 |

- Molecule 2 is SODIUM ION (three-letter code: NA) (formula: Na).

| Mol | Chain | Residues | Atoms | | ZeroOcc | AltConf |
|---|---|---|---|---|---|---|
| | | | Total | Na | | |
| 2 | A | 1 | 1 | 1 | 0 | 0 |

- Molecule 3 is CHLORIDE ION (three-letter code: CL) (formula: Cl).

| Mol | Chain | Residues | Atoms | | ZeroOcc | AltConf |
|---|---|---|---|---|---|---|
| | | | Total | Cl | | |
| 3 | A | 7 | 7 | 7 | 0 | 0 |

- Molecule 4 is GLYCEROL (three-letter code: GOL) (formula: $C_3H_8O_3$).

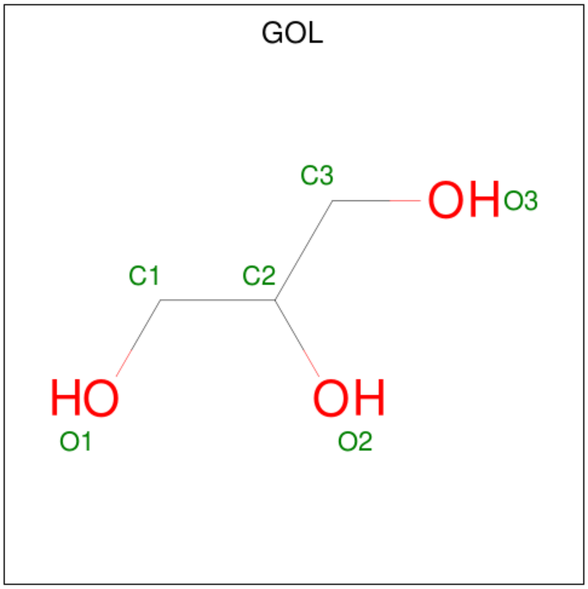


For Manuscript Review

| Mol | Chain | Residues | Atoms | ZeroOcc | AltConf |
|---|---|---|---|---|---|
| 4 | A | 1 | Total C O<br>6 3 3 | 0 | 0 |
| 4 | A | 1 | Total C O<br>6 3 3 | 0 | 0 |

- Molecule 5 is water.

| Mol | Chain | Residues | Atoms | ZeroOcc | AltConf |
|---|---|---|---|---|---|
| 5 | A | 112 | Total O<br>112 112 | 0 | 0 |

For Manuscript Review

### 3 Residue-property plots (i)

These plots are drawn for all protein, RNA, DNA and oligosaccharide chains in the entry. The first graphic for a chain summarises the proportions of the various outlier classes displayed in the second graphic. The second graphic shows the sequence view annotated by issues in geometry and electron density. Residues are color-coded according to the number of geometric quality criteria for which they contain at least one outlier: green = 0, yellow = 1, orange = 2 and red = 3 or more. A red dot above a residue indicates a poor fit to the electron density (RSRZ > 2). Stretches of 2 or more consecutive residues without any outlier are shown as a green connector. Residues present in the sample, but not in the model, are shown in grey.

- Molecule 1: Lysozyme C

Chain A:

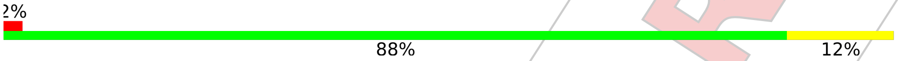


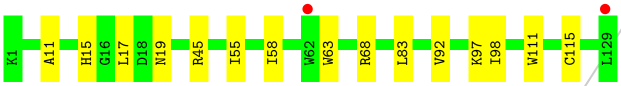


For Manuscript Review

### 4 Data and refinement statistics (i)

| Property | Value | Source |
|---|---|---|
| Space group | P 43 21 2 | Depositor |
| Cell constants<br>a, b, c, $\alpha$, $\beta$, $\gamma$ | 78.68Å 78.68Å 37.11Å<br>90.00° 90.00° 90.00° | Depositor |
| Resolution (Å) | 22.27 – 1.50<br>22.26 – 1.50 | Depositor<br>EDS |
| % Data completeness<br>(in resolution range) | 99.8 (22.27-1.50)<br>99.8 (22.26-1.50) | Depositor<br>EDS |
| $R_{merge}$ | (Not available) | Depositor |
| $R_{sym}$ | (Not available) | Depositor |
| $< I/\sigma(I) >$ [1] | 6.99 (at 1.50Å) | Xtriage |
| Refinement program | REFMAC 5.8.0267 | Depositor |
| R, $R_{free}$ | 0.176 , 0.212<br>0.184 , 0.218 | Depositor<br>DCC |
| $R_{free}$ test set | 986 reflections (5.12%) | wwPDB-VP |
| Wilson B-factor (Å$^2$) | 9.1 | Xtriage |
| Anisotropy | 0.009 | Xtriage |
| Bulk solvent $k_{sol}$(e/Å$^3$), $B_{sol}$(Å$^2$) | 0.36 , 38.1 | EDS |
| L-test for twinning[2] | $< \|L\| > = 0.48$, $< L^2 > = 0.31$ | Xtriage |
| Estimated twinning fraction | No twinning to report. | Xtriage |
| $F_o$,$F_c$ correlation | 0.95 | EDS |
| Total number of atoms | 1145 | wwPDB-VP |
| Average B, all atoms (Å$^2$) | 11.0 | wwPDB-VP |

Xtriage's analysis on translational NCS is as follows: *The largest off-origin peak in the Patterson function is 8.62% of the height of the origin peak. No significant pseudotranslation is detected.*

[1] Intensities estimated from amplitudes.

[2] Theoretical values of $< |L| >$, $< L^2 >$ for acentric reflections are 0.5, 0.333 respectively for untwinned datasets, and 0.375, 0.2 for perfectly twinned datasets.

For Manuscript Review

### 5 Model quality (i)

#### 5.1 Standard geometry (i)

Bond lengths and bond angles in the following residue types are not validated in this section: CL, NA, GOL

The Z score for a bond length (or angle) is the number of standard deviations the observed value is removed from the expected value. A bond length (or angle) with $|Z| > 5$ is considered an outlier worth inspection. RMSZ is the root-mean-square of all Z scores of the bond lengths (or angles).

| Mol | Chain | Bond lengths | | Bond angles | |
|---|---|---|---|---|---|
| | | RMSZ | #$\|Z\|$ >5 | RMSZ | #$\|Z\|$ >5 |
| 1 | A | 0.77 | 0/1033 | 0.92 | 0/1395 |

There are no bond length outliers.

There are no bond angle outliers.

There are no chirality outliers.

There are no planarity outliers.

#### 5.2 Too-close contacts (i)

In the following table, the Non-H and H(model) columns list the number of non-hydrogen atoms and hydrogen atoms in the chain respectively. The H(added) column lists the number of hydrogen atoms added and optimized by MolProbity. The Clashes column lists the number of clashes within the asymmetric unit, whereas Symm-Clashes lists symmetry-related clashes.

| Mol | Chain | Non-H | H(model) | H(added) | Clashes | Symm-Clashes |
|---|---|---|---|---|---|---|
| 1 | A | 1013 | 0 | 967 | 8 | 0 |
| 2 | A | 1 | 0 | 0 | 0 | 0 |
| 3 | A | 7 | 0 | 0 | 0 | 0 |
| 4 | A | 12 | 0 | 16 | 0 | 0 |
| 5 | A | 112 | 0 | 0 | 1 | 0 |
| All | All | 1145 | 0 | 983 | 8 | 0 |

The all-atom clashscore is defined as the number of clashes found per 1000 atoms (including hydrogen atoms). The all-atom clashscore for this structure is 4.

All (8) close contacts within the same asymmetric unit are listed below, sorted by their clash magnitude.

For Manuscript Review

| Atom-1 | Atom-2 | Interatomic distance (Å) | Clash overlap (Å) |
|---|---|---|---|
| 1:A:11:ALA:O | 1:A:15:HIS:HD2 | 1.88 | 0.56 |
| 1:A:45:ARG:NE | 1:A:68:ARG:NH2 | 2.56 | 0.54 |
| 1:A:15:HIS:HE1 | 5:A:360:HOH:O | 1.90 | 0.53 |
| 1:A:45:ARG:CZ | 1:A:68:ARG:NH2 | 2.80 | 0.44 |
| 1:A:63:TRP:CE2 | 1:A:98:ILE:HG12 | 2.54 | 0.42 |
| 1:A:58:ILE:HB | 1:A:83:LEU:HD13 | 2.01 | 0.42 |
| 1:A:17:LEU:HG | 1:A:92:VAL:HG13 | 2.01 | 0.42 |
| 1:A:111:TRP:CD1 | 1:A:115:CYS:HB2 | 2.56 | 0.41 |

There are no symmetry-related clashes.

#### 5.3 Torsion angles (i)

##### 5.3.1 Protein backbone (i)

In the following table, the Percentiles column shows the percent Ramachandran outliers of the chain as a percentile score with respect to all X-ray entries followed by that with respect to entries of similar resolution.

The Analysed column shows the number of residues for which the backbone conformation was analysed, and the total number of residues.

| Mol | Chain | Analysed | Favoured | Allowed | Outliers | Percentiles | |
|---|---|---|---|---|---|---|---|
| 1 | A | 129/129 (100%) | 128 (99%) | 1 (1%) | 0 | 100 | 100 |

There are no Ramachandran outliers to report.

##### 5.3.2 Protein sidechains (i)

In the following table, the Percentiles column shows the percent sidechain outliers of the chain as a percentile score with respect to all X-ray entries followed by that with respect to entries of similar resolution.

The Analysed column shows the number of residues for which the sidechain conformation was analysed, and the total number of residues.

| Mol | Chain | Analysed | Rotameric | Outliers | Percentiles | |
|---|---|---|---|---|---|---|
| 1 | A | 107/105 (102%) | 104 (97%) | 3 (3%) | 43 | 14 |

All (3) residues with a non-rotameric sidechain are listed below:

For Manuscript Review

| Mol | Chain | Res | Type |
|---|---|---|---|
| 1 | A | 19 | ASN |
| 1 | A | 55 | ILE |
| 1 | A | 97 | LYS |

Sometimes sidechains can be flipped to improve hydrogen bonding and reduce clashes. All (3) such sidechains are listed below:

| Mol | Chain | Res | Type |
|---|---|---|---|
| 1 | A | 15 | HIS |
| 1 | A | 19 | ASN |
| 1 | A | 65 | ASN |

###### 5.3.3 RNA (i)

There are no RNA molecules in this entry.

##### 5.4 Non-standard residues in protein, DNA, RNA chains (i)

There are no non-standard protein/DNA/RNA residues in this entry.

##### 5.5 Carbohydrates (i)

There are no monosaccharides in this entry.

##### 5.6 Ligand geometry (i)

Of 10 ligands modelled in this entry, 8 are monoatomic - leaving 2 for Mogul analysis.

In the following table, the Counts columns list the number of bonds (or angles) for which Mogul statistics could be retrieved, the number of bonds (or angles) that are observed in the model and the number of bonds (or angles) that are defined in the Chemical Component Dictionary. The Link column lists molecule types, if any, to which the group is linked. The Z score for a bond length (or angle) is the number of standard deviations the observed value is removed from the expected value. A bond length (or angle) with $|Z| > 2$ is considered an outlier worth inspection. RMSZ is the root-mean-square of all Z scores of the bond lengths (or angles).

| Mol | Type | Chain | Res | Link | Bond lengths | | | Bond angles | | |
|---|---|---|---|---|---|---|---|---|---|---|
| | | | | | Counts | RMSZ | $\#\|Z\| > 2$ | Counts | RMSZ | $\#\|Z\| > 2$ |
| 4 | GOL | A | 210 | - | 5,5,5 | 0.11 | 0 | 5,5,5 | 0.20 | 0 |
| 4 | GOL | A | 209 | - | 5,5,5 | 0.14 | 0 | 5,5,5 | 0.38 | 0 |

For Manuscript Review

In the following table, the Chirals column lists the number of chiral outliers, the number of chiral centers analysed, the number of these observed in the model and the number defined in the Chemical Component Dictionary. Similar counts are reported in the Torsion and Rings columns. '-' means no outliers of that kind were identified.

| Mol | Type | Chain | Res | Link | Chirals | Torsions | Rings |
|---|---|---|---|---|---|---|---|
| 4 | GOL | A | 210 | - | - | 0/4/4/4 | - |
| 4 | GOL | A | 209 | - | - | 0/4/4/4 | - |

There are no bond length outliers.

There are no bond angle outliers.

There are no chirality outliers.

There are no torsion outliers.

There are no ring outliers.

No monomer is involved in short contacts.

#### 5.7 Other polymers (i)

There are no such residues in this entry.

#### 5.8 Polymer linkage issues (i)

There are no chain breaks in this entry.

For Manuscript Review

### 6 Fit of model and data (i)

#### 6.1 Protein, DNA and RNA chains (i)

In the following table, the column labelled '#RSRZ> 2' contains the number (and percentage) of RSRZ outliers, followed by percent RSRZ outliers for the chain as percentile scores relative to all X-ray entries and entries of similar resolution. The OWAB column contains the minimum, median, $95^{th}$ percentile and maximum values of the occupancy-weighted average B-factor per residue. The column labelled 'Q< 0.9' lists the number of (and percentage) of residues with an average occupancy less than 0.9.

| Mol | Chain | Analysed | <RSRZ> | #RSRZ>2 | | | OWAB(Å$^2$) | Q<0.9 |
|---|---|---|---|---|---|---|---|---|
| 1 | A | 129/129 (100%) | -0.02 | 2 (1%) | 72 | 77 | 5, 9, 18, 24 | 0 |

All (2) RSRZ outliers are listed below:

| Mol | Chain | Res | Type | RSRZ |
|---|---|---|---|---|
| 1 | A | 129 | LEU | 3.6 |
| 1 | A | 62 | TRP | 2.3 |

#### 6.2 Non-standard residues in protein, DNA, RNA chains (i)

There are no non-standard protein/DNA/RNA residues in this entry.

#### 6.3 Carbohydrates (i)

There are no monosaccharides in this entry.

#### 6.4 Ligands (i)

In the following table, the Atoms column lists the number of modelled atoms in the group and the number defined in the chemical component dictionary. The B-factors column lists the minimum, median, $95^{th}$ percentile and maximum values of B factors of atoms in the group. The column labelled 'Q< 0.9' lists the number of atoms with occupancy less than 0.9.

| Mol | Type | Chain | Res | Atoms | RSCC | RSR | B-factors(Å$^2$) | Q<0.9 |
|---|---|---|---|---|---|---|---|---|
| 4 | GOL | A | 210 | 6/6 | 0.77 | 0.18 | 44,46,46,47 | 0 |
| 4 | GOL | A | 209 | 6/6 | 0.84 | 0.17 | 21,30,33,38 | 0 |
| 3 | CL | A | 208 | 1/1 | 0.90 | 0.16 | 31,31,31,31 | 0 |
| 3 | CL | A | 205 | 1/1 | 0.97 | 0.06 | 22,22,22,22 | 0 |
| 3 | CL | A | 204 | 1/1 | 0.97 | 0.08 | 15,15,15,15 | 0 |

*Continued on next page...*

For Manuscript Review

*Continued from previous page...*

| Mol | Type | Chain | Res | Atoms | RSCC | RSR | B-factors(Å$^2$) | Q<0.9 |
|---|---|---|---|---|---|---|---|---|
| 3 | CL | A | 207 | 1/1 | 0.98 | 0.07 | 19,19,19,19 | 0 |
| 3 | CL | A | 206 | 1/1 | 0.98 | 0.07 | 20,20,20,20 | 0 |
| 3 | CL | A | 202 | 1/1 | 0.99 | 0.03 | 11,11,11,11 | 0 |
| 2 | NA | A | 201 | 1/1 | 0.99 | 0.08 | 11,11,11,11 | 0 |
| 3 | CL | A | 203 | 1/1 | 1.00 | 0.03 | 13,13,13,13 | 0 |

#### 6.5 Other polymers (i)

There are no such residues in this entry.

For Manuscript Review

### Full wwPDB X-ray Structure Validation Report (i)

Jun 7, 2022 – 12:31 PM EDT

| | | |
|---|---|---|
| PDB ID | : | 8D5W |
| Title | : | Crystal structure of hen egg white lysozyme at 100 Kelvin |
| Deposited on | : | 2022-06-06 |
| Resolution | : | 1.50 Å(reported) |

**This wwPDB validation report is for manuscript review**

This is a Full wwPDB X-ray Structure Validation Report.

This report is produced by the wwPDB biocuration pipeline after annotation of the structure.

We welcome your comments at **
A user guide is available at
https://www.wwpdb.org/validation/2017/XrayValidationReportHelp
with specific help available everywhere you see the (i) symbol.

The following versions of software and data (see references (i)) were used in the production of this report:

| | | |
|---|---|---|
| MolProbity | : | 4.02b-467 |
| Mogul | : | 1.8.5 (274361), CSD as541be (2020) |
| Xtriage (Phenix) | : | 1.13 |
| EDS | : | 2.28 |
| Percentile statistics | : | 20191225.v01 (using entries in the PDB archive December 25th 2019) |
| Refmac | : | 5.8.0158 |
| CCP4 | : | 7.0.044 (Gargrove) |
| Ideal geometry (proteins) | : | Engh & Huber (2001) |
| Ideal geometry (DNA, RNA) | : | Parkinson et al. (1996) |
| Validation Pipeline (wwPDB-VP) | : | 2.28 |

For Manuscript Review

### 1 Overall quality at a glance (i)

The following experimental techniques were used to determine the structure:
*X-RAY DIFFRACTION*

The reported resolution of this entry is 1.50 Å.

Percentile scores (ranging between 0-100) for global validation metrics of the entry are shown in the following graphic. The table shows the number of entries on which the scores are based.

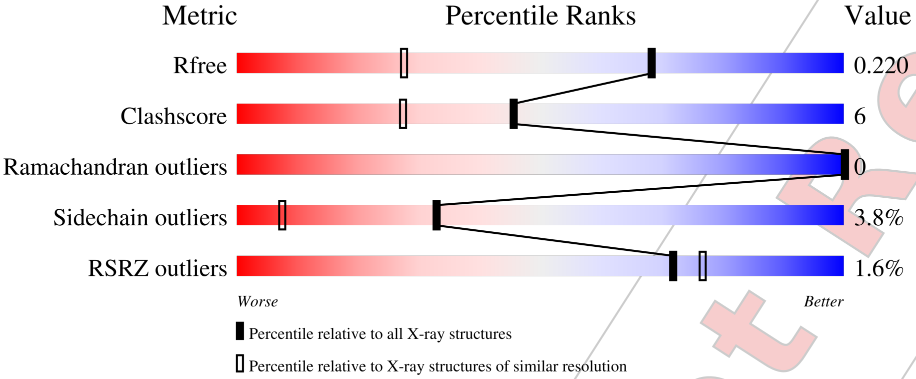


| Metric | Whole archive (#Entries) | Similar resolution (#Entries, resolution range(Å)) |
|---|---|---|
| $R_{free}$ | 130704 | 2936 (1.50-1.50) |
| Clashscore | 141614 | 3144 (1.50-1.50) |
| Ramachandran outliers | 138981 | 3066 (1.50-1.50) |
| Sidechain outliers | 138945 | 3064 (1.50-1.50) |
| RSRZ outliers | 127900 | 2884 (1.50-1.50) |

The table below summarises the geometric issues observed across the polymeric chains and their fit to the electron density. The red, orange, yellow and green segments of the lower bar indicate the fraction of residues that contain outliers for >=3, 2, 1 and 0 types of geometric quality criteria respectively. A grey segment represents the fraction of residues that are not modelled. The numeric value for each fraction is indicated below the corresponding segment, with a dot representing fractions <=5% The upper red bar (where present) indicates the fraction of residues that have poor fit to the electron density. The numeric value is given above the bar.

| Mol | Chain | Length | Quality of chain |
|---|---|---|---|
| 1 | A | 129 | 2% / 87% 11% • |

For Manuscript Review

#### 2 Entry composition (i)

There are 5 unique types of molecules in this entry. The entry contains 1144 atoms, of which 0 are hydrogens and 0 are deuteriums.

In the tables below, the ZeroOcc column contains the number of atoms modelled with zero occupancy, the AltConf column contains the number of residues with at least one atom in alternate conformation and the Trace column contains the number of residues modelled with at most 2 atoms.

- Molecule 1 is a protein called Lysozyme C.

| Mol | Chain | Residues | Atoms | | | | | ZeroOcc | AltConf | Trace |
|---|---|---|---|---|---|---|---|---|---|---|
| | | | Total | C | N | O | S | | | |
| 1 | A | 129 | 1013 | 619 | 195 | 189 | 10 | 0 | 2 | 0 |

- Molecule 2 is SODIUM ION (three-letter code: NA) (formula: Na).

| Mol | Chain | Residues | Atoms | | ZeroOcc | AltConf |
|---|---|---|---|---|---|---|
| | | | Total | Na | | |
| 2 | A | 1 | 1 | 1 | 0 | 0 |

- Molecule 3 is CHLORIDE ION (three-letter code: CL) (formula: Cl).

| Mol | Chain | Residues | Atoms | | ZeroOcc | AltConf |
|---|---|---|---|---|---|---|
| | | | Total | Cl | | |
| 3 | A | 7 | 7 | 7 | 0 | 0 |

- Molecule 4 is GLYCEROL (three-letter code: GOL) (formula: $C_3H_8O_3$).

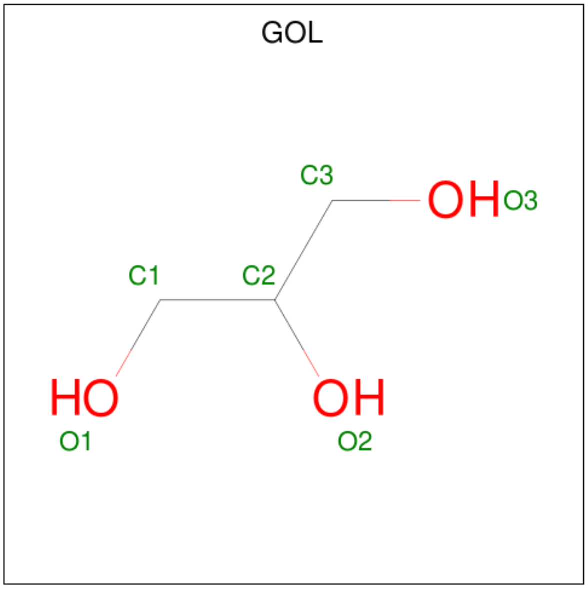


For Manuscript Review

| Mol | Chain | Residues | Atoms | ZeroOcc | AltConf |
|---|---|---|---|---|---|
| 4 | A | 1 | Total C O<br>6 3 3 | 0 | 0 |
| 4 | A | 1 | Total C O<br>6 3 3 | 0 | 0 |

- Molecule 5 is water.

| Mol | Chain | Residues | Atoms | ZeroOcc | AltConf |
|---|---|---|---|---|---|
| 5 | A | 111 | Total O<br>111 111 | 0 | 0 |

For Manuscript Review

### 3 Residue-property plots (i)

These plots are drawn for all protein, RNA, DNA and oligosaccharide chains in the entry. The first graphic for a chain summarises the proportions of the various outlier classes displayed in the second graphic. The second graphic shows the sequence view annotated by issues in geometry and electron density. Residues are color-coded according to the number of geometric quality criteria for which they contain at least one outlier: green = 0, yellow = 1, orange = 2 and red = 3 or more. A red dot above a residue indicates a poor fit to the electron density (RSRZ > 2). Stretches of 2 or more consecutive residues without any outlier are shown as a green connector. Residues present in the sample, but not in the model, are shown in grey.

- Molecule 1: Lysozyme C

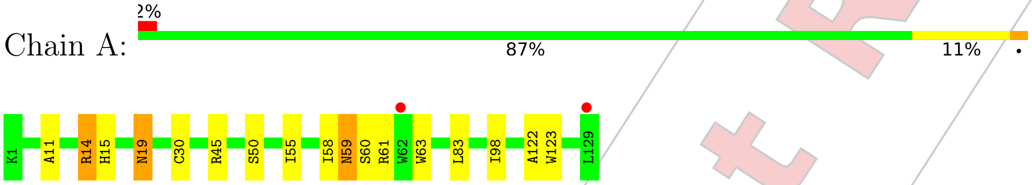


For Manuscript Review

### 4 Data and refinement statistics ⓘ

| Property | Value | Source |
|---|---|---|
| Space group | P 43 21 2 | Depositor |
| Cell constants<br>a, b, c, $\alpha$, $\beta$, $\gamma$ | 78.78Å 78.78Å 37.17Å<br>90.00° 90.00° 90.00° | Depositor |
| Resolution (Å) | 22.30 – 1.50<br>22.29 – 1.50 | Depositor<br>EDS |
| % Data completeness<br>(in resolution range) | 99.6 (22.30-1.50)<br>99.7 (22.29-1.50) | Depositor<br>EDS |
| $R_{merge}$ | (Not available) | Depositor |
| $R_{sym}$ | (Not available) | Depositor |
| $< I/\sigma(I) >$ [1] | 6.33 (at 1.50Å) | Xtriage |
| Refinement program | REFMAC 5.8.0267 | Depositor |
| R, $R_{free}$ | 0.185 , 0.214<br>0.193 , 0.220 | Depositor<br>DCC |
| $R_{free}$ test set | 987 reflections (5.12%) | wwPDB-VP |
| Wilson B-factor (Å$^2$) | 8.6 | Xtriage |
| Anisotropy | 0.069 | Xtriage |
| Bulk solvent $k_{sol}$(e/Å$^3$), $B_{sol}$(Å$^2$) | 0.34 , 37.0 | EDS |
| L-test for twinning[2] | $< \|L\| > = 0.48$, $< L^2 > = 0.31$ | Xtriage |
| Estimated twinning fraction | No twinning to report. | Xtriage |
| $F_o$,$F_c$ correlation | 0.94 | EDS |
| Total number of atoms | 1144 | wwPDB-VP |
| Average B, all atoms (Å$^2$) | 11.0 | wwPDB-VP |

Xtriage's analysis on translational NCS is as follows: *The largest off-origin peak in the Patterson function is 8.81% of the height of the origin peak. No significant pseudotranslation is detected.*

[1] Intensities estimated from amplitudes.

[2] Theoretical values of $< |L| >$, $< L^2 >$ for acentric reflections are 0.5, 0.333 respectively for untwinned datasets, and 0.375, 0.2 for perfectly twinned datasets.

For Manuscript Review

### 5 Model quality (i)

#### 5.1 Standard geometry (i)

Bond lengths and bond angles in the following residue types are not validated in this section: CL, GOL, NA

The Z score for a bond length (or angle) is the number of standard deviations the observed value is removed from the expected value. A bond length (or angle) with $|Z| > 5$ is considered an outlier worth inspection. RMSZ is the root-mean-square of all Z scores of the bond lengths (or angles).

| Mol | Chain | Bond lengths | | Bond angles | |
|---|---|---|---|---|---|
| | | RMSZ | #$\|Z\|$ >5 | RMSZ | #$\|Z\|$ >5 |
| 1 | A | 0.77 | 0/1033 | 0.95 | 2/1395 (0.1%) |

There are no bond length outliers.

All (2) bond angle outliers are listed below:

| Mol | Chain | Res | Type | Atoms | Z | Observed($^o$) | Ideal($^o$) |
|---|---|---|---|---|---|---|---|
| 1 | A | 45 | ARG | NE-CZ-NH1 | -6.41 | 117.09 | 120.30 |
| 1 | A | 45 | ARG | NE-CZ-NH2 | 5.85 | 123.22 | 120.30 |

There are no chirality outliers.

There are no planarity outliers.

#### 5.2 Too-close contacts (i)

In the following table, the Non-H and H(model) columns list the number of non-hydrogen atoms and hydrogen atoms in the chain respectively. The H(added) column lists the number of hydrogen atoms added and optimized by MolProbity. The Clashes column lists the number of clashes within the asymmetric unit, whereas Symm-Clashes lists symmetry-related clashes.

| Mol | Chain | Non-H | H(model) | H(added) | Clashes | Symm-Clashes |
|---|---|---|---|---|---|---|
| 1 | A | 1013 | 0 | 967 | 13 | 0 |
| 2 | A | 1 | 0 | 0 | 0 | 0 |
| 3 | A | 7 | 0 | 0 | 0 | 0 |
| 4 | A | 12 | 0 | 16 | 1 | 0 |
| 5 | A | 111 | 0 | 0 | 1 | 0 |
| All | All | 1144 | 0 | 983 | 13 | 0 |

The all-atom clashscore is defined as the number of clashes found per 1000 atoms (including hydrogen atoms). The all-atom clashscore for this structure is 6.

For Manuscript Review

All (13) close contacts within the same asymmetric unit are listed below, sorted by their clash magnitude.

| Atom-1 | Atom-2 | Interatomic distance (Å) | Clash overlap (Å) |
|---|---|---|---|
| 1:A:15:HIS:HE1 | 5:A:380:HOH:O | 1.85 | 0.59 |
| 1:A:11:ALA:O | 1:A:15:HIS:HD2 | 1.85 | 0.58 |
| 1:A:59:ASN:HD22 | 1:A:60:SER:N | 2.04 | 0.55 |
| 1:A:50:SER:HB2 | 1:A:59:ASN:HD21 | 1.73 | 0.53 |
| 1:A:50:SER:CB | 1:A:59:ASN:HD21 | 2.23 | 0.50 |
| 1:A:14:ARG:HH11 | 1:A:14:ARG:CG | 2.29 | 0.45 |
| 1:A:122:ALA:HB1 | 4:A:209:GOL:H32 | 1.99 | 0.43 |
| 1:A:19:ASN:HD22 | 1:A:19:ASN:HA | 1.61 | 0.43 |
| 1:A:50:SER:HB2 | 1:A:59:ASN:ND2 | 2.34 | 0.43 |
| 1:A:63:TRP:CE2 | 1:A:98:ILE:HG12 | 2.56 | 0.40 |
| 1:A:30:CYS:HB2 | 1:A:123:TRP:CD1 | 2.56 | 0.40 |
| 1:A:58:ILE:HB | 1:A:83:LEU:HD13 | 2.02 | 0.40 |
| 1:A:59:ASN:ND2 | 1:A:61:ARG:H | 2.19 | 0.40 |

There are no symmetry-related clashes.

#### 5.3 Torsion angles (i)

##### 5.3.1 Protein backbone (i)

In the following table, the Percentiles column shows the percent Ramachandran outliers of the chain as a percentile score with respect to all X-ray entries followed by that with respect to entries of similar resolution.

The Analysed column shows the number of residues for which the backbone conformation was analysed, and the total number of residues.

| Mol | Chain | Analysed | Favoured | Allowed | Outliers | Percentiles | |
|---|---|---|---|---|---|---|---|
| 1 | A | 129/129 (100%) | 128 (99%) | 1 (1%) | 0 | 100 | 100 |

There are no Ramachandran outliers to report.

##### 5.3.2 Protein sidechains (i)

In the following table, the Percentiles column shows the percent sidechain outliers of the chain as a percentile score with respect to all X-ray entries followed by that with respect to entries of similar resolution.

The Analysed column shows the number of residues for which the sidechain conformation was

For Manuscript Review

analysed, and the total number of residues.

| Mol | Chain | Analysed | Rotameric | Outliers | Percentiles | |
|---|---|---|---|---|---|---|
| 1 | A | 107/105 (102%) | 103 (96%) | 4 (4%) | 34 | 8 |

All (4) residues with a non-rotameric sidechain are listed below:

| Mol | Chain | Res | Type |
|---|---|---|---|
| 1 | A | 14 | ARG |
| 1 | A | 19 | ASN |
| 1 | A | 55 | ILE |
| 1 | A | 59 | ASN |

Sometimes sidechains can be flipped to improve hydrogen bonding and reduce clashes. All (4) such sidechains are listed below:

| Mol | Chain | Res | Type |
|---|---|---|---|
| 1 | A | 15 | HIS |
| 1 | A | 19 | ASN |
| 1 | A | 59 | ASN |
| 1 | A | 65 | ASN |

##### 5.3.3 RNA (i)

There are no RNA molecules in this entry.

#### 5.4 Non-standard residues in protein, DNA, RNA chains (i)

There are no non-standard protein/DNA/RNA residues in this entry.

#### 5.5 Carbohydrates (i)

There are no monosaccharides in this entry.

#### 5.6 Ligand geometry (i)

Of 10 ligands modelled in this entry, 8 are monoatomic - leaving 2 for Mogul analysis.

In the following table, the Counts columns list the number of bonds (or angles) for which Mogul statistics could be retrieved, the number of bonds (or angles) that are observed in the model and the number of bonds (or angles) that are defined in the Chemical Component Dictionary. The Link column lists molecule types, if any, to which the group is linked. The Z score for a bond

For Manuscript Review

length (or angle) is the number of standard deviations the observed value is removed from the expected value. A bond length (or angle) with $|Z| > 2$ is considered an outlier worth inspection. RMSZ is the root-mean-square of all Z scores of the bond lengths (or angles).

| Mol | Type | Chain | Res | Link | Bond lengths | | | Bond angles | | |
|---|---|---|---|---|---|---|---|---|---|---|
| | | | | | Counts | RMSZ | $\#\|Z\|>2$ | Counts | RMSZ | $\#\|Z\|>2$ |
| 4 | GOL | A | 209 | - | 5,5,5 | 0.13 | 0 | 5,5,5 | 0.40 | 0 |
| 4 | GOL | A | 210 | - | 5,5,5 | 0.10 | 0 | 5,5,5 | 0.18 | 0 |

In the following table, the Chirals column lists the number of chiral outliers, the number of chiral centers analysed, the number of these observed in the model and the number defined in the Chemical Component Dictionary. Similar counts are reported in the Torsion and Rings columns. '-' means no outliers of that kind were identified.

| Mol | Type | Chain | Res | Link | Chirals | Torsions | Rings |
|---|---|---|---|---|---|---|---|
| 4 | GOL | A | 209 | - | - | 0/4/4/4 | - |
| 4 | GOL | A | 210 | - | - | 0/4/4/4 | - |

There are no bond length outliers.

There are no bond angle outliers.

There are no chirality outliers.

There are no torsion outliers.

There are no ring outliers.

1 monomer is involved in 1 short contact:

| Mol | Chain | Res | Type | Clashes | Symm-Clashes |
|---|---|---|---|---|---|
| 4 | A | 209 | GOL | 1 | 0 |

#### 5.7 Other polymers (i)

There are no such residues in this entry.

#### 5.8 Polymer linkage issues (i)

There are no chain breaks in this entry.

For Manuscript Review

### 6 Fit of model and data (i)

#### 6.1 Protein, DNA and RNA chains (i)

In the following table, the column labelled '#RSRZ> 2' contains the number (and percentage) of RSRZ outliers, followed by percent RSRZ outliers for the chain as percentile scores relative to all X-ray entries and entries of similar resolution. The OWAB column contains the minimum, median, 95$^{th}$ percentile and maximum values of the occupancy-weighted average B-factor per residue. The column labelled 'Q< 0.9' lists the number of (and percentage) of residues with an average occupancy less than 0.9.

| Mol | Chain | Analysed | <RSRZ> | #RSRZ>2 | | | OWAB(Å$^2$) | Q<0.9 |
|---|---|---|---|---|---|---|---|---|
| 1 | A | 129/129 (100%) | -0.03 | 2 (1%) | 72 | 77 | 5, 9, 19, 24 | 0 |

All (2) RSRZ outliers are listed below:

| Mol | Chain | Res | Type | RSRZ |
|---|---|---|---|---|
| 1 | A | 129 | LEU | 4.2 |
| 1 | A | 62 | TRP | 2.7 |

#### 6.2 Non-standard residues in protein, DNA, RNA chains (i)

There are no non-standard protein/DNA/RNA residues in this entry.

#### 6.3 Carbohydrates (i)

There are no monosaccharides in this entry.

#### 6.4 Ligands (i)

In the following table, the Atoms column lists the number of modelled atoms in the group and the number defined in the chemical component dictionary. The B-factors column lists the minimum, median, 95$^{th}$ percentile and maximum values of B factors of atoms in the group. The column labelled 'Q< 0.9' lists the number of atoms with occupancy less than 0.9.

| Mol | Type | Chain | Res | Atoms | RSCC | RSR | B-factors(Å$^2$) | Q<0.9 |
|---|---|---|---|---|---|---|---|---|
| 4 | GOL | A | 210 | 6/6 | 0.69 | 0.25 | 47,49,50,50 | 0 |
| 4 | GOL | A | 209 | 6/6 | 0.85 | 0.24 | 19,32,35,40 | 0 |
| 3 | CL | A | 208 | 1/1 | 0.93 | 0.25 | 35,35,35,35 | 0 |
| 3 | CL | A | 205 | 1/1 | 0.96 | 0.06 | 22,22,22,22 | 0 |
| 3 | CL | A | 207 | 1/1 | 0.97 | 0.07 | 18,18,18,18 | 0 |

*Continued on next page...*

For Manuscript Review

*Continued from previous page...*

| Mol | Type | Chain | Res | Atoms | RSCC | RSR | B-factors($Å^2$) | Q<0.9 |
|---|---|---|---|---|---|---|---|---|
| 3 | CL | A | 204 | 1/1 | 0.97 | 0.06 | 15,15,15,15 | 0 |
| 3 | CL | A | 206 | 1/1 | 0.98 | 0.09 | 22,22,22,22 | 0 |
| 3 | CL | A | 202 | 1/1 | 0.99 | 0.03 | 11,11,11,11 | 0 |
| 3 | CL | A | 203 | 1/1 | 0.99 | 0.04 | 14,14,14,14 | 0 |
| 2 | NA | A | 201 | 1/1 | 0.99 | 0.14 | 12,12,12,12 | 0 |

#### 6.5 Other polymers (i)

There are no such residues in this entry.

For Manuscript Review

### Full wwPDB X-ray Structure Validation Report (i)

Jun 7, 2022 – 01:50 PM EDT

| | | |
|---|---|---|
| PDB ID | : | 8D5X |
| Title | : | Crystal structure of hen egg white lysozyme at 125 Kelvin (Duplicate) |
| Deposited on | : | 2022-06-06 |
| Resolution | : | 1.50 Å(reported) |

**This wwPDB validation report is for manuscript review**

This is a Full wwPDB X-ray Structure Validation Report.

This report is produced by the wwPDB biocuration pipeline after annotation of the structure.

We welcome your comments at **
A user guide is available at
https://www.wwpdb.org/validation/2017/XrayValidationReportHelp
with specific help available everywhere you see the (i) symbol.

The following versions of software and data (see references (i)) were used in the production of this report:

| | | |
|---|---|---|
| MolProbity | : | 4.02b-467 |
| Mogul | : | 1.8.5 (274361), CSD as541be (2020) |
| Xtriage (Phenix) | : | 1.13 |
| EDS | : | 2.28 |
| Percentile statistics | : | 20191225.v01 (using entries in the PDB archive December 25th 2019) |
| Refmac | : | 5.8.0158 |
| CCP4 | : | 7.0.044 (Gargrove) |
| Ideal geometry (proteins) | : | Engh & Huber (2001) |
| Ideal geometry (DNA, RNA) | : | Parkinson et al. (1996) |
| Validation Pipeline (wwPDB-VP) | : | 2.28 |

For Manuscript Review

### 1 Overall quality at a glance (i)

The following experimental techniques were used to determine the structure:
*X-RAY DIFFRACTION*

The reported resolution of this entry is 1.50 Å.

Percentile scores (ranging between 0-100) for global validation metrics of the entry are shown in the following graphic. The table shows the number of entries on which the scores are based.

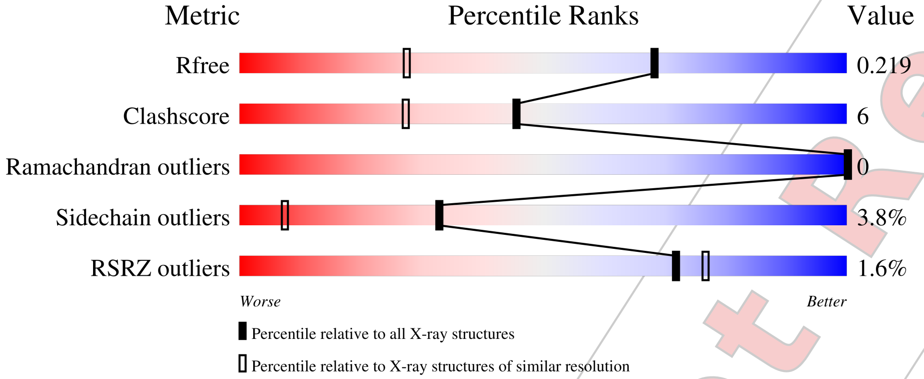


| Metric | Whole archive (#Entries) | Similar resolution (#Entries, resolution range(Å)) |
|---|---|---|
| $R_{free}$ | 130704 | 2936 (1.50-1.50) |
| Clashscore | 141614 | 3144 (1.50-1.50) |
| Ramachandran outliers | 138981 | 3066 (1.50-1.50) |
| Sidechain outliers | 138945 | 3064 (1.50-1.50) |
| RSRZ outliers | 127900 | 2884 (1.50-1.50) |

The table below summarises the geometric issues observed across the polymeric chains and their fit to the electron density. The red, orange, yellow and green segments of the lower bar indicate the fraction of residues that contain outliers for >=3, 2, 1 and 0 types of geometric quality criteria respectively. A grey segment represents the fraction of residues that are not modelled. The numeric value for each fraction is indicated below the corresponding segment, with a dot representing fractions <=5% The upper red bar (where present) indicates the fraction of residues that have poor fit to the electron density. The numeric value is given above the bar.

| Mol | Chain | Length | Quality of chain |
|---|---|---|---|
| 1 | A | 129 | 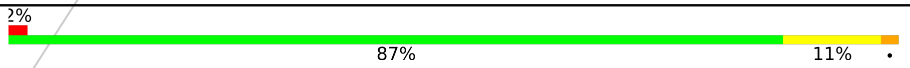  |

For Manuscript Review

#### 2 Entry composition (i)

There are 5 unique types of molecules in this entry. The entry contains 1144 atoms, of which 0 are hydrogens and 0 are deuteriums.

In the tables below, the ZeroOcc column contains the number of atoms modelled with zero occupancy, the AltConf column contains the number of residues with at least one atom in alternate conformation and the Trace column contains the number of residues modelled with at most 2 atoms.

- Molecule 1 is a protein called Lysozyme C.

| Mol | Chain | Residues | Atoms | | | | | ZeroOcc | AltConf | Trace |
|---|---|---|---|---|---|---|---|---|---|---|
| | | | Total | C | N | O | S | | | |
| 1 | A | 129 | 1013 | 619 | 195 | 189 | 10 | 0 | 2 | 0 |

- Molecule 2 is SODIUM ION (three-letter code: NA) (formula: Na).

| Mol | Chain | Residues | Atoms | | ZeroOcc | AltConf |
|---|---|---|---|---|---|---|
| | | | Total | Na | | |
| 2 | A | 1 | 1 | 1 | 0 | 0 |

- Molecule 3 is CHLORIDE ION (three-letter code: CL) (formula: Cl).

| Mol | Chain | Residues | Atoms | | ZeroOcc | AltConf |
|---|---|---|---|---|---|---|
| | | | Total | Cl | | |
| 3 | A | 7 | 7 | 7 | 0 | 0 |

- Molecule 4 is GLYCEROL (three-letter code: GOL) (formula: $C_3H_8O_3$).

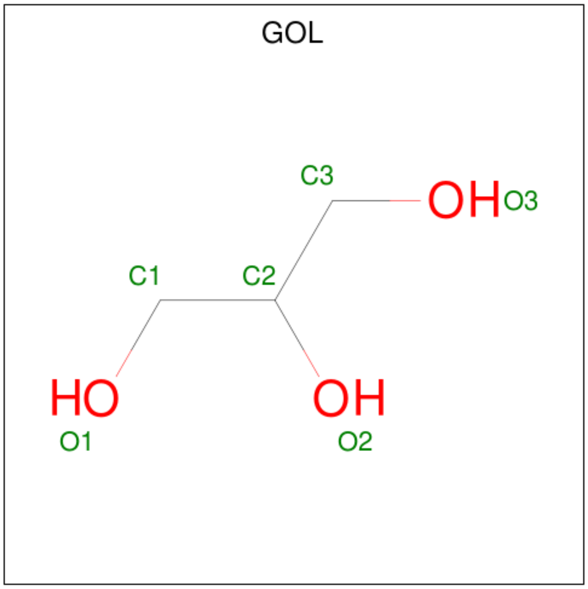


For Manuscript Review

| Mol | Chain | Residues | Atoms | ZeroOcc | AltConf |
|---|---|---|---|---|---|
| 4 | A | 1 | Total C O<br>6 3 3 | 0 | 0 |
| 4 | A | 1 | Total C O<br>6 3 3 | 0 | 0 |

- Molecule 5 is water.

| Mol | Chain | Residues | Atoms | ZeroOcc | AltConf |
|---|---|---|---|---|---|
| 5 | A | 111 | Total O<br>111 111 | 0 | 0 |

For Manuscript Review

### 3 Residue-property plots (i)

These plots are drawn for all protein, RNA, DNA and oligosaccharide chains in the entry. The first graphic for a chain summarises the proportions of the various outlier classes displayed in the second graphic. The second graphic shows the sequence view annotated by issues in geometry and electron density. Residues are color-coded according to the number of geometric quality criteria for which they contain at least one outlier: green = 0, yellow = 1, orange = 2 and red = 3 or more. A red dot above a residue indicates a poor fit to the electron density (RSRZ > 2). Stretches of 2 or more consecutive residues without any outlier are shown as a green connector. Residues present in the sample, but not in the model, are shown in grey.

• Molecule 1: Lysozyme C

Chain A:

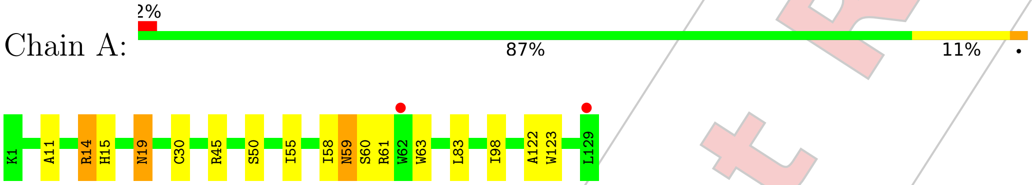


For Manuscript Review

### 4 Data and refinement statistics (i)

| Property | Value | Source |
|---|---|---|
| Space group | P 43 21 2 | Depositor |
| Cell constants<br>a, b, c, $\alpha$, $\beta$, $\gamma$ | 78.78Å 78.78Å 37.17Å<br>90.00° 90.00° 90.00° | Depositor |
| Resolution (Å) | 22.27 – 1.50<br>22.29 – 1.50 | Depositor<br>EDS |
| % Data completeness<br>(in resolution range) | 99.6 (22.27-1.50)<br>99.7 (22.29-1.50) | Depositor<br>EDS |
| $R_{merge}$ | (Not available) | Depositor |
| $R_{sym}$ | (Not available) | Depositor |
| $< I/\sigma(I) >$ [1] | 6.33 (at 1.50Å) | Xtriage |
| Refinement program | REFMAC 5.8.0267 | Depositor |
| R, $R_{free}$ | 0.185 , 0.214<br>0.193 , 0.219 | Depositor<br>DCC |
| $R_{free}$ test set | 987 reflections (5.12%) | wwPDB-VP |
| Wilson B-factor (Å$^2$) | 8.6 | Xtriage |
| Anisotropy | 0.069 | Xtriage |
| Bulk solvent $k_{sol}$(e/Å$^3$), $B_{sol}$(Å$^2$) | 0.34 , 37.0 | EDS |
| L-test for twinning[2] | $< \|L\| > = 0.48$, $< L^2 > = 0.31$ | Xtriage |
| Estimated twinning fraction | No twinning to report. | Xtriage |
| $F_o$,$F_c$ correlation | 0.94 | EDS |
| Total number of atoms | 1144 | wwPDB-VP |
| Average B, all atoms (Å$^2$) | 11.0 | wwPDB-VP |

Xtriage's analysis on translational NCS is as follows: *The largest off-origin peak in the Patterson function is 8.81% of the height of the origin peak. No significant pseudotranslation is detected.*

[1] Intensities estimated from amplitudes.
[2] Theoretical values of $< |L| >$, $< L^2 >$ for acentric reflections are 0.5, 0.333 respectively for untwinned datasets, and 0.375, 0.2 for perfectly twinned datasets.

For Manuscript Review

### 5 Model quality (i)

#### 5.1 Standard geometry (i)

Bond lengths and bond angles in the following residue types are not validated in this section: CL, GOL, NA

The Z score for a bond length (or angle) is the number of standard deviations the observed value is removed from the expected value. A bond length (or angle) with $|Z| > 5$ is considered an outlier worth inspection. RMSZ is the root-mean-square of all Z scores of the bond lengths (or angles).

| Mol | Chain | Bond lengths | | Bond angles | |
|---|---|---|---|---|---|
| | | RMSZ | #\|Z\| >5 | RMSZ | #\|Z\| >5 |
| 1 | A | 0.77 | 0/1033 | 0.95 | 2/1395 (0.1%) |

There are no bond length outliers.

All (2) bond angle outliers are listed below:

| Mol | Chain | Res | Type | Atoms | Z | Observed($^o$) | Ideal($^o$) |
|---|---|---|---|---|---|---|---|
| 1 | A | 45 | ARG | NE-CZ-NH1 | -6.41 | 117.09 | 120.30 |
| 1 | A | 45 | ARG | NE-CZ-NH2 | 5.85 | 123.22 | 120.30 |

There are no chirality outliers.

There are no planarity outliers.

#### 5.2 Too-close contacts (i)

In the following table, the Non-H and H(model) columns list the number of non-hydrogen atoms and hydrogen atoms in the chain respectively. The H(added) column lists the number of hydrogen atoms added and optimized by MolProbity. The Clashes column lists the number of clashes within the asymmetric unit, whereas Symm-Clashes lists symmetry-related clashes.

| Mol | Chain | Non-H | H(model) | H(added) | Clashes | Symm-Clashes |
|---|---|---|---|---|---|---|
| 1 | A | 1013 | 0 | 967 | 13 | 0 |
| 2 | A | 1 | 0 | 0 | 0 | 0 |
| 3 | A | 7 | 0 | 0 | 0 | 0 |
| 4 | A | 12 | 0 | 16 | 1 | 0 |
| 5 | A | 111 | 0 | 0 | 1 | 0 |
| All | All | 1144 | 0 | 983 | 13 | 0 |

The all-atom clashscore is defined as the number of clashes found per 1000 atoms (including hydrogen atoms). The all-atom clashscore for this structure is 6.

For Manuscript Review

All (13) close contacts within the same asymmetric unit are listed below, sorted by their clash magnitude.

| Atom-1 | Atom-2 | Interatomic distance (Å) | Clash overlap (Å) |
|---|---|---|---|
| 1:A:15:HIS:HE1 | 5:A:380:HOH:O | 1.85 | 0.59 |
| 1:A:11:ALA:O | 1:A:15:HIS:HD2 | 1.85 | 0.58 |
| 1:A:59:ASN:HD22 | 1:A:60:SER:N | 2.04 | 0.55 |
| 1:A:50:SER:HB2 | 1:A:59:ASN:HD21 | 1.73 | 0.53 |
| 1:A:50:SER:CB | 1:A:59:ASN:HD21 | 2.23 | 0.50 |
| 1:A:14:ARG:HH11 | 1:A:14:ARG:CG | 2.29 | 0.45 |
| 1:A:122:ALA:HB1 | 4:A:209:GOL:H32 | 1.99 | 0.43 |
| 1:A:19:ASN:HD22 | 1:A:19:ASN:HA | 1.61 | 0.43 |
| 1:A:50:SER:HB2 | 1:A:59:ASN:ND2 | 2.34 | 0.43 |
| 1:A:63:TRP:CE2 | 1:A:98:ILE:HG12 | 2.56 | 0.40 |
| 1:A:30:CYS:HB2 | 1:A:123:TRP:CD1 | 2.56 | 0.40 |
| 1:A:58:ILE:HB | 1:A:83:LEU:HD13 | 2.02 | 0.40 |
| 1:A:59:ASN:ND2 | 1:A:61:ARG:H | 2.19 | 0.40 |

There are no symmetry-related clashes.

#### 5.3 Torsion angles (i)

##### 5.3.1 Protein backbone (i)

In the following table, the Percentiles column shows the percent Ramachandran outliers of the chain as a percentile score with respect to all X-ray entries followed by that with respect to entries of similar resolution.

The Analysed column shows the number of residues for which the backbone conformation was analysed, and the total number of residues.

| Mol | Chain | Analysed | Favoured | Allowed | Outliers | Percentiles | |
|---|---|---|---|---|---|---|---|
| 1 | A | 129/129 (100%) | 128 (99%) | 1 (1%) | 0 | 100 | 100 |

There are no Ramachandran outliers to report.

##### 5.3.2 Protein sidechains (i)

In the following table, the Percentiles column shows the percent sidechain outliers of the chain as a percentile score with respect to all X-ray entries followed by that with respect to entries of similar resolution.

The Analysed column shows the number of residues for which the sidechain conformation was

For Manuscript Review

analysed, and the total number of residues.

| Mol | Chain | Analysed | Rotameric | Outliers | Percentiles | |
|---|---|---|---|---|---|---|
| 1 | A | 107/105 (102%) | 103 (96%) | 4 (4%) | 34 | 8 |

All (4) residues with a non-rotameric sidechain are listed below:

| Mol | Chain | Res | Type |
|---|---|---|---|
| 1 | A | 14 | ARG |
| 1 | A | 19 | ASN |
| 1 | A | 55 | ILE |
| 1 | A | 59 | ASN |

Sometimes sidechains can be flipped to improve hydrogen bonding and reduce clashes. All (4) such sidechains are listed below:

| Mol | Chain | Res | Type |
|---|---|---|---|
| 1 | A | 15 | HIS |
| 1 | A | 19 | ASN |
| 1 | A | 59 | ASN |
| 1 | A | 65 | ASN |

##### 5.3.3 RNA (i)

There are no RNA molecules in this entry.

#### 5.4 Non-standard residues in protein, DNA, RNA chains (i)

There are no non-standard protein/DNA/RNA residues in this entry.

#### 5.5 Carbohydrates (i)

There are no monosaccharides in this entry.

#### 5.6 Ligand geometry (i)

Of 10 ligands modelled in this entry, 8 are monoatomic - leaving 2 for Mogul analysis.

In the following table, the Counts columns list the number of bonds (or angles) for which Mogul statistics could be retrieved, the number of bonds (or angles) that are observed in the model and the number of bonds (or angles) that are defined in the Chemical Component Dictionary. The Link column lists molecule types, if any, to which the group is linked. The Z score for a bond

For Manuscript Review

length (or angle) is the number of standard deviations the observed value is removed from the expected value. A bond length (or angle) with $|Z| > 2$ is considered an outlier worth inspection. RMSZ is the root-mean-square of all Z scores of the bond lengths (or angles).

| Mol | Type | Chain | Res | Link | Bond lengths | | | Bond angles | | |
|---|---|---|---|---|---|---|---|---|---|---|
| | | | | | Counts | RMSZ | $\#\|Z\| > 2$ | Counts | RMSZ | $\#\|Z\| > 2$ |
| 4 | GOL | A | 209 | - | 5,5,5 | 0.13 | 0 | 5,5,5 | 0.40 | 0 |
| 4 | GOL | A | 210 | - | 5,5,5 | 0.10 | 0 | 5,5,5 | 0.18 | 0 |

In the following table, the Chirals column lists the number of chiral outliers, the number of chiral centers analysed, the number of these observed in the model and the number defined in the Chemical Component Dictionary. Similar counts are reported in the Torsion and Rings columns. '-' means no outliers of that kind were identified.

| Mol | Type | Chain | Res | Link | Chirals | Torsions | Rings |
|---|---|---|---|---|---|---|---|
| 4 | GOL | A | 209 | - | - | 0/4/4/4 | - |
| 4 | GOL | A | 210 | - | - | 0/4/4/4 | - |

There are no bond length outliers.

There are no bond angle outliers.

There are no chirality outliers.

There are no torsion outliers.

There are no ring outliers.

1 monomer is involved in 1 short contact:

| Mol | Chain | Res | Type | Clashes | Symm-Clashes |
|---|---|---|---|---|---|
| 4 | A | 209 | GOL | 1 | 0 |

#### 5.7 Other polymers (i)

There are no such residues in this entry.

#### 5.8 Polymer linkage issues (i)

There are no chain breaks in this entry.

For Manuscript Review

### 6 Fit of model and data (i)

#### 6.1 Protein, DNA and RNA chains (i)

In the following table, the column labelled '#RSRZ> 2' contains the number (and percentage) of RSRZ outliers, followed by percent RSRZ outliers for the chain as percentile scores relative to all X-ray entries and entries of similar resolution. The OWAB column contains the minimum, median, $95^{th}$ percentile and maximum values of the occupancy-weighted average B-factor per residue. The column labelled 'Q< 0.9' lists the number of (and percentage) of residues with an average occupancy less than 0.9.

| Mol | Chain | Analysed | <RSRZ> | #RSRZ>2 | | | OWAB(Å$^2$) | Q<0.9 |
|---|---|---|---|---|---|---|---|---|
| 1 | A | 129/129 (100%) | -0.03 | 2 (1%) | 72 | 77 | 5, 9, 19, 24 | 0 |

All (2) RSRZ outliers are listed below:

| Mol | Chain | Res | Type | RSRZ |
|---|---|---|---|---|
| 1 | A | 129 | LEU | 4.2 |
| 1 | A | 62 | TRP | 2.7 |

#### 6.2 Non-standard residues in protein, DNA, RNA chains (i)

There are no non-standard protein/DNA/RNA residues in this entry.

#### 6.3 Carbohydrates (i)

There are no monosaccharides in this entry.

#### 6.4 Ligands (i)

In the following table, the Atoms column lists the number of modelled atoms in the group and the number defined in the chemical component dictionary. The B-factors column lists the minimum, median, $95^{th}$ percentile and maximum values of B factors of atoms in the group. The column labelled 'Q< 0.9' lists the number of atoms with occupancy less than 0.9.

| Mol | Type | Chain | Res | Atoms | RSCC | RSR | B-factors(Å$^2$) | Q<0.9 |
|---|---|---|---|---|---|---|---|---|
| 4 | GOL | A | 210 | 6/6 | 0.69 | 0.25 | 47,49,50,50 | 0 |
| 4 | GOL | A | 209 | 6/6 | 0.85 | 0.24 | 19,32,35,40 | 0 |
| 3 | CL | A | 208 | 1/1 | 0.93 | 0.25 | 35,35,35,35 | 0 |
| 3 | CL | A | 205 | 1/1 | 0.96 | 0.06 | 22,22,22,22 | 0 |
| 3 | CL | A | 207 | 1/1 | 0.97 | 0.07 | 18,18,18,18 | 0 |

*Continued on next page...*

For Manuscript Review

*Continued from previous page...*

| Mol | Type | Chain | Res | Atoms | RSCC | RSR | B-factors($Å^2$) | Q<0.9 |
|---|---|---|---|---|---|---|---|---|
| 3 | CL | A | 204 | 1/1 | 0.97 | 0.06 | 15,15,15,15 | 0 |
| 3 | CL | A | 206 | 1/1 | 0.98 | 0.09 | 22,22,22,22 | 0 |
| 3 | CL | A | 202 | 1/1 | 0.99 | 0.03 | 11,11,11,11 | 0 |
| 3 | CL | A | 203 | 1/1 | 0.99 | 0.04 | 14,14,14,14 | 0 |
| 2 | NA | A | 201 | 1/1 | 0.99 | 0.14 | 12,12,12,12 | 0 |

#### 6.5 Other polymers (i)

There are no such residues in this entry.

For Manuscript Review

### Full wwPDB X-ray Structure Validation Report (i)

Jun 7, 2022 – 01:52 PM EDT

| | | |
|---|---|---|
| PDB ID | : | 8D5Z |
| Title | : | Crystal structure of hen egg white lysozyme at 125 Kelvin (Triplicate) |
| Deposited on | : | 2022-06-06 |
| Resolution | : | 1.50 Å(reported) |

**This wwPDB validation report is for manuscript review**

This is a Full wwPDB X-ray Structure Validation Report.

This report is produced by the wwPDB biocuration pipeline after annotation of the structure.

We welcome your comments at **
A user guide is available at
https://www.wwpdb.org/validation/2017/XrayValidationReportHelp
with specific help available everywhere you see the (i) symbol.

The following versions of software and data (see references (i)) were used in the production of this report:

| | | |
|---|---|---|
| MolProbity | : | 4.02b-467 |
| Mogul | : | 1.8.5 (274361), CSD as541be (2020) |
| Xtriage (Phenix) | : | 1.13 |
| EDS | : | 2.28 |
| Percentile statistics | : | 20191225.v01 (using entries in the PDB archive December 25th 2019) |
| Refmac | : | 5.8.0158 |
| CCP4 | : | 7.0.044 (Gargrove) |
| Ideal geometry (proteins) | : | Engh & Huber (2001) |
| Ideal geometry (DNA, RNA) | : | Parkinson et al. (1996) |
| Validation Pipeline (wwPDB-VP) | : | 2.28 |

For Manuscript Review

### 1 Overall quality at a glance (i)

The following experimental techniques were used to determine the structure:
*X-RAY DIFFRACTION*

The reported resolution of this entry is 1.50 Å.

Percentile scores (ranging between 0-100) for global validation metrics of the entry are shown in the following graphic. The table shows the number of entries on which the scores are based.

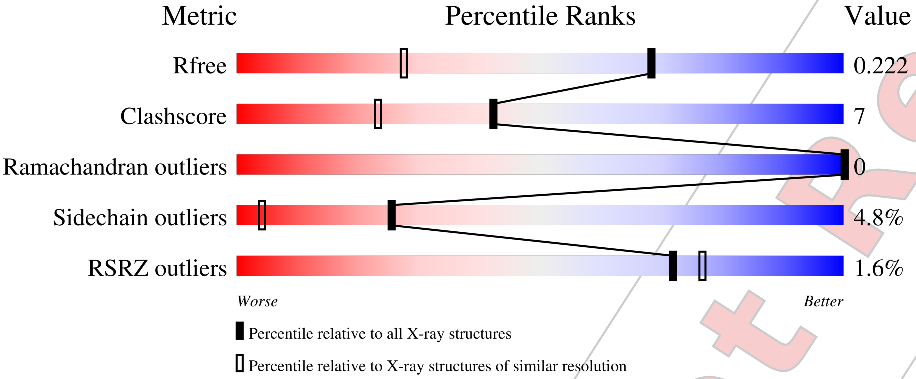


| Metric | Whole archive (#Entries) | Similar resolution (#Entries, resolution range(Å)) |
|---|---|---|
| $R_{free}$ | 130704 | 2936 (1.50-1.50) |
| Clashscore | 141614 | 3144 (1.50-1.50) |
| Ramachandran outliers | 138981 | 3066 (1.50-1.50) |
| Sidechain outliers | 138945 | 3064 (1.50-1.50) |
| RSRZ outliers | 127900 | 2884 (1.50-1.50) |

The table below summarises the geometric issues observed across the polymeric chains and their fit to the electron density. The red, orange, yellow and green segments of the lower bar indicate the fraction of residues that contain outliers for >=3, 2, 1 and 0 types of geometric quality criteria respectively. A grey segment represents the fraction of residues that are not modelled. The numeric value for each fraction is indicated below the corresponding segment, with a dot representing fractions <=5% The upper red bar (where present) indicates the fraction of residues that have poor fit to the electron density. The numeric value is given above the bar.

| Mol | Chain | Length | Quality of chain |
|---|---|---|---|
| 1 | A | 129 | 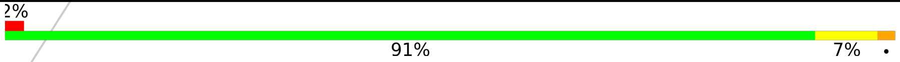 |

For Manuscript Review

#### 2 Entry composition (i)

There are 5 unique types of molecules in this entry. The entry contains 1143 atoms, of which 0 are hydrogens and 0 are deuteriums.

In the tables below, the ZeroOcc column contains the number of atoms modelled with zero occupancy, the AltConf column contains the number of residues with at least one atom in alternate conformation and the Trace column contains the number of residues modelled with at most 2 atoms.

- Molecule 1 is a protein called Lysozyme C.

| Mol | Chain | Residues | Atoms | | | | | ZeroOcc | AltConf | Trace |
|---|---|---|---|---|---|---|---|---|---|---|
| | | | Total | C | N | O | S | | | |
| 1 | A | 129 | 1012 | 619 | 197 | 186 | 10 | 0 | 1 | 0 |

- Molecule 2 is SODIUM ION (three-letter code: NA) (formula: Na).

| Mol | Chain | Residues | Atoms | | ZeroOcc | AltConf |
|---|---|---|---|---|---|---|
| | | | Total | Na | | |
| 2 | A | 1 | 1 | 1 | 0 | 0 |

- Molecule 3 is CHLORIDE ION (three-letter code: CL) (formula: Cl).

| Mol | Chain | Residues | Atoms | | ZeroOcc | AltConf |
|---|---|---|---|---|---|---|
| | | | Total | Cl | | |
| 3 | A | 7 | 7 | 7 | 0 | 0 |

- Molecule 4 is GLYCEROL (three-letter code: GOL) (formula: $C_3H_8O_3$).

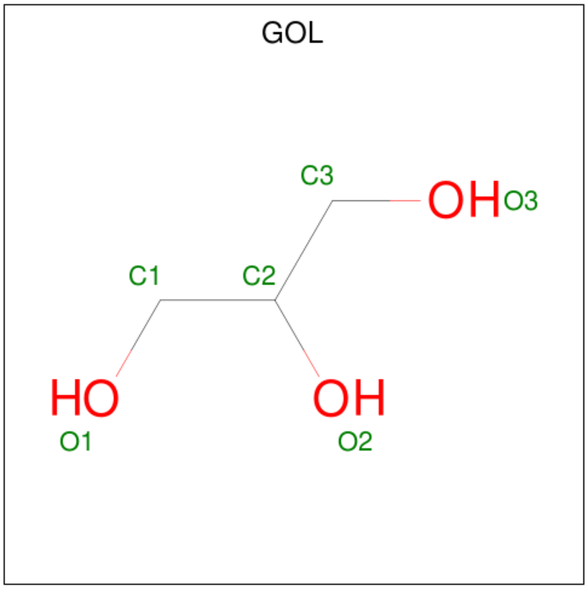


For Manuscript Review

| Mol | Chain | Residues | Atoms | ZeroOcc | AltConf |
|---|---|---|---|---|---|
| 4 | A | 1 | Total C O<br>6 3 3 | 0 | 0 |
| 4 | A | 1 | Total C O<br>6 3 3 | 0 | 0 |

- Molecule 5 is water.

| Mol | Chain | Residues | Atoms | ZeroOcc | AltConf |
|---|---|---|---|---|---|
| 5 | A | 111 | Total O<br>111 111 | 0 | 0 |

For Manuscript Review

### 3 Residue-property plots (i)

These plots are drawn for all protein, RNA, DNA and oligosaccharide chains in the entry. The first graphic for a chain summarises the proportions of the various outlier classes displayed in the second graphic. The second graphic shows the sequence view annotated by issues in geometry and electron density. Residues are color-coded according to the number of geometric quality criteria for which they contain at least one outlier: green = 0, yellow = 1, orange = 2 and red = 3 or more. A red dot above a residue indicates a poor fit to the electron density (RSRZ > 2). Stretches of 2 or more consecutive residues without any outlier are shown as a green connector. Residues present in the sample, but not in the model, are shown in grey.

- Molecule 1: Lysozyme C

Chain A:

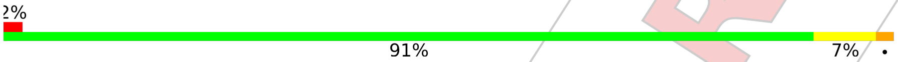


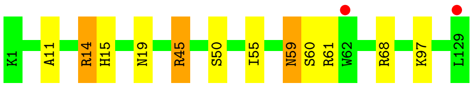


For Manuscript Review

### 4 Data and refinement statistics (i)

| Property | Value | Source |
|---|---|---|
| Space group | P 43 21 2 | Depositor |
| Cell constants<br>a, b, c, $\alpha$, $\beta$, $\gamma$ | 78.70Å 78.70Å 37.10Å<br>90.00° 90.00° 90.00° | Depositor |
| Resolution (Å) | 22.27 – 1.50<br>22.26 – 1.49 | Depositor<br>EDS |
| % Data completeness<br>(in resolution range) | 99.6 (22.27-1.50)<br>99.6 (22.26-1.49) | Depositor<br>EDS |
| $R_{merge}$ | (Not available) | Depositor |
| $R_{sym}$ | (Not available) | Depositor |
| $< I/\sigma(I) >$ [1] | 1.46 (at 1.49Å) | Xtriage |
| Refinement program | REFMAC 5.8.0267 | Depositor |
| R, $R_{free}$ | 0.187 , 0.217<br>0.196 , 0.222 | Depositor<br>DCC |
| $R_{free}$ test set | 992 reflections (5.12%) | wwPDB-VP |
| Wilson B-factor (Å$^2$) | 11.3 | Xtriage |
| Anisotropy | 0.032 | Xtriage |
| Bulk solvent $k_{sol}$(e/Å$^3$), $B_{sol}$(Å$^2$) | 0.34 , 36.2 | EDS |
| L-test for twinning[2] | $< \|L\| > = 0.48$, $< L^2 > = 0.32$ | Xtriage |
| Estimated twinning fraction | No twinning to report. | Xtriage |
| $F_o$,$F_c$ correlation | 0.95 | EDS |
| Total number of atoms | 1143 | wwPDB-VP |
| Average B, all atoms (Å$^2$) | 13.0 | wwPDB-VP |

Xtriage's analysis on translational NCS is as follows: *The largest off-origin peak in the Patterson function is 8.66% of the height of the origin peak. No significant pseudotranslation is detected.*

[1] Intensities estimated from amplitudes.
[2] Theoretical values of $< |L| >$, $< L^2 >$ for acentric reflections are 0.5, 0.333 respectively for untwinned datasets, and 0.375, 0.2 for perfectly twinned datasets.

For Manuscript Review

### 5 Model quality (i)

#### 5.1 Standard geometry (i)

Bond lengths and bond angles in the following residue types are not validated in this section: CL, NA, GOL

The Z score for a bond length (or angle) is the number of standard deviations the observed value is removed from the expected value. A bond length (or angle) with $|Z| > 5$ is considered an outlier worth inspection. RMSZ is the root-mean-square of all Z scores of the bond lengths (or angles).

| Mol | Chain | Bond lengths | | Bond angles | |
|---|---|---|---|---|---|
| | | RMSZ | #$\|Z\|$ >5 | RMSZ | #$\|Z\|$ >5 |
| 1 | A | 0.73 | 0/1032 | 0.96 | 4/1393 (0.3%) |

There are no bond length outliers.

All (4) bond angle outliers are listed below:

| Mol | Chain | Res | Type | Atoms | Z | Observed($^o$) | Ideal($^o$) |
|---|---|---|---|---|---|---|---|
| 1 | A | 45[A] | ARG | NE-CZ-NH2 | -6.59 | 117.00 | 120.30 |
| 1 | A | 45[B] | ARG | NE-CZ-NH2 | -6.59 | 117.00 | 120.30 |
| 1 | A | 45[A] | ARG | NE-CZ-NH1 | 5.29 | 122.95 | 120.30 |
| 1 | A | 45[B] | ARG | NE-CZ-NH1 | 5.29 | 122.95 | 120.30 |

There are no chirality outliers.

There are no planarity outliers.

#### 5.2 Too-close contacts (i)

In the following table, the Non-H and H(model) columns list the number of non-hydrogen atoms and hydrogen atoms in the chain respectively. The H(added) column lists the number of hydrogen atoms added and optimized by MolProbity. The Clashes column lists the number of clashes within the asymmetric unit, whereas Symm-Clashes lists symmetry-related clashes.

| Mol | Chain | Non-H | H(model) | H(added) | Clashes | Symm-Clashes |
|---|---|---|---|---|---|---|
| 1 | A | 1012 | 0 | 970 | 14 | 0 |
| 2 | A | 1 | 0 | 0 | 0 | 0 |
| 3 | A | 7 | 0 | 0 | 0 | 0 |
| 4 | A | 12 | 0 | 16 | 0 | 0 |
| 5 | A | 111 | 0 | 0 | 1 | 0 |
| All | All | 1143 | 0 | 986 | 14 | 0 |

For Manuscript Review

The all-atom clashscore is defined as the number of clashes found per 1000 atoms (including hydrogen atoms). The all-atom clashscore for this structure is 7.

All (14) close contacts within the same asymmetric unit are listed below, sorted by their clash magnitude.

| Atom-1 | Atom-2 | Interatomic distance (Å) | Clash overlap (Å) |
|---|---|---|---|
| 1:A:45[B]:ARG:HH21 | 1:A:45[B]:ARG:CG | 1.69 | 1.05 |
| 1:A:45[B]:ARG:HG3 | 1:A:45[B]:ARG:NH2 | 1.55 | 0.98 |
| 1:A:45[B]:ARG:NH2 | 5:A:301:HOH:O | 1.99 | 0.93 |
| 1:A:45[B]:ARG:HH21 | 1:A:45[B]:ARG:HG3 | 0.76 | 0.85 |
| 1:A:45[B]:ARG:HD3 | 1:A:68:ARG:NH2 | 2.12 | 0.64 |
| 1:A:50:SER:HB2 | 1:A:59:ASN:HD21 | 1.68 | 0.58 |
| 1:A:11:ALA:O | 1:A:15:HIS:HD2 | 1.87 | 0.58 |
| 1:A:50:SER:CB | 1:A:59:ASN:HD21 | 2.17 | 0.57 |
| 1:A:45[B]:ARG:HD3 | 1:A:68:ARG:HH21 | 1.72 | 0.55 |
| 1:A:59:ASN:HD22 | 1:A:60:SER:N | 2.06 | 0.53 |
| 1:A:50:SER:HB2 | 1:A:59:ASN:ND2 | 2.27 | 0.49 |
| 1:A:59:ASN:ND2 | 1:A:61:ARG:H | 2.13 | 0.45 |
| 1:A:14:ARG:HH11 | 1:A:14:ARG:CG | 2.34 | 0.41 |
| 1:A:45[B]:ARG:CD | 1:A:68:ARG:NH2 | 2.82 | 0.40 |

There are no symmetry-related clashes.

#### 5.3 Torsion angles (i)

##### 5.3.1 Protein backbone (i)

In the following table, the Percentiles column shows the percent Ramachandran outliers of the chain as a percentile score with respect to all X-ray entries followed by that with respect to entries of similar resolution.

The Analysed column shows the number of residues for which the backbone conformation was analysed, and the total number of residues.

| Mol | Chain | Analysed | Favoured | Allowed | Outliers | Percentiles | |
|---|---|---|---|---|---|---|---|
| 1 | A | 128/129 (99%) | 127 (99%) | 1 (1%) | 0 | 100 | 100 |

There are no Ramachandran outliers to report.

For Manuscript Review

##### 5.3.2 Protein sidechains (i)

In the following table, the Percentiles column shows the percent sidechain outliers of the chain as a percentile score with respect to all X-ray entries followed by that with respect to entries of similar resolution.

The Analysed column shows the number of residues for which the sidechain conformation was analysed, and the total number of residues.

| Mol | Chain | Analysed | Rotameric | Outliers | Percentiles | |
|---|---|---|---|---|---|---|
| 1 | A | 106/105 (101%) | 101 (95%) | 5 (5%) | 26 | 4 |

All (5) residues with a non-rotameric sidechain are listed below:

| Mol | Chain | Res | Type |
|---|---|---|---|
| 1 | A | 14 | ARG |
| 1 | A | 19 | ASN |
| 1 | A | 55 | ILE |
| 1 | A | 59 | ASN |
| 1 | A | 97 | LYS |

Sometimes sidechains can be flipped to improve hydrogen bonding and reduce clashes. All (4) such sidechains are listed below:

| Mol | Chain | Res | Type |
|---|---|---|---|
| 1 | A | 15 | HIS |
| 1 | A | 19 | ASN |
| 1 | A | 59 | ASN |
| 1 | A | 65 | ASN |

##### 5.3.3 RNA (i)

There are no RNA molecules in this entry.

#### 5.4 Non-standard residues in protein, DNA, RNA chains (i)

There are no non-standard protein/DNA/RNA residues in this entry.

#### 5.5 Carbohydrates (i)

There are no monosaccharides in this entry.

For Manuscript Review

#### 5.6 Ligand geometry (i)

Of 10 ligands modelled in this entry, 8 are monoatomic - leaving 2 for Mogul analysis.

In the following table, the Counts columns list the number of bonds (or angles) for which Mogul statistics could be retrieved, the number of bonds (or angles) that are observed in the model and the number of bonds (or angles) that are defined in the Chemical Component Dictionary. The Link column lists molecule types, if any, to which the group is linked. The Z score for a bond length (or angle) is the number of standard deviations the observed value is removed from the expected value. A bond length (or angle) with $|Z| > 2$ is considered an outlier worth inspection. RMSZ is the root-mean-square of all Z scores of the bond lengths (or angles).

| Mol | Type | Chain | Res | Link | Bond lengths | | | Bond angles | | |
|---|---|---|---|---|---|---|---|---|---|---|
| | | | | | Counts | RMSZ | $\#\|Z\| > 2$ | Counts | RMSZ | $\#\|Z\| > 2$ |
| 4 | GOL | A | 210 | - | 5,5,5 | 0.10 | 0 | 5,5,5 | 0.23 | 0 |
| 4 | GOL | A | 209 | - | 5,5,5 | 0.16 | 0 | 5,5,5 | 0.46 | 0 |

In the following table, the Chirals column lists the number of chiral outliers, the number of chiral centers analysed, the number of these observed in the model and the number defined in the Chemical Component Dictionary. Similar counts are reported in the Torsion and Rings columns. '-' means no outliers of that kind were identified.

| Mol | Type | Chain | Res | Link | Chirals | Torsions | Rings |
|---|---|---|---|---|---|---|---|
| 4 | GOL | A | 210 | - | - | 2/4/4/4 | - |
| 4 | GOL | A | 209 | - | - | 0/4/4/4 | - |

There are no bond length outliers.

There are no bond angle outliers.

There are no chirality outliers.

All (2) torsion outliers are listed below:

| Mol | Chain | Res | Type | Atoms |
|---|---|---|---|---|
| 4 | A | 210 | GOL | C1-C2-C3-O3 |
| 4 | A | 210 | GOL | O2-C2-C3-O3 |

There are no ring outliers.

No monomer is involved in short contacts.

#### 5.7 Other polymers (i)

There are no such residues in this entry.

For Manuscript Review

#### 5.8 Polymer linkage issues (i)

There are no chain breaks in this entry.

For Manuscript Review

### 6 Fit of model and data (i)

#### 6.1 Protein, DNA and RNA chains (i)

In the following table, the column labelled '#RSRZ> 2' contains the number (and percentage) of RSRZ outliers, followed by percent RSRZ outliers for the chain as percentile scores relative to all X-ray entries and entries of similar resolution. The OWAB column contains the minimum, median, $95^{th}$ percentile and maximum values of the occupancy-weighted average B-factor per residue. The column labelled 'Q< 0.9' lists the number of (and percentage) of residues with an average occupancy less than 0.9.

| Mol | Chain | Analysed | <RSRZ> | #RSRZ>2 | | | OWAB(Å$^2$) | Q<0.9 |
|---|---|---|---|---|---|---|---|---|
| 1 | A | 129/129 (100%) | 0.22 | 2 (1%) | 72 | 77 | 6, 11, 20, 26 | 0 |

All (2) RSRZ outliers are listed below:

| Mol | Chain | Res | Type | RSRZ |
|---|---|---|---|---|
| 1 | A | 129 | LEU | 4.9 |
| 1 | A | 62 | TRP | 2.6 |

#### 6.2 Non-standard residues in protein, DNA, RNA chains (i)

There are no non-standard protein/DNA/RNA residues in this entry.

#### 6.3 Carbohydrates (i)

There are no monosaccharides in this entry.

#### 6.4 Ligands (i)

In the following table, the Atoms column lists the number of modelled atoms in the group and the number defined in the chemical component dictionary. The B-factors column lists the minimum, median, $95^{th}$ percentile and maximum values of B factors of atoms in the group. The column labelled 'Q< 0.9' lists the number of atoms with occupancy less than 0.9.

| Mol | Type | Chain | Res | Atoms | RSCC | RSR | B-factors(Å$^2$) | Q<0.9 |
|---|---|---|---|---|---|---|---|---|
| 4 | GOL | A | 209 | 6/6 | 0.81 | 0.21 | 21,32,34,38 | 0 |
| 4 | GOL | A | 210 | 6/6 | 0.83 | 0.13 | 47,49,51,53 | 0 |
| 3 | CL | A | 208 | 1/1 | 0.96 | 0.23 | 33,33,33,33 | 0 |
| 3 | CL | A | 204 | 1/1 | 0.97 | 0.07 | 17,17,17,17 | 0 |
| 3 | CL | A | 205 | 1/1 | 0.97 | 0.05 | 24,24,24,24 | 0 |

*Continued on next page...*

For Manuscript Review

*Continued from previous page...*

| Mol | Type | Chain | Res | Atoms | RSCC | RSR | B-factors($Å^2$) | Q<0.9 |
|---|---|---|---|---|---|---|---|---|
| 3 | CL | A | 207 | 1/1 | 0.97 | 0.09 | 21,21,21,21 | 0 |
| 3 | CL | A | 206 | 1/1 | 0.98 | 0.09 | 24,24,24,24 | 0 |
| 3 | CL | A | 202 | 1/1 | 0.99 | 0.04 | 13,13,13,13 | 0 |
| 3 | CL | A | 203 | 1/1 | 0.99 | 0.03 | 15,15,15,15 | 0 |
| 2 | NA | A | 201 | 1/1 | 0.99 | 0.12 | 14,14,14,14 | 0 |

##### 6.5 Other polymers (i)

There are no such residues in this entry.

For Manuscript Review

### Full wwPDB X-ray Structure Validation Report (i)

Jun 7, 2022 – 01:54 PM EDT

| | | |
|---|---|---|
| PDB ID | : | 8D60 |
| Title | : | Crystal structure of hen egg white lysozyme at 150 Kelvin |
| Deposited on | : | 2022-06-06 |
| Resolution | : | 1.50 Å(reported) |

**This wwPDB validation report is for manuscript review**

This is a Full wwPDB X-ray Structure Validation Report.

This report is produced by the wwPDB biocuration pipeline after annotation of the structure.

We welcome your comments at **
A user guide is available at
https://www.wwpdb.org/validation/2017/XrayValidationReportHelp
with specific help available everywhere you see the (i) symbol.

For Manuscript Review

The following versions of software and data (see references (i)) were used in the production of this report:

| | | |
|---|---|---|
| MolProbity | : | 4.02b-467 |
| Mogul | : | 1.8.5 (274361), CSD as541be (2020) |
| Xtriage (Phenix) | : | 1.13 |
| EDS | : | 2.28 |
| Percentile statistics | : | 20191225.v01 (using entries in the PDB archive December 25th 2019) |
| Refmac | : | 5.8.0158 |
| CCP4 | : | 7.0.044 (Gargrove) |
| Ideal geometry (proteins) | : | Engh & Huber (2001) |
| Ideal geometry (DNA, RNA) | : | Parkinson et al. (1996) |
| Validation Pipeline (wwPDB-VP) | : | 2.28 |

### 1 Overall quality at a glance (i)

The following experimental techniques were used to determine the structure:
*X-RAY DIFFRACTION*

The reported resolution of this entry is 1.50 Å.

Percentile scores (ranging between 0-100) for global validation metrics of the entry are shown in the following graphic. The table shows the number of entries on which the scores are based.

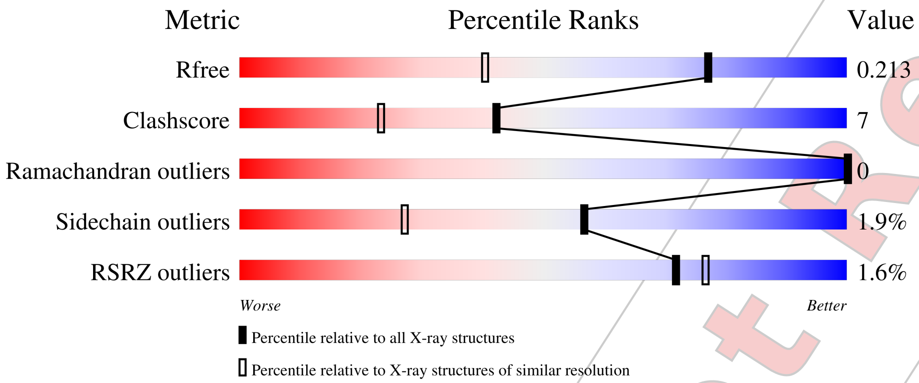


| Metric | Whole archive (#Entries) | Similar resolution (#Entries, resolution range(Å)) |
|---|---|---|
| $R_{free}$ | 130704 | 2936 (1.50-1.50) |
| Clashscore | 141614 | 3144 (1.50-1.50) |
| Ramachandran outliers | 138981 | 3066 (1.50-1.50) |
| Sidechain outliers | 138945 | 3064 (1.50-1.50) |
| RSRZ outliers | 127900 | 2884 (1.50-1.50) |

The table below summarises the geometric issues observed across the polymeric chains and their fit to the electron density. The red, orange, yellow and green segments of the lower bar indicate the fraction of residues that contain outliers for >=3, 2, 1 and 0 types of geometric quality criteria respectively. A grey segment represents the fraction of residues that are not modelled. The numeric value for each fraction is indicated below the corresponding segment, with a dot representing fractions <=5% The upper red bar (where present) indicates the fraction of residues that have poor fit to the electron density. The numeric value is given above the bar.

| Mol | Chain | Length | Quality of chain |
|---|---|---|---|
| 1 | A | 129 | 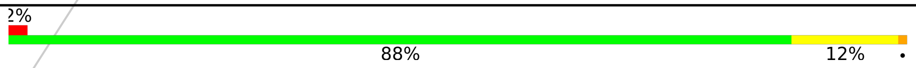  |

For Manuscript Review

### 2 Entry composition (i)

There are 5 unique types of molecules in this entry. The entry contains 1154 atoms, of which 0 are hydrogens and 0 are deuteriums.

In the tables below, the ZeroOcc column contains the number of atoms modelled with zero occupancy, the AltConf column contains the number of residues with at least one atom in alternate conformation and the Trace column contains the number of residues modelled with at most 2 atoms.

- Molecule 1 is a protein called Lysozyme C.

| Mol | Chain | Residues | Atoms | | | | | ZeroOcc | AltConf | Trace |
|---|---|---|---|---|---|---|---|---|---|---|
| | | | Total | C | N | O | S | | | |
| 1 | A | 129 | 1012 | 619 | 197 | 186 | 10 | 0 | 1 | 0 |

- Molecule 2 is SODIUM ION (three-letter code: NA) (formula: Na).

| Mol | Chain | Residues | Atoms | | ZeroOcc | AltConf |
|---|---|---|---|---|---|---|
| | | | Total | Na | | |
| 2 | A | 1 | 1 | 1 | 0 | 0 |

- Molecule 3 is CHLORIDE ION (three-letter code: CL) (formula: Cl).

| Mol | Chain | Residues | Atoms | | ZeroOcc | AltConf |
|---|---|---|---|---|---|---|
| | | | Total | Cl | | |
| 3 | A | 7 | 7 | 7 | 0 | 0 |

- Molecule 4 is GLYCEROL (three-letter code: GOL) (formula: $C_3H_8O_3$).

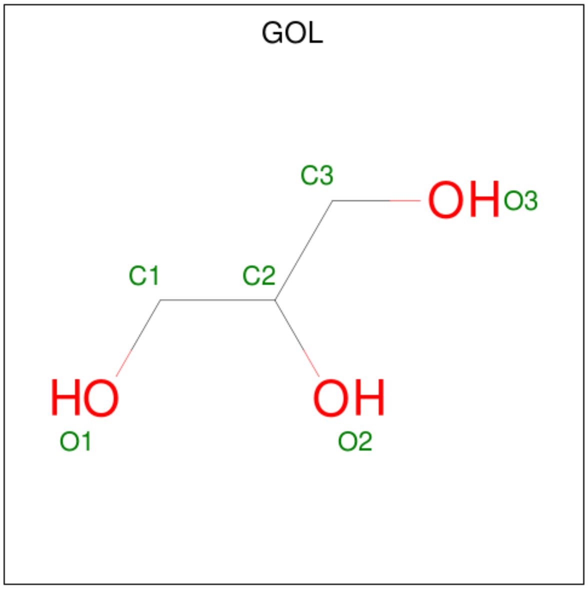


For Manuscript Review

| Mol | Chain | Residues | Atoms | ZeroOcc | AltConf |
|---|---|---|---|---|---|
| 4 | A | 1 | Total C O<br>6 3 3 | 0 | 0 |
| 4 | A | 1 | Total C O<br>6 3 3 | 0 | 0 |

- Molecule 5 is water.

| Mol | Chain | Residues | Atoms | ZeroOcc | AltConf |
|---|---|---|---|---|---|
| 5 | A | 122 | Total O<br>122 122 | 0 | 0 |

For Manuscript Review

### 3 Residue-property plots (i)

These plots are drawn for all protein, RNA, DNA and oligosaccharide chains in the entry. The first graphic for a chain summarises the proportions of the various outlier classes displayed in the second graphic. The second graphic shows the sequence view annotated by issues in geometry and electron density. Residues are color-coded according to the number of geometric quality criteria for which they contain at least one outlier: green = 0, yellow = 1, orange = 2 and red = 3 or more. A red dot above a residue indicates a poor fit to the electron density (RSRZ > 2). Stretches of 2 or more consecutive residues without any outlier are shown as a green connector. Residues present in the sample, but not in the model, are shown in grey.

• Molecule 1: Lysozyme C

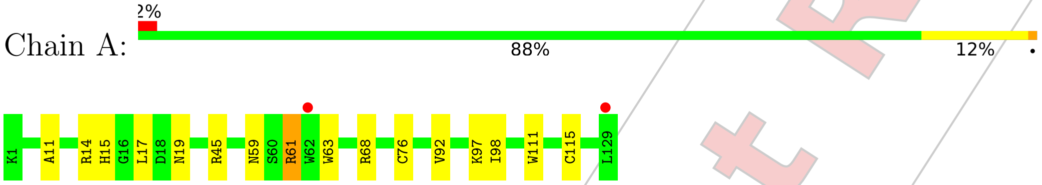


For Manuscript Review

### 4 Data and refinement statistics (i)

| Property | Value | Source |
|---|---|---|
| Space group | P 43 21 2 | Depositor |
| Cell constants<br>a, b, c, $\alpha$, $\beta$, $\gamma$ | 78.72Å 78.72Å 37.16Å<br>90.00° 90.00° 90.00° | Depositor |
| Resolution (Å) | 22.29 – 1.50<br>22.28 – 1.50 | Depositor<br>EDS |
| % Data completeness<br>(in resolution range) | 99.7 (22.29-1.50)<br>99.7 (22.28-1.50) | Depositor<br>EDS |
| $R_{merge}$ | (Not available) | Depositor |
| $R_{sym}$ | (Not available) | Depositor |
| $< I/\sigma(I) >$ [1] | 6.97 (at 1.50Å) | Xtriage |
| Refinement program | REFMAC 5.8.0267 | Depositor |
| R, $R_{free}$ | 0.170 , 0.209<br>0.178 , 0.213 | Depositor<br>DCC |
| $R_{free}$ test set | 988 reflections (5.12%) | wwPDB-VP |
| Wilson B-factor (Å$^2$) | 10.1 | Xtriage |
| Anisotropy | 0.013 | Xtriage |
| Bulk solvent $k_{sol}$(e/Å$^3$), $B_{sol}$(Å$^2$) | 0.35 , 38.7 | EDS |
| L-test for twinning[2] | $< \|L\| > = 0.50$, $< L^2 > = 0.33$ | Xtriage |
| Estimated twinning fraction | No twinning to report. | Xtriage |
| $F_o$,$F_c$ correlation | 0.95 | EDS |
| Total number of atoms | 1154 | wwPDB-VP |
| Average B, all atoms (Å$^2$) | 12.0 | wwPDB-VP |

Xtriage's analysis on translational NCS is as follows: *The largest off-origin peak in the Patterson function is 8.51% of the height of the origin peak. No significant pseudotranslation is detected.*

[1] Intensities estimated from amplitudes.

[2] Theoretical values of $< |L| >$, $< L^2 >$ for acentric reflections are 0.5, 0.333 respectively for untwinned datasets, and 0.375, 0.2 for perfectly twinned datasets.

For Manuscript Review

### 5 Model quality (i)

#### 5.1 Standard geometry (i)

Bond lengths and bond angles in the following residue types are not validated in this section: CL, GOL, NA

The Z score for a bond length (or angle) is the number of standard deviations the observed value is removed from the expected value. A bond length (or angle) with $|Z| > 5$ is considered an outlier worth inspection. RMSZ is the root-mean-square of all Z scores of the bond lengths (or angles).

| Mol | Chain | Bond lengths | | Bond angles | |
|---|---|---|---|---|---|
| | | RMSZ | #\|Z\| >5 | RMSZ | #\|Z\| >5 |
| 1 | A | 0.75 | 0/1032 | 0.96 | 2/1393 (0.1%) |

There are no bond length outliers.

All (2) bond angle outliers are listed below:

| Mol | Chain | Res | Type | Atoms | Z | Observed(°) | Ideal(°) |
|---|---|---|---|---|---|---|---|
| 1 | A | 14 | ARG | NE-CZ-NH2 | 5.64 | 123.12 | 120.30 |
| 1 | A | 61 | ARG | NE-CZ-NH1 | 5.09 | 122.84 | 120.30 |

There are no chirality outliers.

There are no planarity outliers.

#### 5.2 Too-close contacts (i)

In the following table, the Non-H and H(model) columns list the number of non-hydrogen atoms and hydrogen atoms in the chain respectively. The H(added) column lists the number of hydrogen atoms added and optimized by MolProbity. The Clashes column lists the number of clashes within the asymmetric unit, whereas Symm-Clashes lists symmetry-related clashes.

| Mol | Chain | Non-H | H(model) | H(added) | Clashes | Symm-Clashes |
|---|---|---|---|---|---|---|
| 1 | A | 1012 | 0 | 970 | 14 | 0 |
| 2 | A | 1 | 0 | 0 | 0 | 0 |
| 3 | A | 7 | 0 | 0 | 0 | 0 |
| 4 | A | 12 | 0 | 16 | 0 | 0 |
| 5 | A | 122 | 0 | 0 | 1 | 0 |
| All | All | 1154 | 0 | 986 | 14 | 0 |

The all-atom clashscore is defined as the number of clashes found per 1000 atoms (including hydrogen atoms). The all-atom clashscore for this structure is 7.

For Manuscript Review

All (14) close contacts within the same asymmetric unit are listed below, sorted by their clash magnitude.

| Atom-1 | Atom-2 | Interatomic distance (Å) | Clash overlap (Å) |
|---|---|---|---|
| 1:A:59:ASN:ND2 | 1:A:61:ARG:HB3 | 1.97 | 0.78 |
| 1:A:59:ASN:HD22 | 1:A:61:ARG:HB3 | 1.61 | 0.66 |
| 1:A:45:ARG:NE | 1:A:68[B]:ARG:NH2 | 2.46 | 0.64 |
| 1:A:59:ASN:ND2 | 1:A:61:ARG:H | 1.97 | 0.63 |
| 1:A:45:ARG:HE | 1:A:68[B]:ARG:NH2 | 1.96 | 0.63 |
| 1:A:59:ASN:ND2 | 1:A:61:ARG:CB | 2.61 | 0.62 |
| 1:A:11:ALA:O | 1:A:15:HIS:HD2 | 1.87 | 0.57 |
| 1:A:59:ASN:HD21 | 1:A:61:ARG:CB | 2.20 | 0.54 |
| 1:A:15:HIS:HE1 | 5:A:327:HOH:O | 1.92 | 0.53 |
| 1:A:17:LEU:HG | 1:A:92:VAL:HG13 | 1.99 | 0.44 |
| 1:A:59:ASN:HD21 | 1:A:61:ARG:HB2 | 1.82 | 0.43 |
| 1:A:63:TRP:CE2 | 1:A:98:ILE:HG12 | 2.55 | 0.41 |
| 1:A:111:TRP:CD1 | 1:A:115:CYS:HB2 | 2.55 | 0.41 |
| 1:A:63:TRP:O | 1:A:76:CYS:HB2 | 2.21 | 0.40 |

There are no symmetry-related clashes.

#### 5.3 Torsion angles (i)

##### 5.3.1 Protein backbone (i)

In the following table, the Percentiles column shows the percent Ramachandran outliers of the chain as a percentile score with respect to all X-ray entries followed by that with respect to entries of similar resolution.

The Analysed column shows the number of residues for which the backbone conformation was analysed, and the total number of residues.

| Mol | Chain | Analysed | Favoured | Allowed | Outliers | Percentiles | |
|---|---|---|---|---|---|---|---|
| 1 | A | 128/129 (99%) | 127 (99%) | 1 (1%) | 0 | 100 | 100 |

There are no Ramachandran outliers to report.

##### 5.3.2 Protein sidechains (i)

In the following table, the Percentiles column shows the percent sidechain outliers of the chain as a percentile score with respect to all X-ray entries followed by that with respect to entries of similar resolution.

For Manuscript Review

The Analysed column shows the number of residues for which the sidechain conformation was analysed, and the total number of residues.

| Mol | Chain | Analysed | Rotameric | Outliers | Percentiles | |
|---|---|---|---|---|---|---|
| 1 | A | 106/105 (101%) | 104 (98%) | 2 (2%) | 57 | 27 |

All (2) residues with a non-rotameric sidechain are listed below:

| Mol | Chain | Res | Type |
|---|---|---|---|
| 1 | A | 19 | ASN |
| 1 | A | 97 | LYS |

Sometimes sidechains can be flipped to improve hydrogen bonding and reduce clashes. All (5) such sidechains are listed below:

| Mol | Chain | Res | Type |
|---|---|---|---|
| 1 | A | 15 | HIS |
| 1 | A | 19 | ASN |
| 1 | A | 46 | ASN |
| 1 | A | 59 | ASN |
| 1 | A | 65 | ASN |

##### 5.3.3 RNA (i)

There are no RNA molecules in this entry.

#### 5.4 Non-standard residues in protein, DNA, RNA chains (i)

There are no non-standard protein/DNA/RNA residues in this entry.

#### 5.5 Carbohydrates (i)

There are no monosaccharides in this entry.

#### 5.6 Ligand geometry (i)

Of 10 ligands modelled in this entry, 8 are monoatomic - leaving 2 for Mogul analysis.

In the following table, the Counts columns list the number of bonds (or angles) for which Mogul statistics could be retrieved, the number of bonds (or angles) that are observed in the model and the number of bonds (or angles) that are defined in the Chemical Component Dictionary. The Link column lists molecule types, if any, to which the group is linked. The Z score for a bond

For Manuscript Review

length (or angle) is the number of standard deviations the observed value is removed from the expected value. A bond length (or angle) with $|Z| > 2$ is considered an outlier worth inspection. RMSZ is the root-mean-square of all Z scores of the bond lengths (or angles).

| Mol | Type | Chain | Res | Link | Bond lengths | | | Bond angles | | |
|---|---|---|---|---|---|---|---|---|---|---|
| | | | | | Counts | RMSZ | $\#\|Z\| > 2$ | Counts | RMSZ | $\#\|Z\| > 2$ |
| 4 | GOL | A | 210 | - | 5,5,5 | 0.10 | 0 | 5,5,5 | 0.20 | 0 |
| 4 | GOL | A | 209 | - | 5,5,5 | 0.13 | 0 | 5,5,5 | 0.39 | 0 |

In the following table, the Chirals column lists the number of chiral outliers, the number of chiral centers analysed, the number of these observed in the model and the number defined in the Chemical Component Dictionary. Similar counts are reported in the Torsion and Rings columns. '-' means no outliers of that kind were identified.

| Mol | Type | Chain | Res | Link | Chirals | Torsions | Rings |
|---|---|---|---|---|---|---|---|
| 4 | GOL | A | 210 | - | - | 0/4/4/4 | - |
| 4 | GOL | A | 209 | - | - | 0/4/4/4 | - |

There are no bond length outliers.

There are no bond angle outliers.

There are no chirality outliers.

There are no torsion outliers.

There are no ring outliers.

No monomer is involved in short contacts.

#### 5.7 Other polymers (i)

There are no such residues in this entry.

#### 5.8 Polymer linkage issues (i)

There are no chain breaks in this entry.

For Manuscript Review

### 6 Fit of model and data (i)

#### 6.1 Protein, DNA and RNA chains (i)

In the following table, the column labelled '#RSRZ> 2' contains the number (and percentage) of RSRZ outliers, followed by percent RSRZ outliers for the chain as percentile scores relative to all X-ray entries and entries of similar resolution. The OWAB column contains the minimum, median, $95^{th}$ percentile and maximum values of the occupancy-weighted average B-factor per residue. The column labelled 'Q< 0.9' lists the number of (and percentage) of residues with an average occupancy less than 0.9.

| Mol | Chain | Analysed | <RSRZ> | #RSRZ>2 | | | OWAB(Å$^2$) | Q<0.9 |
|---|---|---|---|---|---|---|---|---|
| 1 | A | 129/129 (100%) | 0.06 | 2 (1%) | 72 | 77 | 5, 10, 19, 24 | 0 |

All (2) RSRZ outliers are listed below:

| Mol | Chain | Res | Type | RSRZ |
|---|---|---|---|---|
| 1 | A | 129 | LEU | 3.1 |
| 1 | A | 62 | TRP | 2.5 |

#### 6.2 Non-standard residues in protein, DNA, RNA chains (i)

There are no non-standard protein/DNA/RNA residues in this entry.

#### 6.3 Carbohydrates (i)

There are no monosaccharides in this entry.

#### 6.4 Ligands (i)

In the following table, the Atoms column lists the number of modelled atoms in the group and the number defined in the chemical component dictionary. The B-factors column lists the minimum, median, $95^{th}$ percentile and maximum values of B factors of atoms in the group. The column labelled 'Q< 0.9' lists the number of atoms with occupancy less than 0.9.

| Mol | Type | Chain | Res | Atoms | RSCC | RSR | B-factors(Å$^2$) | Q<0.9 |
|---|---|---|---|---|---|---|---|---|
| 4 | GOL | A | 210 | 6/6 | 0.76 | 0.20 | 44,44,46,49 | 0 |
| 4 | GOL | A | 209 | 6/6 | 0.83 | 0.16 | 24,31,36,37 | 0 |
| 3 | CL | A | 208 | 1/1 | 0.93 | 0.15 | 29,29,29,29 | 0 |
| 3 | CL | A | 204 | 1/1 | 0.96 | 0.07 | 16,16,16,16 | 0 |
| 3 | CL | A | 205 | 1/1 | 0.97 | 0.05 | 23,23,23,23 | 0 |

*Continued on next page...*

For Manuscript Review

*Continued from previous page...*

| Mol | Type | Chain | Res | Atoms | RSCC | RSR | B-factors(Å$^2$) | Q<0.9 |
|---|---|---|---|---|---|---|---|---|
| 3 | CL | A | 207 | 1/1 | 0.98 | 0.12 | 20,20,20,20 | 0 |
| 3 | CL | A | 206 | 1/1 | 0.98 | 0.08 | 20,20,20,20 | 0 |
| 3 | CL | A | 202 | 1/1 | 0.99 | 0.05 | 12,12,12,12 | 0 |
| 3 | CL | A | 203 | 1/1 | 1.00 | 0.02 | 12,12,12,12 | 0 |
| 2 | NA | A | 201 | 1/1 | 1.00 | 0.05 | 10,10,10,10 | 0 |

#### 6.5 Other polymers ⓘ

There are no such residues in this entry.

For Manuscript Review

### Full wwPDB X-ray Structure Validation Report (i)

Jun 7, 2022 – 01:56 PM EDT

| | | |
|---|---|---|
| PDB ID | : | 8D61 |
| Title | : | Crystal structure of hen egg white lysozyme at 150 Kelvin (Duplicate) |
| Deposited on | : | 2022-06-06 |
| Resolution | : | 1.50 Å(reported) |

**This wwPDB validation report is for manuscript review**

This is a Full wwPDB X-ray Structure Validation Report.

This report is produced by the wwPDB biocuration pipeline after annotation of the structure.

We welcome your comments at **
A user guide is available at
https://www.wwpdb.org/validation/2017/XrayValidationReportHelp
with specific help available everywhere you see the (i) symbol.

The following versions of software and data (see references (i)) were used in the production of this report:

| | | |
|---|---|---|
| MolProbity | : | 4.02b-467 |
| Mogul | : | 1.8.5 (274361), CSD as541be (2020) |
| Xtriage (Phenix) | : | 1.13 |
| EDS | : | 2.28 |
| Percentile statistics | : | 20191225.v01 (using entries in the PDB archive December 25th 2019) |
| Refmac | : | 5.8.0158 |
| CCP4 | : | 7.0.044 (Gargrove) |
| Ideal geometry (proteins) | : | Engh & Huber (2001) |
| Ideal geometry (DNA, RNA) | : | Parkinson et al. (1996) |
| Validation Pipeline (wwPDB-VP) | : | 2.28 |

For Manuscript Review

### 1 Overall quality at a glance (i)

The following experimental techniques were used to determine the structure:
*X-RAY DIFFRACTION*

The reported resolution of this entry is 1.50 Å.

Percentile scores (ranging between 0-100) for global validation metrics of the entry are shown in the following graphic. The table shows the number of entries on which the scores are based.

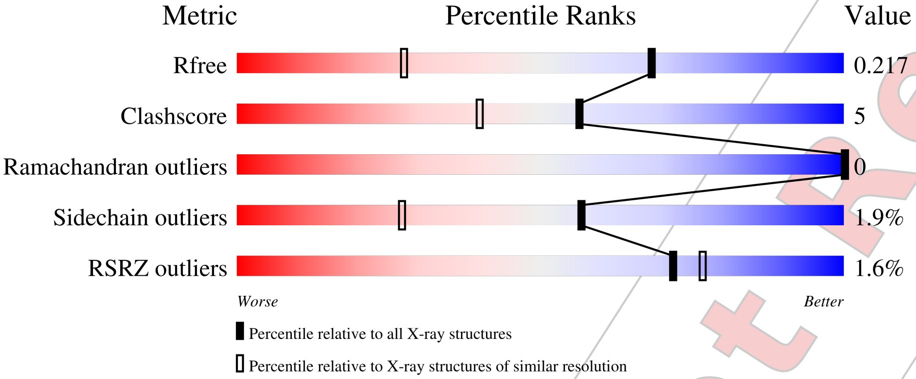


| Metric | Whole archive (#Entries) | Similar resolution (#Entries, resolution range(Å)) |
|---|---|---|
| $R_{free}$ | 130704 | 2936 (1.50-1.50) |
| Clashscore | 141614 | 3144 (1.50-1.50) |
| Ramachandran outliers | 138981 | 3066 (1.50-1.50) |
| Sidechain outliers | 138945 | 3064 (1.50-1.50) |
| RSRZ outliers | 127900 | 2884 (1.50-1.50) |

The table below summarises the geometric issues observed across the polymeric chains and their fit to the electron density. The red, orange, yellow and green segments of the lower bar indicate the fraction of residues that contain outliers for >=3, 2, 1 and 0 types of geometric quality criteria respectively. A grey segment represents the fraction of residues that are not modelled. The numeric value for each fraction is indicated below the corresponding segment, with a dot representing fractions <=5% The upper red bar (where present) indicates the fraction of residues that have poor fit to the electron density. The numeric value is given above the bar.

| Mol | Chain | Length | Quality of chain |
|---|---|---|---|
| 1 | A | 129 | 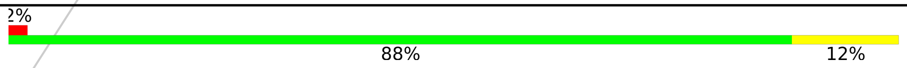 |

For Manuscript Review

#### 2 Entry composition (i)

There are 5 unique types of molecules in this entry. The entry contains 1134 atoms, of which 0 are hydrogens and 0 are deuteriums.

In the tables below, the ZeroOcc column contains the number of atoms modelled with zero occupancy, the AltConf column contains the number of residues with at least one atom in alternate conformation and the Trace column contains the number of residues modelled with at most 2 atoms.

- Molecule 1 is a protein called Lysozyme C.

| Mol | Chain | Residues | Atoms | | | | | ZeroOcc | AltConf | Trace |
|---|---|---|---|---|---|---|---|---|---|---|
| | | | Total | C | N | O | S | | | |
| 1 | A | 129 | 1001 | 613 | 193 | 185 | 10 | 0 | 0 | 0 |

- Molecule 2 is SODIUM ION (three-letter code: NA) (formula: Na).

| Mol | Chain | Residues | Atoms | | ZeroOcc | AltConf |
|---|---|---|---|---|---|---|
| | | | Total | Na | | |
| 2 | A | 1 | 1 | 1 | 0 | 0 |

- Molecule 3 is CHLORIDE ION (three-letter code: CL) (formula: Cl).

| Mol | Chain | Residues | Atoms | | ZeroOcc | AltConf |
|---|---|---|---|---|---|---|
| | | | Total | Cl | | |
| 3 | A | 7 | 7 | 7 | 0 | 0 |

- Molecule 4 is GLYCEROL (three-letter code: GOL) (formula: $C_3H_8O_3$).

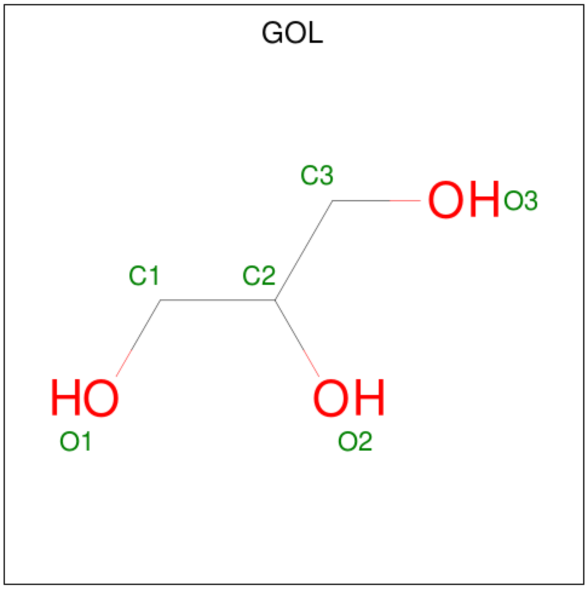


For Manuscript Review

| Mol | Chain | Residues | Atoms | | | ZeroOcc | AltConf |
|---|---|---|---|---|---|---|---|
| 4 | A | 1 | Total 6 | C 3 | O 3 | 0 | 0 |
| 4 | A | 1 | Total 6 | C 3 | O 3 | 0 | 0 |

- Molecule 5 is water.

| Mol | Chain | Residues | Atoms | | ZeroOcc | AltConf |
|---|---|---|---|---|---|---|
| 5 | A | 113 | Total 113 | O 113 | 0 | 0 |

For Manuscript Review

### 3 Residue-property plots (i)

These plots are drawn for all protein, RNA, DNA and oligosaccharide chains in the entry. The first graphic for a chain summarises the proportions of the various outlier classes displayed in the second graphic. The second graphic shows the sequence view annotated by issues in geometry and electron density. Residues are color-coded according to the number of geometric quality criteria for which they contain at least one outlier: green = 0, yellow = 1, orange = 2 and red = 3 or more. A red dot above a residue indicates a poor fit to the electron density (RSRZ > 2). Stretches of 2 or more consecutive residues without any outlier are shown as a green connector. Residues present in the sample, but not in the model, are shown in grey.

• Molecule 1: Lysozyme C

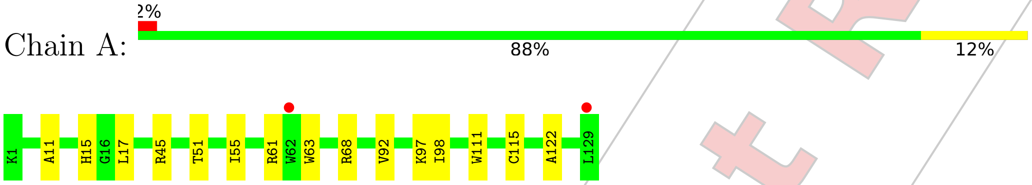


For Manuscript Review

### 4 Data and refinement statistics (i)

| Property | Value | Source |
|---|---|---|
| Space group | P 43 21 2 | Depositor |
| Cell constants<br>a, b, c, $\alpha$, $\beta$, $\gamma$ | 78.52Å 78.52Å 37.12Å<br>90.00° 90.00° 90.00° | Depositor |
| Resolution (Å) | 22.24 – 1.50<br>22.23 – 1.50 | Depositor<br>EDS |
| % Data completeness<br>(in resolution range) | 99.6 (22.24-1.50)<br>99.7 (22.23-1.50) | Depositor<br>EDS |
| $R_{merge}$ | (Not available) | Depositor |
| $R_{sym}$ | (Not available) | Depositor |
| $< I/\sigma(I) >$ [1] | 5.08 (at 1.50Å) | Xtriage |
| Refinement program | REFMAC 5.8.0267 | Depositor |
| R, $R_{free}$ | 0.176 , 0.211<br>0.185 , 0.217 | Depositor<br>DCC |
| $R_{free}$ test set | 983 reflections (5.13%) | wwPDB-VP |
| Wilson B-factor (Å$^2$) | 11.3 | Xtriage |
| Anisotropy | 0.031 | Xtriage |
| Bulk solvent $k_{sol}$(e/Å$^3$), $B_{sol}$(Å$^2$) | 0.36 , 41.2 | EDS |
| L-test for twinning[2] | $< \|L\| >$ = 0.47, $< L^2 >$ = 0.30 | Xtriage |
| Estimated twinning fraction | No twinning to report. | Xtriage |
| $F_o$,$F_c$ correlation | 0.95 | EDS |
| Total number of atoms | 1134 | wwPDB-VP |
| Average B, all atoms (Å$^2$) | 13.0 | wwPDB-VP |

Xtriage's analysis on translational NCS is as follows: *The largest off-origin peak in the Patterson function is 8.39% of the height of the origin peak. No significant pseudotranslation is detected.*

[1]Intensities estimated from amplitudes.
[2]Theoretical values of $< |L| >$, $< L^2 >$ for acentric reflections are 0.5, 0.333 respectively for untwinned datasets, and 0.375, 0.2 for perfectly twinned datasets.

For Manuscript Review

### 5 Model quality (i)

#### 5.1 Standard geometry (i)

Bond lengths and bond angles in the following residue types are not validated in this section: GOL, NA, CL

The Z score for a bond length (or angle) is the number of standard deviations the observed value is removed from the expected value. A bond length (or angle) with $|Z| > 5$ is considered an outlier worth inspection. RMSZ is the root-mean-square of all Z scores of the bond lengths (or angles).

| Mol | Chain | Bond lengths | | Bond angles | |
|---|---|---|---|---|---|
| | | RMSZ | #$|Z|$ >5 | RMSZ | #$|Z|$ >5 |
| 1 | A | 0.77 | 0/1021 | 0.94 | 2/1379 (0.1%) |

There are no bond length outliers.

All (2) bond angle outliers are listed below:

| Mol | Chain | Res | Type | Atoms | Z | Observed($^o$) | Ideal($^o$) |
|---|---|---|---|---|---|---|---|
| 1 | A | 61 | ARG | NE-CZ-NH1 | 5.26 | 122.93 | 120.30 |
| 1 | A | 61 | ARG | NE-CZ-NH2 | -5.01 | 117.80 | 120.30 |

There are no chirality outliers.

There are no planarity outliers.

#### 5.2 Too-close contacts (i)

In the following table, the Non-H and H(model) columns list the number of non-hydrogen atoms and hydrogen atoms in the chain respectively. The H(added) column lists the number of hydrogen atoms added and optimized by MolProbity. The Clashes column lists the number of clashes within the asymmetric unit, whereas Symm-Clashes lists symmetry-related clashes.

| Mol | Chain | Non-H | H(model) | H(added) | Clashes | Symm-Clashes |
|---|---|---|---|---|---|---|
| 1 | A | 1001 | 0 | 958 | 10 | 0 |
| 2 | A | 1 | 0 | 0 | 0 | 0 |
| 3 | A | 7 | 0 | 0 | 0 | 0 |
| 4 | A | 12 | 0 | 16 | 1 | 0 |
| 5 | A | 113 | 0 | 0 | 2 | 1 |
| All | All | 1134 | 0 | 974 | 10 | 1 |

The all-atom clashscore is defined as the number of clashes found per 1000 atoms (including hydrogen atoms). The all-atom clashscore for this structure is 5.

For Manuscript Review

All (10) close contacts within the same asymmetric unit are listed below, sorted by their clash magnitude.

| Atom-1 | Atom-2 | Interatomic distance (Å) | Clash overlap (Å) |
|---|---|---|---|
| 1:A:45:ARG:NE | 1:A:68:ARG:NH2 | 2.49 | 0.61 |
| 1:A:68:ARG:HD2 | 5:A:305:HOH:O | 2.01 | 0.60 |
| 1:A:11:ALA:O | 1:A:15:HIS:HD2 | 1.89 | 0.55 |
| 1:A:45:ARG:CZ | 1:A:68:ARG:NH2 | 2.77 | 0.47 |
| 1:A:45:ARG:HD3 | 1:A:51:THR:OG1 | 2.16 | 0.45 |
| 1:A:63:TRP:CE2 | 1:A:98:ILE:HG12 | 2.53 | 0.44 |
| 1:A:17:LEU:HG | 1:A:92:VAL:HG13 | 2.00 | 0.43 |
| 1:A:122:ALA:HB1 | 4:A:209:GOL:H32 | 2.02 | 0.42 |
| 1:A:111:TRP:CD1 | 1:A:115:CYS:HB2 | 2.56 | 0.41 |
| 1:A:45:ARG:NH2 | 5:A:305:HOH:O | 2.54 | 0.41 |

All (1) symmetry-related close contacts are listed below. The label for Atom-2 includes the symmetry operator and encoded unit-cell translations to be applied.

| Atom-1 | Atom-2 | Interatomic distance (Å) | Clash overlap (Å) |
|---|---|---|---|
| 5:A:404:HOH:O | 5:A:404:HOH:O[7_556] | 2.11 | 0.09 |

#### 5.3 Torsion angles (i)

##### 5.3.1 Protein backbone (i)

In the following table, the Percentiles column shows the percent Ramachandran outliers of the chain as a percentile score with respect to all X-ray entries followed by that with respect to entries of similar resolution.

The Analysed column shows the number of residues for which the backbone conformation was analysed, and the total number of residues.

| Mol | Chain | Analysed | Favoured | Allowed | Outliers | Percentiles | |
|---|---|---|---|---|---|---|---|
| 1 | A | 127/129 (98%) | 126 (99%) | 1 (1%) | 0 | 100 | 100 |

There are no Ramachandran outliers to report.

##### 5.3.2 Protein sidechains (i)

In the following table, the Percentiles column shows the percent sidechain outliers of the chain as a percentile score with respect to all X-ray entries followed by that with respect to entries of similar resolution.

For Manuscript Review

The Analysed column shows the number of residues for which the sidechain conformation was analysed, and the total number of residues.

| Mol | Chain | Analysed | Rotameric | Outliers | Percentiles | |
|---|---|---|---|---|---|---|
| 1 | A | 105/105 (100%) | 103 (98%) | 2 (2%) | 57 | 27 |

All (2) residues with a non-rotameric sidechain are listed below:

| Mol | Chain | Res | Type |
|---|---|---|---|
| 1 | A | 55 | ILE |
| 1 | A | 97 | LYS |

Sometimes sidechains can be flipped to improve hydrogen bonding and reduce clashes. All (2) such sidechains are listed below:

| Mol | Chain | Res | Type |
|---|---|---|---|
| 1 | A | 15 | HIS |
| 1 | A | 65 | ASN |

##### 5.3.3 RNA (i)

There are no RNA molecules in this entry.

#### 5.4 Non-standard residues in protein, DNA, RNA chains (i)

There are no non-standard protein/DNA/RNA residues in this entry.

#### 5.5 Carbohydrates (i)

There are no monosaccharides in this entry.

#### 5.6 Ligand geometry (i)

Of 10 ligands modelled in this entry, 8 are monoatomic - leaving 2 for Mogul analysis.

In the following table, the Counts columns list the number of bonds (or angles) for which Mogul statistics could be retrieved, the number of bonds (or angles) that are observed in the model and the number of bonds (or angles) that are defined in the Chemical Component Dictionary. The Link column lists molecule types, if any, to which the group is linked. The Z score for a bond length (or angle) is the number of standard deviations the observed value is removed from the expected value. A bond length (or angle) with $|Z| > 2$ is considered an outlier worth inspection. RMSZ is the root-mean-square of all Z scores of the bond lengths (or angles).

For Manuscript Review

| Mol | Type | Chain | Res | Link | Bond lengths | | | Bond angles | | |
|---|---|---|---|---|---|---|---|---|---|---|
| | | | | | Counts | RMSZ | #\|Z\| > 2 | Counts | RMSZ | #\|Z\| > 2 |
| 4 | GOL | A | 209 | - | 5,5,5 | 0.15 | 0 | 5,5,5 | 0.37 | 0 |
| 4 | GOL | A | 210 | - | 5,5,5 | 0.12 | 0 | 5,5,5 | 0.19 | 0 |

In the following table, the Chirals column lists the number of chiral outliers, the number of chiral centers analysed, the number of these observed in the model and the number defined in the Chemical Component Dictionary. Similar counts are reported in the Torsion and Rings columns. '-' means no outliers of that kind were identified.

| Mol | Type | Chain | Res | Link | Chirals | Torsions | Rings |
|---|---|---|---|---|---|---|---|
| 4 | GOL | A | 209 | - | - | 0/4/4/4 | - |
| 4 | GOL | A | 210 | - | - | 0/4/4/4 | - |

There are no bond length outliers.

There are no bond angle outliers.

There are no chirality outliers.

There are no torsion outliers.

There are no ring outliers.

1 monomer is involved in 1 short contact:

| Mol | Chain | Res | Type | Clashes | Symm-Clashes |
|---|---|---|---|---|---|
| 4 | A | 209 | GOL | 1 | 0 |

#### 5.7 Other polymers (i)

There are no such residues in this entry.

#### 5.8 Polymer linkage issues (i)

There are no chain breaks in this entry.

For Manuscript Review

### 6 Fit of model and data (i)

#### 6.1 Protein, DNA and RNA chains (i)

In the following table, the column labelled '#RSRZ> 2' contains the number (and percentage) of RSRZ outliers, followed by percent RSRZ outliers for the chain as percentile scores relative to all X-ray entries and entries of similar resolution. The OWAB column contains the minimum, median, $95^{th}$ percentile and maximum values of the occupancy-weighted average B-factor per residue. The column labelled 'Q< 0.9' lists the number of (and percentage) of residues with an average occupancy less than 0.9.

| Mol | Chain | Analysed | <RSRZ> | #RSRZ>2 | | | OWAB(Å$^2$) | Q<0.9 |
|---|---|---|---|---|---|---|---|---|
| 1 | A | 129/129 (100%) | -0.03 | 2 (1%) | 72 | 77 | 6, 11, 20, 26 | 0 |

All (2) RSRZ outliers are listed below:

| Mol | Chain | Res | Type | RSRZ |
|---|---|---|---|---|
| 1 | A | 129 | LEU | 2.7 |
| 1 | A | 62 | TRP | 2.3 |

#### 6.2 Non-standard residues in protein, DNA, RNA chains (i)

There are no non-standard protein/DNA/RNA residues in this entry.

#### 6.3 Carbohydrates (i)

There are no monosaccharides in this entry.

#### 6.4 Ligands (i)

In the following table, the Atoms column lists the number of modelled atoms in the group and the number defined in the chemical component dictionary. The B-factors column lists the minimum, median, $95^{th}$ percentile and maximum values of B factors of atoms in the group. The column labelled 'Q< 0.9' lists the number of atoms with occupancy less than 0.9.

| Mol | Type | Chain | Res | Atoms | RSCC | RSR | B-factors(Å$^2$) | Q<0.9 |
|---|---|---|---|---|---|---|---|---|
| 4 | GOL | A | 210 | 6/6 | 0.82 | 0.16 | 41,41,43,44 | 0 |
| 4 | GOL | A | 209 | 6/6 | 0.84 | 0.20 | 23,33,37,40 | 0 |
| 3 | CL | A | 208 | 1/1 | 0.93 | 0.21 | 33,33,33,33 | 0 |
| 3 | CL | A | 204 | 1/1 | 0.95 | 0.07 | 17,17,17,17 | 0 |
| 3 | CL | A | 207 | 1/1 | 0.96 | 0.07 | 22,22,22,22 | 0 |

*Continued on next page...*

For Manuscript Review

*Continued from previous page...*

| Mol | Type | Chain | Res | Atoms | RSCC | RSR | B-factors($Å^2$) | Q<0.9 |
|---|---|---|---|---|---|---|---|---|
| 3 | CL | A | 205 | 1/1 | 0.96 | 0.06 | 24,24,24,24 | 0 |
| 3 | CL | A | 206 | 1/1 | 0.98 | 0.07 | 22,22,22,22 | 0 |
| 3 | CL | A | 202 | 1/1 | 0.99 | 0.04 | 14,14,14,14 | 0 |
| 2 | NA | A | 201 | 1/1 | 0.99 | 0.06 | 11,11,11,11 | 0 |
| 3 | CL | A | 203 | 1/1 | 1.00 | 0.03 | 14,14,14,14 | 0 |

#### 6.5 Other polymers (i)

There are no such residues in this entry.

For Manuscript Review

### Full wwPDB X-ray Structure Validation Report (i)

Jun 7, 2022 – 01:57 PM EDT

| | | |
|---|---|---|
| PDB ID | : | 8D62 |
| Title | : | Crystal structure of hen egg white lysozyme at 150 Kelvin (Triplicate) |
| Deposited on | : | 2022-06-06 |
| Resolution | : | 1.50 Å(reported) |

**This wwPDB validation report is for manuscript review**

This is a Full wwPDB X-ray Structure Validation Report.

This report is produced by the wwPDB biocuration pipeline after annotation of the structure.

We welcome your comments at **
A user guide is available at
https://www.wwpdb.org/validation/2017/XrayValidationReportHelp
with specific help available everywhere you see the (i) symbol.

The following versions of software and data (see references (i)) were used in the production of this report:

| | | |
|---|---|---|
| MolProbity | : | 4.02b-467 |
| Mogul | : | 1.8.5 (274361), CSD as541be (2020) |
| Xtriage (Phenix) | : | 1.13 |
| EDS | : | 2.28 |
| Percentile statistics | : | 20191225.v01 (using entries in the PDB archive December 25th 2019) |
| Refmac | : | 5.8.0158 |
| CCP4 | : | 7.0.044 (Gargrove) |
| Ideal geometry (proteins) | : | Engh & Huber (2001) |
| Ideal geometry (DNA, RNA) | : | Parkinson et al. (1996) |
| Validation Pipeline (wwPDB-VP) | : | 2.28 |

For Manuscript Review

### 1 Overall quality at a glance (i)

The following experimental techniques were used to determine the structure:
*X-RAY DIFFRACTION*

The reported resolution of this entry is 1.50 Å.

Percentile scores (ranging between 0-100) for global validation metrics of the entry are shown in the following graphic. The table shows the number of entries on which the scores are based.

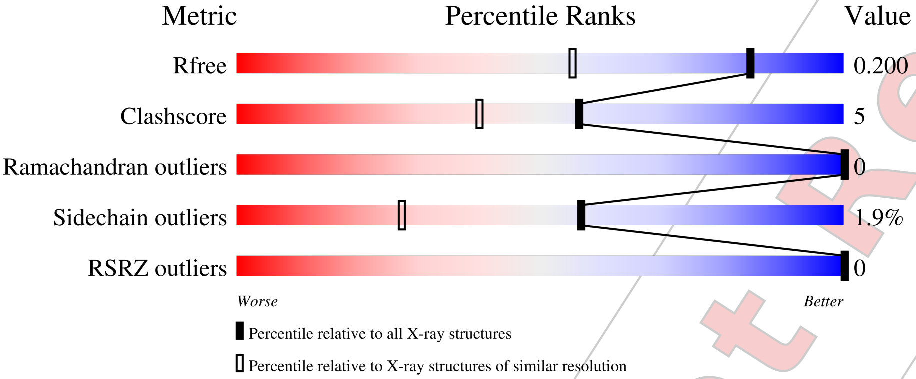


| Metric | Whole archive (#Entries) | Similar resolution (#Entries, resolution range(Å)) |
|---|---|---|
| $R_{free}$ | 130704 | 2936 (1.50-1.50) |
| Clashscore | 141614 | 3144 (1.50-1.50) |
| Ramachandran outliers | 138981 | 3066 (1.50-1.50) |
| Sidechain outliers | 138945 | 3064 (1.50-1.50) |
| RSRZ outliers | 127900 | 2884 (1.50-1.50) |

The table below summarises the geometric issues observed across the polymeric chains and their fit to the electron density. The red, orange, yellow and green segments of the lower bar indicate the fraction of residues that contain outliers for >=3, 2, 1 and 0 types of geometric quality criteria respectively. A grey segment represents the fraction of residues that are not modelled. The numeric value for each fraction is indicated below the corresponding segment, with a dot representing fractions <=5% The upper red bar (where present) indicates the fraction of residues that have poor fit to the electron density. The numeric value is given above the bar.

| Mol | Chain | Length | Quality of chain |
|---|---|---|---|
| 1 | A | 129 | 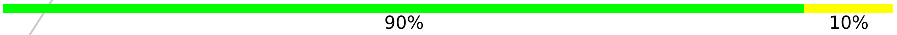 90% 10% |

For Manuscript Review

### 2 Entry composition (i)

There are 5 unique types of molecules in this entry. The entry contains 1147 atoms, of which 0 are hydrogens and 0 are deuteriums.

In the tables below, the ZeroOcc column contains the number of atoms modelled with zero occupancy, the AltConf column contains the number of residues with at least one atom in alternate conformation and the Trace column contains the number of residues modelled with at most 2 atoms.

- Molecule 1 is a protein called Lysozyme C.

| Mol | Chain | Residues | Atoms | | | | | ZeroOcc | AltConf | Trace |
|---|---|---|---|---|---|---|---|---|---|---|
| | | | Total | C | N | O | S | | | |
| 1 | A | 129 | 1001 | 613 | 193 | 185 | 10 | 0 | 0 | 0 |

- Molecule 2 is SODIUM ION (three-letter code: NA) (formula: Na).

| Mol | Chain | Residues | Atoms | | ZeroOcc | AltConf |
|---|---|---|---|---|---|---|
| | | | Total | Na | | |
| 2 | A | 1 | 1 | 1 | 0 | 0 |

- Molecule 3 is CHLORIDE ION (three-letter code: CL) (formula: Cl).

| Mol | Chain | Residues | Atoms | | ZeroOcc | AltConf |
|---|---|---|---|---|---|---|
| | | | Total | Cl | | |
| 3 | A | 7 | 7 | 7 | 0 | 0 |

- Molecule 4 is GLYCEROL (three-letter code: GOL) (formula: $C_3H_8O_3$).

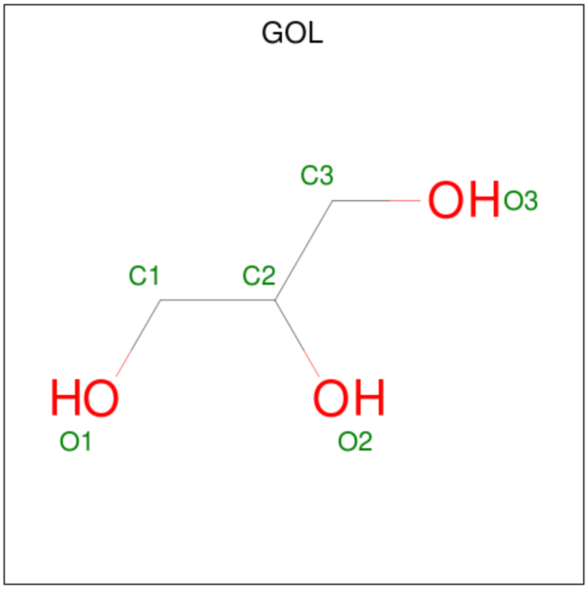


For Manuscript Review

| Mol | Chain | Residues | Atoms | ZeroOcc | AltConf |
|---|---|---|---|---|---|
| 4 | A | 1 | Total C O<br>6 3 3 | 0 | 0 |
| 4 | A | 1 | Total C O<br>6 3 3 | 0 | 0 |

- Molecule 5 is water.

| Mol | Chain | Residues | Atoms | ZeroOcc | AltConf |
|---|---|---|---|---|---|
| 5 | A | 126 | Total O<br>126 126 | 0 | 0 |

For Manuscript Review

### 3 Residue-property plots (i)

These plots are drawn for all protein, RNA, DNA and oligosaccharide chains in the entry. The first graphic for a chain summarises the proportions of the various outlier classes displayed in the second graphic. The second graphic shows the sequence view annotated by issues in geometry and electron density. Residues are color-coded according to the number of geometric quality criteria for which they contain at least one outlier: green = 0, yellow = 1, orange = 2 and red = 3 or more. A red dot above a residue indicates a poor fit to the electron density (RSRZ > 2). Stretches of 2 or more consecutive residues without any outlier are shown as a green connector. Residues present in the sample, but not in the model, are shown in grey.

• Molecule 1: Lysozyme C

Chain A:

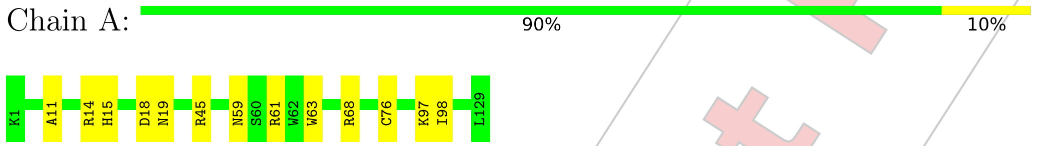


For Manuscript Review

### 4 Data and refinement statistics (i)

| Property | Value | Source |
|---|---|---|
| Space group | P 43 21 2 | Depositor |
| Cell constants<br>a, b, c, $\alpha$, $\beta$, $\gamma$ | 78.72Å 78.72Å 37.08Å<br>90.00° 90.00° 90.00° | Depositor |
| Resolution (Å) | 24.91 – 1.50<br>24.89 – 1.50 | Depositor<br>EDS |
| % Data completeness<br>(in resolution range) | 99.9 (24.91-1.50)<br>99.9 (24.89-1.50) | Depositor<br>EDS |
| $R_{merge}$ | (Not available) | Depositor |
| $R_{sym}$ | (Not available) | Depositor |
| $< I/\sigma(I) >$ [1] | 6.03 (at 1.50Å) | Xtriage |
| Refinement program | REFMAC 5.8.0267 | Depositor |
| R, $R_{free}$ | 0.165 , 0.190<br>0.174 , 0.200 | Depositor<br>DCC |
| $R_{free}$ test set | 986 reflections (5.12%) | wwPDB-VP |
| Wilson B-factor (Å$^2$) | 8.1 | Xtriage |
| Anisotropy | 0.038 | Xtriage |
| Bulk solvent $k_{sol}$(e/Å$^3$), $B_{sol}$(Å$^2$) | 0.38 , 41.2 | EDS |
| L-test for twinning[2] | $< \|L\| > = 0.48$, $< L^2 > = 0.31$ | Xtriage |
| Estimated twinning fraction | No twinning to report. | Xtriage |
| $F_o$,$F_c$ correlation | 0.95 | EDS |
| Total number of atoms | 1147 | wwPDB-VP |
| Average B, all atoms (Å$^2$) | 11.0 | wwPDB-VP |

Xtriage's analysis on translational NCS is as follows: *The largest off-origin peak in the Patterson function is 8.17% of the height of the origin peak. No significant pseudotranslation is detected.*

[1] Intensities estimated from amplitudes.
[2] Theoretical values of $< |L| >$, $< L^2 >$ for acentric reflections are 0.5, 0.333 respectively for untwinned datasets, and 0.375, 0.2 for perfectly twinned datasets.

For Manuscript Review

### 5 Model quality (i)

#### 5.1 Standard geometry (i)

Bond lengths and bond angles in the following residue types are not validated in this section: CL, GOL, NA

The Z score for a bond length (or angle) is the number of standard deviations the observed value is removed from the expected value. A bond length (or angle) with $|Z| > 5$ is considered an outlier worth inspection. RMSZ is the root-mean-square of all Z scores of the bond lengths (or angles).

| Mol | Chain | Bond lengths | | Bond angles | |
|---|---|---|---|---|---|
| | | RMSZ | #$\|Z\|$ >5 | RMSZ | #$\|Z\|$ >5 |
| 1 | A | 0.76 | 0/1021 | 0.94 | 2/1379 (0.1%) |

There are no bond length outliers.

All (2) bond angle outliers are listed below:

| Mol | Chain | Res | Type | Atoms | Z | Observed($^o$) | Ideal($^o$) |
|---|---|---|---|---|---|---|---|
| 1 | A | 18 | ASP | CB-CG-OD2 | -6.08 | 112.83 | 118.30 |
| 1 | A | 14 | ARG | NE-CZ-NH2 | 5.07 | 122.84 | 120.30 |

There are no chirality outliers.

There are no planarity outliers.

#### 5.2 Too-close contacts (i)

In the following table, the Non-H and H(model) columns list the number of non-hydrogen atoms and hydrogen atoms in the chain respectively. The H(added) column lists the number of hydrogen atoms added and optimized by MolProbity. The Clashes column lists the number of clashes within the asymmetric unit, whereas Symm-Clashes lists symmetry-related clashes.

| Mol | Chain | Non-H | H(model) | H(added) | Clashes | Symm-Clashes |
|---|---|---|---|---|---|---|
| 1 | A | 1001 | 0 | 958 | 10 | 0 |
| 2 | A | 1 | 0 | 0 | 0 | 0 |
| 3 | A | 7 | 0 | 0 | 0 | 0 |
| 4 | A | 12 | 0 | 16 | 0 | 0 |
| 5 | A | 126 | 0 | 0 | 3 | 0 |
| All | All | 1147 | 0 | 974 | 10 | 0 |

The all-atom clashscore is defined as the number of clashes found per 1000 atoms (including hydrogen atoms). The all-atom clashscore for this structure is 5.

For Manuscript Review

All (10) close contacts within the same asymmetric unit are listed below, sorted by their clash magnitude.

| Atom-1 | Atom-2 | Interatomic distance (Å) | Clash overlap (Å) |
|---|---|---|---|
| 1:A:45:ARG:NH1 | 5:A:301:HOH:O | 2.23 | 0.62 |
| 1:A:59:ASN:ND2 | 1:A:61:ARG:HB3 | 2.15 | 0.61 |
| 1:A:68:ARG:CD | 5:A:301:HOH:O | 2.50 | 0.59 |
| 1:A:59:ASN:ND2 | 1:A:61:ARG:H | 2.02 | 0.56 |
| 1:A:59:ASN:HD22 | 1:A:61:ARG:HB3 | 1.75 | 0.52 |
| 1:A:11:ALA:O | 1:A:15:HIS:HD2 | 1.94 | 0.51 |
| 1:A:59:ASN:ND2 | 1:A:61:ARG:CB | 2.75 | 0.50 |
| 1:A:63:TRP:O | 1:A:76:CYS:HB2 | 2.14 | 0.47 |
| 1:A:68:ARG:HD3 | 5:A:301:HOH:O | 2.14 | 0.44 |
| 1:A:63:TRP:CE2 | 1:A:98:ILE:HG12 | 2.54 | 0.43 |

There are no symmetry-related clashes.

#### 5.3 Torsion angles (i)

##### 5.3.1 Protein backbone (i)

In the following table, the Percentiles column shows the percent Ramachandran outliers of the chain as a percentile score with respect to all X-ray entries followed by that with respect to entries of similar resolution.

The Analysed column shows the number of residues for which the backbone conformation was analysed, and the total number of residues.

| Mol | Chain | Analysed | Favoured | Allowed | Outliers | Percentiles | |
|---|---|---|---|---|---|---|---|
| 1 | A | 127/129 (98%) | 124 (98%) | 3 (2%) | 0 | 100 | 100 |

There are no Ramachandran outliers to report.

##### 5.3.2 Protein sidechains (i)

In the following table, the Percentiles column shows the percent sidechain outliers of the chain as a percentile score with respect to all X-ray entries followed by that with respect to entries of similar resolution.

The Analysed column shows the number of residues for which the sidechain conformation was analysed, and the total number of residues.

For Manuscript Review

| Mol | Chain | Analysed | Rotameric | Outliers | Percentiles | |
|---|---|---|---|---|---|---|
| 1 | A | 105/105 (100%) | 103 (98%) | 2 (2%) | 57 | 27 |

All (2) residues with a non-rotameric sidechain are listed below:

| Mol | Chain | Res | Type |
|---|---|---|---|
| 1 | A | 19 | ASN |
| 1 | A | 97 | LYS |

Sometimes sidechains can be flipped to improve hydrogen bonding and reduce clashes. All (5) such sidechains are listed below:

| Mol | Chain | Res | Type |
|---|---|---|---|
| 1 | A | 15 | HIS |
| 1 | A | 19 | ASN |
| 1 | A | 46 | ASN |
| 1 | A | 59 | ASN |
| 1 | A | 65 | ASN |

###### 5.3.3 RNA (i)

There are no RNA molecules in this entry.

##### 5.4 Non-standard residues in protein, DNA, RNA chains (i)

There are no non-standard protein/DNA/RNA residues in this entry.

##### 5.5 Carbohydrates (i)

There are no monosaccharides in this entry.

##### 5.6 Ligand geometry (i)

Of 10 ligands modelled in this entry, 8 are monoatomic - leaving 2 for Mogul analysis.

In the following table, the Counts columns list the number of bonds (or angles) for which Mogul statistics could be retrieved, the number of bonds (or angles) that are observed in the model and the number of bonds (or angles) that are defined in the Chemical Component Dictionary. The Link column lists molecule types, if any, to which the group is linked. The Z score for a bond length (or angle) is the number of standard deviations the observed value is removed from the expected value. A bond length (or angle) with $|Z| > 2$ is considered an outlier worth inspection. RMSZ is the root-mean-square of all Z scores of the bond lengths (or angles).

For Manuscript Review

| Mol | Type | Chain | Res | Link | Bond lengths | | | Bond angles | | |
|---|---|---|---|---|---|---|---|---|---|---|
| | | | | | Counts | RMSZ | #\|Z\| > 2 | Counts | RMSZ | #\|Z\| > 2 |
| 4 | GOL | A | 210 | - | 5,5,5 | 0.13 | 0 | 5,5,5 | 0.27 | 0 |
| 4 | GOL | A | 209 | - | 5,5,5 | 0.14 | 0 | 5,5,5 | 0.33 | 0 |

In the following table, the Chirals column lists the number of chiral outliers, the number of chiral centers analysed, the number of these observed in the model and the number defined in the Chemical Component Dictionary. Similar counts are reported in the Torsion and Rings columns. '-' means no outliers of that kind were identified.

| Mol | Type | Chain | Res | Link | Chirals | Torsions | Rings |
|---|---|---|---|---|---|---|---|
| 4 | GOL | A | 210 | - | - | 0/4/4/4 | - |
| 4 | GOL | A | 209 | - | - | 0/4/4/4 | - |

There are no bond length outliers.

There are no bond angle outliers.

There are no chirality outliers.

There are no torsion outliers.

There are no ring outliers.

No monomer is involved in short contacts.

#### 5.7 Other polymers (i)

There are no such residues in this entry.

#### 5.8 Polymer linkage issues (i)

There are no chain breaks in this entry.

For Manuscript Review

### 6 Fit of model and data (i)

#### 6.1 Protein, DNA and RNA chains (i)

In the following table, the column labelled '#RSRZ> 2' contains the number (and percentage) of RSRZ outliers, followed by percent RSRZ outliers for the chain as percentile scores relative to all X-ray entries and entries of similar resolution. The OWAB column contains the minimum, median, 95$^{th}$ percentile and maximum values of the occupancy-weighted average B-factor per residue. The column labelled 'Q< 0.9' lists the number of (and percentage) of residues with an average occupancy less than 0.9.

| Mol | Chain | Analysed | <RSRZ> | #RSRZ>2 | | | OWAB(Å$^2$) | Q<0.9 |
|---|---|---|---|---|---|---|---|---|
| 1 | A | 129/129 (100%) | -0.22 | 0 | 100 | 100 | 5, 8, 18, 23 | 0 |

There are no RSRZ outliers to report.

#### 6.2 Non-standard residues in protein, DNA, RNA chains (i)

There are no non-standard protein/DNA/RNA residues in this entry.

#### 6.3 Carbohydrates (i)

There are no monosaccharides in this entry.

#### 6.4 Ligands (i)

In the following table, the Atoms column lists the number of modelled atoms in the group and the number defined in the chemical component dictionary. The B-factors column lists the minimum, median, 95$^{th}$ percentile and maximum values of B factors of atoms in the group. The column labelled 'Q< 0.9' lists the number of atoms with occupancy less than 0.9.

| Mol | Type | Chain | Res | Atoms | RSCC | RSR | B-factors(Å$^2$) | Q<0.9 |
|---|---|---|---|---|---|---|---|---|
| 4 | GOL | A | 210 | 6/6 | 0.77 | 0.17 | 31,31,33,41 | 0 |
| 4 | GOL | A | 209 | 6/6 | 0.82 | 0.14 | 26,30,36,38 | 0 |
| 3 | CL | A | 208 | 1/1 | 0.93 | 0.14 | 23,23,23,23 | 0 |
| 3 | CL | A | 204 | 1/1 | 0.96 | 0.09 | 15,15,15,15 | 0 |
| 3 | CL | A | 205 | 1/1 | 0.97 | 0.05 | 20,20,20,20 | 0 |
| 3 | CL | A | 207 | 1/1 | 0.98 | 0.07 | 22,22,22,22 | 0 |
| 3 | CL | A | 206 | 1/1 | 0.98 | 0.07 | 17,17,17,17 | 0 |
| 3 | CL | A | 202 | 1/1 | 0.99 | 0.03 | 10,10,10,10 | 0 |
| 2 | NA | A | 201 | 1/1 | 0.99 | 0.04 | 7,7,7,7 | 0 |
| 3 | CL | A | 203 | 1/1 | 1.00 | 0.02 | 10,10,10,10 | 0 |

For Manuscript Review

##### 6.5 Other polymers (i)

There are no such residues in this entry.

For Manuscript Review

### Full wwPDB X-ray Structure Validation Report (i)

Jun 7, 2022 – 01:59 PM EDT

| | | |
|---|---|---|
| PDB ID | : | 8D69 |
| Title | : | Crystal structure of hen egg white lysozyme at 175 Kelvin |
| Deposited on | : | 2022-06-06 |
| Resolution | : | 1.50 Å(reported) |

**This wwPDB validation report is for manuscript review**

This is a Full wwPDB X-ray Structure Validation Report.

This report is produced by the wwPDB biocuration pipeline after annotation of the structure.

We welcome your comments at **
A user guide is available at
https://www.wwpdb.org/validation/2017/XrayValidationReportHelp
with specific help available everywhere you see the (i) symbol.

The following versions of software and data (see references (i)) were used in the production of this report:

| | | |
|---|---|---|
| MolProbity | : | 4.02b-467 |
| Mogul | : | 1.8.5 (274361), CSD as541be (2020) |
| Xtriage (Phenix) | : | 1.13 |
| EDS | : | 2.28 |
| Percentile statistics | : | 20191225.v01 (using entries in the PDB archive December 25th 2019) |
| Refmac | : | 5.8.0158 |
| CCP4 | : | 7.0.044 (Gargrove) |
| Ideal geometry (proteins) | : | Engh & Huber (2001) |
| Ideal geometry (DNA, RNA) | : | Parkinson et al. (1996) |
| Validation Pipeline (wwPDB-VP) | : | 2.28 |

For Manuscript Review

### 1 Overall quality at a glance (i)

The following experimental techniques were used to determine the structure:
*X-RAY DIFFRACTION*

The reported resolution of this entry is 1.50 Å.

Percentile scores (ranging between 0-100) for global validation metrics of the entry are shown in the following graphic. The table shows the number of entries on which the scores are based.

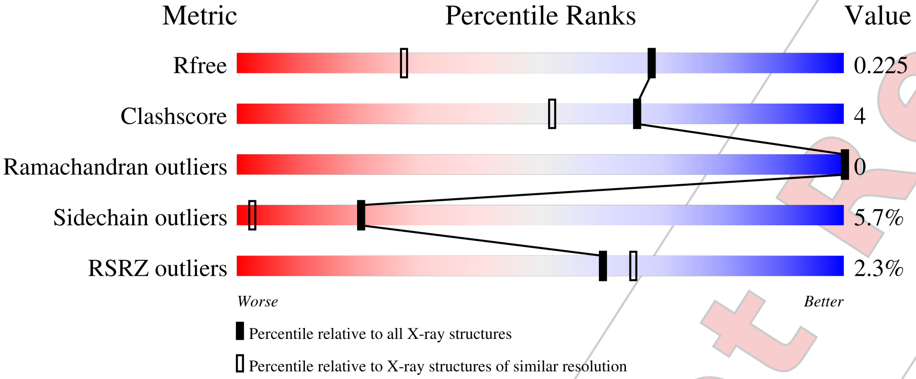


| Metric | Whole archive (#Entries) | Similar resolution (#Entries, resolution range(Å)) |
|---|---|---|
| $R_{free}$ | 130704 | 2936 (1.50-1.50) |
| Clashscore | 141614 | 3144 (1.50-1.50) |
| Ramachandran outliers | 138981 | 3066 (1.50-1.50) |
| Sidechain outliers | 138945 | 3064 (1.50-1.50) |
| RSRZ outliers | 127900 | 2884 (1.50-1.50) |

The table below summarises the geometric issues observed across the polymeric chains and their fit to the electron density. The red, orange, yellow and green segments of the lower bar indicate the fraction of residues that contain outliers for >=3, 2, 1 and 0 types of geometric quality criteria respectively. A grey segment represents the fraction of residues that are not modelled. The numeric value for each fraction is indicated below the corresponding segment, with a dot representing fractions <=5% The upper red bar (where present) indicates the fraction of residues that have poor fit to the electron density. The numeric value is given above the bar.

| Mol | Chain | Length | Quality of chain |
|---|---|---|---|
| 1 | A | 129 | 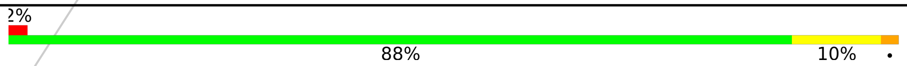 |

For Manuscript Review

#### 2 Entry composition (i)

There are 5 unique types of molecules in this entry. The entry contains 1132 atoms, of which 0 are hydrogens and 0 are deuteriums.

In the tables below, the ZeroOcc column contains the number of atoms modelled with zero occupancy, the AltConf column contains the number of residues with at least one atom in alternate conformation and the Trace column contains the number of residues modelled with at most 2 atoms.

- Molecule 1 is a protein called Lysozyme C.

| Mol | Chain | Residues | Atoms | | | | | ZeroOcc | AltConf | Trace |
|---|---|---|---|---|---|---|---|---|---|---|
| | | | Total | C | N | O | S | | | |
| 1 | A | 129 | 1013 | 619 | 195 | 189 | 10 | 0 | 2 | 0 |

- Molecule 2 is SODIUM ION (three-letter code: NA) (formula: Na).

| Mol | Chain | Residues | Atoms | | ZeroOcc | AltConf |
|---|---|---|---|---|---|---|
| | | | Total | Na | | |
| 2 | A | 1 | 1 | 1 | 0 | 0 |

- Molecule 3 is CHLORIDE ION (three-letter code: CL) (formula: Cl).

| Mol | Chain | Residues | Atoms | | ZeroOcc | AltConf |
|---|---|---|---|---|---|---|
| | | | Total | Cl | | |
| 3 | A | 7 | 7 | 7 | 0 | 0 |

- Molecule 4 is GLYCEROL (three-letter code: GOL) (formula: $C_3H_8O_3$).

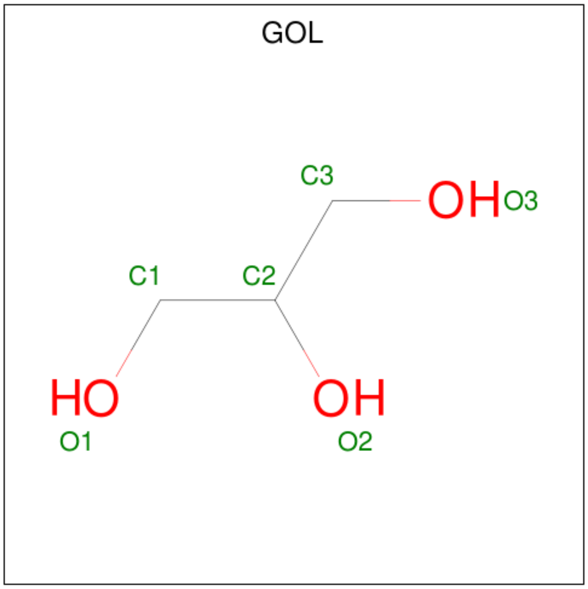


For Manuscript Review

| Mol | Chain | Residues | Atoms | ZeroOcc | AltConf |
|---|---|---|---|---|---|
| 4 | A | 1 | Total C O<br>6 3 3 | 0 | 0 |
| 4 | A | 1 | Total C O<br>6 3 3 | 0 | 0 |

- Molecule 5 is water.

| Mol | Chain | Residues | Atoms | ZeroOcc | AltConf |
|---|---|---|---|---|---|
| 5 | A | 99 | Total O<br>99 99 | 0 | 0 |

For Manuscript Review

### 3 Residue-property plots (i)

These plots are drawn for all protein, RNA, DNA and oligosaccharide chains in the entry. The first graphic for a chain summarises the proportions of the various outlier classes displayed in the second graphic. The second graphic shows the sequence view annotated by issues in geometry and electron density. Residues are color-coded according to the number of geometric quality criteria for which they contain at least one outlier: green = 0, yellow = 1, orange = 2 and red = 3 or more. A red dot above a residue indicates a poor fit to the electron density (RSRZ > 2). Stretches of 2 or more consecutive residues without any outlier are shown as a green connector. Residues present in the sample, but not in the model, are shown in grey.

• Molecule 1: Lysozyme C

Chain A:

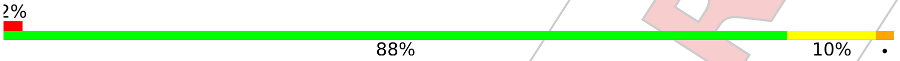


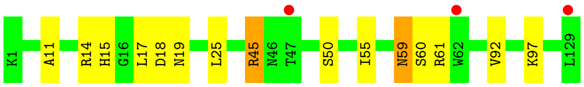


For Manuscript Review

### 4 Data and refinement statistics (i)

| Property | Value | Source |
|---|---|---|
| Space group | P 43 21 2 | Depositor |
| Cell constants<br>a, b, c, $\alpha$, $\beta$, $\gamma$ | 78.75Å 78.75Å 37.11Å<br>90.00° 90.00° 90.00° | Depositor |
| Resolution (Å) | 22.28 – 1.50<br>22.27 – 1.50 | Depositor<br>EDS |
| % Data completeness<br>(in resolution range) | 99.7 (22.28-1.50)<br>99.7 (22.27-1.50) | Depositor<br>EDS |
| $R_{merge}$ | (Not available) | Depositor |
| $R_{sym}$ | (Not available) | Depositor |
| $< I/\sigma(I) >$ [1] | 1.80 (at 1.49Å) | Xtriage |
| Refinement program | REFMAC 5.8.0267 | Depositor |
| R, $R_{free}$ | 0.187 , 0.220<br>0.195 , 0.225 | Depositor<br>DCC |
| $R_{free}$ test set | 988 reflections (5.11%) | wwPDB-VP |
| Wilson B-factor (Å$^2$) | 10.7 | Xtriage |
| Anisotropy | 0.021 | Xtriage |
| Bulk solvent $k_{sol}$(e/Å$^3$), $B_{sol}$(Å$^2$) | 0.35 , 39.5 | EDS |
| L-test for twinning[2] | $< \|L\| > = 0.47$, $< L^2 > = 0.31$ | Xtriage |
| Estimated twinning fraction | No twinning to report. | Xtriage |
| $F_o$,$F_c$ correlation | 0.95 | EDS |
| Total number of atoms | 1132 | wwPDB-VP |
| Average B, all atoms (Å$^2$) | 13.0 | wwPDB-VP |

Xtriage's analysis on translational NCS is as follows: *The largest off-origin peak in the Patterson function is 8.86% of the height of the origin peak. No significant pseudotranslation is detected.*

[1] Intensities estimated from amplitudes.

[2] Theoretical values of $< |L| >$, $< L^2 >$ for acentric reflections are 0.5, 0.333 respectively for untwinned datasets, and 0.375, 0.2 for perfectly twinned datasets.

For Manuscript Review

### 5 Model quality (i)

#### 5.1 Standard geometry (i)

Bond lengths and bond angles in the following residue types are not validated in this section: NA, GOL, CL

The Z score for a bond length (or angle) is the number of standard deviations the observed value is removed from the expected value. A bond length (or angle) with $|Z| > 5$ is considered an outlier worth inspection. RMSZ is the root-mean-square of all Z scores of the bond lengths (or angles).

| Mol | Chain | Bond lengths | | Bond angles | |
|---|---|---|---|---|---|
| | | RMSZ | #$\|Z\|$ >5 | RMSZ | #$\|Z\|$ >5 |
| 1 | A | 0.77 | 0/1033 | 1.06 | 2/1395 (0.1%) |

There are no bond length outliers.

All (2) bond angle outliers are listed below:

| Mol | Chain | Res | Type | Atoms | Z | Observed($^o$) | Ideal($^o$) |
|---|---|---|---|---|---|---|---|
| 1 | A | 45 | ARG | NE-CZ-NH1 | 13.89 | 127.25 | 120.30 |
| 1 | A | 45 | ARG | NE-CZ-NH2 | -13.15 | 113.72 | 120.30 |

There are no chirality outliers.

There are no planarity outliers.

#### 5.2 Too-close contacts (i)

In the following table, the Non-H and H(model) columns list the number of non-hydrogen atoms and hydrogen atoms in the chain respectively. The H(added) column lists the number of hydrogen atoms added and optimized by MolProbity. The Clashes column lists the number of clashes within the asymmetric unit, whereas Symm-Clashes lists symmetry-related clashes.

| Mol | Chain | Non-H | H(model) | H(added) | Clashes | Symm-Clashes |
|---|---|---|---|---|---|---|
| 1 | A | 1013 | 0 | 967 | 8 | 0 |
| 2 | A | 1 | 0 | 0 | 0 | 0 |
| 3 | A | 7 | 0 | 0 | 0 | 0 |
| 4 | A | 12 | 0 | 16 | 0 | 0 |
| 5 | A | 99 | 0 | 0 | 0 | 0 |
| All | All | 1132 | 0 | 983 | 8 | 0 |

The all-atom clashscore is defined as the number of clashes found per 1000 atoms (including hydrogen atoms). The all-atom clashscore for this structure is 4.

For Manuscript Review

All (8) close contacts within the same asymmetric unit are listed below, sorted by their clash magnitude.

| Atom-1 | Atom-2 | Interatomic distance (Å) | Clash overlap (Å) |
|---|---|---|---|
| 1:A:11:ALA:O | 1:A:15:HIS:HD2 | 1.85 | 0.60 |
| 1:A:59:ASN:HD22 | 1:A:60:SER:N | 2.06 | 0.53 |
| 1:A:50:SER:HB2 | 1:A:59:ASN:HD21 | 1.74 | 0.51 |
| 1:A:18:ASP:HB3 | 1:A:25:LEU:HD13 | 1.93 | 0.50 |
| 1:A:50:SER:CB | 1:A:59:ASN:HD21 | 2.25 | 0.49 |
| 1:A:59:ASN:ND2 | 1:A:61:ARG:H | 2.12 | 0.47 |
| 1:A:17:LEU:HG | 1:A:92:VAL:HG13 | 1.98 | 0.45 |
| 1:A:50:SER:HB2 | 1:A:59:ASN:ND2 | 2.34 | 0.42 |

There are no symmetry-related clashes.

#### 5.3 Torsion angles (i)

##### 5.3.1 Protein backbone (i)

In the following table, the Percentiles column shows the percent Ramachandran outliers of the chain as a percentile score with respect to all X-ray entries followed by that with respect to entries of similar resolution.

The Analysed column shows the number of residues for which the backbone conformation was analysed, and the total number of residues.

| Mol | Chain | Analysed | Favoured | Allowed | Outliers | Percentiles | |
|---|---|---|---|---|---|---|---|
| 1 | A | 129/129 (100%) | 128 (99%) | 1 (1%) | 0 | 100 | 100 |

There are no Ramachandran outliers to report.

##### 5.3.2 Protein sidechains (i)

In the following table, the Percentiles column shows the percent sidechain outliers of the chain as a percentile score with respect to all X-ray entries followed by that with respect to entries of similar resolution.

The Analysed column shows the number of residues for which the sidechain conformation was analysed, and the total number of residues.

| Mol | Chain | Analysed | Rotameric | Outliers | Percentiles | |
|---|---|---|---|---|---|---|
| 1 | A | 107/105 (102%) | 101 (94%) | 6 (6%) | 21 | 3 |

For Manuscript Review

All (6) residues with a non-rotameric sidechain are listed below:

| Mol | Chain | Res | Type |
|---|---|---|---|
| 1 | A | 14 | ARG |
| 1 | A | 19 | ASN |
| 1 | A | 45 | ARG |
| 1 | A | 55 | ILE |
| 1 | A | 59 | ASN |
| 1 | A | 97 | LYS |

Sometimes sidechains can be flipped to improve hydrogen bonding and reduce clashes. All (4) such sidechains are listed below:

| Mol | Chain | Res | Type |
|---|---|---|---|
| 1 | A | 15 | HIS |
| 1 | A | 19 | ASN |
| 1 | A | 59 | ASN |
| 1 | A | 65 | ASN |

##### 5.3.3 RNA (i)

There are no RNA molecules in this entry.

#### 5.4 Non-standard residues in protein, DNA, RNA chains (i)

There are no non-standard protein/DNA/RNA residues in this entry.

#### 5.5 Carbohydrates (i)

There are no monosaccharides in this entry.

#### 5.6 Ligand geometry (i)

Of 10 ligands modelled in this entry, 8 are monoatomic - leaving 2 for Mogul analysis.

In the following table, the Counts columns list the number of bonds (or angles) for which Mogul statistics could be retrieved, the number of bonds (or angles) that are observed in the model and the number of bonds (or angles) that are defined in the Chemical Component Dictionary. The Link column lists molecule types, if any, to which the group is linked. The Z score for a bond length (or angle) is the number of standard deviations the observed value is removed from the expected value. A bond length (or angle) with $|Z| > 2$ is considered an outlier worth inspection. RMSZ is the root-mean-square of all Z scores of the bond lengths (or angles).

For Manuscript Review

| Mol | Type | Chain | Res | Link | Bond lengths | | | Bond angles | | |
|---|---|---|---|---|---|---|---|---|---|---|
| | | | | | Counts | RMSZ | #\|Z\| > 2 | Counts | RMSZ | #\|Z\| > 2 |
| 4 | GOL | A | 210 | - | 5,5,5 | 0.10 | 0 | 5,5,5 | 0.22 | 0 |
| 4 | GOL | A | 209 | - | 5,5,5 | 0.15 | 0 | 5,5,5 | 0.46 | 0 |

In the following table, the Chirals column lists the number of chiral outliers, the number of chiral centers analysed, the number of these observed in the model and the number defined in the Chemical Component Dictionary. Similar counts are reported in the Torsion and Rings columns. '-' means no outliers of that kind were identified.

| Mol | Type | Chain | Res | Link | Chirals | Torsions | Rings |
|---|---|---|---|---|---|---|---|
| 4 | GOL | A | 210 | - | - | 2/4/4/4 | - |
| 4 | GOL | A | 209 | - | - | 0/4/4/4 | - |

There are no bond length outliers.

There are no bond angle outliers.

There are no chirality outliers.

All (2) torsion outliers are listed below:

| Mol | Chain | Res | Type | Atoms |
|---|---|---|---|---|
| 4 | A | 210 | GOL | O2-C2-C3-O3 |
| 4 | A | 210 | GOL | C1-C2-C3-O3 |

There are no ring outliers.

No monomer is involved in short contacts.

##### 5.7 Other polymers (i)

There are no such residues in this entry.

##### 5.8 Polymer linkage issues (i)

There are no chain breaks in this entry.

For Manuscript Review

### 6 Fit of model and data (i)

#### 6.1 Protein, DNA and RNA chains (i)

In the following table, the column labelled '#RSRZ> 2' contains the number (and percentage) of RSRZ outliers, followed by percent RSRZ outliers for the chain as percentile scores relative to all X-ray entries and entries of similar resolution. The OWAB column contains the minimum, median, $95^{th}$ percentile and maximum values of the occupancy-weighted average B-factor per residue. The column labelled 'Q< 0.9' lists the number of (and percentage) of residues with an average occupancy less than 0.9.

| Mol | Chain | Analysed | <RSRZ> | #RSRZ>2 | | | OWAB(Å$^2$) | Q<0.9 |
|---|---|---|---|---|---|---|---|---|
| 1 | A | 129/129 (100%) | 0.09 | 3 (2%) | 60 | 65 | 6, 11, 20, 24 | 0 |

All (3) RSRZ outliers are listed below:

| Mol | Chain | Res | Type | RSRZ |
|---|---|---|---|---|
| 1 | A | 129 | LEU | 3.9 |
| 1 | A | 47 | THR | 2.2 |
| 1 | A | 62 | TRP | 2.2 |

#### 6.2 Non-standard residues in protein, DNA, RNA chains (i)

There are no non-standard protein/DNA/RNA residues in this entry.

#### 6.3 Carbohydrates (i)

There are no monosaccharides in this entry.

#### 6.4 Ligands (i)

In the following table, the Atoms column lists the number of modelled atoms in the group and the number defined in the chemical component dictionary. The B-factors column lists the minimum, median, $95^{th}$ percentile and maximum values of B factors of atoms in the group. The column labelled 'Q< 0.9' lists the number of atoms with occupancy less than 0.9.

| Mol | Type | Chain | Res | Atoms | RSCC | RSR | B-factors(Å$^2$) | Q<0.9 |
|---|---|---|---|---|---|---|---|---|
| 4 | GOL | A | 210 | 6/6 | 0.76 | 0.20 | 49,50,52,52 | 0 |
| 4 | GOL | A | 209 | 6/6 | 0.82 | 0.24 | 23,34,36,40 | 0 |
| 3 | CL | A | 204 | 1/1 | 0.95 | 0.07 | 18,18,18,18 | 0 |
| 3 | CL | A | 206 | 1/1 | 0.95 | 0.09 | 30,30,30,30 | 0 |

*Continued on next page...*

For Manuscript Review

*Continued from previous page...*

| Mol | Type | Chain | Res | Atoms | RSCC | RSR | B-factors($Å^2$) | Q<0.9 |
|---|---|---|---|---|---|---|---|---|
| 3 | CL | A | 205 | 1/1 | 0.96 | 0.08 | 23,23,23,23 | 0 |
| 3 | CL | A | 208 | 1/1 | 0.96 | 0.31 | 37,37,37,37 | 0 |
| 3 | CL | A | 207 | 1/1 | 0.98 | 0.09 | 21,21,21,21 | 0 |
| 3 | CL | A | 202 | 1/1 | 0.99 | 0.03 | 14,14,14,14 | 0 |
| 2 | NA | A | 201 | 1/1 | 0.99 | 0.15 | 14,14,14,14 | 0 |
| 3 | CL | A | 203 | 1/1 | 1.00 | 0.05 | 16,16,16,16 | 0 |

#### 6.5 Other polymers ⓘ

There are no such residues in this entry.

For Manuscript Review

### Full wwPDB X-ray Structure Validation Report (i)

Jun 7, 2022 – 02:01 PM EDT

| | | |
|---|---|---|
| PDB ID | : | 8D6B |
| Title | : | Crystal structure of hen egg white lysozyme at 175 Kelvin (Duplicate) |
| Deposited on | : | 2022-06-06 |
| Resolution | : | 1.50 Å(reported) |

**This wwPDB validation report is for manuscript review**

This is a Full wwPDB X-ray Structure Validation Report.

This report is produced by the wwPDB biocuration pipeline after annotation of the structure.

We welcome your comments at **
A user guide is available at
https://www.wwpdb.org/validation/2017/XrayValidationReportHelp
with specific help available everywhere you see the (i) symbol.

For Manuscript Review

The following versions of software and data (see references (i)) were used in the production of this report:

| | | |
|---|---|---|
| MolProbity | : | 4.02b-467 |
| Mogul | : | 1.8.5 (274361), CSD as541be (2020) |
| Xtriage (Phenix) | : | 1.13 |
| EDS | : | 2.28 |
| Percentile statistics | : | 20191225.v01 (using entries in the PDB archive December 25th 2019) |
| Refmac | : | 5.8.0158 |
| CCP4 | : | 7.0.044 (Gargrove) |
| Ideal geometry (proteins) | : | Engh & Huber (2001) |
| Ideal geometry (DNA, RNA) | : | Parkinson et al. (1996) |
| Validation Pipeline (wwPDB-VP) | : | 2.28 |

### 1 Overall quality at a glance (i)

The following experimental techniques were used to determine the structure:
*X-RAY DIFFRACTION*

The reported resolution of this entry is 1.50 Å.

Percentile scores (ranging between 0-100) for global validation metrics of the entry are shown in the following graphic. The table shows the number of entries on which the scores are based.

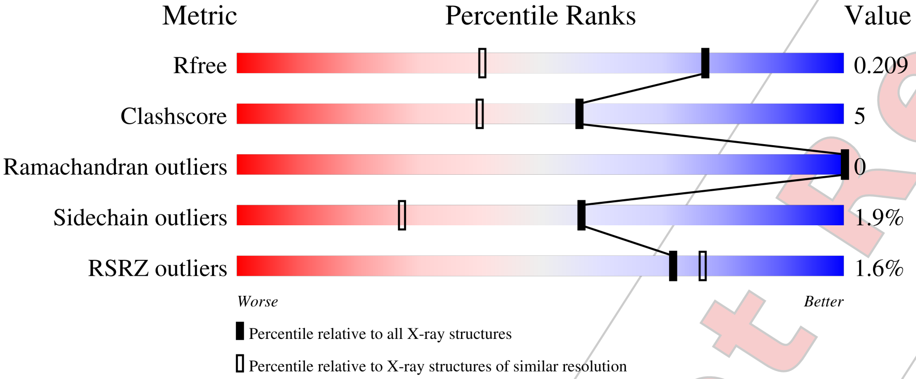


| Metric | Whole archive (#Entries) | Similar resolution (#Entries, resolution range(Å)) |
|---|---|---|
| $R_{free}$ | 130704 | 2936 (1.50-1.50) |
| Clashscore | 141614 | 3144 (1.50-1.50) |
| Ramachandran outliers | 138981 | 3066 (1.50-1.50) |
| Sidechain outliers | 138945 | 3064 (1.50-1.50) |
| RSRZ outliers | 127900 | 2884 (1.50-1.50) |

The table below summarises the geometric issues observed across the polymeric chains and their fit to the electron density. The red, orange, yellow and green segments of the lower bar indicate the fraction of residues that contain outliers for >=3, 2, 1 and 0 types of geometric quality criteria respectively. A grey segment represents the fraction of residues that are not modelled. The numeric value for each fraction is indicated below the corresponding segment, with a dot representing fractions <=5% The upper red bar (where present) indicates the fraction of residues that have poor fit to the electron density. The numeric value is given above the bar.

| Mol | Chain | Length | Quality of chain |
|---|---|---|---|
| 1 | A | 129 | 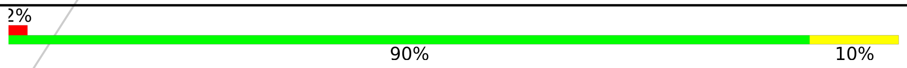  |

For Manuscript Review

### 2 Entry composition (i)

There are 5 unique types of molecules in this entry. The entry contains 1135 atoms, of which 0 are hydrogens and 0 are deuteriums.

In the tables below, the ZeroOcc column contains the number of atoms modelled with zero occupancy, the AltConf column contains the number of residues with at least one atom in alternate conformation and the Trace column contains the number of residues modelled with at most 2 atoms.

- Molecule 1 is a protein called Lysozyme C.

| Mol | Chain | Residues | Atoms | | | | | ZeroOcc | AltConf | Trace |
|---|---|---|---|---|---|---|---|---|---|---|
| | | | Total | C | N | O | S | | | |
| 1 | A | 129 | 1001 | 613 | 193 | 185 | 10 | 0 | 0 | 0 |

- Molecule 2 is SODIUM ION (three-letter code: NA) (formula: Na).

| Mol | Chain | Residues | Atoms | | ZeroOcc | AltConf |
|---|---|---|---|---|---|---|
| | | | Total | Na | | |
| 2 | A | 1 | 1 | 1 | 0 | 0 |

- Molecule 3 is CHLORIDE ION (three-letter code: CL) (formula: Cl).

| Mol | Chain | Residues | Atoms | | ZeroOcc | AltConf |
|---|---|---|---|---|---|---|
| | | | Total | Cl | | |
| 3 | A | 7 | 7 | 7 | 0 | 0 |

- Molecule 4 is GLYCEROL (three-letter code: GOL) (formula: $C_3H_8O_3$).

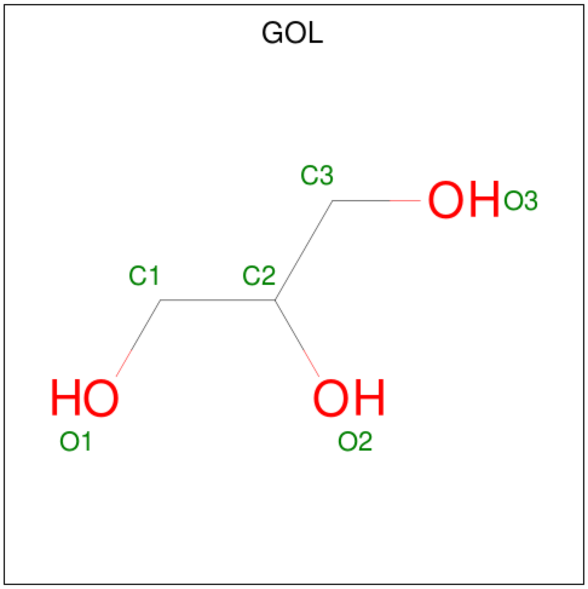


For Manuscript Review

| Mol | Chain | Residues | Atoms | ZeroOcc | AltConf |
|---|---|---|---|---|---|
| 4 | A | 1 | Total C O<br>6 3 3 | 0 | 0 |
| 4 | A | 1 | Total C O<br>6 3 3 | 0 | 0 |

- Molecule 5 is water.

| Mol | Chain | Residues | Atoms | ZeroOcc | AltConf |
|---|---|---|---|---|---|
| 5 | A | 114 | Total O<br>114 114 | 0 | 0 |

For Manuscript Review

### 3 Residue-property plots (i)

These plots are drawn for all protein, RNA, DNA and oligosaccharide chains in the entry. The first graphic for a chain summarises the proportions of the various outlier classes displayed in the second graphic. The second graphic shows the sequence view annotated by issues in geometry and electron density. Residues are color-coded according to the number of geometric quality criteria for which they contain at least one outlier: green = 0, yellow = 1, orange = 2 and red = 3 or more. A red dot above a residue indicates a poor fit to the electron density (RSRZ > 2). Stretches of 2 or more consecutive residues without any outlier are shown as a green connector. Residues present in the sample, but not in the model, are shown in grey.

- Molecule 1: Lysozyme C

Chain A:

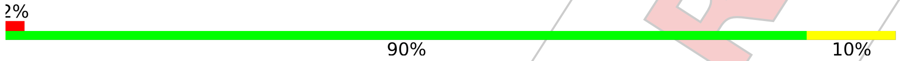


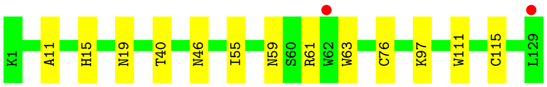


For Manuscript Review

### 4 Data and refinement statistics (i)

| Property | Value | Source |
|---|---|---|
| Space group | P 43 21 2 | Depositor |
| Cell constants<br>a, b, c, $\alpha$, $\beta$, $\gamma$ | 78.67Å 78.67Å 37.06Å<br>90.00° 90.00° 90.00° | Depositor |
| Resolution (Å) | 25.53 – 1.50<br>25.52 – 1.50 | Depositor<br>EDS |
| % Data completeness<br>(in resolution range) | 99.6 (25.53-1.50)<br>99.7 (25.52-1.50) | Depositor<br>EDS |
| $R_{merge}$ | (Not available) | Depositor |
| $R_{sym}$ | (Not available) | Depositor |
| $< I/\sigma(I) >$ [1] | 2.60 (at 1.49Å) | Xtriage |
| Refinement program | REFMAC 5.8.0267 | Depositor |
| R, $R_{free}$ | 0.172 , 0.204<br>0.180 , 0.209 | Depositor<br>DCC |
| $R_{free}$ test set | 984 reflections (5.12%) | wwPDB-VP |
| Wilson B-factor (Å$^2$) | 11.2 | Xtriage |
| Anisotropy | 0.034 | Xtriage |
| Bulk solvent $k_{sol}$(e/Å$^3$), $B_{sol}$(Å$^2$) | 0.37 , 40.6 | EDS |
| L-test for twinning[2] | $< \|L\| >= 0.48$, $< L^2 >= 0.32$ | Xtriage |
| Estimated twinning fraction | No twinning to report. | Xtriage |
| $F_o$,$F_c$ correlation | 0.95 | EDS |
| Total number of atoms | 1135 | wwPDB-VP |
| Average B, all atoms (Å$^2$) | 13.0 | wwPDB-VP |

Xtriage's analysis on translational NCS is as follows: *The largest off-origin peak in the Patterson function is 8.24% of the height of the origin peak. No significant pseudotranslation is detected.*

[1]Intensities estimated from amplitudes.
[2]Theoretical values of $< |L| >$, $< L^2 >$ for acentric reflections are 0.5, 0.333 respectively for untwinned datasets, and 0.375, 0.2 for perfectly twinned datasets.

For Manuscript Review

### 5 Model quality (i)

#### 5.1 Standard geometry (i)

Bond lengths and bond angles in the following residue types are not validated in this section: CL, NA, GOL

The Z score for a bond length (or angle) is the number of standard deviations the observed value is removed from the expected value. A bond length (or angle) with $|Z| > 5$ is considered an outlier worth inspection. RMSZ is the root-mean-square of all Z scores of the bond lengths (or angles).

| Mol | Chain | Bond lengths | | Bond angles | |
|---|---|---|---|---|---|
| | | RMSZ | $\#\|Z\| > 5$ | RMSZ | $\#\|Z\| > 5$ |
| 1 | A | 0.74 | 0/1021 | 0.88 | 0/1379 |

There are no bond length outliers.

There are no bond angle outliers.

There are no chirality outliers.

There are no planarity outliers.

#### 5.2 Too-close contacts (i)

In the following table, the Non-H and H(model) columns list the number of non-hydrogen atoms and hydrogen atoms in the chain respectively. The H(added) column lists the number of hydrogen atoms added and optimized by MolProbity. The Clashes column lists the number of clashes within the asymmetric unit, whereas Symm-Clashes lists symmetry-related clashes.

| Mol | Chain | Non-H | H(model) | H(added) | Clashes | Symm-Clashes |
|---|---|---|---|---|---|---|
| 1 | A | 1001 | 0 | 958 | 9 | 0 |
| 2 | A | 1 | 0 | 0 | 0 | 0 |
| 3 | A | 7 | 0 | 0 | 1 | 0 |
| 4 | A | 12 | 0 | 16 | 0 | 0 |
| 5 | A | 114 | 0 | 0 | 1 | 0 |
| All | All | 1135 | 0 | 974 | 10 | 0 |

The all-atom clashscore is defined as the number of clashes found per 1000 atoms (including hydrogen atoms). The all-atom clashscore for this structure is 5.

All (10) close contacts within the same asymmetric unit are listed below, sorted by their clash magnitude.

For Manuscript Review

| Atom-1 | Atom-2 | Interatomic distance (Å) | Clash overlap (Å) |
|---|---|---|---|
| 1:A:59:ASN:ND2 | 1:A:61:ARG:HB3 | 2.13 | 0.62 |
| 3:A:204:CL:CL | 5:A:385:HOH:O | 2.54 | 0.58 |
| 1:A:59:ASN:ND2 | 1:A:61:ARG:H | 2.04 | 0.56 |
| 1:A:59:ASN:HD22 | 1:A:61:ARG:HB3 | 1.69 | 0.56 |
| 1:A:46:ASN:ND2 | 1:A:59:ASN:OD1 | 2.41 | 0.53 |
| 1:A:59:ASN:ND2 | 1:A:61:ARG:CB | 2.71 | 0.52 |
| 1:A:11:ALA:O | 1:A:15:HIS:HD2 | 1.96 | 0.48 |
| 1:A:40:THR:HA | 1:A:55:ILE:HG23 | 1.98 | 0.46 |
| 1:A:111:TRP:CD1 | 1:A:115:CYS:HB2 | 2.54 | 0.42 |
| 1:A:63:TRP:O | 1:A:76:CYS:HB2 | 2.21 | 0.41 |

There are no symmetry-related clashes.

#### 5.3 Torsion angles ⓘ

##### 5.3.1 Protein backbone ⓘ

In the following table, the Percentiles column shows the percent Ramachandran outliers of the chain as a percentile score with respect to all X-ray entries followed by that with respect to entries of similar resolution.

The Analysed column shows the number of residues for which the backbone conformation was analysed, and the total number of residues.

| Mol | Chain | Analysed | Favoured | Allowed | Outliers | Percentiles | |
|---|---|---|---|---|---|---|---|
| 1 | A | 127/129 (98%) | 126 (99%) | 1 (1%) | 0 | 100 | 100 |

There are no Ramachandran outliers to report.

##### 5.3.2 Protein sidechains ⓘ

In the following table, the Percentiles column shows the percent sidechain outliers of the chain as a percentile score with respect to all X-ray entries followed by that with respect to entries of similar resolution.

The Analysed column shows the number of residues for which the sidechain conformation was analysed, and the total number of residues.

| Mol | Chain | Analysed | Rotameric | Outliers | Percentiles | |
|---|---|---|---|---|---|---|
| 1 | A | 105/105 (100%) | 103 (98%) | 2 (2%) | 57 | 27 |

For Manuscript Review

All (2) residues with a non-rotameric sidechain are listed below:

| Mol | Chain | Res | Type |
|---|---|---|---|
| 1 | A | 19 | ASN |
| 1 | A | 97 | LYS |

Sometimes sidechains can be flipped to improve hydrogen bonding and reduce clashes. All (5) such sidechains are listed below:

| Mol | Chain | Res | Type |
|---|---|---|---|
| 1 | A | 15 | HIS |
| 1 | A | 19 | ASN |
| 1 | A | 46 | ASN |
| 1 | A | 59 | ASN |
| 1 | A | 65 | ASN |

##### 5.3.3 RNA (i)

There are no RNA molecules in this entry.

#### 5.4 Non-standard residues in protein, DNA, RNA chains (i)

There are no non-standard protein/DNA/RNA residues in this entry.

#### 5.5 Carbohydrates (i)

There are no monosaccharides in this entry.

#### 5.6 Ligand geometry (i)

Of 10 ligands modelled in this entry, 8 are monoatomic - leaving 2 for Mogul analysis.

In the following table, the Counts columns list the number of bonds (or angles) for which Mogul statistics could be retrieved, the number of bonds (or angles) that are observed in the model and the number of bonds (or angles) that are defined in the Chemical Component Dictionary. The Link column lists molecule types, if any, to which the group is linked. The Z score for a bond length (or angle) is the number of standard deviations the observed value is removed from the expected value. A bond length (or angle) with $|Z| > 2$ is considered an outlier worth inspection. RMSZ is the root-mean-square of all Z scores of the bond lengths (or angles).

| Mol | Type | Chain | Res | Link | Bond lengths | | | Bond angles | | |
|---|---|---|---|---|---|---|---|---|---|---|
| | | | | | Counts | RMSZ | $\#\|Z\| > 2$ | Counts | RMSZ | $\#\|Z\| > 2$ |
| 4 | GOL | A | 210 | - | 5,5,5 | 0.13 | 0 | 5,5,5 | 0.19 | 0 |

For Manuscript Review

| Mol | Type | Chain | Res | Link | Bond lengths | | | Bond angles | | |
|---|---|---|---|---|---|---|---|---|---|---|
| | | | | | Counts | RMSZ | #\|Z\| > 2 | Counts | RMSZ | #\|Z\| > 2 |
| 4 | GOL | A | 209 | - | 5,5,5 | 0.16 | 0 | 5,5,5 | 0.38 | 0 |

In the following table, the Chirals column lists the number of chiral outliers, the number of chiral centers analysed, the number of these observed in the model and the number defined in the Chemical Component Dictionary. Similar counts are reported in the Torsion and Rings columns. '-' means no outliers of that kind were identified.

| Mol | Type | Chain | Res | Link | Chirals | Torsions | Rings |
|---|---|---|---|---|---|---|---|
| 4 | GOL | A | 210 | - | - | 0/4/4/4 | - |
| 4 | GOL | A | 209 | - | - | 0/4/4/4 | - |

There are no bond length outliers.

There are no bond angle outliers.

There are no chirality outliers.

There are no torsion outliers.

There are no ring outliers.

No monomer is involved in short contacts.

#### 5.7 Other polymers (i)

There are no such residues in this entry.

#### 5.8 Polymer linkage issues (i)

There are no chain breaks in this entry.

For Manuscript Review

### 6 Fit of model and data (i)

#### 6.1 Protein, DNA and RNA chains (i)

In the following table, the column labelled '#RSRZ> 2' contains the number (and percentage) of RSRZ outliers, followed by percent RSRZ outliers for the chain as percentile scores relative to all X-ray entries and entries of similar resolution. The OWAB column contains the minimum, median, $95^{th}$ percentile and maximum values of the occupancy-weighted average B-factor per residue. The column labelled 'Q< 0.9' lists the number of (and percentage) of residues with an average occupancy less than 0.9.

| Mol | Chain | Analysed | <RSRZ> | #RSRZ>2 | | | OWAB(Å$^2$) | Q<0.9 |
|---|---|---|---|---|---|---|---|---|
| 1 | A | 129/129 (100%) | 0.06 | 2 (1%) | 72 | 77 | 6, 11, 21, 26 | 0 |

All (2) RSRZ outliers are listed below:

| Mol | Chain | Res | Type | RSRZ |
|---|---|---|---|---|
| 1 | A | 129 | LEU | 3.6 |
| 1 | A | 62 | TRP | 2.0 |

#### 6.2 Non-standard residues in protein, DNA, RNA chains (i)

There are no non-standard protein/DNA/RNA residues in this entry.

#### 6.3 Carbohydrates (i)

There are no monosaccharides in this entry.

#### 6.4 Ligands (i)

In the following table, the Atoms column lists the number of modelled atoms in the group and the number defined in the chemical component dictionary. The B-factors column lists the minimum, median, $95^{th}$ percentile and maximum values of B factors of atoms in the group. The column labelled 'Q< 0.9' lists the number of atoms with occupancy less than 0.9.

| Mol | Type | Chain | Res | Atoms | RSCC | RSR | B-factors(Å$^2$) | Q<0.9 |
|---|---|---|---|---|---|---|---|---|
| 4 | GOL | A | 210 | 6/6 | 0.84 | 0.12 | 43,44,46,49 | 0 |
| 4 | GOL | A | 209 | 6/6 | 0.87 | 0.14 | 24,28,31,34 | 0 |
| 3 | CL | A | 204 | 1/1 | 0.94 | 0.08 | 19,19,19,19 | 0 |
| 3 | CL | A | 208 | 1/1 | 0.94 | 0.17 | 31,31,31,31 | 0 |
| 3 | CL | A | 205 | 1/1 | 0.96 | 0.06 | 22,22,22,22 | 0 |

*Continued on next page...*

For Manuscript Review

*Continued from previous page...*

| Mol | Type | Chain | Res | Atoms | RSCC | RSR | B-factors($Å^2$) | Q<0.9 |
|---|---|---|---|---|---|---|---|---|
| 3 | CL | A | 207 | 1/1 | 0.98 | 0.09 | 25,25,25,25 | 0 |
| 3 | CL | A | 202 | 1/1 | 0.99 | 0.04 | 14,14,14,14 | 0 |
| 3 | CL | A | 206 | 1/1 | 0.99 | 0.07 | 22,22,22,22 | 0 |
| 2 | NA | A | 201 | 1/1 | 0.99 | 0.08 | 12,12,12,12 | 0 |
| 3 | CL | A | 203 | 1/1 | 1.00 | 0.03 | 15,15,15,15 | 0 |

#### 6.5 Other polymers (i)

There are no such residues in this entry.

For Manuscript Review

For Manuscript Review

### Full wwPDB X-ray Structure Validation Report (i)

Jun 7, 2022 – 02:02 PM EDT

| | | |
|---|---|---|
| PDB ID | : | 8D6I |
| Title | : | Crystal structure of hen egg white lysozyme at 175 Kelvin (Triplicate) |
| Deposited on | : | 2022-06-06 |
| Resolution | : | 1.50 Å(reported) |

**This wwPDB validation report is for manuscript review**

This is a Full wwPDB X-ray Structure Validation Report.

This report is produced by the wwPDB biocuration pipeline after annotation of the structure.

We welcome your comments at **
A user guide is available at
https://www.wwpdb.org/validation/2017/XrayValidationReportHelp
with specific help available everywhere you see the (i) symbol.

The following versions of software and data (see references (i)) were used in the production of this report:

| | | |
|---|---|---|
| MolProbity | : | 4.02b-467 |
| Mogul | : | 1.8.5 (274361), CSD as541be (2020) |
| Xtriage (Phenix) | : | 1.13 |
| EDS | : | 2.28 |
| Percentile statistics | : | 20191225.v01 (using entries in the PDB archive December 25th 2019) |
| Refmac | : | 5.8.0158 |
| CCP4 | : | 7.0.044 (Gargrove) |
| Ideal geometry (proteins) | : | Engh & Huber (2001) |
| Ideal geometry (DNA, RNA) | : | Parkinson et al. (1996) |
| Validation Pipeline (wwPDB-VP) | : | 2.28 |

### 1 Overall quality at a glance (i)

The following experimental techniques were used to determine the structure:
*X-RAY DIFFRACTION*

The reported resolution of this entry is 1.50 Å.

Percentile scores (ranging between 0-100) for global validation metrics of the entry are shown in the following graphic. The table shows the number of entries on which the scores are based.

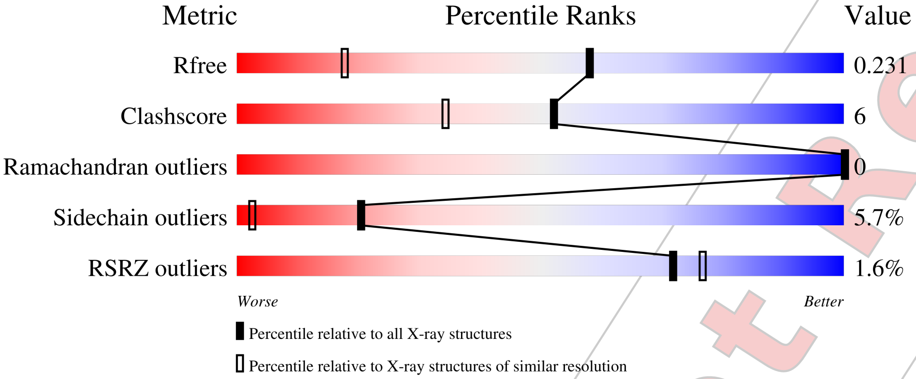


| Metric | Whole archive (#Entries) | Similar resolution (#Entries, resolution range(Å)) |
|---|---|---|
| $R_{free}$ | 130704 | 2936 (1.50-1.50) |
| Clashscore | 141614 | 3144 (1.50-1.50) |
| Ramachandran outliers | 138981 | 3066 (1.50-1.50) |
| Sidechain outliers | 138945 | 3064 (1.50-1.50) |
| RSRZ outliers | 127900 | 2884 (1.50-1.50) |

The table below summarises the geometric issues observed across the polymeric chains and their fit to the electron density. The red, orange, yellow and green segments of the lower bar indicate the fraction of residues that contain outliers for >=3, 2, 1 and 0 types of geometric quality criteria respectively. A grey segment represents the fraction of residues that are not modelled. The numeric value for each fraction is indicated below the corresponding segment, with a dot representing fractions <=5% The upper red bar (where present) indicates the fraction of residues that have poor fit to the electron density. The numeric value is given above the bar.

| Mol | Chain | Length | Quality of chain |
|---|---|---|---|
| 1 | A | 129 | 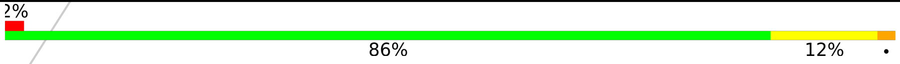 |

For Manuscript Review

#### 2 Entry composition (i)

There are 5 unique types of molecules in this entry. The entry contains 1123 atoms, of which 0 are hydrogens and 0 are deuteriums.

In the tables below, the ZeroOcc column contains the number of atoms modelled with zero occupancy, the AltConf column contains the number of residues with at least one atom in alternate conformation and the Trace column contains the number of residues modelled with at most 2 atoms.

- Molecule 1 is a protein called Lysozyme C.

| Mol | Chain | Residues | Atoms | | | | | ZeroOcc | AltConf | Trace |
|---|---|---|---|---|---|---|---|---|---|---|
| | | | Total | C | N | O | S | | | |
| 1 | A | 129 | 1001 | 613 | 193 | 185 | 10 | 0 | 0 | 0 |

- Molecule 2 is SODIUM ION (three-letter code: NA) (formula: Na).

| Mol | Chain | Residues | Atoms | | ZeroOcc | AltConf |
|---|---|---|---|---|---|---|
| | | | Total | Na | | |
| 2 | A | 1 | 1 | 1 | 0 | 0 |

- Molecule 3 is CHLORIDE ION (three-letter code: CL) (formula: Cl).

| Mol | Chain | Residues | Atoms | | ZeroOcc | AltConf |
|---|---|---|---|---|---|---|
| | | | Total | Cl | | |
| 3 | A | 7 | 7 | 7 | 0 | 0 |

- Molecule 4 is GLYCEROL (three-letter code: GOL) (formula: $C_3H_8O_3$).

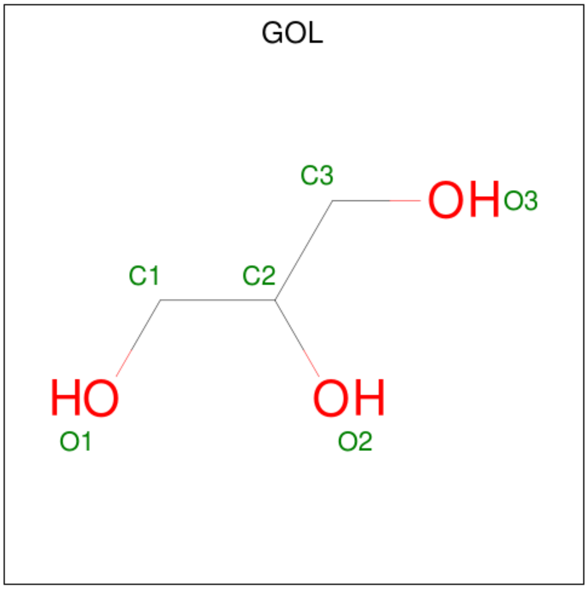


For Manuscript Review

| Mol | Chain | Residues | Atoms | | | ZeroOcc | AltConf |
|---|---|---|---|---|---|---|---|
| 4 | A | 1 | Total 6 | C 3 | O 3 | 0 | 0 |
| 4 | A | 1 | Total 6 | C 3 | O 3 | 0 | 0 |

- Molecule 5 is water.

| Mol | Chain | Residues | Atoms | | ZeroOcc | AltConf |
|---|---|---|---|---|---|---|
| 5 | A | 102 | Total 102 | O 102 | 0 | 0 |

For Manuscript Review

### 3 Residue-property plots (i)

These plots are drawn for all protein, RNA, DNA and oligosaccharide chains in the entry. The first graphic for a chain summarises the proportions of the various outlier classes displayed in the second graphic. The second graphic shows the sequence view annotated by issues in geometry and electron density. Residues are color-coded according to the number of geometric quality criteria for which they contain at least one outlier: green = 0, yellow = 1, orange = 2 and red = 3 or more. A red dot above a residue indicates a poor fit to the electron density (RSRZ > 2). Stretches of 2 or more consecutive residues without any outlier are shown as a green connector. Residues present in the sample, but not in the model, are shown in grey.

- Molecule 1: Lysozyme C

Chain A: 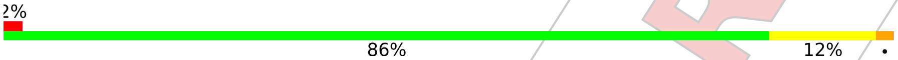


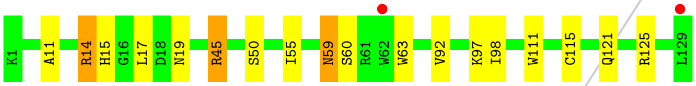


For Manuscript Review

### 4 Data and refinement statistics (i)

| Property | Value | Source |
|---|---|---|
| Space group | P 43 21 2 | Depositor |
| Cell constants<br>a, b, c, $\alpha$, $\beta$, $\gamma$ | 78.78Å 78.78Å 37.16Å<br>90.00° 90.00° 90.00° | Depositor |
| Resolution (Å) | 24.92 – 1.50<br>24.91 – 1.50 | Depositor<br>EDS |
| % Data completeness<br>(in resolution range) | 99.7 (24.92-1.50)<br>99.7 (24.91-1.50) | Depositor<br>EDS |
| $R_{merge}$ | (Not available) | Depositor |
| $R_{sym}$ | (Not available) | Depositor |
| $< I/\sigma(I) >$ [1] | 4.86 (at 1.50Å) | Xtriage |
| Refinement program | REFMAC 5.8.0267 | Depositor |
| R, $R_{free}$ | 0.185 , 0.223<br>0.195 , 0.231 | Depositor<br>DCC |
| $R_{free}$ test set | 986 reflections (5.12%) | wwPDB-VP |
| Wilson B-factor (Å$^2$) | 11.4 | Xtriage |
| Anisotropy | 0.149 | Xtriage |
| Bulk solvent $k_{sol}$(e/Å$^3$), $B_{sol}$(Å$^2$) | 0.36 , 38.9 | EDS |
| L-test for twinning[2] | $< \|L\| >$ = 0.48, $< L^2 >$ = 0.31 | Xtriage |
| Estimated twinning fraction | No twinning to report. | Xtriage |
| $F_o$,$F_c$ correlation | 0.95 | EDS |
| Total number of atoms | 1123 | wwPDB-VP |
| Average B, all atoms (Å$^2$) | 14.0 | wwPDB-VP |

Xtriage's analysis on translational NCS is as follows: *The largest off-origin peak in the Patterson function is 8.43% of the height of the origin peak. No significant pseudotranslation is detected.*

[1] Intensities estimated from amplitudes.

[2] Theoretical values of $< |L| >$, $< L^2 >$ for acentric reflections are 0.5, 0.333 respectively for untwinned datasets, and 0.375, 0.2 for perfectly twinned datasets.

For Manuscript Review

### 5 Model quality (i)

#### 5.1 Standard geometry (i)

Bond lengths and bond angles in the following residue types are not validated in this section: GOL, NA, CL

The Z score for a bond length (or angle) is the number of standard deviations the observed value is removed from the expected value. A bond length (or angle) with $|Z| > 5$ is considered an outlier worth inspection. RMSZ is the root-mean-square of all Z scores of the bond lengths (or angles).

| Mol | Chain | Bond lengths | | Bond angles | |
|---|---|---|---|---|---|
| | | RMSZ | #\|Z\| >5 | RMSZ | #\|Z\| >5 |
| 1 | A | 0.77 | 0/1021 | 0.95 | 2/1379 (0.1%) |

There are no bond length outliers.

All (2) bond angle outliers are listed below:

| Mol | Chain | Res | Type | Atoms | Z | Observed(°) | Ideal(°) |
|---|---|---|---|---|---|---|---|
| 1 | A | 45 | ARG | NE-CZ-NH1 | 9.15 | 124.87 | 120.30 |
| 1 | A | 45 | ARG | NE-CZ-NH2 | -7.70 | 116.45 | 120.30 |

There are no chirality outliers.

There are no planarity outliers.

#### 5.2 Too-close contacts (i)

In the following table, the Non-H and H(model) columns list the number of non-hydrogen atoms and hydrogen atoms in the chain respectively. The H(added) column lists the number of hydrogen atoms added and optimized by MolProbity. The Clashes column lists the number of clashes within the asymmetric unit, whereas Symm-Clashes lists symmetry-related clashes.

| Mol | Chain | Non-H | H(model) | H(added) | Clashes | Symm-Clashes |
|---|---|---|---|---|---|---|
| 1 | A | 1001 | 0 | 958 | 11 | 0 |
| 2 | A | 1 | 0 | 0 | 0 | 0 |
| 3 | A | 7 | 0 | 0 | 0 | 0 |
| 4 | A | 12 | 0 | 16 | 0 | 0 |
| 5 | A | 102 | 0 | 0 | 0 | 0 |
| All | All | 1123 | 0 | 974 | 11 | 0 |

The all-atom clashscore is defined as the number of clashes found per 1000 atoms (including hydrogen atoms). The all-atom clashscore for this structure is 6.

For Manuscript Review

All (11) close contacts within the same asymmetric unit are listed below, sorted by their clash magnitude.

| Atom-1 | Atom-2 | Interatomic distance (Å) | Clash overlap (Å) |
|---|---|---|---|
| 1:A:59:ASN:HD22 | 1:A:60:SER:N | 2.02 | 0.57 |
| 1:A:50:SER:HB2 | 1:A:59:ASN:HD21 | 1.70 | 0.56 |
| 1:A:121:GLN:OE1 | 1:A:125:ARG:NH2 | 2.33 | 0.54 |
| 1:A:50:SER:CB | 1:A:59:ASN:HD21 | 2.24 | 0.51 |
| 1:A:17:LEU:HG | 1:A:92:VAL:HG13 | 1.93 | 0.50 |
| 1:A:59:ASN:HD22 | 1:A:60:SER:H | 1.63 | 0.46 |
| 1:A:11:ALA:O | 1:A:15:HIS:HD2 | 1.98 | 0.46 |
| 1:A:14:ARG:HH11 | 1:A:14:ARG:HG2 | 1.79 | 0.46 |
| 1:A:63:TRP:CE2 | 1:A:98:ILE:HG12 | 2.54 | 0.43 |
| 1:A:14:ARG:HG2 | 1:A:14:ARG:NH1 | 2.33 | 0.42 |
| 1:A:111:TRP:CD1 | 1:A:115:CYS:HB2 | 2.55 | 0.41 |

There are no symmetry-related clashes.

#### 5.3 Torsion angles (i)

##### 5.3.1 Protein backbone (i)

In the following table, the Percentiles column shows the percent Ramachandran outliers of the chain as a percentile score with respect to all X-ray entries followed by that with respect to entries of similar resolution.

The Analysed column shows the number of residues for which the backbone conformation was analysed, and the total number of residues.

| Mol | Chain | Analysed | Favoured | Allowed | Outliers | Percentiles | |
|---|---|---|---|---|---|---|---|
| 1 | A | 127/129 (98%) | 126 (99%) | 1 (1%) | 0 | 100 | 100 |

There are no Ramachandran outliers to report.

##### 5.3.2 Protein sidechains (i)

In the following table, the Percentiles column shows the percent sidechain outliers of the chain as a percentile score with respect to all X-ray entries followed by that with respect to entries of similar resolution.

The Analysed column shows the number of residues for which the sidechain conformation was analysed, and the total number of residues.

For Manuscript Review

| Mol | Chain | Analysed | Rotameric | Outliers | Percentiles |
|---|---|---|---|---|---|
| 1 | A | 105/105 (100%) | 99 (94%) | 6 (6%) | 20 2 |

All (6) residues with a non-rotameric sidechain are listed below:

| Mol | Chain | Res | Type |
|---|---|---|---|
| 1 | A | 14 | ARG |
| 1 | A | 19 | ASN |
| 1 | A | 45 | ARG |
| 1 | A | 55 | ILE |
| 1 | A | 59 | ASN |
| 1 | A | 97 | LYS |

Sometimes sidechains can be flipped to improve hydrogen bonding and reduce clashes. All (4) such sidechains are listed below:

| Mol | Chain | Res | Type |
|---|---|---|---|
| 1 | A | 15 | HIS |
| 1 | A | 19 | ASN |
| 1 | A | 59 | ASN |
| 1 | A | 65 | ASN |

###### 5.3.3 RNA (i)

There are no RNA molecules in this entry.

##### 5.4 Non-standard residues in protein, DNA, RNA chains (i)

There are no non-standard protein/DNA/RNA residues in this entry.

##### 5.5 Carbohydrates (i)

There are no monosaccharides in this entry.

##### 5.6 Ligand geometry (i)

Of 10 ligands modelled in this entry, 8 are monoatomic - leaving 2 for Mogul analysis.

In the following table, the Counts columns list the number of bonds (or angles) for which Mogul statistics could be retrieved, the number of bonds (or angles) that are observed in the model and the number of bonds (or angles) that are defined in the Chemical Component Dictionary. The Link column lists molecule types, if any, to which the group is linked. The Z score for a bond

For Manuscript Review

length (or angle) is the number of standard deviations the observed value is removed from the expected value. A bond length (or angle) with $|Z| > 2$ is considered an outlier worth inspection. RMSZ is the root-mean-square of all Z scores of the bond lengths (or angles).

| Mol | Type | Chain | Res | Link | Bond lengths | | | Bond angles | | |
|---|---|---|---|---|---|---|---|---|---|---|
| | | | | | Counts | RMSZ | $\#\|Z\| > 2$ | Counts | RMSZ | $\#\|Z\| > 2$ |
| 4 | GOL | A | 209 | - | 5,5,5 | 0.16 | 0 | 5,5,5 | 0.39 | 0 |
| 4 | GOL | A | 210 | - | 5,5,5 | 0.13 | 0 | 5,5,5 | 0.19 | 0 |

In the following table, the Chirals column lists the number of chiral outliers, the number of chiral centers analysed, the number of these observed in the model and the number defined in the Chemical Component Dictionary. Similar counts are reported in the Torsion and Rings columns. '-' means no outliers of that kind were identified.

| Mol | Type | Chain | Res | Link | Chirals | Torsions | Rings |
|---|---|---|---|---|---|---|---|
| 4 | GOL | A | 209 | - | - | 0/4/4/4 | - |
| 4 | GOL | A | 210 | - | - | 0/4/4/4 | - |

There are no bond length outliers.

There are no bond angle outliers.

There are no chirality outliers.

There are no torsion outliers.

There are no ring outliers.

No monomer is involved in short contacts.

#### 5.7 Other polymers (i)

There are no such residues in this entry.

#### 5.8 Polymer linkage issues (i)

There are no chain breaks in this entry.

For Manuscript Review

### 6 Fit of model and data (i)

#### 6.1 Protein, DNA and RNA chains (i)

In the following table, the column labelled '#RSRZ> 2' contains the number (and percentage) of RSRZ outliers, followed by percent RSRZ outliers for the chain as percentile scores relative to all X-ray entries and entries of similar resolution. The OWAB column contains the minimum, median, $95^{th}$ percentile and maximum values of the occupancy-weighted average B-factor per residue. The column labelled 'Q< 0.9' lists the number of (and percentage) of residues with an average occupancy less than 0.9.

| Mol | Chain | Analysed | <RSRZ> | #RSRZ>2 | | | OWAB(Å$^2$) | Q<0.9 |
|---|---|---|---|---|---|---|---|---|
| 1 | A | 129/129 (100%) | 0.06 | 2 (1%) | 72 | 77 | 6, 12, 22, 30 | 0 |

All (2) RSRZ outliers are listed below:

| Mol | Chain | Res | Type | RSRZ |
|---|---|---|---|---|
| 1 | A | 129 | LEU | 3.0 |
| 1 | A | 62 | TRP | 2.3 |

#### 6.2 Non-standard residues in protein, DNA, RNA chains (i)

There are no non-standard protein/DNA/RNA residues in this entry.

#### 6.3 Carbohydrates (i)

There are no monosaccharides in this entry.

#### 6.4 Ligands (i)

In the following table, the Atoms column lists the number of modelled atoms in the group and the number defined in the chemical component dictionary. The B-factors column lists the minimum, median, $95^{th}$ percentile and maximum values of B factors of atoms in the group. The column labelled 'Q< 0.9' lists the number of atoms with occupancy less than 0.9.

| Mol | Type | Chain | Res | Atoms | RSCC | RSR | B-factors(Å$^2$) | Q<0.9 |
|---|---|---|---|---|---|---|---|---|
| 4 | GOL | A | 210 | 6/6 | 0.83 | 0.15 | 45,46,46,49 | 0 |
| 4 | GOL | A | 209 | 6/6 | 0.87 | 0.15 | 23,30,31,36 | 0 |
| 3 | CL | A | 208 | 1/1 | 0.94 | 0.22 | 34,34,34,34 | 0 |
| 3 | CL | A | 205 | 1/1 | 0.95 | 0.06 | 23,23,23,23 | 0 |
| 3 | CL | A | 204 | 1/1 | 0.95 | 0.07 | 19,19,19,19 | 0 |

*Continued on next page...*

For Manuscript Review

*Continued from previous page...*

| Mol | Type | Chain | Res | Atoms | RSCC | RSR | B-factors($Å^2$) | Q<0.9 |
|---|---|---|---|---|---|---|---|---|
| 3 | CL | A | 207 | 1/1 | 0.97 | 0.10 | 25,25,25,25 | 0 |
| 3 | CL | A | 202 | 1/1 | 0.99 | 0.04 | 15,15,15,15 | 0 |
| 3 | CL | A | 206 | 1/1 | 0.99 | 0.05 | 24,24,24,24 | 0 |
| 2 | NA | A | 201 | 1/1 | 0.99 | 0.09 | 13,13,13,13 | 0 |
| 3 | CL | A | 203 | 1/1 | 1.00 | 0.03 | 16,16,16,16 | 0 |

#### 6.5 Other polymers (i)

There are no such residues in this entry.

For Manuscript Review

### Full wwPDB X-ray Structure Validation Report (i)

Jun 8, 2022 – 03:46 PM EDT

| | | |
|---|---|---|
| PDB ID | : | 8D75 |
| Title | : | Crystal structure of hen egg white lysozyme at 200 Kelvin |
| Deposited on | : | 2022-06-07 |
| Resolution | : | 1.50 Å(reported) |

**This wwPDB validation report is for manuscript review**

This is a Full wwPDB X-ray Structure Validation Report.

This report is produced by the wwPDB biocuration pipeline after annotation of the structure.

We welcome your comments at **
A user guide is available at
https://www.wwpdb.org/validation/2017/XrayValidationReportHelp
with specific help available everywhere you see the (i) symbol.

The following versions of software and data (see references (i)) were used in the production of this report:

| | | |
|---|---|---|
| MolProbity | : | 4.02b-467 |
| Mogul | : | 1.8.5 (274361), CSD as541be (2020) |
| Xtriage (Phenix) | : | 1.13 |
| EDS | : | 2.28 |
| Percentile statistics | : | 20191225.v01 (using entries in the PDB archive December 25th 2019) |
| Refmac | : | 5.8.0158 |
| CCP4 | : | 7.0.044 (Gargrove) |
| Ideal geometry (proteins) | : | Engh & Huber (2001) |
| Ideal geometry (DNA, RNA) | : | Parkinson et al. (1996) |
| Validation Pipeline (wwPDB-VP) | : | 2.28 |

For Manuscript Review

### 1 Overall quality at a glance (i)

The following experimental techniques were used to determine the structure:
*X-RAY DIFFRACTION*

The reported resolution of this entry is 1.50 Å.

Percentile scores (ranging between 0-100) for global validation metrics of the entry are shown in the following graphic. The table shows the number of entries on which the scores are based.

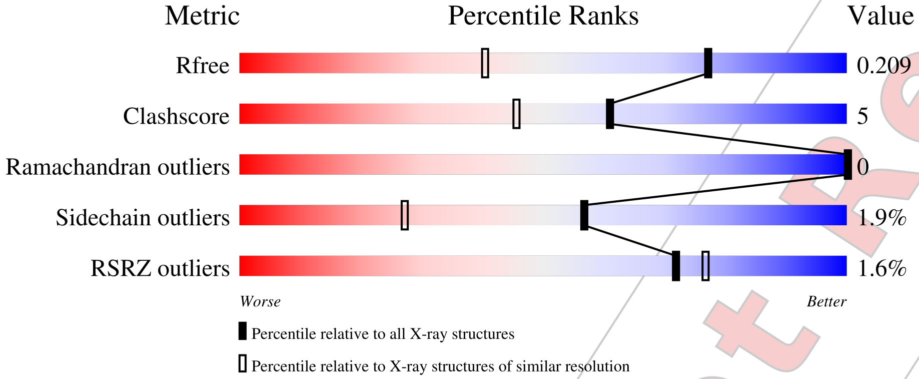


| Metric | Whole archive (#Entries) | Similar resolution (#Entries, resolution range(Å)) |
|---|---|---|
| $R_{free}$ | 130704 | 2936 (1.50-1.50) |
| Clashscore | 141614 | 3144 (1.50-1.50) |
| Ramachandran outliers | 138981 | 3066 (1.50-1.50) |
| Sidechain outliers | 138945 | 3064 (1.50-1.50) |
| RSRZ outliers | 127900 | 2884 (1.50-1.50) |

The table below summarises the geometric issues observed across the polymeric chains and their fit to the electron density. The red, orange, yellow and green segments of the lower bar indicate the fraction of residues that contain outliers for >=3, 2, 1 and 0 types of geometric quality criteria respectively. A grey segment represents the fraction of residues that are not modelled. The numeric value for each fraction is indicated below the corresponding segment, with a dot representing fractions <=5% The upper red bar (where present) indicates the fraction of residues that have poor fit to the electron density. The numeric value is given above the bar.

| Mol | Chain | Length | Quality of chain |
|---|---|---|---|
| 1 | A | 129 | 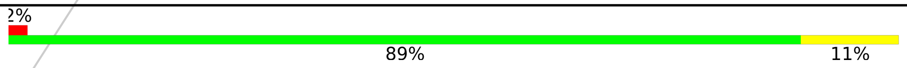 |

For Manuscript Review

#### 2 Entry composition (i)

There are 5 unique types of molecules in this entry. The entry contains 1164 atoms, of which 0 are hydrogens and 0 are deuteriums.

In the tables below, the ZeroOcc column contains the number of atoms modelled with zero occupancy, the AltConf column contains the number of residues with at least one atom in alternate conformation and the Trace column contains the number of residues modelled with at most 2 atoms.

- Molecule 1 is a protein called Lysozyme C.

| Mol | Chain | Residues | Atoms | | | | | ZeroOcc | AltConf | Trace |
|---|---|---|---|---|---|---|---|---|---|---|
| | | | Total | C | N | O | S | | | |
| 1 | A | 129 | 1001 | 613 | 193 | 185 | 10 | 0 | 0 | 0 |

- Molecule 2 is SODIUM ION (three-letter code: NA) (formula: Na).

| Mol | Chain | Residues | Atoms | | ZeroOcc | AltConf |
|---|---|---|---|---|---|---|
| | | | Total | Na | | |
| 2 | A | 1 | 1 | 1 | 0 | 0 |

- Molecule 3 is CHLORIDE ION (three-letter code: CL) (formula: Cl).

| Mol | Chain | Residues | Atoms | | ZeroOcc | AltConf |
|---|---|---|---|---|---|---|
| | | | Total | Cl | | |
| 3 | A | 7 | 7 | 7 | 0 | 0 |

- Molecule 4 is GLYCEROL (three-letter code: GOL) (formula: $C_3H_8O_3$).

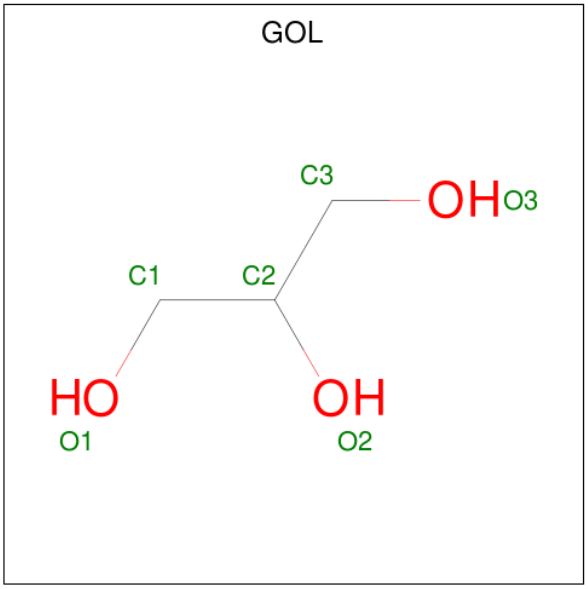


For Manuscript Review

| Mol | Chain | Residues | Atoms | ZeroOcc | AltConf |
|---|---|---|---|---|---|
| 4 | A | 1 | Total C O<br>6 3 3 | 0 | 0 |
| 4 | A | 1 | Total C O<br>6 3 3 | 0 | 0 |

- Molecule 5 is water.

| Mol | Chain | Residues | Atoms | ZeroOcc | AltConf |
|---|---|---|---|---|---|
| 5 | A | 143 | Total O<br>143 143 | 0 | 0 |

For Manuscript Review

### 3 Residue-property plots (i)

These plots are drawn for all protein, RNA, DNA and oligosaccharide chains in the entry. The first graphic for a chain summarises the proportions of the various outlier classes displayed in the second graphic. The second graphic shows the sequence view annotated by issues in geometry and electron density. Residues are color-coded according to the number of geometric quality criteria for which they contain at least one outlier: green = 0, yellow = 1, orange = 2 and red = 3 or more. A red dot above a residue indicates a poor fit to the electron density (RSRZ > 2). Stretches of 2 or more consecutive residues without any outlier are shown as a green connector. Residues present in the sample, but not in the model, are shown in grey.

• Molecule 1: Lysozyme C

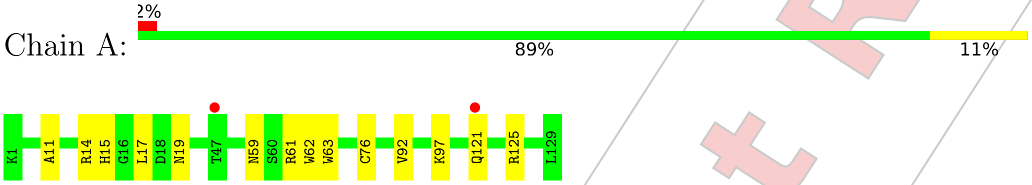


For Manuscript Review

### 4 Data and refinement statistics (i)

| Property | Value | Source |
|---|---|---|
| Space group | P 43 21 2 | Depositor |
| Cell constants<br>a, b, c, $\alpha$, $\beta$, $\gamma$ | 79.11Å 79.11Å 37.18Å<br>90.00° 90.00° 90.00° | Depositor |
| Resolution (Å) | 22.36 – 1.50<br>22.35 – 1.50 | Depositor<br>EDS |
| % Data completeness<br>(in resolution range) | 99.5 (22.36-1.50)<br>99.5 (22.35-1.50) | Depositor<br>EDS |
| $R_{merge}$ | (Not available) | Depositor |
| $R_{sym}$ | (Not available) | Depositor |
| $< I/\sigma(I) >$ [1] | 1.70 (at 1.49Å) | Xtriage |
| Refinement program | REFMAC 5.8.0267 | Depositor |
| R, $R_{free}$ | 0.167 , 0.202<br>0.175 , 0.209 | Depositor<br>DCC |
| $R_{free}$ test set | 997 reflections (5.11%) | wwPDB-VP |
| Wilson B-factor (Å$^2$) | 8.5 | Xtriage |
| Anisotropy | 0.014 | Xtriage |
| Bulk solvent $k_{sol}$(e/Å$^3$), $B_{sol}$(Å$^2$) | 0.39 , 40.6 | EDS |
| L-test for twinning[2] | $< \|L\| > = 0.49$, $< L^2 > = 0.32$ | Xtriage |
| Estimated twinning fraction | No twinning to report. | Xtriage |
| $F_o$,$F_c$ correlation | 0.95 | EDS |
| Total number of atoms | 1164 | wwPDB-VP |
| Average B, all atoms (Å$^2$) | 10.0 | wwPDB-VP |

Xtriage's analysis on translational NCS is as follows: *The largest off-origin peak in the Patterson function is 8.13% of the height of the origin peak. No significant pseudotranslation is detected.*

[1] Intensities estimated from amplitudes.

[2] Theoretical values of $< |L| >$, $< L^2 >$ for acentric reflections are 0.5, 0.333 respectively for untwinned datasets, and 0.375, 0.2 for perfectly twinned datasets.

For Manuscript Review

### 5 Model quality (i)

#### 5.1 Standard geometry (i)

Bond lengths and bond angles in the following residue types are not validated in this section: GOL, CL, NA

The Z score for a bond length (or angle) is the number of standard deviations the observed value is removed from the expected value. A bond length (or angle) with $|Z| > 5$ is considered an outlier worth inspection. RMSZ is the root-mean-square of all Z scores of the bond lengths (or angles).

| Mol | Chain | Bond lengths | | Bond angles | |
|---|---|---|---|---|---|
| | | RMSZ | #$\|Z\|$ >5 | RMSZ | #$\|Z\|$ >5 |
| 1 | A | 0.72 | 0/1021 | 0.93 | 0/1379 |

Chiral center outliers are detected by calculating the chiral volume of a chiral center and verifying if the center is modelled as a planar moiety or with the opposite hand.A planarity outlier is detected by checking planarity of atoms in a peptide group, atoms in a mainchain group or atoms of a sidechain that are expected to be planar.

| Mol | Chain | #Chirality outliers | #Planarity outliers |
|---|---|---|---|
| 1 | A | 0 | 1 |

There are no bond length outliers.

There are no bond angle outliers.

There are no chirality outliers.

All (1) planarity outliers are listed below:

| Mol | Chain | Res | Type | Group |
|---|---|---|---|---|
| 1 | A | 62 | TRP | Peptide |

#### 5.2 Too-close contacts (i)

In the following table, the Non-H and H(model) columns list the number of non-hydrogen atoms and hydrogen atoms in the chain respectively. The H(added) column lists the number of hydrogen atoms added and optimized by MolProbity. The Clashes column lists the number of clashes within the asymmetric unit, whereas Symm-Clashes lists symmetry-related clashes.

| Mol | Chain | Non-H | H(model) | H(added) | Clashes | Symm-Clashes |
|---|---|---|---|---|---|---|
| 1 | A | 1001 | 0 | 958 | 8 | 0 |
| 2 | A | 1 | 0 | 0 | 0 | 0 |

*Continued on next page...*

For Manuscript Review

*Continued from previous page...*

| Mol | Chain | Non-H | H(model) | H(added) | Clashes | Symm-Clashes |
|---|---|---|---|---|---|---|
| 3 | A | 7 | 0 | 0 | 1 | 0 |
| 4 | A | 12 | 0 | 16 | 0 | 0 |
| 5 | A | 143 | 0 | 0 | 1 | 0 |
| All | All | 1164 | 0 | 974 | 9 | 0 |

The all-atom clashscore is defined as the number of clashes found per 1000 atoms (including hydrogen atoms). The all-atom clashscore for this structure is 5.

All (9) close contacts within the same asymmetric unit are listed below, sorted by their clash magnitude.

| Atom-1 | Atom-2 | Interatomic distance (Å) | Clash overlap (Å) |
|---|---|---|---|
| 3:A:204:CL:CL | 5:A:396:HOH:O | 2.35 | 0.82 |
| 1:A:121:GLN:OE1 | 1:A:125:ARG:NH1 | 2.33 | 0.61 |
| 1:A:59:ASN:ND2 | 1:A:61:ARG:H | 2.02 | 0.56 |
| 1:A:121:GLN:OE1 | 1:A:125:ARG:CZ | 2.58 | 0.52 |
| 1:A:11:ALA:O | 1:A:15:HIS:HD2 | 1.96 | 0.49 |
| 1:A:17:LEU:HG | 1:A:92:VAL:HG13 | 1.94 | 0.48 |
| 1:A:59:ASN:ND2 | 1:A:61:ARG:HB3 | 2.28 | 0.48 |
| 1:A:63:TRP:O | 1:A:76:CYS:HB2 | 2.17 | 0.44 |
| 1:A:19:ASN:HD22 | 1:A:19:ASN:HA | 1.71 | 0.41 |

There are no symmetry-related clashes.

#### 5.3 Torsion angles (i)

##### 5.3.1 Protein backbone (i)

In the following table, the Percentiles column shows the percent Ramachandran outliers of the chain as a percentile score with respect to all X-ray entries followed by that with respect to entries of similar resolution.

The Analysed column shows the number of residues for which the backbone conformation was analysed, and the total number of residues.

| Mol | Chain | Analysed | Favoured | Allowed | Outliers | Percentiles | |
|---|---|---|---|---|---|---|---|
| 1 | A | 127/129 (98%) | 126 (99%) | 1 (1%) | 0 | 100 | 100 |

There are no Ramachandran outliers to report.

For Manuscript Review

##### 5.3.2 Protein sidechains (i)

In the following table, the Percentiles column shows the percent sidechain outliers of the chain as a percentile score with respect to all X-ray entries followed by that with respect to entries of similar resolution.

The Analysed column shows the number of residues for which the sidechain conformation was analysed, and the total number of residues.

| Mol | Chain | Analysed | Rotameric | Outliers | Percentiles | |
|---|---|---|---|---|---|---|
| 1 | A | 105/105 (100%) | 103 (98%) | 2 (2%) | 57 | 27 |

All (2) residues with a non-rotameric sidechain are listed below:

| Mol | Chain | Res | Type |
|---|---|---|---|
| 1 | A | 14 | ARG |
| 1 | A | 97 | LYS |

Sometimes sidechains can be flipped to improve hydrogen bonding and reduce clashes. All (4) such sidechains are listed below:

| Mol | Chain | Res | Type |
|---|---|---|---|
| 1 | A | 15 | HIS |
| 1 | A | 19 | ASN |
| 1 | A | 59 | ASN |
| 1 | A | 65 | ASN |

##### 5.3.3 RNA (i)

There are no RNA molecules in this entry.

#### 5.4 Non-standard residues in protein, DNA, RNA chains (i)

There are no non-standard protein/DNA/RNA residues in this entry.

#### 5.5 Carbohydrates (i)

There are no monosaccharides in this entry.

#### 5.6 Ligand geometry (i)

Of 10 ligands modelled in this entry, 8 are monoatomic - leaving 2 for Mogul analysis.

For Manuscript Review

In the following table, the Counts columns list the number of bonds (or angles) for which Mogul statistics could be retrieved, the number of bonds (or angles) that are observed in the model and the number of bonds (or angles) that are defined in the Chemical Component Dictionary. The Link column lists molecule types, if any, to which the group is linked. The Z score for a bond length (or angle) is the number of standard deviations the observed value is removed from the expected value. A bond length (or angle) with $|Z| > 2$ is considered an outlier worth inspection. RMSZ is the root-mean-square of all Z scores of the bond lengths (or angles).

| Mol | Type | Chain | Res | Link | Bond lengths | | | Bond angles | | |
|---|---|---|---|---|---|---|---|---|---|---|
| | | | | | Counts | RMSZ | $\#\|Z\| > 2$ | Counts | RMSZ | $\#\|Z\| > 2$ |
| 4 | GOL | A | 210 | - | 5,5,5 | 0.11 | 0 | 5,5,5 | 0.23 | 0 |
| 4 | GOL | A | 209 | - | 5,5,5 | 0.20 | 0 | 5,5,5 | 0.42 | 0 |

In the following table, the Chirals column lists the number of chiral outliers, the number of chiral centers analysed, the number of these observed in the model and the number defined in the Chemical Component Dictionary. Similar counts are reported in the Torsion and Rings columns. '-' means no outliers of that kind were identified.

| Mol | Type | Chain | Res | Link | Chirals | Torsions | Rings |
|---|---|---|---|---|---|---|---|
| 4 | GOL | A | 210 | - | - | 0/4/4/4 | - |
| 4 | GOL | A | 209 | - | - | 0/4/4/4 | - |

There are no bond length outliers.

There are no bond angle outliers.

There are no chirality outliers.

There are no torsion outliers.

There are no ring outliers.

No monomer is involved in short contacts.

#### 5.7 Other polymers (i)

There are no such residues in this entry.

#### 5.8 Polymer linkage issues (i)

There are no chain breaks in this entry.

For Manuscript Review

### 6 Fit of model and data (i)

#### 6.1 Protein, DNA and RNA chains (i)

In the following table, the column labelled '#RSRZ> 2' contains the number (and percentage) of RSRZ outliers, followed by percent RSRZ outliers for the chain as percentile scores relative to all X-ray entries and entries of similar resolution. The OWAB column contains the minimum, median, $95^{th}$ percentile and maximum values of the occupancy-weighted average B-factor per residue. The column labelled 'Q< 0.9' lists the number of (and percentage) of residues with an average occupancy less than 0.9.

| Mol | Chain | Analysed | <RSRZ> | #RSRZ>2 | | | OWAB(Å$^2$) | Q<0.9 |
|---|---|---|---|---|---|---|---|---|
| 1 | A | 129/129 (100%) | -0.01 | 2 (1%) | 72 | 77 | 5, 8, 17, 23 | 0 |

All (2) RSRZ outliers are listed below:

| Mol | Chain | Res | Type | RSRZ |
|---|---|---|---|---|
| 1 | A | 121 | GLN | 2.1 |
| 1 | A | 47 | THR | 2.1 |

#### 6.2 Non-standard residues in protein, DNA, RNA chains (i)

There are no non-standard protein/DNA/RNA residues in this entry.

#### 6.3 Carbohydrates (i)

There are no monosaccharides in this entry.

#### 6.4 Ligands (i)

In the following table, the Atoms column lists the number of modelled atoms in the group and the number defined in the chemical component dictionary. The B-factors column lists the minimum, median, $95^{th}$ percentile and maximum values of B factors of atoms in the group. The column labelled 'Q< 0.9' lists the number of atoms with occupancy less than 0.9.

| Mol | Type | Chain | Res | Atoms | RSCC | RSR | B-factors(Å$^2$) | Q<0.9 |
|---|---|---|---|---|---|---|---|---|
| 4 | GOL | A | 209 | 6/6 | 0.80 | 0.15 | 19,22,22,23 | 0 |
| 4 | GOL | A | 210 | 6/6 | 0.88 | 0.15 | 16,17,18,20 | 0 |
| 3 | CL | A | 208 | 1/1 | 0.93 | 0.07 | 19,19,19,19 | 0 |
| 3 | CL | A | 207 | 1/1 | 0.97 | 0.07 | 18,18,18,18 | 0 |
| 3 | CL | A | 204 | 1/1 | 0.97 | 0.10 | 16,16,16,16 | 0 |

*Continued on next page...*

For Manuscript Review

*Continued from previous page...*

| Mol | Type | Chain | Res | Atoms | RSCC | RSR | B-factors($Å^2$) | Q<0.9 |
|---|---|---|---|---|---|---|---|---|
| 3 | CL | A | 206 | 1/1 | 0.98 | 0.13 | 18,18,18,18 | 0 |
| 3 | CL | A | 205 | 1/1 | 0.99 | 0.05 | 14,14,14,14 | 0 |
| 3 | CL | A | 203 | 1/1 | 0.99 | 0.04 | 8,8,8,8 | 0 |
| 3 | CL | A | 202 | 1/1 | 0.99 | 0.05 | 8,8,8,8 | 0 |
| 2 | NA | A | 201 | 1/1 | 1.00 | 0.04 | 6,6,6,6 | 0 |

#### 6.5 Other polymers (i)

There are no such residues in this entry.

For Manuscript Review

### Full wwPDB X-ray Structure Validation Report (i)

Jun 8, 2022 – 03:48 PM EDT

| | | |
|---|---|---|
| PDB ID | : | 8D77 |
| Title | : | Crystal structure of hen egg white lysozyme at 200 Kelvin (Duplicate) |
| Deposited on | : | 2022-06-07 |
| Resolution | : | 1.50 Å(reported) |

**This wwPDB validation report is for manuscript review**

This is a Full wwPDB X-ray Structure Validation Report.

This report is produced by the wwPDB biocuration pipeline after annotation of the structure.

We welcome your comments at **
A user guide is available at
https://www.wwpdb.org/validation/2017/XrayValidationReportHelp
with specific help available everywhere you see the (i) symbol.

The following versions of software and data (see references (i)) were used in the production of this report:

| | | |
|---|---|---|
| MolProbity | : | 4.02b-467 |
| Mogul | : | 1.8.5 (274361), CSD as541be (2020) |
| Xtriage (Phenix) | : | 1.13 |
| EDS | : | 2.28 |
| Percentile statistics | : | 20191225.v01 (using entries in the PDB archive December 25th 2019) |
| Refmac | : | 5.8.0158 |
| CCP4 | : | 7.0.044 (Gargrove) |
| Ideal geometry (proteins) | : | Engh & Huber (2001) |
| Ideal geometry (DNA, RNA) | : | Parkinson et al. (1996) |
| Validation Pipeline (wwPDB-VP) | : | 2.28 |

For Manuscript Review

### 1 Overall quality at a glance (i)

The following experimental techniques were used to determine the structure:
*X-RAY DIFFRACTION*

The reported resolution of this entry is 1.50 Å.

Percentile scores (ranging between 0-100) for global validation metrics of the entry are shown in the following graphic. The table shows the number of entries on which the scores are based.

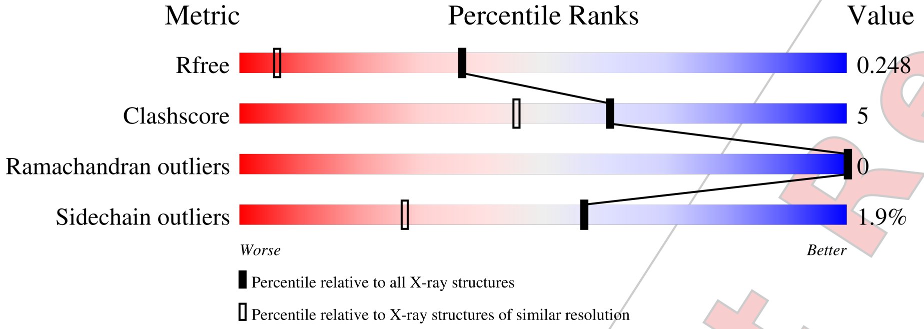


| Metric | Whole archive (#Entries) | Similar resolution (#Entries, resolution range(Å)) |
|---|---|---|
| $R_{free}$ | 130704 | 2936 (1.50-1.50) |
| Clashscore | 141614 | 3144 (1.50-1.50) |
| Ramachandran outliers | 138981 | 3066 (1.50-1.50) |
| Sidechain outliers | 138945 | 3064 (1.50-1.50) |

The table below summarises the geometric issues observed across the polymeric chains and their fit to the electron density. The red, orange, yellow and green segments of the lower bar indicate the fraction of residues that contain outliers for >=3, 2, 1 and 0 types of geometric quality criteria respectively. A grey segment represents the fraction of residues that are not modelled. The numeric value for each fraction is indicated below the corresponding segment, with a dot representing fractions <=5%

| Mol | Chain | Length | Quality of chain |
|---|---|---|---|
| 1 | A | 129 | 89% 11% |

For Manuscript Review

#### 2 Entry composition (i)

There are 5 unique types of molecules in this entry. The entry contains 1164 atoms, of which 0 are hydrogens and 0 are deuteriums.

In the tables below, the ZeroOcc column contains the number of atoms modelled with zero occupancy, the AltConf column contains the number of residues with at least one atom in alternate conformation and the Trace column contains the number of residues modelled with at most 2 atoms.

- Molecule 1 is a protein called Lysozyme C.

| Mol | Chain | Residues | Atoms | | | | | ZeroOcc | AltConf | Trace |
|---|---|---|---|---|---|---|---|---|---|---|
| | | | Total | C | N | O | S | | | |
| 1 | A | 129 | 1001 | 613 | 193 | 185 | 10 | 0 | 0 | 0 |

- Molecule 2 is SODIUM ION (three-letter code: NA) (formula: Na).

| Mol | Chain | Residues | Atoms | | ZeroOcc | AltConf |
|---|---|---|---|---|---|---|
| | | | Total | Na | | |
| 2 | A | 1 | 1 | 1 | 0 | 0 |

- Molecule 3 is CHLORIDE ION (three-letter code: CL) (formula: Cl).

| Mol | Chain | Residues | Atoms | | ZeroOcc | AltConf |
|---|---|---|---|---|---|---|
| | | | Total | Cl | | |
| 3 | A | 7 | 7 | 7 | 0 | 0 |

- Molecule 4 is GLYCEROL (three-letter code: GOL) (formula: $C_3H_8O_3$).

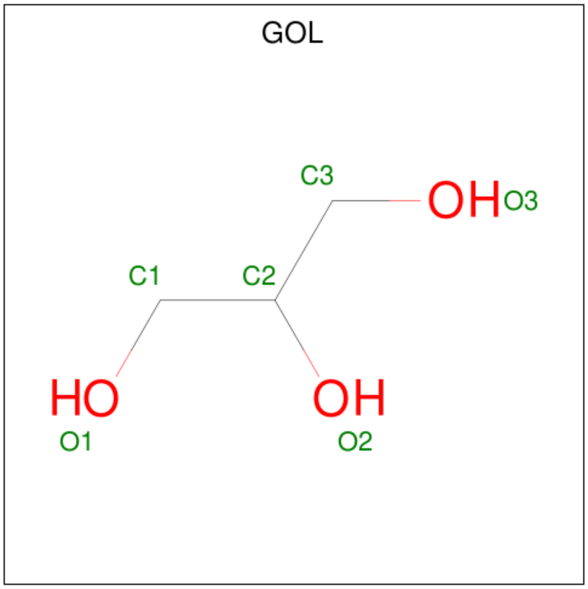


For Manuscript Review

| Mol | Chain | Residues | Atoms | ZeroOcc | AltConf |
|---|---|---|---|---|---|
| 4 | A | 1 | Total C O<br>6 3 3 | 0 | 0 |
| 4 | A | 1 | Total C O<br>6 3 3 | 0 | 0 |

- Molecule 5 is water.

| Mol | Chain | Residues | Atoms | ZeroOcc | AltConf |
|---|---|---|---|---|---|
| 5 | A | 143 | Total O<br>143 143 | 0 | 0 |

For Manuscript Review

### 3 Residue-property plots (i)

These plots are drawn for all protein, RNA, DNA and oligosaccharide chains in the entry. The first graphic for a chain summarises the proportions of the various outlier classes displayed in the second graphic. The second graphic shows the sequence view annotated by issues in geometry. Residues are color-coded according to the number of geometric quality criteria for which they contain at least one outlier: green = 0, yellow = 1, orange = 2 and red = 3 or more. Stretches of 2 or more consecutive residues without any outlier are shown as a green connector. Residues present in the sample, but not in the model, are shown in grey.

- Molecule 1: Lysozyme C

Chain A:

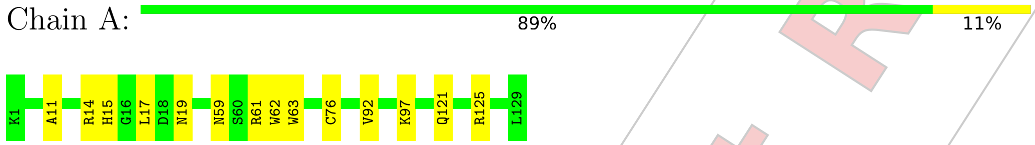


For Manuscript Review

### 4 Data and refinement statistics (i)

| Property | Value | Source |
|---|---|---|
| Space group | P 43 21 2 | Depositor |
| Cell constants<br>a, b, c, $\alpha$, $\beta$, $\gamma$ | 79.11Å 79.11Å 37.18Å<br>90.00° 90.00° 90.00° | Depositor |
| Resolution (Å) | 22.29 – 1.50<br>22.35 – 1.50 | Depositor<br>EDS |
| % Data completeness<br>(in resolution range) | 99.5 (22.29-1.50)<br>99.3 (22.35-1.50) | Depositor<br>EDS |
| $R_{merge}$ | (Not available) | Depositor |
| $R_{sym}$ | (Not available) | Depositor |
| $< I/\sigma(I) >$ [1] | 2.49 (at 1.50Å) | Xtriage |
| Refinement program | REFMAC 5.8.0267 | Depositor |
| R, $R_{free}$ | 0.167 , 0.202<br>0.230 , 0.248 | Depositor<br>DCC |
| $R_{free}$ test set | 990 reflections (5.12%) | wwPDB-VP |
| Wilson B-factor (Å$^2$) | 11.0 | Xtriage |
| Anisotropy | 0.094 | Xtriage |
| Bulk solvent $k_{sol}$(e/Å$^3$), $B_{sol}$(Å$^2$) | 0.37 , 38.0 | EDS |
| L-test for twinning[2] | $< \|L\| > = 0.48$, $< L^2 > = 0.31$ | Xtriage |
| Estimated twinning fraction | No twinning to report. | Xtriage |
| $F_o$,$F_c$ correlation | 0.93 | EDS |
| Total number of atoms | 1164 | wwPDB-VP |
| Average B, all atoms (Å$^2$) | 10.0 | wwPDB-VP |

Xtriage's analysis on translational NCS is as follows: *The largest off-origin peak in the Patterson function is 8.36% of the height of the origin peak. No significant pseudotranslation is detected.*

[1]Intensities estimated from amplitudes.

[2]Theoretical values of $< |L| >$, $< L^2 >$ for acentric reflections are 0.5, 0.333 respectively for untwinned datasets, and 0.375, 0.2 for perfectly twinned datasets.

For Manuscript Review

### 5 Model quality (i)

#### 5.1 Standard geometry (i)

Bond lengths and bond angles in the following residue types are not validated in this section: CL, NA, GOL

The Z score for a bond length (or angle) is the number of standard deviations the observed value is removed from the expected value. A bond length (or angle) with $|Z| > 5$ is considered an outlier worth inspection. RMSZ is the root-mean-square of all Z scores of the bond lengths (or angles).

| Mol | Chain | Bond lengths | | Bond angles | |
|---|---|---|---|---|---|
| | | RMSZ | $\#|Z| >5$ | RMSZ | $\#|Z| >5$ |
| 1 | A | 0.72 | 0/1021 | 0.93 | 0/1379 |

Chiral center outliers are detected by calculating the chiral volume of a chiral center and verifying if the center is modelled as a planar moiety or with the opposite hand.A planarity outlier is detected by checking planarity of atoms in a peptide group, atoms in a mainchain group or atoms of a sidechain that are expected to be planar.

| Mol | Chain | #Chirality outliers | #Planarity outliers |
|---|---|---|---|
| 1 | A | 0 | 1 |

There are no bond length outliers.

There are no bond angle outliers.

There are no chirality outliers.

All (1) planarity outliers are listed below:

| Mol | Chain | Res | Type | Group |
|---|---|---|---|---|
| 1 | A | 62 | TRP | Peptide |

#### 5.2 Too-close contacts (i)

In the following table, the Non-H and H(model) columns list the number of non-hydrogen atoms and hydrogen atoms in the chain respectively. The H(added) column lists the number of hydrogen atoms added and optimized by MolProbity. The Clashes column lists the number of clashes within the asymmetric unit, whereas Symm-Clashes lists symmetry-related clashes.

| Mol | Chain | Non-H | H(model) | H(added) | Clashes | Symm-Clashes |
|---|---|---|---|---|---|---|
| 1 | A | 1001 | 0 | 958 | 8 | 0 |
| 2 | A | 1 | 0 | 0 | 0 | 0 |

*Continued on next page...*

For Manuscript Review

*Continued from previous page...*

| Mol | Chain | Non-H | H(model) | H(added) | Clashes | Symm-Clashes |
|---|---|---|---|---|---|---|
| 3 | A | 7 | 0 | 0 | 1 | 0 |
| 4 | A | 12 | 0 | 16 | 0 | 0 |
| 5 | A | 143 | 0 | 0 | 1 | 0 |
| All | All | 1164 | 0 | 974 | 9 | 0 |

The all-atom clashscore is defined as the number of clashes found per 1000 atoms (including hydrogen atoms). The all-atom clashscore for this structure is 5.

All (9) close contacts within the same asymmetric unit are listed below, sorted by their clash magnitude.

| Atom-1 | Atom-2 | Interatomic distance (Å) | Clash overlap (Å) |
|---|---|---|---|
| 3:A:204:CL:CL | 5:A:396:HOH:O | 2.35 | 0.82 |
| 1:A:121:GLN:OE1 | 1:A:125:ARG:NH1 | 2.33 | 0.61 |
| 1:A:59:ASN:ND2 | 1:A:61:ARG:H | 2.02 | 0.56 |
| 1:A:121:GLN:OE1 | 1:A:125:ARG:CZ | 2.58 | 0.52 |
| 1:A:11:ALA:O | 1:A:15:HIS:HD2 | 1.96 | 0.49 |
| 1:A:17:LEU:HG | 1:A:92:VAL:HG13 | 1.94 | 0.48 |
| 1:A:59:ASN:ND2 | 1:A:61:ARG:HB3 | 2.28 | 0.48 |
| 1:A:63:TRP:O | 1:A:76:CYS:HB2 | 2.17 | 0.44 |
| 1:A:19:ASN:HD22 | 1:A:19:ASN:HA | 1.71 | 0.41 |

There are no symmetry-related clashes.

#### 5.3 Torsion angles (i)

##### 5.3.1 Protein backbone (i)

In the following table, the Percentiles column shows the percent Ramachandran outliers of the chain as a percentile score with respect to all X-ray entries followed by that with respect to entries of similar resolution.

The Analysed column shows the number of residues for which the backbone conformation was analysed, and the total number of residues.

| Mol | Chain | Analysed | Favoured | Allowed | Outliers | Percentiles | |
|---|---|---|---|---|---|---|---|
| 1 | A | 127/129 (98%) | 126 (99%) | 1 (1%) | 0 | 100 | 100 |

There are no Ramachandran outliers to report.

For Manuscript Review

##### 5.3.2 Protein sidechains (i)

In the following table, the Percentiles column shows the percent sidechain outliers of the chain as a percentile score with respect to all X-ray entries followed by that with respect to entries of similar resolution.

The Analysed column shows the number of residues for which the sidechain conformation was analysed, and the total number of residues.

| Mol | Chain | Analysed | Rotameric | Outliers | Percentiles | |
|---|---|---|---|---|---|---|
| 1 | A | 105/105 (100%) | 103 (98%) | 2 (2%) | 57 | 27 |

All (2) residues with a non-rotameric sidechain are listed below:

| Mol | Chain | Res | Type |
|---|---|---|---|
| 1 | A | 14 | ARG |
| 1 | A | 97 | LYS |

Sometimes sidechains can be flipped to improve hydrogen bonding and reduce clashes. All (4) such sidechains are listed below:

| Mol | Chain | Res | Type |
|---|---|---|---|
| 1 | A | 15 | HIS |
| 1 | A | 19 | ASN |
| 1 | A | 59 | ASN |
| 1 | A | 65 | ASN |

##### 5.3.3 RNA (i)

There are no RNA molecules in this entry.

#### 5.4 Non-standard residues in protein, DNA, RNA chains (i)

There are no non-standard protein/DNA/RNA residues in this entry.

#### 5.5 Carbohydrates (i)

There are no monosaccharides in this entry.

#### 5.6 Ligand geometry (i)

Of 10 ligands modelled in this entry, 8 are monoatomic - leaving 2 for Mogul analysis.

For Manuscript Review

In the following table, the Counts columns list the number of bonds (or angles) for which Mogul statistics could be retrieved, the number of bonds (or angles) that are observed in the model and the number of bonds (or angles) that are defined in the Chemical Component Dictionary. The Link column lists molecule types, if any, to which the group is linked. The Z score for a bond length (or angle) is the number of standard deviations the observed value is removed from the expected value. A bond length (or angle) with $|Z| > 2$ is considered an outlier worth inspection. RMSZ is the root-mean-square of all Z scores of the bond lengths (or angles).

| Mol | Type | Chain | Res | Link | Bond lengths | | | Bond angles | | |
|---|---|---|---|---|---|---|---|---|---|---|
| | | | | | Counts | RMSZ | $\#|Z|>2$ | Counts | RMSZ | $\#|Z|>2$ |
| 4 | GOL | A | 209 | - | 5,5,5 | 0.20 | 0 | 5,5,5 | 0.42 | 0 |
| 4 | GOL | A | 210 | - | 5,5,5 | 0.11 | 0 | 5,5,5 | 0.23 | 0 |

In the following table, the Chirals column lists the number of chiral outliers, the number of chiral centers analysed, the number of these observed in the model and the number defined in the Chemical Component Dictionary. Similar counts are reported in the Torsion and Rings columns. '-' means no outliers of that kind were identified.

| Mol | Type | Chain | Res | Link | Chirals | Torsions | Rings |
|---|---|---|---|---|---|---|---|
| 4 | GOL | A | 209 | - | - | 0/4/4/4 | - |
| 4 | GOL | A | 210 | - | - | 0/4/4/4 | - |

There are no bond length outliers.

There are no bond angle outliers.

There are no chirality outliers.

There are no torsion outliers.

There are no ring outliers.

No monomer is involved in short contacts.

#### 5.7 Other polymers (i)

There are no such residues in this entry.

#### 5.8 Polymer linkage issues (i)

There are no chain breaks in this entry.

For Manuscript Review

### 6 Fit of model and data (i)

#### 6.1 Protein, DNA and RNA chains (i)

Unable to reproduce the depositors R factor - this section is therefore empty.

#### 6.2 Non-standard residues in protein, DNA, RNA chains (i)

Unable to reproduce the depositors R factor - this section is therefore empty.

#### 6.3 Carbohydrates (i)

Unable to reproduce the depositors R factor - this section is therefore empty.

#### 6.4 Ligands (i)

Unable to reproduce the depositors R factor - this section is therefore empty.

#### 6.5 Other polymers (i)

Unable to reproduce the depositors R factor - this section is therefore empty.

For Manuscript Review

### Full wwPDB X-ray Structure Validation Report (i)

Jun 8, 2022 – 03:49 PM EDT

| | | |
|---|---|---|
| PDB ID | : | 8D7A |
| Title | : | Crystal structure of hen egg white lysozyme at 200 Kelvin (Triplicate) |
| Deposited on | : | 2022-06-07 |
| Resolution | : | 1.50 Å(reported) |

**This wwPDB validation report is for manuscript review**

This is a Full wwPDB X-ray Structure Validation Report.

This report is produced by the wwPDB biocuration pipeline after annotation of the structure.

We welcome your comments at **
A user guide is available at
https://www.wwpdb.org/validation/2017/XrayValidationReportHelp
with specific help available everywhere you see the (i) symbol.

The following versions of software and data (see references (i)) were used in the production of this report:

| | | |
|---|---|---|
| MolProbity | : | 4.02b-467 |
| Mogul | : | 1.8.5 (274361), CSD as541be (2020) |
| Xtriage (Phenix) | : | 1.13 |
| EDS | : | 2.28 |
| Percentile statistics | : | 20191225.v01 (using entries in the PDB archive December 25th 2019) |
| Refmac | : | 5.8.0158 |
| CCP4 | : | 7.0.044 (Gargrove) |
| Ideal geometry (proteins) | : | Engh & Huber (2001) |
| Ideal geometry (DNA, RNA) | : | Parkinson et al. (1996) |
| Validation Pipeline (wwPDB-VP) | : | 2.28 |

For Manuscript Review

### 1 Overall quality at a glance (i)

The following experimental techniques were used to determine the structure:
*X-RAY DIFFRACTION*

The reported resolution of this entry is 1.50 Å.

Percentile scores (ranging between 0-100) for global validation metrics of the entry are shown in the following graphic. The table shows the number of entries on which the scores are based.

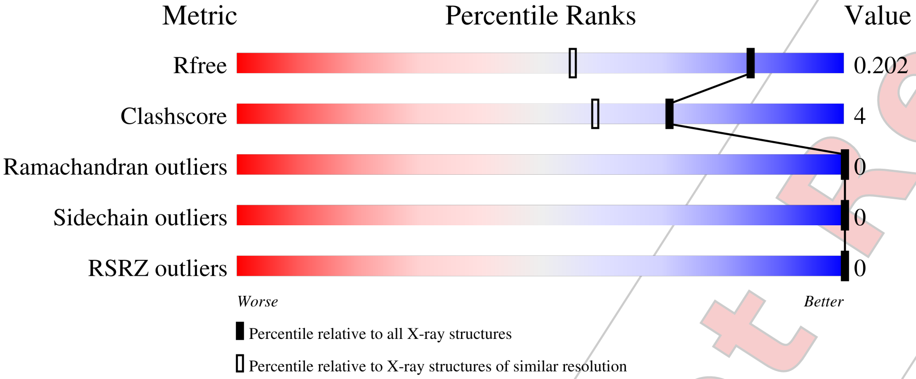


| Metric | Whole archive (#Entries) | Similar resolution (#Entries, resolution range(Å)) |
|---|---|---|
| $R_{free}$ | 130704 | 2936 (1.50-1.50) |
| Clashscore | 141614 | 3144 (1.50-1.50) |
| Ramachandran outliers | 138981 | 3066 (1.50-1.50) |
| Sidechain outliers | 138945 | 3064 (1.50-1.50) |
| RSRZ outliers | 127900 | 2884 (1.50-1.50) |

The table below summarises the geometric issues observed across the polymeric chains and their fit to the electron density. The red, orange, yellow and green segments of the lower bar indicate the fraction of residues that contain outliers for >=3, 2, 1 and 0 types of geometric quality criteria respectively. A grey segment represents the fraction of residues that are not modelled. The numeric value for each fraction is indicated below the corresponding segment, with a dot representing fractions <=5% The upper red bar (where present) indicates the fraction of residues that have poor fit to the electron density. The numeric value is given above the bar.

| Mol | Chain | Length | Quality of chain |
|---|---|---|---|
| 1 | A | 129 | 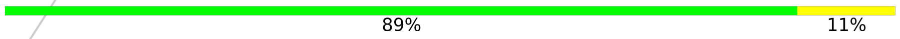 |

For Manuscript Review

### 2 Entry composition (i)

There are 5 unique types of molecules in this entry. The entry contains 1163 atoms, of which 0 are hydrogens and 0 are deuteriums.

In the tables below, the ZeroOcc column contains the number of atoms modelled with zero occupancy, the AltConf column contains the number of residues with at least one atom in alternate conformation and the Trace column contains the number of residues modelled with at most 2 atoms.

- Molecule 1 is a protein called Lysozyme C.

| Mol | Chain | Residues | Atoms | | | | | ZeroOcc | AltConf | Trace |
|---|---|---|---|---|---|---|---|---|---|---|
| | | | Total | C | N | O | S | | | |
| 1 | A | 129 | 1001 | 613 | 193 | 185 | 10 | 0 | 0 | 0 |

- Molecule 2 is SODIUM ION (three-letter code: NA) (formula: Na).

| Mol | Chain | Residues | Atoms | | ZeroOcc | AltConf |
|---|---|---|---|---|---|---|
| | | | Total | Na | | |
| 2 | A | 1 | 1 | 1 | 0 | 0 |

- Molecule 3 is CHLORIDE ION (three-letter code: CL) (formula: Cl).

| Mol | Chain | Residues | Atoms | | ZeroOcc | AltConf |
|---|---|---|---|---|---|---|
| | | | Total | Cl | | |
| 3 | A | 7 | 7 | 7 | 0 | 0 |

- Molecule 4 is GLYCEROL (three-letter code: GOL) (formula: $C_3H_8O_3$).

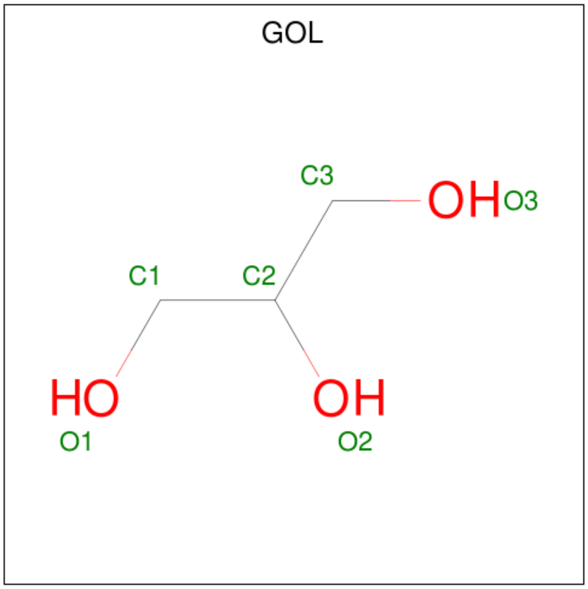


For Manuscript Review

| Mol | Chain | Residues | Atoms | ZeroOcc | AltConf |
|---|---|---|---|---|---|
| 4 | A | 1 | Total C O<br>6 3 3 | 0 | 0 |
| 4 | A | 1 | Total C O<br>6 3 3 | 0 | 0 |

- Molecule 5 is water.

| Mol | Chain | Residues | Atoms | ZeroOcc | AltConf |
|---|---|---|---|---|---|
| 5 | A | 142 | Total O<br>142 142 | 0 | 0 |

For Manuscript Review

### 3 Residue-property plots (i)

These plots are drawn for all protein, RNA, DNA and oligosaccharide chains in the entry. The first graphic for a chain summarises the proportions of the various outlier classes displayed in the second graphic. The second graphic shows the sequence view annotated by issues in geometry and electron density. Residues are color-coded according to the number of geometric quality criteria for which they contain at least one outlier: green = 0, yellow = 1, orange = 2 and red = 3 or more. A red dot above a residue indicates a poor fit to the electron density (RSRZ > 2). Stretches of 2 or more consecutive residues without any outlier are shown as a green connector. Residues present in the sample, but not in the model, are shown in grey.

- Molecule 1: Lysozyme C

Chain A:

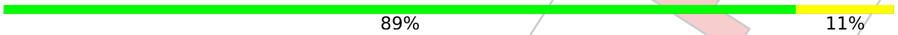


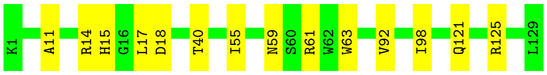


For Manuscript Review

### 4 Data and refinement statistics (i)

| Property | Value | Source |
|---|---|---|
| Space group | P 43 21 2 | Depositor |
| Cell constants<br>a, b, c, $\alpha$, $\beta$, $\gamma$ | 78.90Å 78.90Å 37.11Å<br>90.00° 90.00° 90.00° | Depositor |
| Resolution (Å) | 22.31 – 1.50<br>22.30 – 1.50 | Depositor<br>EDS |
| % Data completeness<br>(in resolution range) | 99.9 (22.31-1.50)<br>100.0 (22.30-1.50) | Depositor<br>EDS |
| $R_{merge}$ | (Not available) | Depositor |
| $R_{sym}$ | (Not available) | Depositor |
| $< I/\sigma(I) >$ [1] | 8.68 (at 1.50Å) | Xtriage |
| Refinement program | REFMAC 5.8.0267 | Depositor |
| R, $R_{free}$ | 0.168 , 0.195<br>0.177 , 0.202 | Depositor<br>DCC |
| $R_{free}$ test set | 988 reflections (5.11%) | wwPDB-VP |
| Wilson B-factor (Å$^2$) | 8.7 | Xtriage |
| Anisotropy | 0.048 | Xtriage |
| Bulk solvent $k_{sol}$(e/Å$^3$), $B_{sol}$(Å$^2$) | 0.39 , 41.2 | EDS |
| L-test for twinning[2] | $< \|L\| > = 0.49$, $< L^2 > = 0.32$ | Xtriage |
| Estimated twinning fraction | No twinning to report. | Xtriage |
| $F_o$,$F_c$ correlation | 0.95 | EDS |
| Total number of atoms | 1163 | wwPDB-VP |
| Average B, all atoms (Å$^2$) | 11.0 | wwPDB-VP |

Xtriage's analysis on translational NCS is as follows: *The largest off-origin peak in the Patterson function is 8.02% of the height of the origin peak. No significant pseudotranslation is detected.*

[1]Intensities estimated from amplitudes.

[2]Theoretical values of $< |L| >$, $< L^2 >$ for acentric reflections are 0.5, 0.333 respectively for untwinned datasets, and 0.375, 0.2 for perfectly twinned datasets.

For Manuscript Review

### 5 Model quality (i)

#### 5.1 Standard geometry (i)

Bond lengths and bond angles in the following residue types are not validated in this section: CL, GOL, NA

The Z score for a bond length (or angle) is the number of standard deviations the observed value is removed from the expected value. A bond length (or angle) with $|Z| > 5$ is considered an outlier worth inspection. RMSZ is the root-mean-square of all Z scores of the bond lengths (or angles).

| Mol | Chain | Bond lengths | | Bond angles | |
|---|---|---|---|---|---|
| | | RMSZ | $\#\|Z\| > 5$ | RMSZ | $\#\|Z\| > 5$ |
| 1 | A | 0.75 | 0/1021 | 0.98 | 2/1379 (0.1%) |

There are no bond length outliers.

All (2) bond angle outliers are listed below:

| Mol | Chain | Res | Type | Atoms | Z | Observed($^o$) | Ideal($^o$) |
|---|---|---|---|---|---|---|---|
| 1 | A | 14 | ARG | NE-CZ-NH2 | 6.24 | 123.42 | 120.30 |
| 1 | A | 18 | ASP | CB-CG-OD2 | -5.04 | 113.76 | 118.30 |

There are no chirality outliers.

There are no planarity outliers.

#### 5.2 Too-close contacts (i)

In the following table, the Non-H and H(model) columns list the number of non-hydrogen atoms and hydrogen atoms in the chain respectively. The H(added) column lists the number of hydrogen atoms added and optimized by MolProbity. The Clashes column lists the number of clashes within the asymmetric unit, whereas Symm-Clashes lists symmetry-related clashes.

| Mol | Chain | Non-H | H(model) | H(added) | Clashes | Symm-Clashes |
|---|---|---|---|---|---|---|
| 1 | A | 1001 | 0 | 958 | 7 | 0 |
| 2 | A | 1 | 0 | 0 | 0 | 0 |
| 3 | A | 7 | 0 | 0 | 0 | 0 |
| 4 | A | 12 | 0 | 16 | 0 | 0 |
| 5 | A | 142 | 0 | 0 | 0 | 0 |
| All | All | 1163 | 0 | 974 | 7 | 0 |

The all-atom clashscore is defined as the number of clashes found per 1000 atoms (including hydrogen atoms). The all-atom clashscore for this structure is 4.

For Manuscript Review

All (7) close contacts within the same asymmetric unit are listed below, sorted by their clash magnitude.

| Atom-1 | Atom-2 | Interatomic distance (Å) | Clash overlap (Å) |
|---|---|---|---|
| 1:A:121:GLN:OE1 | 1:A:125:ARG:NH2 | 2.29 | 0.64 |
| 1:A:59:ASN:ND2 | 1:A:61:ARG:H | 1.98 | 0.60 |
| 1:A:11:ALA:O | 1:A:15:HIS:HD2 | 1.90 | 0.54 |
| 1:A:17:LEU:HG | 1:A:92:VAL:HG13 | 1.91 | 0.52 |
| 1:A:59:ASN:ND2 | 1:A:61:ARG:HB3 | 2.30 | 0.47 |
| 1:A:40:THR:HA | 1:A:55:ILE:HG23 | 1.98 | 0.45 |
| 1:A:63:TRP:CE2 | 1:A:98:ILE:HG12 | 2.55 | 0.42 |

There are no symmetry-related clashes.

#### 5.3 Torsion angles (i)

##### 5.3.1 Protein backbone (i)

In the following table, the Percentiles column shows the percent Ramachandran outliers of the chain as a percentile score with respect to all X-ray entries followed by that with respect to entries of similar resolution.

The Analysed column shows the number of residues for which the backbone conformation was analysed, and the total number of residues.

| Mol | Chain | Analysed | Favoured | Allowed | Outliers | Percentiles | |
|---|---|---|---|---|---|---|---|
| 1 | A | 127/129 (98%) | 125 (98%) | 2 (2%) | 0 | 100 | 100 |

There are no Ramachandran outliers to report.

##### 5.3.2 Protein sidechains (i)

In the following table, the Percentiles column shows the percent sidechain outliers of the chain as a percentile score with respect to all X-ray entries followed by that with respect to entries of similar resolution.

The Analysed column shows the number of residues for which the sidechain conformation was analysed, and the total number of residues.

| Mol | Chain | Analysed | Rotameric | Outliers | Percentiles | |
|---|---|---|---|---|---|---|
| 1 | A | 105/105 (100%) | 105 (100%) | 0 | 100 | 100 |

There are no protein residues with a non-rotameric sidechain to report.

For Manuscript Review

Sometimes sidechains can be flipped to improve hydrogen bonding and reduce clashes. All (5) such sidechains are listed below:

| Mol | Chain | Res | Type |
|---|---|---|---|
| 1 | A | 15 | HIS |
| 1 | A | 19 | ASN |
| 1 | A | 46 | ASN |
| 1 | A | 59 | ASN |
| 1 | A | 65 | ASN |

##### 5.3.3 RNA (i)

There are no RNA molecules in this entry.

#### 5.4 Non-standard residues in protein, DNA, RNA chains (i)

There are no non-standard protein/DNA/RNA residues in this entry.

#### 5.5 Carbohydrates (i)

There are no monosaccharides in this entry.

#### 5.6 Ligand geometry (i)

Of 10 ligands modelled in this entry, 8 are monoatomic - leaving 2 for Mogul analysis.

In the following table, the Counts columns list the number of bonds (or angles) for which Mogul statistics could be retrieved, the number of bonds (or angles) that are observed in the model and the number of bonds (or angles) that are defined in the Chemical Component Dictionary. The Link column lists molecule types, if any, to which the group is linked. The Z score for a bond length (or angle) is the number of standard deviations the observed value is removed from the expected value. A bond length (or angle) with $|Z| > 2$ is considered an outlier worth inspection. RMSZ is the root-mean-square of all Z scores of the bond lengths (or angles).

| Mol | Type | Chain | Res | Link | Bond lengths | | | Bond angles | | |
|---|---|---|---|---|---|---|---|---|---|---|
| | | | | | Counts | RMSZ | $\#|Z| > 2$ | Counts | RMSZ | $\#|Z| > 2$ |
| 4 | GOL | A | 210 | - | 5,5,5 | 0.15 | 0 | 5,5,5 | 0.41 | 0 |
| 4 | GOL | A | 209 | - | 5,5,5 | 0.10 | 0 | 5,5,5 | 0.51 | 0 |

In the following table, the Chirals column lists the number of chiral outliers, the number of chiral centers analysed, the number of these observed in the model and the number defined in the Chemical Component Dictionary. Similar counts are reported in the Torsion and Rings columns. '-' means no outliers of that kind were identified.

For Manuscript Review

| Mol | Type | Chain | Res | Link | Chirals | Torsions | Rings |
|---|---|---|---|---|---|---|---|
| 4 | GOL | A | 210 | - | - | 0/4/4/4 | - |
| 4 | GOL | A | 209 | - | - | 0/4/4/4 | - |

There are no bond length outliers.

There are no bond angle outliers.

There are no chirality outliers.

There are no torsion outliers.

There are no ring outliers.

No monomer is involved in short contacts.

#### 5.7 Other polymers (i)

There are no such residues in this entry.

#### 5.8 Polymer linkage issues (i)

There are no chain breaks in this entry.

For Manuscript Review

### 6 Fit of model and data (i)

#### 6.1 Protein, DNA and RNA chains (i)

In the following table, the column labelled '#RSRZ> 2' contains the number (and percentage) of RSRZ outliers, followed by percent RSRZ outliers for the chain as percentile scores relative to all X-ray entries and entries of similar resolution. The OWAB column contains the minimum, median, $95^{th}$ percentile and maximum values of the occupancy-weighted average B-factor per residue. The column labelled 'Q< 0.9' lists the number of (and percentage) of residues with an average occupancy less than 0.9.

| Mol | Chain | Analysed | <RSRZ> | #RSRZ>2 | | | OWAB(Å$^2$) | Q<0.9 |
|---|---|---|---|---|---|---|---|---|
| 1 | A | 129/129 (100%) | -0.15 | 0 | 100 | 100 | 5, 8, 17, 24 | 0 |

There are no RSRZ outliers to report.

#### 6.2 Non-standard residues in protein, DNA, RNA chains (i)

There are no non-standard protein/DNA/RNA residues in this entry.

#### 6.3 Carbohydrates (i)

There are no monosaccharides in this entry.

#### 6.4 Ligands (i)

In the following table, the Atoms column lists the number of modelled atoms in the group and the number defined in the chemical component dictionary. The B-factors column lists the minimum, median, $95^{th}$ percentile and maximum values of B factors of atoms in the group. The column labelled 'Q< 0.9' lists the number of atoms with occupancy less than 0.9.

| Mol | Type | Chain | Res | Atoms | RSCC | RSR | B-factors(Å$^2$) | Q<0.9 |
|---|---|---|---|---|---|---|---|---|
| 4 | GOL | A | 210 | 6/6 | 0.76 | 0.18 | 23,24,25,29 | 0 |
| 4 | GOL | A | 209 | 6/6 | 0.79 | 0.15 | 27,28,30,30 | 0 |
| 3 | CL | A | 208 | 1/1 | 0.90 | 0.08 | 21,21,21,21 | 0 |
| 3 | CL | A | 204 | 1/1 | 0.95 | 0.09 | 16,16,16,16 | 0 |
| 3 | CL | A | 205 | 1/1 | 0.97 | 0.06 | 17,17,17,17 | 0 |
| 3 | CL | A | 207 | 1/1 | 0.98 | 0.05 | 18,18,18,18 | 0 |
| 3 | CL | A | 206 | 1/1 | 0.98 | 0.12 | 18,18,18,18 | 0 |
| 3 | CL | A | 202 | 1/1 | 0.99 | 0.04 | 9,9,9,9 | 0 |
| 3 | CL | A | 203 | 1/1 | 0.99 | 0.03 | 8,8,8,8 | 0 |
| 2 | NA | A | 201 | 1/1 | 0.99 | 0.04 | 6,6,6,6 | 0 |

For Manuscript Review

#### 6.5 Other polymers (i)

There are no such residues in this entry.

For Manuscript Review

### Full wwPDB X-ray Structure Validation Report (i)

Jun 8, 2022 – 03:55 PM EDT

| | | |
|---|---|---|
| PDB ID | : | 8D7B |
| Title | : | Crystal structure of hen egg white lysozyme at 225 Kelvin |
| Deposited on | : | 2022-06-07 |
| Resolution | : | 1.50 Å(reported) |

**This wwPDB validation report is for manuscript review**

This is a Full wwPDB X-ray Structure Validation Report.

This report is produced by the wwPDB biocuration pipeline after annotation of the structure.

We welcome your comments at **
A user guide is available at
https://www.wwpdb.org/validation/2017/XrayValidationReportHelp
with specific help available everywhere you see the (i) symbol.

The following versions of software and data (see references (i)) were used in the production of this report:

| | | |
|---|---|---|
| MolProbity | : | 4.02b-467 |
| Mogul | : | 1.8.5 (274361), CSD as541be (2020) |
| Xtriage (Phenix) | : | 1.13 |
| EDS | : | 2.28 |
| Percentile statistics | : | 20191225.v01 (using entries in the PDB archive December 25th 2019) |
| Refmac | : | 5.8.0158 |
| CCP4 | : | 7.0.044 (Gargrove) |
| Ideal geometry (proteins) | : | Engh & Huber (2001) |
| Ideal geometry (DNA, RNA) | : | Parkinson et al. (1996) |
| Validation Pipeline (wwPDB-VP) | : | 2.28 |

For Manuscript Review

### 1 Overall quality at a glance (i)

The following experimental techniques were used to determine the structure:
*X-RAY DIFFRACTION*

The reported resolution of this entry is 1.50 Å.

Percentile scores (ranging between 0-100) for global validation metrics of the entry are shown in the following graphic. The table shows the number of entries on which the scores are based.

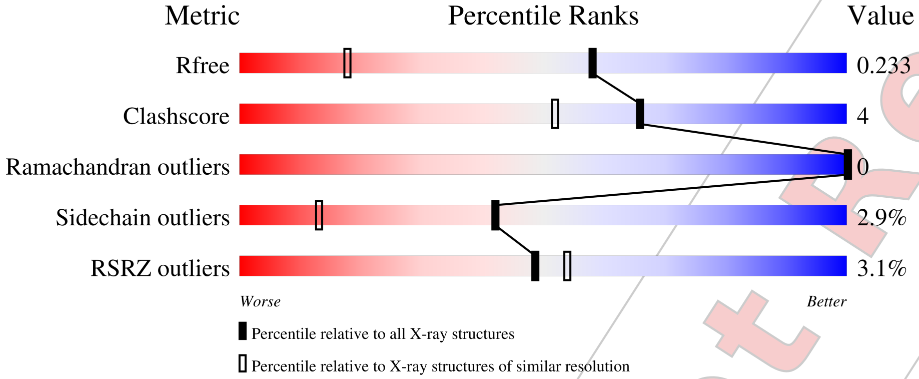


| Metric | Whole archive (#Entries) | Similar resolution (#Entries, resolution range(Å)) |
|---|---|---|
| $R_{free}$ | 130704 | 2936 (1.50-1.50) |
| Clashscore | 141614 | 3144 (1.50-1.50) |
| Ramachandran outliers | 138981 | 3066 (1.50-1.50) |
| Sidechain outliers | 138945 | 3064 (1.50-1.50) |
| RSRZ outliers | 127900 | 2884 (1.50-1.50) |

The table below summarises the geometric issues observed across the polymeric chains and their fit to the electron density. The red, orange, yellow and green segments of the lower bar indicate the fraction of residues that contain outliers for >=3, 2, 1 and 0 types of geometric quality criteria respectively. A grey segment represents the fraction of residues that are not modelled. The numeric value for each fraction is indicated below the corresponding segment, with a dot representing fractions <=5% The upper red bar (where present) indicates the fraction of residues that have poor fit to the electron density. The numeric value is given above the bar.

| Mol | Chain | Length | Quality of chain |
|---|---|---|---|
| 1 | A | 129 | 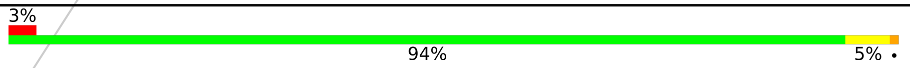 |

For Manuscript Review

#### 2 Entry composition (i)

There are 5 unique types of molecules in this entry. The entry contains 1090 atoms, of which 0 are hydrogens and 0 are deuteriums.

In the tables below, the ZeroOcc column contains the number of atoms modelled with zero occupancy, the AltConf column contains the number of residues with at least one atom in alternate conformation and the Trace column contains the number of residues modelled with at most 2 atoms.

- Molecule 1 is a protein called Lysozyme C.

| Mol | Chain | Residues | Atoms | | | | | ZeroOcc | AltConf | Trace |
|---|---|---|---|---|---|---|---|---|---|---|
| | | | Total | C | N | O | S | | | |
| 1 | A | 129 | 1001 | 613 | 193 | 185 | 10 | 0 | 0 | 0 |

- Molecule 2 is SODIUM ION (three-letter code: NA) (formula: Na).

| Mol | Chain | Residues | Atoms | | ZeroOcc | AltConf |
|---|---|---|---|---|---|---|
| | | | Total | Na | | |
| 2 | A | 1 | 1 | 1 | 0 | 0 |

- Molecule 3 is CHLORIDE ION (three-letter code: CL) (formula: Cl).

| Mol | Chain | Residues | Atoms | | ZeroOcc | AltConf |
|---|---|---|---|---|---|---|
| | | | Total | Cl | | |
| 3 | A | 7 | 7 | 7 | 0 | 0 |

- Molecule 4 is GLYCEROL (three-letter code: GOL) (formula: $C_3H_8O_3$).

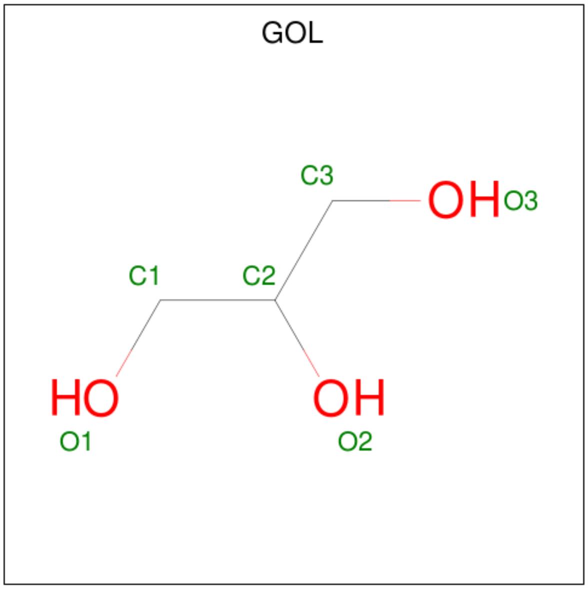


For Manuscript Review

| Mol | Chain | Residues | Atoms | ZeroOcc | AltConf |
|---|---|---|---|---|---|
| 4 | A | 1 | Total C O<br>6 3 3 | 0 | 0 |
| 4 | A | 1 | Total C O<br>6 3 3 | 0 | 0 |

- Molecule 5 is water.

| Mol | Chain | Residues | Atoms | ZeroOcc | AltConf |
|---|---|---|---|---|---|
| 5 | A | 69 | Total O<br>69 69 | 0 | 0 |

For Manuscript Review

### 3 Residue-property plots (i)

These plots are drawn for all protein, RNA, DNA and oligosaccharide chains in the entry. The first graphic for a chain summarises the proportions of the various outlier classes displayed in the second graphic. The second graphic shows the sequence view annotated by issues in geometry and electron density. Residues are color-coded according to the number of geometric quality criteria for which they contain at least one outlier: green = 0, yellow = 1, orange = 2 and red = 3 or more. A red dot above a residue indicates a poor fit to the electron density (RSRZ > 2). Stretches of 2 or more consecutive residues without any outlier are shown as a green connector. Residues present in the sample, but not in the model, are shown in grey.

• Molecule 1: Lysozyme C

Chain A:

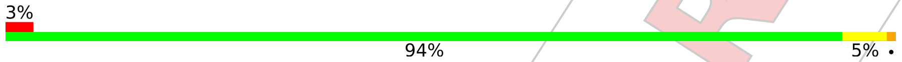


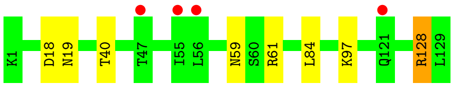


For Manuscript Review

### 4 Data and refinement statistics (i)

| Property | Value | Source |
|---|---|---|
| Space group | P 43 21 2 | Depositor |
| Cell constants<br>a, b, c, $\alpha$, $\beta$, $\gamma$ | 79.18Å 79.18Å 37.14Å<br>90.00° 90.00° 90.00° | Depositor |
| Resolution (Å) | 22.36 – 1.50<br>22.35 – 1.50 | Depositor<br>EDS |
| % Data completeness<br>(in resolution range) | 99.6 (22.36-1.50)<br>99.7 (22.35-1.50) | Depositor<br>EDS |
| $R_{merge}$ | (Not available) | Depositor |
| $R_{sym}$ | (Not available) | Depositor |
| $< I/\sigma(I) >$ [1] | 1.75 (at 1.49Å) | Xtriage |
| Refinement program | REFMAC 5.8.0267 | Depositor |
| R, $R_{free}$ | 0.191 , 0.226<br>0.198 , 0.233 | Depositor<br>DCC |
| $R_{free}$ test set | 998 reflections (5.11%) | wwPDB-VP |
| Wilson B-factor (Å$^2$) | 13.5 | Xtriage |
| Anisotropy | 0.014 | Xtriage |
| Bulk solvent $k_{sol}$(e/Å$^3$), $B_{sol}$(Å$^2$) | 0.40 , 39.2 | EDS |
| L-test for twinning[2] | $< \|L\| > = 0.42$, $< L^2 > = 0.25$ | Xtriage |
| Estimated twinning fraction | No twinning to report. | Xtriage |
| $F_o$,$F_c$ correlation | 0.95 | EDS |
| Total number of atoms | 1090 | wwPDB-VP |
| Average B, all atoms (Å$^2$) | 16.0 | wwPDB-VP |

Xtriage's analysis on translational NCS is as follows: *The largest off-origin peak in the Patterson function is 8.44% of the height of the origin peak. No significant pseudotranslation is detected.*

[1] Intensities estimated from amplitudes.
[2] Theoretical values of $< |L| >$, $< L^2 >$ for acentric reflections are 0.5, 0.333 respectively for untwinned datasets, and 0.375, 0.2 for perfectly twinned datasets.

For Manuscript Review

### 5 Model quality (i)

#### 5.1 Standard geometry (i)

Bond lengths and bond angles in the following residue types are not validated in this section: NA, GOL, CL

The Z score for a bond length (or angle) is the number of standard deviations the observed value is removed from the expected value. A bond length (or angle) with $|Z| > 5$ is considered an outlier worth inspection. RMSZ is the root-mean-square of all Z scores of the bond lengths (or angles).

| Mol | Chain | Bond lengths | | Bond angles | |
|---|---|---|---|---|---|
| | | RMSZ | #$|Z|$ >5 | RMSZ | #$|Z|$ >5 |
| 1 | A | 0.75 | 0/1021 | 0.91 | 0/1379 |

There are no bond length outliers.

There are no bond angle outliers.

There are no chirality outliers.

There are no planarity outliers.

#### 5.2 Too-close contacts (i)

In the following table, the Non-H and H(model) columns list the number of non-hydrogen atoms and hydrogen atoms in the chain respectively. The H(added) column lists the number of hydrogen atoms added and optimized by MolProbity. The Clashes column lists the number of clashes within the asymmetric unit, whereas Symm-Clashes lists symmetry-related clashes.

| Mol | Chain | Non-H | H(model) | H(added) | Clashes | Symm-Clashes |
|---|---|---|---|---|---|---|
| 1 | A | 1001 | 0 | 958 | 8 | 0 |
| 2 | A | 1 | 0 | 0 | 0 | 0 |
| 3 | A | 7 | 0 | 0 | 0 | 0 |
| 4 | A | 12 | 0 | 16 | 0 | 0 |
| 5 | A | 69 | 0 | 0 | 1 | 0 |
| All | All | 1090 | 0 | 974 | 8 | 0 |

The all-atom clashscore is defined as the number of clashes found per 1000 atoms (including hydrogen atoms). The all-atom clashscore for this structure is 4.

All (8) close contacts within the same asymmetric unit are listed below, sorted by their clash magnitude.

For Manuscript Review

| Atom-1 | Atom-2 | Interatomic distance (Å) | Clash overlap (Å) |
|---|---|---|---|
| 1:A:18:ASP:HB2 | 5:A:340:HOH:O | 1.94 | 0.66 |
| 1:A:128:ARG:HH11 | 1:A:128:ARG:CG | 2.15 | 0.59 |
| 1:A:59:ASN:ND2 | 1:A:61:ARG:HB3 | 2.22 | 0.54 |
| 1:A:128:ARG:HH11 | 1:A:128:ARG:HG2 | 1.74 | 0.52 |
| 1:A:59:ASN:ND2 | 1:A:61:ARG:H | 2.07 | 0.52 |
| 1:A:128:ARG:CG | 1:A:128:ARG:NH1 | 2.77 | 0.46 |
| 1:A:59:ASN:HD22 | 1:A:61:ARG:HB3 | 1.85 | 0.41 |
| 1:A:40:THR:O | 1:A:84:LEU:HA | 2.20 | 0.41 |

There are no symmetry-related clashes.

#### 5.3 Torsion angles (i)

##### 5.3.1 Protein backbone (i)

In the following table, the Percentiles column shows the percent Ramachandran outliers of the chain as a percentile score with respect to all X-ray entries followed by that with respect to entries of similar resolution.

The Analysed column shows the number of residues for which the backbone conformation was analysed, and the total number of residues.

| Mol | Chain | Analysed | Favoured | Allowed | Outliers | Percentiles |
|---|---|---|---|---|---|---|
| 1 | A | 127/129 (98%) | 126 (99%) | 1 (1%) | 0 | 100 100 |

There are no Ramachandran outliers to report.

##### 5.3.2 Protein sidechains (i)

In the following table, the Percentiles column shows the percent sidechain outliers of the chain as a percentile score with respect to all X-ray entries followed by that with respect to entries of similar resolution.

The Analysed column shows the number of residues for which the sidechain conformation was analysed, and the total number of residues.

| Mol | Chain | Analysed | Rotameric | Outliers | Percentiles |
|---|---|---|---|---|---|
| 1 | A | 105/105 (100%) | 102 (97%) | 3 (3%) | 42 13 |

All (3) residues with a non-rotameric sidechain are listed below:

For Manuscript Review

| Mol | Chain | Res | Type |
|---|---|---|---|
| 1 | A | 19 | ASN |
| 1 | A | 97 | LYS |
| 1 | A | 128 | ARG |

Sometimes sidechains can be flipped to improve hydrogen bonding and reduce clashes. All (3) such sidechains are listed below:

| Mol | Chain | Res | Type |
|---|---|---|---|
| 1 | A | 46 | ASN |
| 1 | A | 59 | ASN |
| 1 | A | 65 | ASN |

##### 5.3.3 RNA (i)

There are no RNA molecules in this entry.

#### 5.4 Non-standard residues in protein, DNA, RNA chains (i)

There are no non-standard protein/DNA/RNA residues in this entry.

#### 5.5 Carbohydrates (i)

There are no monosaccharides in this entry.

#### 5.6 Ligand geometry (i)

Of 10 ligands modelled in this entry, 8 are monoatomic - leaving 2 for Mogul analysis.

In the following table, the Counts columns list the number of bonds (or angles) for which Mogul statistics could be retrieved, the number of bonds (or angles) that are observed in the model and the number of bonds (or angles) that are defined in the Chemical Component Dictionary. The Link column lists molecule types, if any, to which the group is linked. The Z score for a bond length (or angle) is the number of standard deviations the observed value is removed from the expected value. A bond length (or angle) with $|Z| > 2$ is considered an outlier worth inspection. RMSZ is the root-mean-square of all Z scores of the bond lengths (or angles).

| Mol | Type | Chain | Res | Link | Bond lengths | | | Bond angles | | |
|---|---|---|---|---|---|---|---|---|---|---|
| | | | | | Counts | RMSZ | $\#\|Z\| > 2$ | Counts | RMSZ | $\#\|Z\| > 2$ |
| 4 | GOL | A | 210 | - | 5,5,5 | 0.11 | 0 | 5,5,5 | 0.22 | 0 |
| 4 | GOL | A | 209 | - | 5,5,5 | 0.11 | 0 | 5,5,5 | 0.31 | 0 |

For Manuscript Review

In the following table, the Chirals column lists the number of chiral outliers, the number of chiral centers analysed, the number of these observed in the model and the number defined in the Chemical Component Dictionary. Similar counts are reported in the Torsion and Rings columns. '-' means no outliers of that kind were identified.

| Mol | Type | Chain | Res | Link | Chirals | Torsions | Rings |
|---|---|---|---|---|---|---|---|
| 4 | GOL | A | 210 | - | - | 0/4/4/4 | - |
| 4 | GOL | A | 209 | - | - | 2/4/4/4 | - |

There are no bond length outliers.

There are no bond angle outliers.

There are no chirality outliers.

All (2) torsion outliers are listed below:

| Mol | Chain | Res | Type | Atoms |
|---|---|---|---|---|
| 4 | A | 209 | GOL | O1-C1-C2-C3 |
| 4 | A | 209 | GOL | O1-C1-C2-O2 |

There are no ring outliers.

No monomer is involved in short contacts.

#### 5.7 Other polymers ⓘ

There are no such residues in this entry.

#### 5.8 Polymer linkage issues ⓘ

There are no chain breaks in this entry.

For Manuscript Review

### 6 Fit of model and data (i)

#### 6.1 Protein, DNA and RNA chains (i)

In the following table, the column labelled '#RSRZ> 2' contains the number (and percentage) of RSRZ outliers, followed by percent RSRZ outliers for the chain as percentile scores relative to all X-ray entries and entries of similar resolution. The OWAB column contains the minimum, median, $95^{th}$ percentile and maximum values of the occupancy-weighted average B-factor per residue. The column labelled 'Q< 0.9' lists the number of (and percentage) of residues with an average occupancy less than 0.9.

| Mol | Chain | Analysed | <RSRZ> | #RSRZ>2 | | | OWAB(Å$^2$) | Q<0.9 |
|---|---|---|---|---|---|---|---|---|
| 1 | A | 129/129 (100%) | 0.12 | 4 (3%) | 49 | 54 | 9, 15, 26, 37 | 0 |

All (4) RSRZ outliers are listed below:

| Mol | Chain | Res | Type | RSRZ |
|---|---|---|---|---|
| 1 | A | 121 | GLN | 2.1 |
| 1 | A | 47 | THR | 2.1 |
| 1 | A | 56 | LEU | 2.0 |
| 1 | A | 55 | ILE | 2.0 |

#### 6.2 Non-standard residues in protein, DNA, RNA chains (i)

There are no non-standard protein/DNA/RNA residues in this entry.

#### 6.3 Carbohydrates (i)

There are no monosaccharides in this entry.

#### 6.4 Ligands (i)

In the following table, the Atoms column lists the number of modelled atoms in the group and the number defined in the chemical component dictionary. The B-factors column lists the minimum, median, $95^{th}$ percentile and maximum values of B factors of atoms in the group. The column labelled 'Q< 0.9' lists the number of atoms with occupancy less than 0.9.

| Mol | Type | Chain | Res | Atoms | RSCC | RSR | B-factors(Å$^2$) | Q<0.9 |
|---|---|---|---|---|---|---|---|---|
| 4 | GOL | A | 209 | 6/6 | 0.62 | 0.28 | 46,48,50,51 | 0 |
| 4 | GOL | A | 210 | 6/6 | 0.68 | 0.22 | 40,43,46,47 | 0 |
| 3 | CL | A | 208 | 1/1 | 0.86 | 0.17 | 36,36,36,36 | 0 |

*Continued on next page...*

For Manuscript Review

*Continued from previous page...*

| Mol | Type | Chain | Res | Atoms | RSCC | RSR | B-factors($Å^2$) | Q<0.9 |
|---|---|---|---|---|---|---|---|---|
| 3 | CL | A | 205 | 1/1 | 0.94 | 0.07 | 35,35,35,35 | 0 |
| 3 | CL | A | 207 | 1/1 | 0.95 | 0.08 | 30,30,30,30 | 0 |
| 3 | CL | A | 206 | 1/1 | 0.96 | 0.15 | 34,34,34,34 | 0 |
| 2 | NA | A | 201 | 1/1 | 0.98 | 0.04 | 12,12,12,12 | 0 |
| 3 | CL | A | 204 | 1/1 | 0.98 | 0.06 | 25,25,25,25 | 0 |
| 3 | CL | A | 203 | 1/1 | 0.99 | 0.04 | 18,18,18,18 | 0 |
| 3 | CL | A | 202 | 1/1 | 0.99 | 0.04 | 16,16,16,16 | 0 |

#### 6.5 Other polymers ⓘ

There are no such residues in this entry.

For Manuscript Review

### Full wwPDB X-ray Structure Validation Report (i)

Jun 8, 2022 – 04:01 PM EDT

| | | |
|---|---|---|
| PDB ID | : | 8D7C |
| Title | : | Crystal structure of hen egg white lysozyme at 225 Kelvin (Duplicate) |
| Deposited on | : | 2022-06-07 |
| Resolution | : | 1.50 Å(reported) |

**This wwPDB validation report is for manuscript review**

This is a Full wwPDB X-ray Structure Validation Report.

This report is produced by the wwPDB biocuration pipeline after annotation of the structure.

We welcome your comments at **
A user guide is available at
https://www.wwpdb.org/validation/2017/XrayValidationReportHelp
with specific help available everywhere you see the (i) symbol.

The following versions of software and data (see references (i)) were used in the production of this report:

| | | |
|---|---|---|
| MolProbity | : | 4.02b-467 |
| Mogul | : | 1.8.5 (274361), CSD as541be (2020) |
| Xtriage (Phenix) | : | 1.13 |
| EDS | : | 2.28 |
| Percentile statistics | : | 20191225.v01 (using entries in the PDB archive December 25th 2019) |
| Refmac | : | 5.8.0158 |
| CCP4 | : | 7.0.044 (Gargrove) |
| Ideal geometry (proteins) | : | Engh & Huber (2001) |
| Ideal geometry (DNA, RNA) | : | Parkinson et al. (1996) |
| Validation Pipeline (wwPDB-VP) | : | 2.28 |

For Manuscript Review

### 1 Overall quality at a glance (i)

The following experimental techniques were used to determine the structure:
*X-RAY DIFFRACTION*

The reported resolution of this entry is 1.50 Å.

Percentile scores (ranging between 0-100) for global validation metrics of the entry are shown in the following graphic. The table shows the number of entries on which the scores are based.

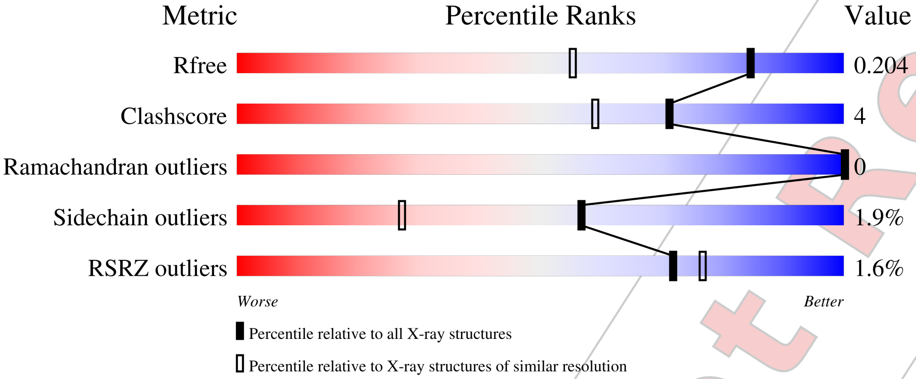


| Metric | Whole archive (#Entries) | Similar resolution (#Entries, resolution range(Å)) |
|---|---|---|
| $R_{free}$ | 130704 | 2936 (1.50-1.50) |
| Clashscore | 141614 | 3144 (1.50-1.50) |
| Ramachandran outliers | 138981 | 3066 (1.50-1.50) |
| Sidechain outliers | 138945 | 3064 (1.50-1.50) |
| RSRZ outliers | 127900 | 2884 (1.50-1.50) |

The table below summarises the geometric issues observed across the polymeric chains and their fit to the electron density. The red, orange, yellow and green segments of the lower bar indicate the fraction of residues that contain outliers for >=3, 2, 1 and 0 types of geometric quality criteria respectively. A grey segment represents the fraction of residues that are not modelled. The numeric value for each fraction is indicated below the corresponding segment, with a dot representing fractions <=5% The upper red bar (where present) indicates the fraction of residues that have poor fit to the electron density. The numeric value is given above the bar.

| Mol | Chain | Length | Quality of chain |
|---|---|---|---|
| 1 | A | 129 | 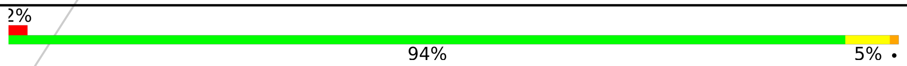  |

For Manuscript Review

### 2 Entry composition (i)

There are 5 unique types of molecules in this entry. The entry contains 1096 atoms, of which 0 are hydrogens and 0 are deuteriums.

In the tables below, the ZeroOcc column contains the number of atoms modelled with zero occupancy, the AltConf column contains the number of residues with at least one atom in alternate conformation and the Trace column contains the number of residues modelled with at most 2 atoms.

- Molecule 1 is a protein called Lysozyme C.

| Mol | Chain | Residues | Atoms | | | | | ZeroOcc | AltConf | Trace |
|---|---|---|---|---|---|---|---|---|---|---|
| | | | Total | C | N | O | S | | | |
| 1 | A | 129 | 1001 | 613 | 193 | 185 | 10 | 0 | 0 | 0 |

- Molecule 2 is SODIUM ION (three-letter code: NA) (formula: Na).

| Mol | Chain | Residues | Atoms | | ZeroOcc | AltConf |
|---|---|---|---|---|---|---|
| | | | Total | Na | | |
| 2 | A | 1 | 1 | 1 | 0 | 0 |

- Molecule 3 is CHLORIDE ION (three-letter code: CL) (formula: Cl).

| Mol | Chain | Residues | Atoms | | ZeroOcc | AltConf |
|---|---|---|---|---|---|---|
| | | | Total | Cl | | |
| 3 | A | 7 | 7 | 7 | 0 | 0 |

- Molecule 4 is GLYCEROL (three-letter code: GOL) (formula: $C_3H_8O_3$).

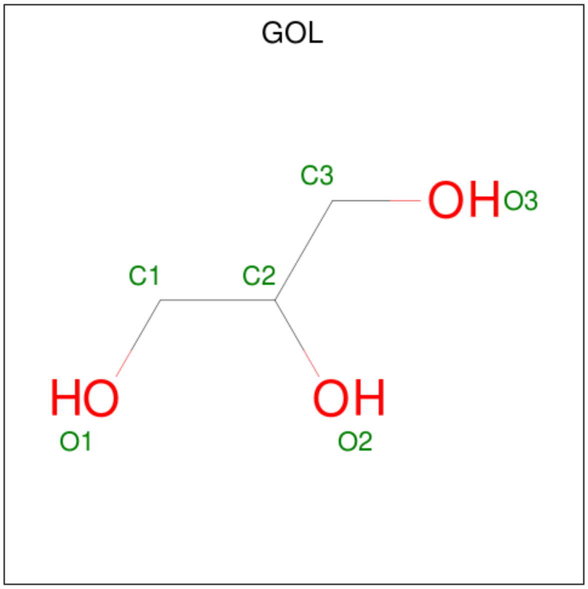


For Manuscript Review

| Mol | Chain | Residues | Atoms | ZeroOcc | AltConf |
|---|---|---|---|---|---|
| 4 | A | 1 | Total C O<br>6 3 3 | 0 | 0 |
| 4 | A | 1 | Total C O<br>6 3 3 | 0 | 0 |

- Molecule 5 is water.

| Mol | Chain | Residues | Atoms | ZeroOcc | AltConf |
|---|---|---|---|---|---|
| 5 | A | 75 | Total O<br>75 75 | 0 | 0 |

For Manuscript Review

### 3 Residue-property plots (i)

These plots are drawn for all protein, RNA, DNA and oligosaccharide chains in the entry. The first graphic for a chain summarises the proportions of the various outlier classes displayed in the second graphic. The second graphic shows the sequence view annotated by issues in geometry and electron density. Residues are color-coded according to the number of geometric quality criteria for which they contain at least one outlier: green = 0, yellow = 1, orange = 2 and red = 3 or more. A red dot above a residue indicates a poor fit to the electron density (RSRZ > 2). Stretches of 2 or more consecutive residues without any outlier are shown as a green connector. Residues present in the sample, but not in the model, are shown in grey.

- Molecule 1: Lysozyme C

Chain A:

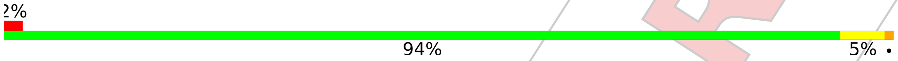


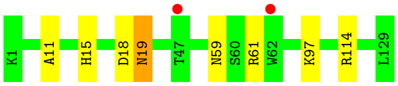


For Manuscript Review

### 4 Data and refinement statistics (i)

| Property | Value | Source |
|---|---|---|
| Space group | P 43 21 2 | Depositor |
| Cell constants<br>a, b, c, $\alpha$, $\beta$, $\gamma$ | 79.43Å 79.43Å 37.39Å<br>90.00° 90.00° 90.00° | Depositor |
| Resolution (Å) | 25.13 – 1.50<br>25.12 – 1.50 | Depositor<br>EDS |
| % Data completeness<br>(in resolution range) | 99.7 (25.13-1.50)<br>99.8 (25.12-1.50) | Depositor<br>EDS |
| $R_{merge}$ | (Not available) | Depositor |
| $R_{sym}$ | (Not available) | Depositor |
| $< I/\sigma(I) >$ [1] | 1.61 (at 1.49Å) | Xtriage |
| Refinement program | REFMAC 5.8.0267 | Depositor |
| R, $R_{free}$ | 0.168 , 0.195<br>0.176 , 0.204 | Depositor<br>DCC |
| $R_{free}$ test set | 1017 reflections (5.14%) | wwPDB-VP |
| Wilson B-factor (Å$^2$) | 12.1 | Xtriage |
| Anisotropy | 0.049 | Xtriage |
| Bulk solvent $k_{sol}$(e/Å$^3$), $B_{sol}$(Å$^2$) | 0.38 , 41.2 | EDS |
| L-test for twinning[2] | $< \|L\| > = 0.48$, $< L^2 > = 0.32$ | Xtriage |
| Estimated twinning fraction | No twinning to report. | Xtriage |
| $F_o$,$F_c$ correlation | 0.96 | EDS |
| Total number of atoms | 1096 | wwPDB-VP |
| Average B, all atoms (Å$^2$) | 15.0 | wwPDB-VP |

Xtriage's analysis on translational NCS is as follows: *The largest off-origin peak in the Patterson function is 9.16% of the height of the origin peak. No significant pseudotranslation is detected.*

[1]Intensities estimated from amplitudes.

[2]Theoretical values of $< |L| >$, $< L^2 >$ for acentric reflections are 0.5, 0.333 respectively for untwinned datasets, and 0.375, 0.2 for perfectly twinned datasets.

For Manuscript Review

### 5 Model quality (i)

#### 5.1 Standard geometry (i)

Bond lengths and bond angles in the following residue types are not validated in this section: NA, GOL, CL

The Z score for a bond length (or angle) is the number of standard deviations the observed value is removed from the expected value. A bond length (or angle) with $|Z| > 5$ is considered an outlier worth inspection. RMSZ is the root-mean-square of all Z scores of the bond lengths (or angles).

| Mol | Chain | Bond lengths | | Bond angles | |
|---|---|---|---|---|---|
| | | RMSZ | #$\|Z\|$ >5 | RMSZ | #$\|Z\|$ >5 |
| 1 | A | 0.78 | 0/1021 | 0.90 | 2/1379 (0.1%) |

There are no bond length outliers.

All (2) bond angle outliers are listed below:

| Mol | Chain | Res | Type | Atoms | Z | Observed($^o$) | Ideal($^o$) |
|---|---|---|---|---|---|---|---|
| 1 | A | 114 | ARG | NE-CZ-NH2 | -5.82 | 117.39 | 120.30 |
| 1 | A | 18 | ASP | CB-CG-OD1 | 5.31 | 123.08 | 118.30 |

There are no chirality outliers.

There are no planarity outliers.

#### 5.2 Too-close contacts (i)

In the following table, the Non-H and H(model) columns list the number of non-hydrogen atoms and hydrogen atoms in the chain respectively. The H(added) column lists the number of hydrogen atoms added and optimized by MolProbity. The Clashes column lists the number of clashes within the asymmetric unit, whereas Symm-Clashes lists symmetry-related clashes.

| Mol | Chain | Non-H | H(model) | H(added) | Clashes | Symm-Clashes |
|---|---|---|---|---|---|---|
| 1 | A | 1001 | 0 | 958 | 7 | 0 |
| 2 | A | 1 | 0 | 0 | 0 | 0 |
| 3 | A | 7 | 0 | 0 | 0 | 0 |
| 4 | A | 12 | 0 | 16 | 0 | 0 |
| 5 | A | 75 | 0 | 0 | 0 | 0 |
| All | All | 1096 | 0 | 974 | 7 | 0 |

The all-atom clashscore is defined as the number of clashes found per 1000 atoms (including hydrogen atoms). The all-atom clashscore for this structure is 4.

For Manuscript Review

All (7) close contacts within the same asymmetric unit are listed below, sorted by their clash magnitude.

| Atom-1 | Atom-2 | Interatomic distance (Å) | Clash overlap (Å) |
|---|---|---|---|
| 1:A:59:ASN:ND2 | 1:A:61:ARG:H | 1.89 | 0.70 |
| 1:A:59:ASN:ND2 | 1:A:61:ARG:HB3 | 2.17 | 0.60 |
| 1:A:59:ASN:ND2 | 1:A:61:ARG:CB | 2.72 | 0.52 |
| 1:A:11:ALA:O | 1:A:15:HIS:HD2 | 1.95 | 0.48 |
| 1:A:59:ASN:HD22 | 1:A:61:ARG:HB3 | 1.79 | 0.46 |
| 1:A:19:ASN:HD22 | 1:A:19:ASN:HA | 1.66 | 0.42 |
| 1:A:59:ASN:HD21 | 1:A:61:ARG:CB | 2.33 | 0.42 |

There are no symmetry-related clashes.

#### 5.3 Torsion angles (i)

##### 5.3.1 Protein backbone (i)

In the following table, the Percentiles column shows the percent Ramachandran outliers of the chain as a percentile score with respect to all X-ray entries followed by that with respect to entries of similar resolution.

The Analysed column shows the number of residues for which the backbone conformation was analysed, and the total number of residues.

| Mol | Chain | Analysed | Favoured | Allowed | Outliers | Percentiles | |
|---|---|---|---|---|---|---|---|
| 1 | A | 127/129 (98%) | 126 (99%) | 1 (1%) | 0 | 100 | 100 |

There are no Ramachandran outliers to report.

##### 5.3.2 Protein sidechains (i)

In the following table, the Percentiles column shows the percent sidechain outliers of the chain as a percentile score with respect to all X-ray entries followed by that with respect to entries of similar resolution.

The Analysed column shows the number of residues for which the sidechain conformation was analysed, and the total number of residues.

| Mol | Chain | Analysed | Rotameric | Outliers | Percentiles | |
|---|---|---|---|---|---|---|
| 1 | A | 105/105 (100%) | 103 (98%) | 2 (2%) | 57 | 27 |

All (2) residues with a non-rotameric sidechain are listed below:

For Manuscript Review

| Mol | Chain | Res | Type |
|---|---|---|---|
| 1 | A | 19 | ASN |
| 1 | A | 97 | LYS |

Sometimes sidechains can be flipped to improve hydrogen bonding and reduce clashes. All (5) such sidechains are listed below:

| Mol | Chain | Res | Type |
|---|---|---|---|
| 1 | A | 15 | HIS |
| 1 | A | 19 | ASN |
| 1 | A | 46 | ASN |
| 1 | A | 59 | ASN |
| 1 | A | 65 | ASN |

##### 5.3.3 RNA (i)

There are no RNA molecules in this entry.

#### 5.4 Non-standard residues in protein, DNA, RNA chains (i)

There are no non-standard protein/DNA/RNA residues in this entry.

#### 5.5 Carbohydrates (i)

There are no monosaccharides in this entry.

#### 5.6 Ligand geometry (i)

Of 10 ligands modelled in this entry, 8 are monoatomic - leaving 2 for Mogul analysis.

In the following table, the Counts columns list the number of bonds (or angles) for which Mogul statistics could be retrieved, the number of bonds (or angles) that are observed in the model and the number of bonds (or angles) that are defined in the Chemical Component Dictionary. The Link column lists molecule types, if any, to which the group is linked. The Z score for a bond length (or angle) is the number of standard deviations the observed value is removed from the expected value. A bond length (or angle) with $|Z| > 2$ is considered an outlier worth inspection. RMSZ is the root-mean-square of all Z scores of the bond lengths (or angles).

| Mol | Type | Chain | Res | Link | Bond lengths | | | Bond angles | | |
|---|---|---|---|---|---|---|---|---|---|---|
| | | | | | Counts | RMSZ | $\#\|Z\| > 2$ | Counts | RMSZ | $\#\|Z\| > 2$ |
| 4 | GOL | A | 210 | - | 5,5,5 | 0.09 | 0 | 5,5,5 | 0.20 | 0 |
| 4 | GOL | A | 209 | - | 5,5,5 | 0.20 | 0 | 5,5,5 | 0.50 | 0 |

For Manuscript Review

In the following table, the Chirals column lists the number of chiral outliers, the number of chiral centers analysed, the number of these observed in the model and the number defined in the Chemical Component Dictionary. Similar counts are reported in the Torsion and Rings columns. '-' means no outliers of that kind were identified.

| Mol | Type | Chain | Res | Link | Chirals | Torsions | Rings |
|---|---|---|---|---|---|---|---|
| 4 | GOL | A | 210 | - | - | 4/4/4/4 | - |
| 4 | GOL | A | 209 | - | - | 2/4/4/4 | - |

There are no bond length outliers.

There are no bond angle outliers.

There are no chirality outliers.

All (6) torsion outliers are listed below:

| Mol | Chain | Res | Type | Atoms |
|---|---|---|---|---|
| 4 | A | 209 | GOL | C1-C2-C3-O3 |
| 4 | A | 210 | GOL | C1-C2-C3-O3 |
| 4 | A | 210 | GOL | O2-C2-C3-O3 |
| 4 | A | 209 | GOL | O2-C2-C3-O3 |
| 4 | A | 210 | GOL | O1-C1-C2-O2 |
| 4 | A | 210 | GOL | O1-C1-C2-C3 |

There are no ring outliers.

No monomer is involved in short contacts.

#### 5.7 Other polymers (i)

There are no such residues in this entry.

#### 5.8 Polymer linkage issues (i)

There are no chain breaks in this entry.

For Manuscript Review

### 6 Fit of model and data (i)

#### 6.1 Protein, DNA and RNA chains (i)

In the following table, the column labelled '#RSRZ> 2' contains the number (and percentage) of RSRZ outliers, followed by percent RSRZ outliers for the chain as percentile scores relative to all X-ray entries and entries of similar resolution. The OWAB column contains the minimum, median, $95^{th}$ percentile and maximum values of the occupancy-weighted average B-factor per residue. The column labelled 'Q< 0.9' lists the number of (and percentage) of residues with an average occupancy less than 0.9.

| Mol | Chain | Analysed | <RSRZ> | #RSRZ>2 | | | OWAB(Å$^2$) | Q<0.9 |
|---|---|---|---|---|---|---|---|---|
| 1 | A | 129/129 (100%) | 0.00 | 2 (1%) | 72 | 77 | 8, 12, 25, 34 | 0 |

All (2) RSRZ outliers are listed below:

| Mol | Chain | Res | Type | RSRZ |
|---|---|---|---|---|
| 1 | A | 62 | TRP | 2.2 |
| 1 | A | 47 | THR | 2.1 |

#### 6.2 Non-standard residues in protein, DNA, RNA chains (i)

There are no non-standard protein/DNA/RNA residues in this entry.

#### 6.3 Carbohydrates (i)

There are no monosaccharides in this entry.

#### 6.4 Ligands (i)

In the following table, the Atoms column lists the number of modelled atoms in the group and the number defined in the chemical component dictionary. The B-factors column lists the minimum, median, $95^{th}$ percentile and maximum values of B factors of atoms in the group. The column labelled 'Q< 0.9' lists the number of atoms with occupancy less than 0.9.

| Mol | Type | Chain | Res | Atoms | RSCC | RSR | B-factors(Å$^2$) | Q<0.9 |
|---|---|---|---|---|---|---|---|---|
| 4 | GOL | A | 210 | 6/6 | 0.52 | 0.27 | 47,50,51,52 | 0 |
| 4 | GOL | A | 209 | 6/6 | 0.85 | 0.21 | 33,38,41,46 | 0 |
| 3 | CL | A | 205 | 1/1 | 0.94 | 0.09 | 37,37,37,37 | 0 |
| 3 | CL | A | 208 | 1/1 | 0.94 | 0.23 | 40,40,40,40 | 0 |
| 3 | CL | A | 207 | 1/1 | 0.96 | 0.12 | 35,35,35,35 | 0 |

*Continued on next page...*

For Manuscript Review

*Continued from previous page...*

| Mol | Type | Chain | Res | Atoms | RSCC | RSR | B-factors(Å$^2$) | Q<0.9 |
|---|---|---|---|---|---|---|---|---|
| 3 | CL | A | 206 | 1/1 | 0.96 | 0.14 | 33,33,33,33 | 0 |
| 3 | CL | A | 204 | 1/1 | 0.98 | 0.08 | 20,20,20,20 | 0 |
| 3 | CL | A | 203 | 1/1 | 0.99 | 0.04 | 18,18,18,18 | 0 |
| 3 | CL | A | 202 | 1/1 | 0.99 | 0.03 | 16,16,16,16 | 0 |
| 2 | NA | A | 201 | 1/1 | 1.00 | 0.05 | 12,12,12,12 | 0 |

#### 6.5 Other polymers (i)

There are no such residues in this entry.

For Manuscript Review

### Full wwPDB X-ray Structure Validation Report (i)

Jun 8, 2022 – 03:43 PM EDT

| | | |
|---|---|---|
| PDB ID | : | 8D7D |
| Title | : | Crystal structure of hen egg white lysozyme at 225 Kelvin (Triplicate) |
| Deposited on | : | 2022-06-07 |
| Resolution | : | 1.50 Å(reported) |

**This wwPDB validation report is for manuscript review**

This is a Full wwPDB X-ray Structure Validation Report.

This report is produced by the wwPDB biocuration pipeline after annotation of the structure.

We welcome your comments at **
A user guide is available at
https://www.wwpdb.org/validation/2017/XrayValidationReportHelp
with specific help available everywhere you see the (i) symbol.

The following versions of software and data (see references (i)) were used in the production of this report:

| | | |
|---|---|---|
| MolProbity | : | 4.02b-467 |
| Mogul | : | 1.8.5 (274361), CSD as541be (2020) |
| Xtriage (Phenix) | : | 1.13 |
| EDS | : | 2.28 |
| Percentile statistics | : | 20191225.v01 (using entries in the PDB archive December 25th 2019) |
| Refmac | : | 5.8.0158 |
| CCP4 | : | 7.0.044 (Gargrove) |
| Ideal geometry (proteins) | : | Engh & Huber (2001) |
| Ideal geometry (DNA, RNA) | : | Parkinson et al. (1996) |
| Validation Pipeline (wwPDB-VP) | : | 2.28 |

For Manuscript Review

### 1 Overall quality at a glance (i)

The following experimental techniques were used to determine the structure:
*X-RAY DIFFRACTION*

The reported resolution of this entry is 1.50 Å.

Percentile scores (ranging between 0-100) for global validation metrics of the entry are shown in the following graphic. The table shows the number of entries on which the scores are based.

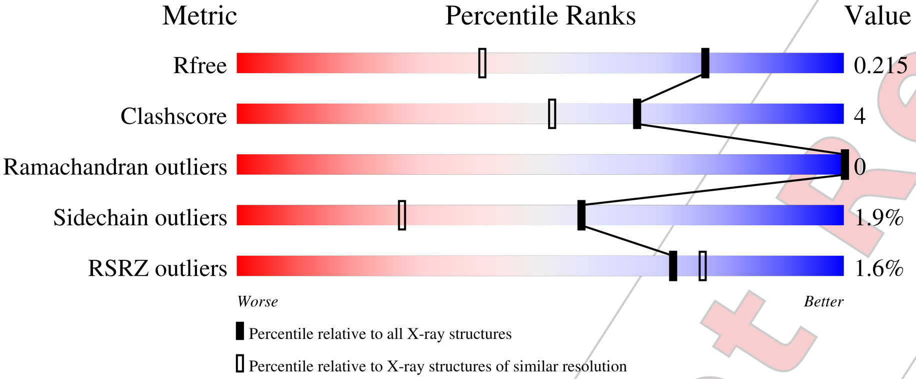


| Metric | Whole archive (#Entries) | Similar resolution (#Entries, resolution range(Å)) |
|---|---|---|
| $R_{free}$ | 130704 | 2936 (1.50-1.50) |
| Clashscore | 141614 | 3144 (1.50-1.50) |
| Ramachandran outliers | 138981 | 3066 (1.50-1.50) |
| Sidechain outliers | 138945 | 3064 (1.50-1.50) |
| RSRZ outliers | 127900 | 2884 (1.50-1.50) |

The table below summarises the geometric issues observed across the polymeric chains and their fit to the electron density. The red, orange, yellow and green segments of the lower bar indicate the fraction of residues that contain outliers for >=3, 2, 1 and 0 types of geometric quality criteria respectively. A grey segment represents the fraction of residues that are not modelled. The numeric value for each fraction is indicated below the corresponding segment, with a dot representing fractions <=5% The upper red bar (where present) indicates the fraction of residues that have poor fit to the electron density. The numeric value is given above the bar.

| Mol | Chain | Length | Quality of chain |
|---|---|---|---|
| 1 | A | 129 | 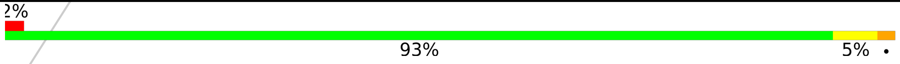  |

For Manuscript Review

#### 2 Entry composition (i)

There are 5 unique types of molecules in this entry. The entry contains 1103 atoms, of which 0 are hydrogens and 0 are deuteriums.

In the tables below, the ZeroOcc column contains the number of atoms modelled with zero occupancy, the AltConf column contains the number of residues with at least one atom in alternate conformation and the Trace column contains the number of residues modelled with at most 2 atoms.

- Molecule 1 is a protein called Lysozyme C.

| Mol | Chain | Residues | Atoms | | | | | ZeroOcc | AltConf | Trace |
|---|---|---|---|---|---|---|---|---|---|---|
| | | | Total | C | N | O | S | | | |
| 1 | A | 129 | 1001 | 613 | 193 | 185 | 10 | 0 | 0 | 0 |

- Molecule 2 is SODIUM ION (three-letter code: NA) (formula: Na).

| Mol | Chain | Residues | Atoms | | ZeroOcc | AltConf |
|---|---|---|---|---|---|---|
| | | | Total | Na | | |
| 2 | A | 1 | 1 | 1 | 0 | 0 |

- Molecule 3 is CHLORIDE ION (three-letter code: CL) (formula: Cl).

| Mol | Chain | Residues | Atoms | | ZeroOcc | AltConf |
|---|---|---|---|---|---|---|
| | | | Total | Cl | | |
| 3 | A | 7 | 7 | 7 | 0 | 0 |

- Molecule 4 is GLYCEROL (three-letter code: GOL) (formula: $C_3H_8O_3$).

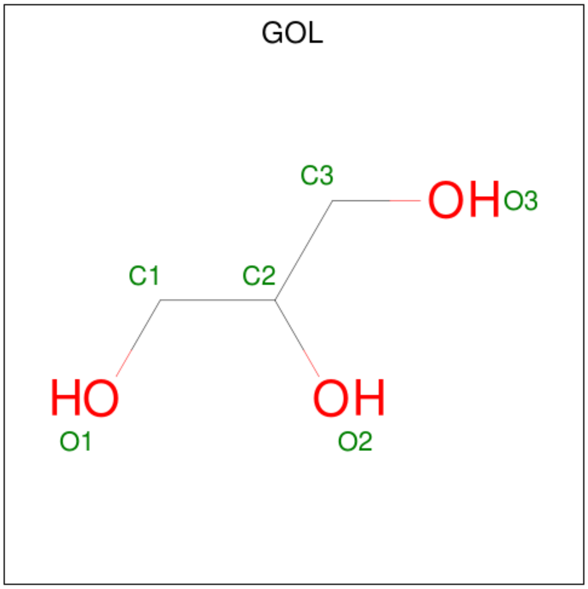


For Manuscript Review

| Mol | Chain | Residues | Atoms | ZeroOcc | AltConf |
|---|---|---|---|---|---|
| 4 | A | 1 | Total C O<br>6 3 3 | 0 | 0 |
| 4 | A | 1 | Total C O<br>6 3 3 | 0 | 0 |

- Molecule 5 is water.

| Mol | Chain | Residues | Atoms | ZeroOcc | AltConf |
|---|---|---|---|---|---|
| 5 | A | 82 | Total O<br>82 82 | 0 | 0 |

For Manuscript Review

### 3 Residue-property plots (i)

These plots are drawn for all protein, RNA, DNA and oligosaccharide chains in the entry. The first graphic for a chain summarises the proportions of the various outlier classes displayed in the second graphic. The second graphic shows the sequence view annotated by issues in geometry and electron density. Residues are color-coded according to the number of geometric quality criteria for which they contain at least one outlier: green = 0, yellow = 1, orange = 2 and red = 3 or more. A red dot above a residue indicates a poor fit to the electron density (RSRZ > 2). Stretches of 2 or more consecutive residues without any outlier are shown as a green connector. Residues present in the sample, but not in the model, are shown in grey.

- Molecule 1: Lysozyme C

Chain A:

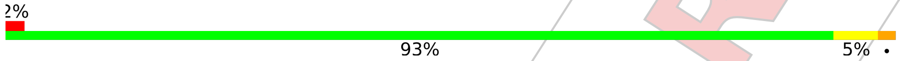


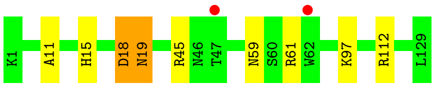


For Manuscript Review

### 4 Data and refinement statistics (i)

| Property | Value | Source |
|---|---|---|
| Space group | P 43 21 2 | Depositor |
| Cell constants<br>a, b, c, $\alpha$, $\beta$, $\gamma$ | 79.49Å 79.49Å 37.49Å<br>90.00° 90.00° 90.00° | Depositor |
| Resolution (Å) | 22.50 – 1.50<br>22.49 – 1.50 | Depositor<br>EDS |
| % Data completeness<br>(in resolution range) | 99.6 (22.50-1.50)<br>99.6 (22.49-1.50) | Depositor<br>EDS |
| $R_{merge}$ | (Not available) | Depositor |
| $R_{sym}$ | (Not available) | Depositor |
| $< I/\sigma(I) >$ [1] | 3.04 (at 1.50Å) | Xtriage |
| Refinement program | REFMAC 5.8.0267 | Depositor |
| R, $R_{free}$ | 0.171 , 0.207<br>0.180 , 0.215 | Depositor<br>DCC |
| $R_{free}$ test set | 1022 reflections (5.17%) | wwPDB-VP |
| Wilson B-factor (Å$^2$) | 12.1 | Xtriage |
| Anisotropy | 0.075 | Xtriage |
| Bulk solvent $k_{sol}$(e/Å$^3$), $B_{sol}$(Å$^2$) | 0.38 , 40.0 | EDS |
| L-test for twinning[2] | $< \|L\| > = 0.50$, $< L^2 > = 0.34$ | Xtriage |
| Estimated twinning fraction | No twinning to report. | Xtriage |
| $F_o$,$F_c$ correlation | 0.96 | EDS |
| Total number of atoms | 1103 | wwPDB-VP |
| Average B, all atoms (Å$^2$) | 15.0 | wwPDB-VP |

Xtriage's analysis on translational NCS is as follows: *The largest off-origin peak in the Patterson function is 9.07% of the height of the origin peak. No significant pseudotranslation is detected.*

[1] Intensities estimated from amplitudes.

[2] Theoretical values of $< |L| >$, $< L^2 >$ for acentric reflections are 0.5, 0.333 respectively for untwinned datasets, and 0.375, 0.2 for perfectly twinned datasets.

For Manuscript Review

### 5 Model quality (i)

#### 5.1 Standard geometry (i)

Bond lengths and bond angles in the following residue types are not validated in this section: CL, NA, GOL

The Z score for a bond length (or angle) is the number of standard deviations the observed value is removed from the expected value. A bond length (or angle) with $|Z| > 5$ is considered an outlier worth inspection. RMSZ is the root-mean-square of all Z scores of the bond lengths (or angles).

| Mol | Chain | Bond lengths | | Bond angles | |
|---|---|---|---|---|---|
| | | RMSZ | #$|Z|$ >5 | RMSZ | #$|Z|$ >5 |
| 1 | A | 0.79 | 0/1021 | 0.95 | 3/1379 (0.2%) |

There are no bond length outliers.

All (3) bond angle outliers are listed below:

| Mol | Chain | Res | Type | Atoms | Z | Observed($^o$) | Ideal($^o$) |
|---|---|---|---|---|---|---|---|
| 1 | A | 18 | ASP | CB-CG-OD1 | 6.41 | 124.06 | 118.30 |
| 1 | A | 45 | ARG | NE-CZ-NH1 | -5.66 | 117.47 | 120.30 |
| 1 | A | 112 | ARG | NE-CZ-NH2 | -5.01 | 117.79 | 120.30 |

There are no chirality outliers.

There are no planarity outliers.

#### 5.2 Too-close contacts (i)

In the following table, the Non-H and H(model) columns list the number of non-hydrogen atoms and hydrogen atoms in the chain respectively. The H(added) column lists the number of hydrogen atoms added and optimized by MolProbity. The Clashes column lists the number of clashes within the asymmetric unit, whereas Symm-Clashes lists symmetry-related clashes.

| Mol | Chain | Non-H | H(model) | H(added) | Clashes | Symm-Clashes |
|---|---|---|---|---|---|---|
| 1 | A | 1001 | 0 | 958 | 8 | 0 |
| 2 | A | 1 | 0 | 0 | 0 | 0 |
| 3 | A | 7 | 0 | 0 | 0 | 0 |
| 4 | A | 12 | 0 | 16 | 0 | 0 |
| 5 | A | 82 | 0 | 0 | 1 | 0 |
| All | All | 1103 | 0 | 974 | 8 | 0 |

The all-atom clashscore is defined as the number of clashes found per 1000 atoms (including

For Manuscript Review

hydrogen atoms). The all-atom clashscore for this structure is 4.

All (8) close contacts within the same asymmetric unit are listed below, sorted by their clash magnitude.

| Atom-1 | Atom-2 | Interatomic distance (Å) | Clash overlap (Å) |
|---|---|---|---|
| 1:A:59:ASN:ND2 | 1:A:61:ARG:H | 1.90 | 0.69 |
| 1:A:59:ASN:ND2 | 1:A:61:ARG:HB3 | 2.08 | 0.67 |
| 1:A:59:ASN:ND2 | 1:A:61:ARG:CB | 2.73 | 0.51 |
| 1:A:59:ASN:HD21 | 1:A:61:ARG:CB | 2.23 | 0.51 |
| 1:A:59:ASN:HD21 | 1:A:61:ARG:HB3 | 1.80 | 0.46 |
| 1:A:11:ALA:O | 1:A:15:HIS:HD2 | 1.98 | 0.45 |
| 1:A:19:ASN:HD22 | 1:A:19:ASN:HA | 1.71 | 0.42 |
| 1:A:18:ASP:HB3 | 5:A:321:HOH:O | 2.21 | 0.40 |

There are no symmetry-related clashes.

#### 5.3 Torsion angles (i)

##### 5.3.1 Protein backbone (i)

In the following table, the Percentiles column shows the percent Ramachandran outliers of the chain as a percentile score with respect to all X-ray entries followed by that with respect to entries of similar resolution.

The Analysed column shows the number of residues for which the backbone conformation was analysed, and the total number of residues.

| Mol | Chain | Analysed | Favoured | Allowed | Outliers | Percentiles | |
|---|---|---|---|---|---|---|---|
| 1 | A | 127/129 (98%) | 125 (98%) | 2 (2%) | 0 | 100 | 100 |

There are no Ramachandran outliers to report.

##### 5.3.2 Protein sidechains (i)

In the following table, the Percentiles column shows the percent sidechain outliers of the chain as a percentile score with respect to all X-ray entries followed by that with respect to entries of similar resolution.

The Analysed column shows the number of residues for which the sidechain conformation was analysed, and the total number of residues.

For Manuscript Review

| Mol | Chain | Analysed | Rotameric | Outliers | Percentiles | |
|---|---|---|---|---|---|---|
| 1 | A | 105/105 (100%) | 103 (98%) | 2 (2%) | 57 | 27 |

All (2) residues with a non-rotameric sidechain are listed below:

| Mol | Chain | Res | Type |
|---|---|---|---|
| 1 | A | 19 | ASN |
| 1 | A | 97 | LYS |

Sometimes sidechains can be flipped to improve hydrogen bonding and reduce clashes. All (5) such sidechains are listed below:

| Mol | Chain | Res | Type |
|---|---|---|---|
| 1 | A | 15 | HIS |
| 1 | A | 19 | ASN |
| 1 | A | 46 | ASN |
| 1 | A | 59 | ASN |
| 1 | A | 65 | ASN |

###### 5.3.3 RNA (i)

There are no RNA molecules in this entry.

##### 5.4 Non-standard residues in protein, DNA, RNA chains (i)

There are no non-standard protein/DNA/RNA residues in this entry.

##### 5.5 Carbohydrates (i)

There are no monosaccharides in this entry.

##### 5.6 Ligand geometry (i)

Of 10 ligands modelled in this entry, 8 are monoatomic - leaving 2 for Mogul analysis.

In the following table, the Counts columns list the number of bonds (or angles) for which Mogul statistics could be retrieved, the number of bonds (or angles) that are observed in the model and the number of bonds (or angles) that are defined in the Chemical Component Dictionary. The Link column lists molecule types, if any, to which the group is linked. The Z score for a bond length (or angle) is the number of standard deviations the observed value is removed from the expected value. A bond length (or angle) with $|Z| > 2$ is considered an outlier worth inspection. RMSZ is the root-mean-square of all Z scores of the bond lengths (or angles).

For Manuscript Review

| Mol | Type | Chain | Res | Link | Bond lengths | | | Bond angles | | |
|---|---|---|---|---|---|---|---|---|---|---|
| | | | | | Counts | RMSZ | #\|Z\| > 2 | Counts | RMSZ | #\|Z\| > 2 |
| 4 | GOL | A | 210 | - | 5,5,5 | 0.11 | 0 | 5,5,5 | 0.25 | 0 |
| 4 | GOL | A | 209 | - | 5,5,5 | 0.23 | 0 | 5,5,5 | 0.49 | 0 |

In the following table, the Chirals column lists the number of chiral outliers, the number of chiral centers analysed, the number of these observed in the model and the number defined in the Chemical Component Dictionary. Similar counts are reported in the Torsion and Rings columns. '-' means no outliers of that kind were identified.

| Mol | Type | Chain | Res | Link | Chirals | Torsions | Rings |
|---|---|---|---|---|---|---|---|
| 4 | GOL | A | 210 | - | - | 4/4/4/4 | - |
| 4 | GOL | A | 209 | - | - | 2/4/4/4 | - |

There are no bond length outliers.

There are no bond angle outliers.

There are no chirality outliers.

All (6) torsion outliers are listed below:

| Mol | Chain | Res | Type | Atoms |
|---|---|---|---|---|
| 4 | A | 209 | GOL | C1-C2-C3-O3 |
| 4 | A | 210 | GOL | C1-C2-C3-O3 |
| 4 | A | 209 | GOL | O2-C2-C3-O3 |
| 4 | A | 210 | GOL | O2-C2-C3-O3 |
| 4 | A | 210 | GOL | O1-C1-C2-O2 |
| 4 | A | 210 | GOL | O1-C1-C2-C3 |

There are no ring outliers.

No monomer is involved in short contacts.

#### 5.7 Other polymers (i)

There are no such residues in this entry.

#### 5.8 Polymer linkage issues (i)

There are no chain breaks in this entry.

For Manuscript Review

### 6 Fit of model and data (i)

#### 6.1 Protein, DNA and RNA chains (i)

In the following table, the column labelled '#RSRZ> 2' contains the number (and percentage) of RSRZ outliers, followed by percent RSRZ outliers for the chain as percentile scores relative to all X-ray entries and entries of similar resolution. The OWAB column contains the minimum, median, $95^{th}$ percentile and maximum values of the occupancy-weighted average B-factor per residue. The column labelled 'Q< 0.9' lists the number of (and percentage) of residues with an average occupancy less than 0.9.

| Mol | Chain | Analysed | <RSRZ> | #RSRZ>2 | | | OWAB(Å$^2$) | Q<0.9 |
|---|---|---|---|---|---|---|---|---|
| 1 | A | 129/129 (100%) | 0.04 | 2 (1%) | 72 | 77 | 8, 13, 26, 29 | 0 |

All (2) RSRZ outliers are listed below:

| Mol | Chain | Res | Type | RSRZ |
|---|---|---|---|---|
| 1 | A | 62 | TRP | 2.6 |
| 1 | A | 47 | THR | 2.5 |

#### 6.2 Non-standard residues in protein, DNA, RNA chains (i)

There are no non-standard protein/DNA/RNA residues in this entry.

#### 6.3 Carbohydrates (i)

There are no monosaccharides in this entry.

#### 6.4 Ligands (i)

In the following table, the Atoms column lists the number of modelled atoms in the group and the number defined in the chemical component dictionary. The B-factors column lists the minimum, median, $95^{th}$ percentile and maximum values of B factors of atoms in the group. The column labelled 'Q< 0.9' lists the number of atoms with occupancy less than 0.9.

| Mol | Type | Chain | Res | Atoms | RSCC | RSR | B-factors(Å$^2$) | Q<0.9 |
|---|---|---|---|---|---|---|---|---|
| 4 | GOL | A | 210 | 6/6 | 0.77 | 0.26 | 58,59,60,61 | 0 |
| 4 | GOL | A | 209 | 6/6 | 0.80 | 0.21 | 35,41,45,47 | 0 |
| 3 | CL | A | 208 | 1/1 | 0.93 | 0.20 | 39,39,39,39 | 0 |
| 3 | CL | A | 207 | 1/1 | 0.95 | 0.11 | 33,33,33,33 | 0 |
| 3 | CL | A | 205 | 1/1 | 0.95 | 0.12 | 35,35,35,35 | 0 |

*Continued on next page...*

For Manuscript Review

*Continued from previous page...*

| Mol | Type | Chain | Res | Atoms | RSCC | RSR | B-factors(Å$^2$) | Q<0.9 |
|---|---|---|---|---|---|---|---|---|
| 3 | CL | A | 206 | 1/1 | 0.97 | 0.17 | 31,31,31,31 | 0 |
| 3 | CL | A | 203 | 1/1 | 0.99 | 0.02 | 17,17,17,17 | 0 |
| 3 | CL | A | 204 | 1/1 | 0.99 | 0.08 | 20,20,20,20 | 0 |
| 2 | NA | A | 201 | 1/1 | 0.99 | 0.04 | 12,12,12,12 | 0 |
| 3 | CL | A | 202 | 1/1 | 0.99 | 0.03 | 17,17,17,17 | 0 |

#### 6.5 Other polymers ⓘ

There are no such residues in this entry.

For Manuscript Review

For Manuscript Review

### Full wwPDB X-ray Structure Validation Report (i)

Jun 8, 2022 – 03:56 PM EDT

| | | |
|---|---|---|
| PDB ID | : | 8D7J |
| Title | : | Crystal structure of hen egg white lysozyme at 250 Kelvin |
| Deposited on | : | 2022-06-07 |
| Resolution | : | 1.50 Å(reported) |

**This wwPDB validation report is for manuscript review**

This is a Full wwPDB X-ray Structure Validation Report.

This report is produced by the wwPDB biocuration pipeline after annotation of the structure.

We welcome your comments at **
A user guide is available at
https://www.wwpdb.org/validation/2017/XrayValidationReportHelp
with specific help available everywhere you see the (i) symbol.

The following versions of software and data (see references (i)) were used in the production of this report:

| | | |
|---|---|---|
| MolProbity | : | 4.02b-467 |
| Mogul | : | 1.8.5 (274361), CSD as541be (2020) |
| Xtriage (Phenix) | : | 1.13 |
| EDS | : | 2.28 |
| Percentile statistics | : | 20191225.v01 (using entries in the PDB archive December 25th 2019) |
| Refmac | : | 5.8.0158 |
| CCP4 | : | 7.0.044 (Gargrove) |
| Ideal geometry (proteins) | : | Engh & Huber (2001) |
| Ideal geometry (DNA, RNA) | : | Parkinson et al. (1996) |
| Validation Pipeline (wwPDB-VP) | : | 2.28 |

### 1 Overall quality at a glance (i)

The following experimental techniques were used to determine the structure:
*X-RAY DIFFRACTION*

The reported resolution of this entry is 1.50 Å.

Percentile scores (ranging between 0-100) for global validation metrics of the entry are shown in the following graphic. The table shows the number of entries on which the scores are based.

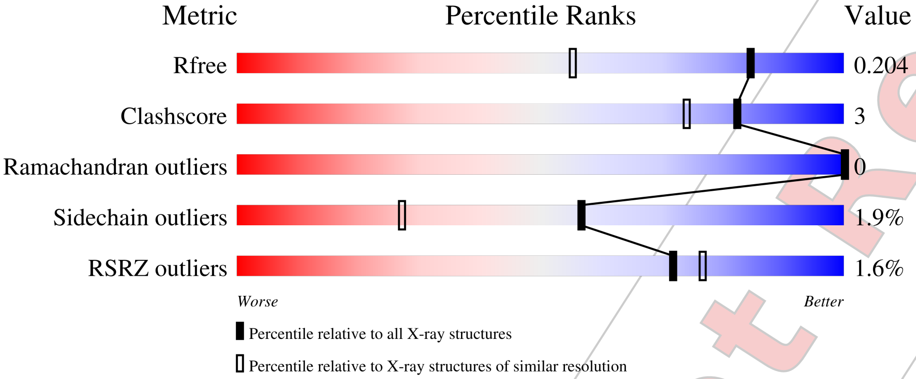


| Metric | Whole archive (#Entries) | Similar resolution (#Entries, resolution range(Å)) |
|---|---|---|
| $R_{free}$ | 130704 | 2936 (1.50-1.50) |
| Clashscore | 141614 | 3144 (1.50-1.50) |
| Ramachandran outliers | 138981 | 3066 (1.50-1.50) |
| Sidechain outliers | 138945 | 3064 (1.50-1.50) |
| RSRZ outliers | 127900 | 2884 (1.50-1.50) |

The table below summarises the geometric issues observed across the polymeric chains and their fit to the electron density. The red, orange, yellow and green segments of the lower bar indicate the fraction of residues that contain outliers for >=3, 2, 1 and 0 types of geometric quality criteria respectively. A grey segment represents the fraction of residues that are not modelled. The numeric value for each fraction is indicated below the corresponding segment, with a dot representing fractions <=5% The upper red bar (where present) indicates the fraction of residues that have poor fit to the electron density. The numeric value is given above the bar.

| Mol | Chain | Length | Quality of chain |
|---|---|---|---|
| 1 | A | 129 | 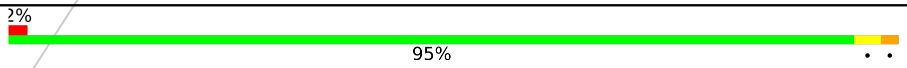  |

For Manuscript Review

### 2 Entry composition (i)

There are 5 unique types of molecules in this entry. The entry contains 1088 atoms, of which 0 are hydrogens and 0 are deuteriums.

In the tables below, the ZeroOcc column contains the number of atoms modelled with zero occupancy, the AltConf column contains the number of residues with at least one atom in alternate conformation and the Trace column contains the number of residues modelled with at most 2 atoms.

- Molecule 1 is a protein called Lysozyme C.

| Mol | Chain | Residues | Atoms | | | | | ZeroOcc | AltConf | Trace |
|---|---|---|---|---|---|---|---|---|---|---|
| | | | Total | C | N | O | S | | | |
| 1 | A | 129 | 1001 | 613 | 193 | 185 | 10 | 0 | 0 | 0 |

- Molecule 2 is SODIUM ION (three-letter code: NA) (formula: Na).

| Mol | Chain | Residues | Atoms | | ZeroOcc | AltConf |
|---|---|---|---|---|---|---|
| | | | Total | Na | | |
| 2 | A | 1 | 1 | 1 | 0 | 0 |

- Molecule 3 is CHLORIDE ION (three-letter code: CL) (formula: Cl).

| Mol | Chain | Residues | Atoms | | ZeroOcc | AltConf |
|---|---|---|---|---|---|---|
| | | | Total | Cl | | |
| 3 | A | 7 | 7 | 7 | 0 | 0 |

- Molecule 4 is GLYCEROL (three-letter code: GOL) (formula: $C_3H_8O_3$).

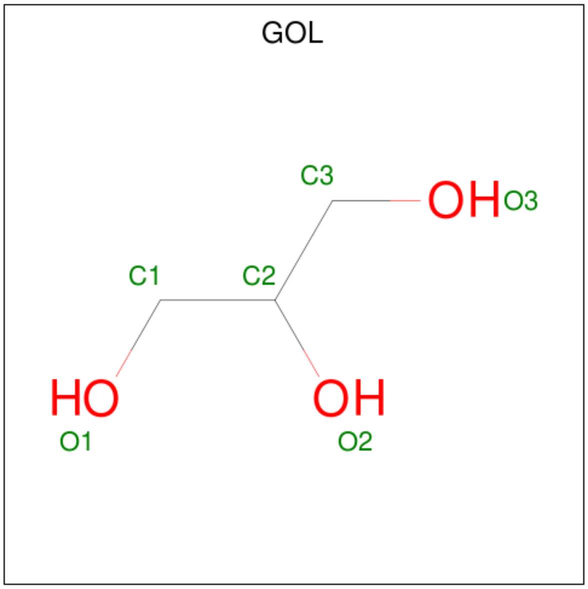


For Manuscript Review

| Mol | Chain | Residues | Atoms | ZeroOcc | AltConf |
|---|---|---|---|---|---|
| 4 | A | 1 | Total C O<br>6 3 3 | 0 | 0 |
| 4 | A | 1 | Total C O<br>6 3 3 | 0 | 0 |

- Molecule 5 is water.

| Mol | Chain | Residues | Atoms | ZeroOcc | AltConf |
|---|---|---|---|---|---|
| 5 | A | 67 | Total O<br>67 67 | 0 | 0 |

For Manuscript Review

### 3 Residue-property plots (i)

These plots are drawn for all protein, RNA, DNA and oligosaccharide chains in the entry. The first graphic for a chain summarises the proportions of the various outlier classes displayed in the second graphic. The second graphic shows the sequence view annotated by issues in geometry and electron density. Residues are color-coded according to the number of geometric quality criteria for which they contain at least one outlier: green = 0, yellow = 1, orange = 2 and red = 3 or more. A red dot above a residue indicates a poor fit to the electron density (RSRZ > 2). Stretches of 2 or more consecutive residues without any outlier are shown as a green connector. Residues present in the sample, but not in the model, are shown in grey.

• Molecule 1: Lysozyme C

Chain A:

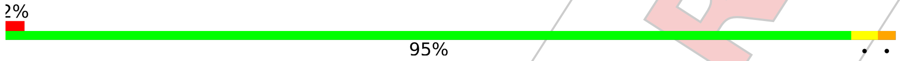


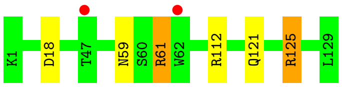


For Manuscript Review

### 4 Data and refinement statistics (i)

| Property | Value | Source |
|---|---|---|
| Space group | P 43 21 2 | Depositor |
| Cell constants<br>a, b, c, $\alpha$, $\beta$, $\gamma$ | 79.32Å 79.32Å 37.76Å<br>90.00° 90.00° 90.00° | Depositor |
| Resolution (Å) | 25.09 – 1.50<br>25.08 – 1.50 | Depositor<br>EDS |
| % Data completeness<br>(in resolution range) | 99.7 (25.09-1.50)<br>99.7 (25.08-1.50) | Depositor<br>EDS |
| $R_{merge}$ | (Not available) | Depositor |
| $R_{sym}$ | (Not available) | Depositor |
| $< I/\sigma(I) >$ [1] | 5.57 (at 1.50Å) | Xtriage |
| Refinement program | REFMAC 5.8.0267 | Depositor |
| R, $R_{free}$ | 0.164 , 0.191<br>0.174 , 0.204 | Depositor<br>DCC |
| $R_{free}$ test set | 1014 reflections (5.10%) | wwPDB-VP |
| Wilson B-factor ($Å^2$) | 12.1 | Xtriage |
| Anisotropy | 0.032 | Xtriage |
| Bulk solvent $k_{sol}$(e/$Å^3$), $B_{sol}$($Å^2$) | 0.35 , 39.7 | EDS |
| L-test for twinning[2] | $< \|L\| >$ = 0.50, $< L^2 >$ = 0.33 | Xtriage |
| Estimated twinning fraction | No twinning to report. | Xtriage |
| $F_o$,$F_c$ correlation | 0.96 | EDS |
| Total number of atoms | 1088 | wwPDB-VP |
| Average B, all atoms ($Å^2$) | 16.0 | wwPDB-VP |

Xtriage's analysis on translational NCS is as follows: *The largest off-origin peak in the Patterson function is 10.33% of the height of the origin peak. No significant pseudotranslation is detected.*

[1]Intensities estimated from amplitudes.

[2]Theoretical values of $< |L| >$, $< L^2 >$ for acentric reflections are 0.5, 0.333 respectively for untwinned datasets, and 0.375, 0.2 for perfectly twinned datasets.

For Manuscript Review

### 5 Model quality (i)

#### 5.1 Standard geometry (i)

Bond lengths and bond angles in the following residue types are not validated in this section: NA, CL, GOL

The Z score for a bond length (or angle) is the number of standard deviations the observed value is removed from the expected value. A bond length (or angle) with $|Z| > 5$ is considered an outlier worth inspection. RMSZ is the root-mean-square of all Z scores of the bond lengths (or angles).

| Mol | Chain | Bond lengths | | Bond angles | |
|---|---|---|---|---|---|
| | | RMSZ | #\|Z\| >5 | RMSZ | #\|Z\| >5 |
| 1 | A | 0.76 | 0/1021 | 0.93 | 2/1379 (0.1%) |

There are no bond length outliers.

All (2) bond angle outliers are listed below:

| Mol | Chain | Res | Type | Atoms | Z | Observed(°) | Ideal(°) |
|---|---|---|---|---|---|---|---|
| 1 | A | 18 | ASP | CB-CG-OD1 | 5.15 | 122.94 | 118.30 |
| 1 | A | 112 | ARG | NE-CZ-NH2 | -5.07 | 117.77 | 120.30 |

There are no chirality outliers.

There are no planarity outliers.

#### 5.2 Too-close contacts (i)

In the following table, the Non-H and H(model) columns list the number of non-hydrogen atoms and hydrogen atoms in the chain respectively. The H(added) column lists the number of hydrogen atoms added and optimized by MolProbity. The Clashes column lists the number of clashes within the asymmetric unit, whereas Symm-Clashes lists symmetry-related clashes.

| Mol | Chain | Non-H | H(model) | H(added) | Clashes | Symm-Clashes |
|---|---|---|---|---|---|---|
| 1 | A | 1001 | 0 | 958 | 5 | 0 |
| 2 | A | 1 | 0 | 0 | 0 | 0 |
| 3 | A | 7 | 0 | 0 | 0 | 0 |
| 4 | A | 12 | 0 | 16 | 0 | 0 |
| 5 | A | 67 | 0 | 0 | 0 | 0 |
| All | All | 1088 | 0 | 974 | 5 | 0 |

The all-atom clashscore is defined as the number of clashes found per 1000 atoms (including hydrogen atoms). The all-atom clashscore for this structure is 3.

For Manuscript Review

All (5) close contacts within the same asymmetric unit are listed below, sorted by their clash magnitude.

| Atom-1 | Atom-2 | Interatomic distance (Å) | Clash overlap (Å) |
|---|---|---|---|
| 1:A:121:GLN:OE1 | 1:A:125:ARG:NH1 | 2.20 | 0.74 |
| 1:A:59:ASN:ND2 | 1:A:61:ARG:H | 1.95 | 0.64 |
| 1:A:59:ASN:ND2 | 1:A:61:ARG:HB3 | 2.24 | 0.53 |
| 1:A:59:ASN:ND2 | 1:A:61:ARG:CB | 2.79 | 0.45 |
| 1:A:59:ASN:HD22 | 1:A:61:ARG:HB3 | 1.83 | 0.44 |

There are no symmetry-related clashes.

#### 5.3 Torsion angles (i)

##### 5.3.1 Protein backbone (i)

In the following table, the Percentiles column shows the percent Ramachandran outliers of the chain as a percentile score with respect to all X-ray entries followed by that with respect to entries of similar resolution.

The Analysed column shows the number of residues for which the backbone conformation was analysed, and the total number of residues.

| Mol | Chain | Analysed | Favoured | Allowed | Outliers | Percentiles |
|---|---|---|---|---|---|---|
| 1 | A | 127/129 (98%) | 125 (98%) | 2 (2%) | 0 | 100 100 |

There are no Ramachandran outliers to report.

##### 5.3.2 Protein sidechains (i)

In the following table, the Percentiles column shows the percent sidechain outliers of the chain as a percentile score with respect to all X-ray entries followed by that with respect to entries of similar resolution.

The Analysed column shows the number of residues for which the sidechain conformation was analysed, and the total number of residues.

| Mol | Chain | Analysed | Rotameric | Outliers | Percentiles |
|---|---|---|---|---|---|
| 1 | A | 105/105 (100%) | 103 (98%) | 2 (2%) | 57 27 |

All (2) residues with a non-rotameric sidechain are listed below:

For Manuscript Review

| Mol | Chain | Res | Type |
|---|---|---|---|
| 1 | A | 61 | ARG |
| 1 | A | 125 | ARG |

Sometimes sidechains can be flipped to improve hydrogen bonding and reduce clashes. All (4) such sidechains are listed below:

| Mol | Chain | Res | Type |
|---|---|---|---|
| 1 | A | 15 | HIS |
| 1 | A | 46 | ASN |
| 1 | A | 59 | ASN |
| 1 | A | 65 | ASN |

##### 5.3.3 RNA (i)

There are no RNA molecules in this entry.

#### 5.4 Non-standard residues in protein, DNA, RNA chains (i)

There are no non-standard protein/DNA/RNA residues in this entry.

#### 5.5 Carbohydrates (i)

There are no monosaccharides in this entry.

#### 5.6 Ligand geometry (i)

Of 10 ligands modelled in this entry, 8 are monoatomic - leaving 2 for Mogul analysis.

In the following table, the Counts columns list the number of bonds (or angles) for which Mogul statistics could be retrieved, the number of bonds (or angles) that are observed in the model and the number of bonds (or angles) that are defined in the Chemical Component Dictionary. The Link column lists molecule types, if any, to which the group is linked. The Z score for a bond length (or angle) is the number of standard deviations the observed value is removed from the expected value. A bond length (or angle) with $|Z| > 2$ is considered an outlier worth inspection. RMSZ is the root-mean-square of all Z scores of the bond lengths (or angles).

| Mol | Type | Chain | Res | Link | Bond lengths | | | Bond angles | | |
|---|---|---|---|---|---|---|---|---|---|---|
| | | | | | Counts | RMSZ | $\#\|Z\| > 2$ | Counts | RMSZ | $\#\|Z\| > 2$ |
| 4 | GOL | A | 209 | - | 5,5,5 | 0.14 | 0 | 5,5,5 | 0.38 | 0 |
| 4 | GOL | A | 210 | - | 5,5,5 | 0.07 | 0 | 5,5,5 | 0.27 | 0 |

For Manuscript Review

In the following table, the Chirals column lists the number of chiral outliers, the number of chiral centers analysed, the number of these observed in the model and the number defined in the Chemical Component Dictionary. Similar counts are reported in the Torsion and Rings columns. '-' means no outliers of that kind were identified.

| Mol | Type | Chain | Res | Link | Chirals | Torsions | Rings |
|---|---|---|---|---|---|---|---|
| 4 | GOL | A | 209 | - | - | 2/4/4/4 | - |
| 4 | GOL | A | 210 | - | - | 2/4/4/4 | - |

There are no bond length outliers.

There are no bond angle outliers.

There are no chirality outliers.

All (4) torsion outliers are listed below:

| Mol | Chain | Res | Type | Atoms |
|---|---|---|---|---|
| 4 | A | 210 | GOL | C1-C2-C3-O3 |
| 4 | A | 210 | GOL | O2-C2-C3-O3 |
| 4 | A | 209 | GOL | O2-C2-C3-O3 |
| 4 | A | 209 | GOL | C1-C2-C3-O3 |

There are no ring outliers.

No monomer is involved in short contacts.

#### 5.7 Other polymers (i)

There are no such residues in this entry.

#### 5.8 Polymer linkage issues (i)

There are no chain breaks in this entry.

For Manuscript Review

### 6 Fit of model and data (i)

#### 6.1 Protein, DNA and RNA chains (i)

In the following table, the column labelled '#RSRZ> 2' contains the number (and percentage) of RSRZ outliers, followed by percent RSRZ outliers for the chain as percentile scores relative to all X-ray entries and entries of similar resolution. The OWAB column contains the minimum, median, $95^{th}$ percentile and maximum values of the occupancy-weighted average B-factor per residue. The column labelled 'Q< 0.9' lists the number of (and percentage) of residues with an average occupancy less than 0.9.

| Mol | Chain | Analysed | <RSRZ> | #RSRZ>2 | | | OWAB(Å$^2$) | Q<0.9 |
|---|---|---|---|---|---|---|---|---|
| 1 | A | 129/129 (100%) | -0.00 | 2 (1%) | 72 | 77 | 7, 13, 26, 34 | 0 |

All (2) RSRZ outliers are listed below:

| Mol | Chain | Res | Type | RSRZ |
|---|---|---|---|---|
| 1 | A | 62 | TRP | 2.1 |
| 1 | A | 47 | THR | 2.1 |

#### 6.2 Non-standard residues in protein, DNA, RNA chains (i)

There are no non-standard protein/DNA/RNA residues in this entry.

#### 6.3 Carbohydrates (i)

There are no monosaccharides in this entry.

#### 6.4 Ligands (i)

In the following table, the Atoms column lists the number of modelled atoms in the group and the number defined in the chemical component dictionary. The B-factors column lists the minimum, median, $95^{th}$ percentile and maximum values of B factors of atoms in the group. The column labelled 'Q< 0.9' lists the number of atoms with occupancy less than 0.9.

| Mol | Type | Chain | Res | Atoms | RSCC | RSR | B-factors(Å$^2$) | Q<0.9 |
|---|---|---|---|---|---|---|---|---|
| 4 | GOL | A | 210 | 6/6 | 0.58 | 0.36 | 66,66,67,67 | 0 |
| 4 | GOL | A | 209 | 6/6 | 0.61 | 0.28 | 46,49,50,55 | 0 |
| 3 | CL | A | 206 | 1/1 | 0.84 | 0.24 | 46,46,46,46 | 0 |
| 3 | CL | A | 208 | 1/1 | 0.85 | 0.23 | 51,51,51,51 | 0 |
| 3 | CL | A | 205 | 1/1 | 0.90 | 0.18 | 44,44,44,44 | 0 |

*Continued on next page...*

For Manuscript Review

*Continued from previous page...*

| Mol | Type | Chain | Res | Atoms | RSCC | RSR | B-factors($Å^2$) | Q<0.9 |
|---|---|---|---|---|---|---|---|---|
| 3 | CL | A | 207 | 1/1 | 0.94 | 0.15 | 40,40,40,40 | 0 |
| 3 | CL | A | 204 | 1/1 | 0.98 | 0.06 | 23,23,23,23 | 0 |
| 3 | CL | A | 202 | 1/1 | 0.99 | 0.04 | 17,17,17,17 | 0 |
| 2 | NA | A | 201 | 1/1 | 0.99 | 0.06 | 13,13,13,13 | 0 |
| 3 | CL | A | 203 | 1/1 | 1.00 | 0.04 | 23,23,23,23 | 0 |

#### 6.5 Other polymers ⓘ

There are no such residues in this entry.

For Manuscript Review

### Full wwPDB X-ray Structure Validation Report (i)

Jun 8, 2022 – 04:03 PM EDT

| | | |
|---|---|---|
| PDB ID | : | 8D7L |
| Title | : | Crystal structure of hen egg white lysozyme at 250 Kelvin |
| Deposited on | : | 2022-06-07 |
| Resolution | : | 1.50 Å(reported) |

**This wwPDB validation report is for manuscript review**

This is a Full wwPDB X-ray Structure Validation Report.

This report is produced by the wwPDB biocuration pipeline after annotation of the structure.

We welcome your comments at **
A user guide is available at
https://www.wwpdb.org/validation/2017/XrayValidationReportHelp
with specific help available everywhere you see the (i) symbol.

For Manuscript Review

The following versions of software and data (see references (i)) were used in the production of this report:

| | | |
|---|---|---|
| MolProbity | : | 4.02b-467 |
| Mogul | : | 1.8.5 (274361), CSD as541be (2020) |
| Xtriage (Phenix) | : | 1.13 |
| EDS | : | 2.28 |
| Percentile statistics | : | 20191225.v01 (using entries in the PDB archive December 25th 2019) |
| Refmac | : | 5.8.0158 |
| CCP4 | : | 7.0.044 (Gargrove) |
| Ideal geometry (proteins) | : | Engh & Huber (2001) |
| Ideal geometry (DNA, RNA) | : | Parkinson et al. (1996) |
| Validation Pipeline (wwPDB-VP) | : | 2.28 |

### 1 Overall quality at a glance (i)

The following experimental techniques were used to determine the structure:
*X-RAY DIFFRACTION*

The reported resolution of this entry is 1.50 Å.

Percentile scores (ranging between 0-100) for global validation metrics of the entry are shown in the following graphic. The table shows the number of entries on which the scores are based.

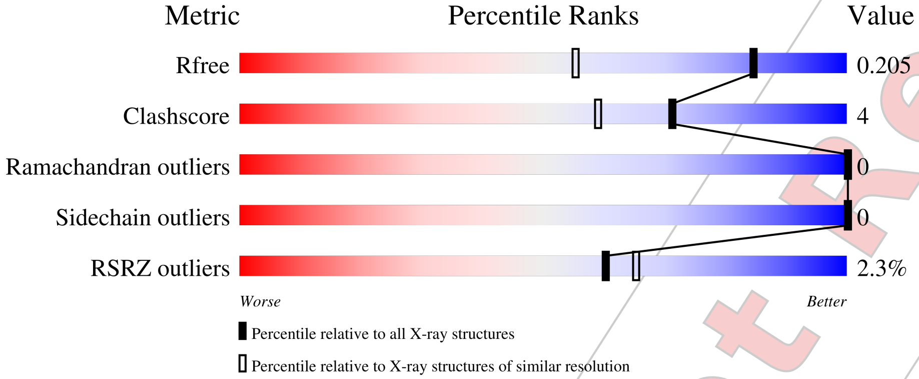


| Metric | Whole archive (#Entries) | Similar resolution (#Entries, resolution range(Å)) |
|---|---|---|
| $R_{free}$ | 130704 | 2936 (1.50-1.50) |
| Clashscore | 141614 | 3144 (1.50-1.50) |
| Ramachandran outliers | 138981 | 3066 (1.50-1.50) |
| Sidechain outliers | 138945 | 3064 (1.50-1.50) |
| RSRZ outliers | 127900 | 2884 (1.50-1.50) |

The table below summarises the geometric issues observed across the polymeric chains and their fit to the electron density. The red, orange, yellow and green segments of the lower bar indicate the fraction of residues that contain outliers for >=3, 2, 1 and 0 types of geometric quality criteria respectively. A grey segment represents the fraction of residues that are not modelled. The numeric value for each fraction is indicated below the corresponding segment, with a dot representing fractions <=5% The upper red bar (where present) indicates the fraction of residues that have poor fit to the electron density. The numeric value is given above the bar.

| Mol | Chain | Length | Quality of chain |
|---|---|---|---|
| 1 | A | 129 | 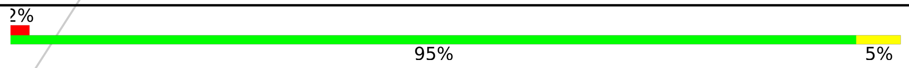 |

For Manuscript Review

### 2 Entry composition (i)

There are 5 unique types of molecules in this entry. The entry contains 1086 atoms, of which 0 are hydrogens and 0 are deuteriums.

In the tables below, the ZeroOcc column contains the number of atoms modelled with zero occupancy, the AltConf column contains the number of residues with at least one atom in alternate conformation and the Trace column contains the number of residues modelled with at most 2 atoms.

- Molecule 1 is a protein called Lysozyme C.

| Mol | Chain | Residues | Atoms | | | | | ZeroOcc | AltConf | Trace |
|---|---|---|---|---|---|---|---|---|---|---|
| | | | Total | C | N | O | S | | | |
| 1 | A | 129 | 1001 | 613 | 193 | 185 | 10 | 0 | 0 | 0 |

- Molecule 2 is SODIUM ION (three-letter code: NA) (formula: Na).

| Mol | Chain | Residues | Atoms | | ZeroOcc | AltConf |
|---|---|---|---|---|---|---|
| | | | Total | Na | | |
| 2 | A | 1 | 1 | 1 | 0 | 0 |

- Molecule 3 is CHLORIDE ION (three-letter code: CL) (formula: Cl).

| Mol | Chain | Residues | Atoms | | ZeroOcc | AltConf |
|---|---|---|---|---|---|---|
| | | | Total | Cl | | |
| 3 | A | 7 | 7 | 7 | 0 | 0 |

- Molecule 4 is GLYCEROL (three-letter code: GOL) (formula: $C_3H_8O_3$).

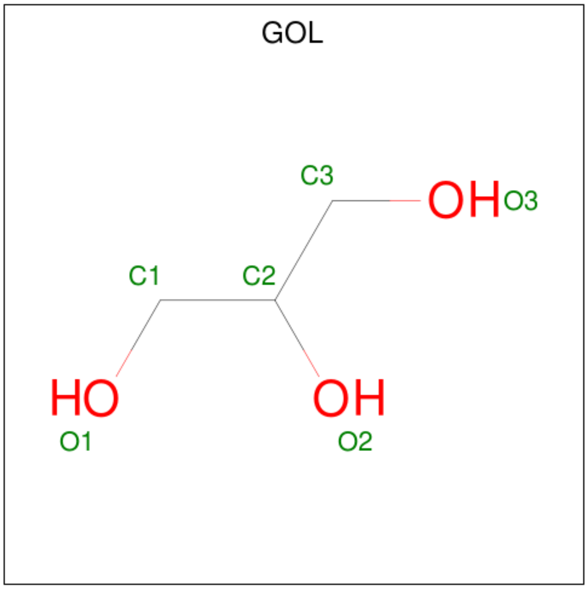


For Manuscript Review

| Mol | Chain | Residues | Atoms | ZeroOcc | AltConf |
|---|---|---|---|---|---|
| 4 | A | 1 | Total C O<br>6 3 3 | 0 | 0 |
| 4 | A | 1 | Total C O<br>6 3 3 | 0 | 0 |

- Molecule 5 is water.

| Mol | Chain | Residues | Atoms | ZeroOcc | AltConf |
|---|---|---|---|---|---|
| 5 | A | 65 | Total O<br>65 65 | 0 | 0 |

For Manuscript Review

### 3 Residue-property plots (i)

These plots are drawn for all protein, RNA, DNA and oligosaccharide chains in the entry. The first graphic for a chain summarises the proportions of the various outlier classes displayed in the second graphic. The second graphic shows the sequence view annotated by issues in geometry and electron density. Residues are color-coded according to the number of geometric quality criteria for which they contain at least one outlier: green = 0, yellow = 1, orange = 2 and red = 3 or more. A red dot above a residue indicates a poor fit to the electron density (RSRZ > 2). Stretches of 2 or more consecutive residues without any outlier are shown as a green connector. Residues present in the sample, but not in the model, are shown in grey.

• Molecule 1: Lysozyme C

Chain A:
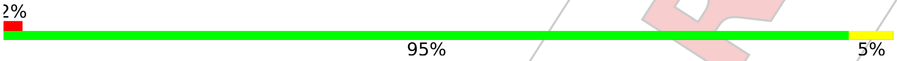


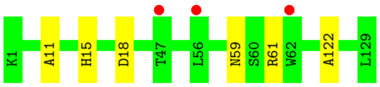


For Manuscript Review

### 4 Data and refinement statistics (i)

| Property | Value | Source |
|---|---|---|
| Space group | P 43 21 2 | Depositor |
| Cell constants<br>a, b, c, $\alpha$, $\beta$, $\gamma$ | 79.45Å 79.45Å 37.77Å<br>90.00° 90.00° 90.00° | Depositor |
| Resolution (Å) | 22.55 – 1.50<br>22.54 – 1.50 | Depositor<br>EDS |
| % Data completeness<br>(in resolution range) | 99.7 (22.55-1.50)<br>99.7 (22.54-1.50) | Depositor<br>EDS |
| $R_{merge}$ | (Not available) | Depositor |
| $R_{sym}$ | (Not available) | Depositor |
| $< I/\sigma(I) >$ [1] | 4.19 (at 1.50Å) | Xtriage |
| Refinement program | REFMAC 5.8.0267 | Depositor |
| R, $R_{free}$ | 0.165 , 0.196<br>0.174 , 0.205 | Depositor<br>DCC |
| $R_{free}$ test set | 1028 reflections (5.17%) | wwPDB-VP |
| Wilson B-factor (Å$^2$) | 14.0 | Xtriage |
| Anisotropy | 0.007 | Xtriage |
| Bulk solvent $k_{sol}$(e/Å$^3$), $B_{sol}$(Å$^2$) | 0.36 , 41.4 | EDS |
| L-test for twinning[2] | $< \|L\| >$ = 0.50, $< L^2 >$ = 0.33 | Xtriage |
| Estimated twinning fraction | No twinning to report. | Xtriage |
| $F_o$,$F_c$ correlation | 0.96 | EDS |
| Total number of atoms | 1086 | wwPDB-VP |
| Average B, all atoms (Å$^2$) | 17.0 | wwPDB-VP |

Xtriage's analysis on translational NCS is as follows: *The largest off-origin peak in the Patterson function is 10.68% of the height of the origin peak. No significant pseudotranslation is detected.*

[1]Intensities estimated from amplitudes.

[2]Theoretical values of $< |L| >$, $< L^2 >$ for acentric reflections are 0.5, 0.333 respectively for untwinned datasets, and 0.375, 0.2 for perfectly twinned datasets.

For Manuscript Review

### 5 Model quality (i)

#### 5.1 Standard geometry (i)

Bond lengths and bond angles in the following residue types are not validated in this section: GOL, NA, CL

The Z score for a bond length (or angle) is the number of standard deviations the observed value is removed from the expected value. A bond length (or angle) with $|Z| > 5$ is considered an outlier worth inspection. RMSZ is the root-mean-square of all Z scores of the bond lengths (or angles).

| Mol | Chain | Bond lengths | | Bond angles | |
|---|---|---|---|---|---|
| | | RMSZ | #\|Z\| >5 | RMSZ | #\|Z\| >5 |
| 1 | A | 0.75 | 0/1021 | 0.91 | 1/1379 (0.1%) |

There are no bond length outliers.

All (1) bond angle outliers are listed below:

| Mol | Chain | Res | Type | Atoms | Z | Observed($^o$) | Ideal($^o$) |
|---|---|---|---|---|---|---|---|
| 1 | A | 18 | ASP | CB-CG-OD1 | 5.52 | 123.27 | 118.30 |

There are no chirality outliers.

There are no planarity outliers.

#### 5.2 Too-close contacts (i)

In the following table, the Non-H and H(model) columns list the number of non-hydrogen atoms and hydrogen atoms in the chain respectively. The H(added) column lists the number of hydrogen atoms added and optimized by MolProbity. The Clashes column lists the number of clashes within the asymmetric unit, whereas Symm-Clashes lists symmetry-related clashes.

| Mol | Chain | Non-H | H(model) | H(added) | Clashes | Symm-Clashes |
|---|---|---|---|---|---|---|
| 1 | A | 1001 | 0 | 958 | 7 | 0 |
| 2 | A | 1 | 0 | 0 | 0 | 0 |
| 3 | A | 7 | 0 | 0 | 0 | 0 |
| 4 | A | 12 | 0 | 16 | 1 | 0 |
| 5 | A | 65 | 0 | 0 | 0 | 0 |
| All | All | 1086 | 0 | 974 | 7 | 0 |

The all-atom clashscore is defined as the number of clashes found per 1000 atoms (including hydrogen atoms). The all-atom clashscore for this structure is 4.

For Manuscript Review

All (7) close contacts within the same asymmetric unit are listed below, sorted by their clash magnitude.

| Atom-1 | Atom-2 | Interatomic distance (Å) | Clash overlap (Å) |
|---|---|---|---|
| 1:A:59:ASN:ND2 | 1:A:61:ARG:HB3 | 2.08 | 0.67 |
| 1:A:59:ASN:ND2 | 1:A:61:ARG:H | 1.95 | 0.64 |
| 1:A:59:ASN:HD22 | 1:A:61:ARG:HB3 | 1.75 | 0.51 |
| 1:A:59:ASN:ND2 | 1:A:61:ARG:CB | 2.73 | 0.50 |
| 1:A:122:ALA:HB1 | 4:A:209:GOL:H32 | 1.94 | 0.49 |
| 1:A:59:ASN:HD21 | 1:A:61:ARG:CB | 2.29 | 0.45 |
| 1:A:11:ALA:O | 1:A:15:HIS:HD2 | 2.03 | 0.42 |

There are no symmetry-related clashes.

#### 5.3 Torsion angles (i)

##### 5.3.1 Protein backbone (i)

In the following table, the Percentiles column shows the percent Ramachandran outliers of the chain as a percentile score with respect to all X-ray entries followed by that with respect to entries of similar resolution.

The Analysed column shows the number of residues for which the backbone conformation was analysed, and the total number of residues.

| Mol | Chain | Analysed | Favoured | Allowed | Outliers | Percentiles | |
|---|---|---|---|---|---|---|---|
| 1 | A | 127/129 (98%) | 126 (99%) | 1 (1%) | 0 | 100 | 100 |

There are no Ramachandran outliers to report.

##### 5.3.2 Protein sidechains (i)

In the following table, the Percentiles column shows the percent sidechain outliers of the chain as a percentile score with respect to all X-ray entries followed by that with respect to entries of similar resolution.

The Analysed column shows the number of residues for which the sidechain conformation was analysed, and the total number of residues.

| Mol | Chain | Analysed | Rotameric | Outliers | Percentiles | |
|---|---|---|---|---|---|---|
| 1 | A | 105/105 (100%) | 105 (100%) | 0 | 100 | 100 |

There are no protein residues with a non-rotameric sidechain to report.

For Manuscript Review

Sometimes sidechains can be flipped to improve hydrogen bonding and reduce clashes. All (4) such sidechains are listed below:

| Mol | Chain | Res | Type |
|---|---|---|---|
| 1 | A | 15 | HIS |
| 1 | A | 46 | ASN |
| 1 | A | 59 | ASN |
| 1 | A | 65 | ASN |

###### 5.3.3 RNA (i)

There are no RNA molecules in this entry.

##### 5.4 Non-standard residues in protein, DNA, RNA chains (i)

There are no non-standard protein/DNA/RNA residues in this entry.

##### 5.5 Carbohydrates (i)

There are no monosaccharides in this entry.

##### 5.6 Ligand geometry (i)

Of 10 ligands modelled in this entry, 8 are monoatomic - leaving 2 for Mogul analysis.

In the following table, the Counts columns list the number of bonds (or angles) for which Mogul statistics could be retrieved, the number of bonds (or angles) that are observed in the model and the number of bonds (or angles) that are defined in the Chemical Component Dictionary. The Link column lists molecule types, if any, to which the group is linked. The Z score for a bond length (or angle) is the number of standard deviations the observed value is removed from the expected value. A bond length (or angle) with $|Z| > 2$ is considered an outlier worth inspection. RMSZ is the root-mean-square of all Z scores of the bond lengths (or angles).

| Mol | Type | Chain | Res | Link | Bond lengths | | | Bond angles | | |
|---|---|---|---|---|---|---|---|---|---|---|
| | | | | | Counts | RMSZ | $\#\|Z\| > 2$ | Counts | RMSZ | $\#\|Z\| > 2$ |
| 4 | GOL | A | 210 | - | 5,5,5 | 0.08 | 0 | 5,5,5 | 0.25 | 0 |
| 4 | GOL | A | 209 | - | 5,5,5 | 0.16 | 0 | 5,5,5 | 0.40 | 0 |

In the following table, the Chirals column lists the number of chiral outliers, the number of chiral centers analysed, the number of these observed in the model and the number defined in the Chemical Component Dictionary. Similar counts are reported in the Torsion and Rings columns. '-' means no outliers of that kind were identified.

For Manuscript Review

| Mol | Type | Chain | Res | Link | Chirals | Torsions | Rings |
|---|---|---|---|---|---|---|---|
| 4 | GOL | A | 210 | - | - | 2/4/4/4 | - |
| 4 | GOL | A | 209 | - | - | 2/4/4/4 | - |

There are no bond length outliers.

There are no bond angle outliers.

There are no chirality outliers.

All (4) torsion outliers are listed below:

| Mol | Chain | Res | Type | Atoms |
|---|---|---|---|---|
| 4 | A | 209 | GOL | C1-C2-C3-O3 |
| 4 | A | 210 | GOL | C1-C2-C3-O3 |
| 4 | A | 210 | GOL | O2-C2-C3-O3 |
| 4 | A | 209 | GOL | O2-C2-C3-O3 |

There are no ring outliers.

1 monomer is involved in 1 short contact:

| Mol | Chain | Res | Type | Clashes | Symm-Clashes |
|---|---|---|---|---|---|
| 4 | A | 209 | GOL | 1 | 0 |

#### 5.7 Other polymers ⓘ

There are no such residues in this entry.

#### 5.8 Polymer linkage issues ⓘ

There are no chain breaks in this entry.

For Manuscript Review

### 6 Fit of model and data (i)

#### 6.1 Protein, DNA and RNA chains (i)

In the following table, the column labelled '#RSRZ> 2' contains the number (and percentage) of RSRZ outliers, followed by percent RSRZ outliers for the chain as percentile scores relative to all X-ray entries and entries of similar resolution. The OWAB column contains the minimum, median, $95^{th}$ percentile and maximum values of the occupancy-weighted average B-factor per residue. The column labelled 'Q< 0.9' lists the number of (and percentage) of residues with an average occupancy less than 0.9.

| Mol | Chain | Analysed | <RSRZ> | #RSRZ>2 | | | OWAB(Å$^2$) | Q<0.9 |
|---|---|---|---|---|---|---|---|---|
| 1 | A | 129/129 (100%) | 0.16 | 3 (2%) | 60 | 65 | 9, 14, 28, 35 | 0 |

All (3) RSRZ outliers are listed below:

| Mol | Chain | Res | Type | RSRZ |
|---|---|---|---|---|
| 1 | A | 47 | THR | 2.7 |
| 1 | A | 62 | TRP | 2.4 |
| 1 | A | 56 | LEU | 2.1 |

#### 6.2 Non-standard residues in protein, DNA, RNA chains (i)

There are no non-standard protein/DNA/RNA residues in this entry.

#### 6.3 Carbohydrates (i)

There are no monosaccharides in this entry.

#### 6.4 Ligands (i)

In the following table, the Atoms column lists the number of modelled atoms in the group and the number defined in the chemical component dictionary. The B-factors column lists the minimum, median, $95^{th}$ percentile and maximum values of B factors of atoms in the group. The column labelled 'Q< 0.9' lists the number of atoms with occupancy less than 0.9.

| Mol | Type | Chain | Res | Atoms | RSCC | RSR | B-factors(Å$^2$) | Q<0.9 |
|---|---|---|---|---|---|---|---|---|
| 4 | GOL | A | 209 | 6/6 | 0.69 | 0.30 | 49,51,53,53 | 0 |
| 4 | GOL | A | 210 | 6/6 | 0.74 | 0.35 | 63,65,67,69 | 0 |
| 3 | CL | A | 206 | 1/1 | 0.86 | 0.25 | 46,46,46,46 | 0 |
| 3 | CL | A | 205 | 1/1 | 0.90 | 0.19 | 42,42,42,42 | 0 |

*Continued on next page...*

For Manuscript Review

*Continued from previous page...*

| Mol | Type | Chain | Res | Atoms | RSCC | RSR | B-factors($Å^2$) | Q<0.9 |
|---|---|---|---|---|---|---|---|---|
| 3 | CL | A | 208 | 1/1 | 0.93 | 0.29 | 46,46,46,46 | 0 |
| 3 | CL | A | 207 | 1/1 | 0.95 | 0.12 | 37,37,37,37 | 0 |
| 3 | CL | A | 203 | 1/1 | 0.99 | 0.03 | 22,22,22,22 | 0 |
| 3 | CL | A | 204 | 1/1 | 0.99 | 0.10 | 23,23,23,23 | 0 |
| 2 | NA | A | 201 | 1/1 | 0.99 | 0.06 | 14,14,14,14 | 0 |
| 3 | CL | A | 202 | 1/1 | 0.99 | 0.04 | 18,18,18,18 | 0 |

#### 6.5 Other polymers (i)

There are no such residues in this entry.

For Manuscript Review

### Full wwPDB X-ray Structure Validation Report (i)

Jun 8, 2022 – 04:05 PM EDT

| | | |
|---|---|---|
| PDB ID | : | 8D7Q |
| Title | : | Crystal structure of hen egg white lysozyme at 250 Kelvin (Triplicate) |
| Deposited on | : | 2022-06-07 |
| Resolution | : | 1.50 Å(reported) |

**This wwPDB validation report is for manuscript review**

This is a Full wwPDB X-ray Structure Validation Report.

This report is produced by the wwPDB biocuration pipeline after annotation of the structure.

We welcome your comments at **
A user guide is available at
https://www.wwpdb.org/validation/2017/XrayValidationReportHelp
with specific help available everywhere you see the (i) symbol.

The following versions of software and data (see references (i)) were used in the production of this report:

| | | |
|---|---|---|
| MolProbity | : | 4.02b-467 |
| Mogul | : | 1.8.5 (274361), CSD as541be (2020) |
| Xtriage (Phenix) | : | 1.13 |
| EDS | : | 2.28 |
| Percentile statistics | : | 20191225.v01 (using entries in the PDB archive December 25th 2019) |
| Refmac | : | 5.8.0158 |
| CCP4 | : | 7.0.044 (Gargrove) |
| Ideal geometry (proteins) | : | Engh & Huber (2001) |
| Ideal geometry (DNA, RNA) | : | Parkinson et al. (1996) |
| Validation Pipeline (wwPDB-VP) | : | 2.28 |

For Manuscript Review

### 1 Overall quality at a glance (i)

The following experimental techniques were used to determine the structure:
*X-RAY DIFFRACTION*

The reported resolution of this entry is 1.50 Å.

Percentile scores (ranging between 0-100) for global validation metrics of the entry are shown in the following graphic. The table shows the number of entries on which the scores are based.

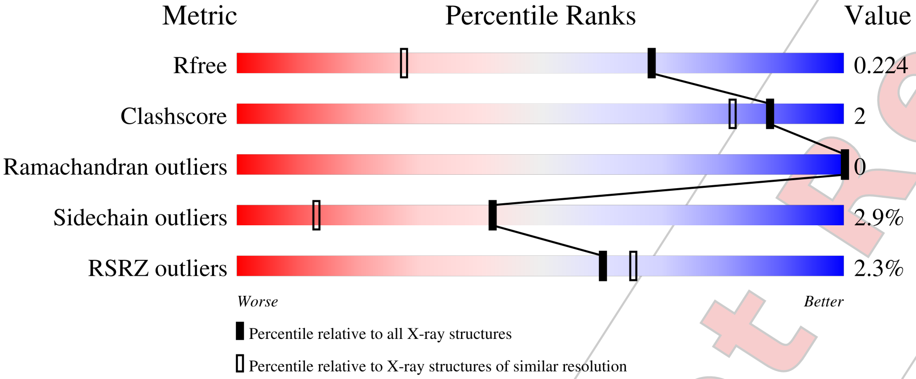


| Metric | Whole archive (#Entries) | Similar resolution (#Entries, resolution range(Å)) |
|---|---|---|
| $R_{free}$ | 130704 | 2936 (1.50-1.50) |
| Clashscore | 141614 | 3144 (1.50-1.50) |
| Ramachandran outliers | 138981 | 3066 (1.50-1.50) |
| Sidechain outliers | 138945 | 3064 (1.50-1.50) |
| RSRZ outliers | 127900 | 2884 (1.50-1.50) |

The table below summarises the geometric issues observed across the polymeric chains and their fit to the electron density. The red, orange, yellow and green segments of the lower bar indicate the fraction of residues that contain outliers for >=3, 2, 1 and 0 types of geometric quality criteria respectively. A grey segment represents the fraction of residues that are not modelled. The numeric value for each fraction is indicated below the corresponding segment, with a dot representing fractions <=5% The upper red bar (where present) indicates the fraction of residues that have poor fit to the electron density. The numeric value is given above the bar.

| Mol | Chain | Length | Quality of chain |
|---|---|---|---|
| 1 | A | 129 | 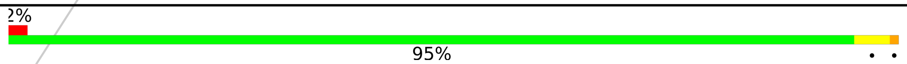 |

The following table lists non-polymeric compounds, carbohydrate monomers and non-standard residues in protein, DNA, RNA chains that are outliers for geometric or electron-density-fit criteria:

| Mol | Type | Chain | Res | Chirality | Geometry | Clashes | Electron density |
|---|---|---|---|---|---|---|---|
| 4 | GOL | A | 209 | - | - | - | X |

*Continued on next page...*

For Manuscript Review

*Continued from previous page...*

| Mol | Type | Chain | Res | Chirality | Geometry | Clashes | Electron density |
|---|---|---|---|---|---|---|---|
| 4 | GOL | A | 210 | - | - | - | X |

For Manuscript Review

### 2 Entry composition (i)

There are 5 unique types of molecules in this entry. The entry contains 1094 atoms, of which 0 are hydrogens and 0 are deuteriums.

In the tables below, the ZeroOcc column contains the number of atoms modelled with zero occupancy, the AltConf column contains the number of residues with at least one atom in alternate conformation and the Trace column contains the number of residues modelled with at most 2 atoms.

- Molecule 1 is a protein called Lysozyme C.

| Mol | Chain | Residues | Atoms | | | | | ZeroOcc | AltConf | Trace |
|---|---|---|---|---|---|---|---|---|---|---|
| | | | Total | C | N | O | S | | | |
| 1 | A | 129 | 1001 | 613 | 193 | 185 | 10 | 0 | 0 | 0 |

- Molecule 2 is SODIUM ION (three-letter code: NA) (formula: Na).

| Mol | Chain | Residues | Atoms | | ZeroOcc | AltConf |
|---|---|---|---|---|---|---|
| | | | Total | Na | | |
| 2 | A | 1 | 1 | 1 | 0 | 0 |

- Molecule 3 is CHLORIDE ION (three-letter code: CL) (formula: Cl).

| Mol | Chain | Residues | Atoms | | ZeroOcc | AltConf |
|---|---|---|---|---|---|---|
| | | | Total | Cl | | |
| 3 | A | 7 | 7 | 7 | 0 | 0 |

- Molecule 4 is GLYCEROL (three-letter code: GOL) (formula: $C_3H_8O_3$).

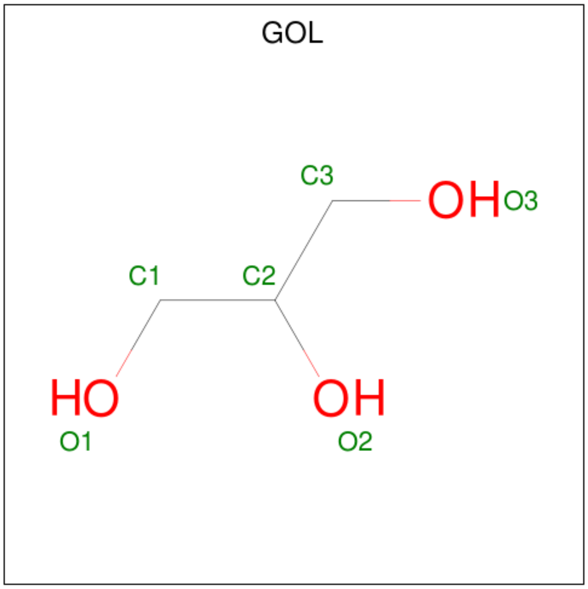


For Manuscript Review

| Mol | Chain | Residues | Atoms | ZeroOcc | AltConf |
|---|---|---|---|---|---|
| 4 | A | 1 | Total C O<br>6 3 3 | 0 | 0 |
| 4 | A | 1 | Total C O<br>6 3 3 | 0 | 0 |

- Molecule 5 is water.

| Mol | Chain | Residues | Atoms | ZeroOcc | AltConf |
|---|---|---|---|---|---|
| 5 | A | 73 | Total O<br>73 73 | 0 | 0 |

For Manuscript Review

### 3 Residue-property plots (i)

These plots are drawn for all protein, RNA, DNA and oligosaccharide chains in the entry. The first graphic for a chain summarises the proportions of the various outlier classes displayed in the second graphic. The second graphic shows the sequence view annotated by issues in geometry and electron density. Residues are color-coded according to the number of geometric quality criteria for which they contain at least one outlier: green = 0, yellow = 1, orange = 2 and red = 3 or more. A red dot above a residue indicates a poor fit to the electron density (RSRZ > 2). Stretches of 2 or more consecutive residues without any outlier are shown as a green connector. Residues present in the sample, but not in the model, are shown in grey.

- Molecule 1: Lysozyme C

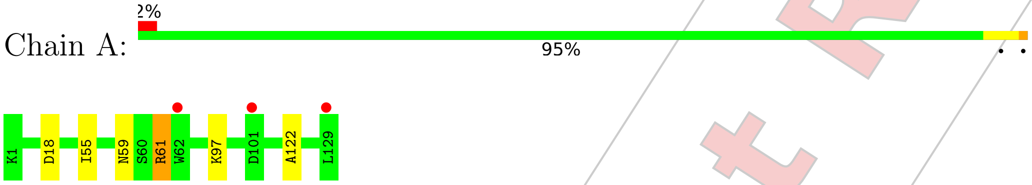


For Manuscript Review

### 4 Data and refinement statistics (i)

| Property | Value | Source |
|---|---|---|
| Space group | P 43 21 2 | Depositor |
| Cell constants<br>a, b, c, $\alpha$, $\beta$, $\gamma$ | 79.17Å 79.17Å 37.72Å<br>90.00° 90.00° 90.00° | Depositor |
| Resolution (Å) | 28.01 – 1.50<br>27.99 – 1.50 | Depositor<br>EDS |
| % Data completeness<br>(in resolution range) | 98.5 (28.01-1.50)<br>98.5 (27.99-1.50) | Depositor<br>EDS |
| $R_{merge}$ | (Not available) | Depositor |
| $R_{sym}$ | (Not available) | Depositor |
| $< I/\sigma(I) >$ [1] | 2.16 (at 1.50Å) | Xtriage |
| Refinement program | REFMAC 5.8.0267 | Depositor |
| R, $R_{free}$ | 0.186 , 0.220<br>0.193 , 0.224 | Depositor<br>DCC |
| $R_{free}$ test set | 1005 reflections (5.16%) | wwPDB-VP |
| Wilson B-factor (Å$^2$) | 16.1 | Xtriage |
| Anisotropy | 0.081 | Xtriage |
| Bulk solvent $k_{sol}$(e/Å$^3$), $B_{sol}$(Å$^2$) | 0.35 , 37.8 | EDS |
| L-test for twinning[2] | $< \|L\| >$ = 0.51, $< L^2 >$ = 0.35 | Xtriage |
| Estimated twinning fraction | No twinning to report. | Xtriage |
| $F_o$,$F_c$ correlation | 0.96 | EDS |
| Total number of atoms | 1094 | wwPDB-VP |
| Average B, all atoms (Å$^2$) | 19.0 | wwPDB-VP |

Xtriage's analysis on translational NCS is as follows: *The largest off-origin peak in the Patterson function is 11.31% of the height of the origin peak. No significant pseudotranslation is detected.*

[1]Intensities estimated from amplitudes.

[2]Theoretical values of $< |L| >$, $< L^2 >$ for acentric reflections are 0.5, 0.333 respectively for untwinned datasets, and 0.375, 0.2 for perfectly twinned datasets.

For Manuscript Review

### 5 Model quality (i)

#### 5.1 Standard geometry (i)

Bond lengths and bond angles in the following residue types are not validated in this section: GOL, NA, CL

The Z score for a bond length (or angle) is the number of standard deviations the observed value is removed from the expected value. A bond length (or angle) with $|Z| > 5$ is considered an outlier worth inspection. RMSZ is the root-mean-square of all Z scores of the bond lengths (or angles).

| Mol | Chain | Bond lengths | | Bond angles | |
|---|---|---|---|---|---|
| | | RMSZ | #\|Z\| >5 | RMSZ | #\|Z\| >5 |
| 1 | A | 0.78 | 0/1021 | 0.94 | 1/1379 (0.1%) |

There are no bond length outliers.

All (1) bond angle outliers are listed below:

| Mol | Chain | Res | Type | Atoms | Z | Observed($^o$) | Ideal($^o$) |
|---|---|---|---|---|---|---|---|
| 1 | A | 18 | ASP | CB-CG-OD1 | 6.75 | 124.37 | 118.30 |

There are no chirality outliers.

There are no planarity outliers.

#### 5.2 Too-close contacts (i)

In the following table, the Non-H and H(model) columns list the number of non-hydrogen atoms and hydrogen atoms in the chain respectively. The H(added) column lists the number of hydrogen atoms added and optimized by MolProbity. The Clashes column lists the number of clashes within the asymmetric unit, whereas Symm-Clashes lists symmetry-related clashes.

| Mol | Chain | Non-H | H(model) | H(added) | Clashes | Symm-Clashes |
|---|---|---|---|---|---|---|
| 1 | A | 1001 | 0 | 958 | 4 | 0 |
| 2 | A | 1 | 0 | 0 | 0 | 0 |
| 3 | A | 7 | 0 | 0 | 0 | 0 |
| 4 | A | 12 | 0 | 16 | 2 | 0 |
| 5 | A | 73 | 0 | 0 | 0 | 1 |
| All | All | 1094 | 0 | 974 | 4 | 1 |

The all-atom clashscore is defined as the number of clashes found per 1000 atoms (including hydrogen atoms). The all-atom clashscore for this structure is 2.

For Manuscript Review

All (4) close contacts within the same asymmetric unit are listed below, sorted by their clash magnitude.

| Atom-1 | Atom-2 | Interatomic distance (Å) | Clash overlap (Å) |
|---|---|---|---|
| 1:A:59:ASN:ND2 | 1:A:61:ARG:H | 1.93 | 0.66 |
| 1:A:59:ASN:ND2 | 1:A:61:ARG:HB3 | 2.15 | 0.62 |
| 1:A:122:ALA:CB | 4:A:209:GOL:H32 | 2.50 | 0.42 |
| 1:A:122:ALA:HB1 | 4:A:209:GOL:H32 | 2.03 | 0.40 |

All (1) symmetry-related close contacts are listed below. The label for Atom-2 includes the symmetry operator and encoded unit-cell translations to be applied.

| Atom-1 | Atom-2 | Interatomic distance (Å) | Clash overlap (Å) |
|---|---|---|---|
| 5:A:353:HOH:O | 5:A:353:HOH:O[8_554] | 2.09 | 0.11 |

#### 5.3 Torsion angles (i)

##### 5.3.1 Protein backbone (i)

In the following table, the Percentiles column shows the percent Ramachandran outliers of the chain as a percentile score with respect to all X-ray entries followed by that with respect to entries of similar resolution.

The Analysed column shows the number of residues for which the backbone conformation was analysed, and the total number of residues.

| Mol | Chain | Analysed | Favoured | Allowed | Outliers | Percentiles | |
|---|---|---|---|---|---|---|---|
| 1 | A | 127/129 (98%) | 126 (99%) | 1 (1%) | 0 | 100 | 100 |

There are no Ramachandran outliers to report.

##### 5.3.2 Protein sidechains (i)

In the following table, the Percentiles column shows the percent sidechain outliers of the chain as a percentile score with respect to all X-ray entries followed by that with respect to entries of similar resolution.

The Analysed column shows the number of residues for which the sidechain conformation was analysed, and the total number of residues.

| Mol | Chain | Analysed | Rotameric | Outliers | Percentiles | |
|---|---|---|---|---|---|---|
| 1 | A | 105/105 (100%) | 102 (97%) | 3 (3%) | 42 | 13 |

For Manuscript Review

All (3) residues with a non-rotameric sidechain are listed below:

| Mol | Chain | Res | Type |
|---|---|---|---|
| 1 | A | 55 | ILE |
| 1 | A | 61 | ARG |
| 1 | A | 97 | LYS |

Sometimes sidechains can be flipped to improve hydrogen bonding and reduce clashes. All (4) such sidechains are listed below:

| Mol | Chain | Res | Type |
|---|---|---|---|
| 1 | A | 15 | HIS |
| 1 | A | 46 | ASN |
| 1 | A | 59 | ASN |
| 1 | A | 65 | ASN |

##### 5.3.3 RNA (i)

There are no RNA molecules in this entry.

#### 5.4 Non-standard residues in protein, DNA, RNA chains (i)

There are no non-standard protein/DNA/RNA residues in this entry.

#### 5.5 Carbohydrates (i)

There are no monosaccharides in this entry.

#### 5.6 Ligand geometry (i)

Of 10 ligands modelled in this entry, 8 are monoatomic - leaving 2 for Mogul analysis.

In the following table, the Counts columns list the number of bonds (or angles) for which Mogul statistics could be retrieved, the number of bonds (or angles) that are observed in the model and the number of bonds (or angles) that are defined in the Chemical Component Dictionary. The Link column lists molecule types, if any, to which the group is linked. The Z score for a bond length (or angle) is the number of standard deviations the observed value is removed from the expected value. A bond length (or angle) with $|Z| > 2$ is considered an outlier worth inspection. RMSZ is the root-mean-square of all Z scores of the bond lengths (or angles).

| Mol | Type | Chain | Res | Link | Bond lengths | | | Bond angles | | |
|---|---|---|---|---|---|---|---|---|---|---|
| | | | | | Counts | RMSZ | $\#\|Z\| > 2$ | Counts | RMSZ | $\#\|Z\| > 2$ |
| 4 | GOL | A | 210 | - | 5,5,5 | 0.09 | 0 | 5,5,5 | 0.29 | 0 |

For Manuscript Review

| Mol | Type | Chain | Res | Link | Bond lengths | | | Bond angles | | |
|---|---|---|---|---|---|---|---|---|---|---|
| | | | | | Counts | RMSZ | #\|Z\| > 2 | Counts | RMSZ | #\|Z\| > 2 |
| 4 | GOL | A | 209 | - | 5,5,5 | 0.18 | 0 | 5,5,5 | 0.45 | 0 |

In the following table, the Chirals column lists the number of chiral outliers, the number of chiral centers analysed, the number of these observed in the model and the number defined in the Chemical Component Dictionary. Similar counts are reported in the Torsion and Rings columns. '-' means no outliers of that kind were identified.

| Mol | Type | Chain | Res | Link | Chirals | Torsions | Rings |
|---|---|---|---|---|---|---|---|
| 4 | GOL | A | 210 | - | - | 2/4/4/4 | - |
| 4 | GOL | A | 209 | - | - | 2/4/4/4 | - |

There are no bond length outliers.

There are no bond angle outliers.

There are no chirality outliers.

All (4) torsion outliers are listed below:

| Mol | Chain | Res | Type | Atoms |
|---|---|---|---|---|
| 4 | A | 209 | GOL | C1-C2-C3-O3 |
| 4 | A | 210 | GOL | C1-C2-C3-O3 |
| 4 | A | 209 | GOL | O2-C2-C3-O3 |
| 4 | A | 210 | GOL | O2-C2-C3-O3 |

There are no ring outliers.

1 monomer is involved in 2 short contacts:

| Mol | Chain | Res | Type | Clashes | Symm-Clashes |
|---|---|---|---|---|---|
| 4 | A | 209 | GOL | 2 | 0 |

#### 5.7 Other polymers (i)

There are no such residues in this entry.

#### 5.8 Polymer linkage issues (i)

There are no chain breaks in this entry.

For Manuscript Review

### 6 Fit of model and data (i)

#### 6.1 Protein, DNA and RNA chains (i)

In the following table, the column labelled '#RSRZ> 2' contains the number (and percentage) of RSRZ outliers, followed by percent RSRZ outliers for the chain as percentile scores relative to all X-ray entries and entries of similar resolution. The OWAB column contains the minimum, median, $95^{th}$ percentile and maximum values of the occupancy-weighted average B-factor per residue. The column labelled 'Q< 0.9' lists the number of (and percentage) of residues with an average occupancy less than 0.9.

| Mol | Chain | Analysed | <RSRZ> | #RSRZ>2 | | | OWAB(Å$^2$) | Q<0.9 |
|---|---|---|---|---|---|---|---|---|
| 1 | A | 129/129 (100%) | 0.22 | 3 (2%) | 60 | 65 | 10, 16, 28, 37 | 0 |

All (3) RSRZ outliers are listed below:

| Mol | Chain | Res | Type | RSRZ |
|---|---|---|---|---|
| 1 | A | 62 | TRP | 3.8 |
| 1 | A | 129 | LEU | 2.9 |
| 1 | A | 101 | ASP | 2.2 |

#### 6.2 Non-standard residues in protein, DNA, RNA chains (i)

There are no non-standard protein/DNA/RNA residues in this entry.

#### 6.3 Carbohydrates (i)

There are no monosaccharides in this entry.

#### 6.4 Ligands (i)

In the following table, the Atoms column lists the number of modelled atoms in the group and the number defined in the chemical component dictionary. The B-factors column lists the minimum, median, $95^{th}$ percentile and maximum values of B factors of atoms in the group. The column labelled 'Q< 0.9' lists the number of atoms with occupancy less than 0.9.

| Mol | Type | Chain | Res | Atoms | RSCC | RSR | B-factors(Å$^2$) | Q<0.9 |
|---|---|---|---|---|---|---|---|---|
| 4 | GOL | A | 210 | 6/6 | 0.62 | 0.47 | 80,82,82,84 | 0 |
| 4 | GOL | A | 209 | 6/6 | 0.64 | 0.42 | 40,50,61,61 | 0 |
| 3 | CL | A | 208 | 1/1 | 0.90 | 0.24 | 50,50,50,50 | 0 |
| 3 | CL | A | 206 | 1/1 | 0.91 | 0.24 | 45,45,45,45 | 0 |

*Continued on next page...*

For Manuscript Review

*Continued from previous page...*

| Mol | Type | Chain | Res | Atoms | RSCC | RSR | B-factors(Å$^2$) | Q<0.9 |
|---|---|---|---|---|---|---|---|---|
| 3 | CL | A | 207 | 1/1 | 0.93 | 0.09 | 37,37,37,37 | 0 |
| 3 | CL | A | 205 | 1/1 | 0.94 | 0.16 | 43,43,43,43 | 0 |
| 3 | CL | A | 204 | 1/1 | 0.97 | 0.05 | 26,26,26,26 | 0 |
| 3 | CL | A | 202 | 1/1 | 0.99 | 0.04 | 19,19,19,19 | 0 |
| 3 | CL | A | 203 | 1/1 | 0.99 | 0.03 | 23,23,23,23 | 0 |
| 2 | NA | A | 201 | 1/1 | 0.99 | 0.05 | 16,16,16,16 | 0 |

#### 6.5 Other polymers (i)

There are no such residues in this entry.

For Manuscript Review

### Full wwPDB X-ray Structure Validation Report (i)

Jun 8, 2022 – 04:07 PM EDT

| | | |
|---|---|---|
| PDB ID | : | 8D7S |
| Title | : | Crystal structure of hen egg white lysozyme at 275 Kelvin |
| Deposited on | : | 2022-06-07 |
| Resolution | : | 1.50 Å(reported) |

**This wwPDB validation report is for manuscript review**

This is a Full wwPDB X-ray Structure Validation Report.

This report is produced by the wwPDB biocuration pipeline after annotation of the structure.

We welcome your comments at **
A user guide is available at
https://www.wwpdb.org/validation/2017/XrayValidationReportHelp
with specific help available everywhere you see the (i) symbol.

The following versions of software and data (see references (i)) were used in the production of this report:

| | | |
|---|---|---|
| MolProbity | : | 4.02b-467 |
| Mogul | : | 1.8.5 (274361), CSD as541be (2020) |
| Xtriage (Phenix) | : | 1.13 |
| EDS | : | 2.28 |
| Percentile statistics | : | 20191225.v01 (using entries in the PDB archive December 25th 2019) |
| Refmac | : | 5.8.0158 |
| CCP4 | : | 7.0.044 (Gargrove) |
| Ideal geometry (proteins) | : | Engh & Huber (2001) |
| Ideal geometry (DNA, RNA) | : | Parkinson et al. (1996) |
| Validation Pipeline (wwPDB-VP) | : | 2.28 |

For Manuscript Review

### 1 Overall quality at a glance (i)

The following experimental techniques were used to determine the structure:
*X-RAY DIFFRACTION*

The reported resolution of this entry is 1.50 Å.

Percentile scores (ranging between 0-100) for global validation metrics of the entry are shown in the following graphic. The table shows the number of entries on which the scores are based.

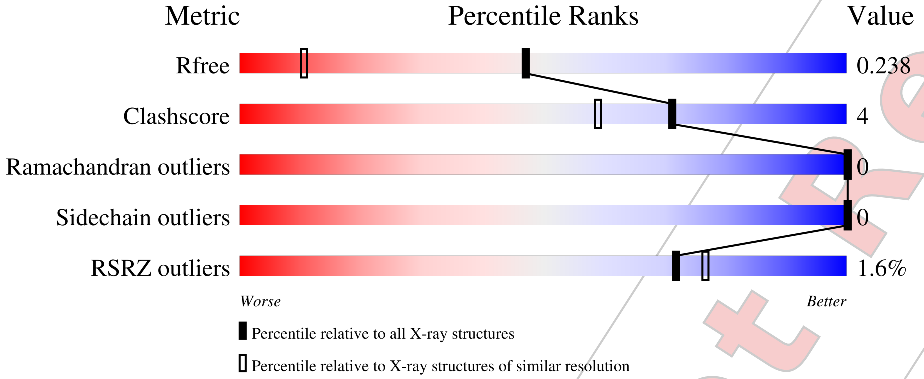


| Metric | Whole archive (#Entries) | Similar resolution (#Entries, resolution range(Å)) |
|---|---|---|
| $R_{free}$ | 130704 | 2936 (1.50-1.50) |
| Clashscore | 141614 | 3144 (1.50-1.50) |
| Ramachandran outliers | 138981 | 3066 (1.50-1.50) |
| Sidechain outliers | 138945 | 3064 (1.50-1.50) |
| RSRZ outliers | 127900 | 2884 (1.50-1.50) |

The table below summarises the geometric issues observed across the polymeric chains and their fit to the electron density. The red, orange, yellow and green segments of the lower bar indicate the fraction of residues that contain outliers for >=3, 2, 1 and 0 types of geometric quality criteria respectively. A grey segment represents the fraction of residues that are not modelled. The numeric value for each fraction is indicated below the corresponding segment, with a dot representing fractions <=5% The upper red bar (where present) indicates the fraction of residues that have poor fit to the electron density. The numeric value is given above the bar.

| Mol | Chain | Length | Quality of chain |
|---|---|---|---|
| 1 | A | 129 | 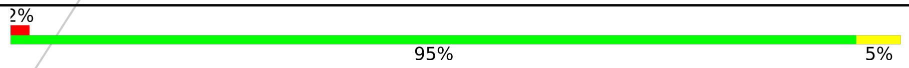 |



For Manuscript Review

### 2 Entry composition (i)

There are 5 unique types of molecules in this entry. The entry contains 1074 atoms, of which 0 are hydrogens and 0 are deuteriums.

In the tables below, the ZeroOcc column contains the number of atoms modelled with zero occupancy, the AltConf column contains the number of residues with at least one atom in alternate conformation and the Trace column contains the number of residues modelled with at most 2 atoms.

- Molecule 1 is a protein called Lysozyme C.

| Mol | Chain | Residues | Atoms | | | | | ZeroOcc | AltConf | Trace |
|---|---|---|---|---|---|---|---|---|---|---|
| | | | Total | C | N | O | S | | | |
| 1 | A | 129 | 1001 | 613 | 193 | 185 | 10 | 0 | 0 | 0 |

- Molecule 2 is SODIUM ION (three-letter code: NA) (formula: Na).

| Mol | Chain | Residues | Atoms | | ZeroOcc | AltConf |
|---|---|---|---|---|---|---|
| | | | Total | Na | | |
| 2 | A | 1 | 1 | 1 | 0 | 0 |

- Molecule 3 is CHLORIDE ION (three-letter code: CL) (formula: Cl).

| Mol | Chain | Residues | Atoms | | ZeroOcc | AltConf |
|---|---|---|---|---|---|---|
| | | | Total | Cl | | |
| 3 | A | 7 | 7 | 7 | 0 | 0 |

- Molecule 4 is GLYCEROL (three-letter code: GOL) (formula: $C_3H_8O_3$).

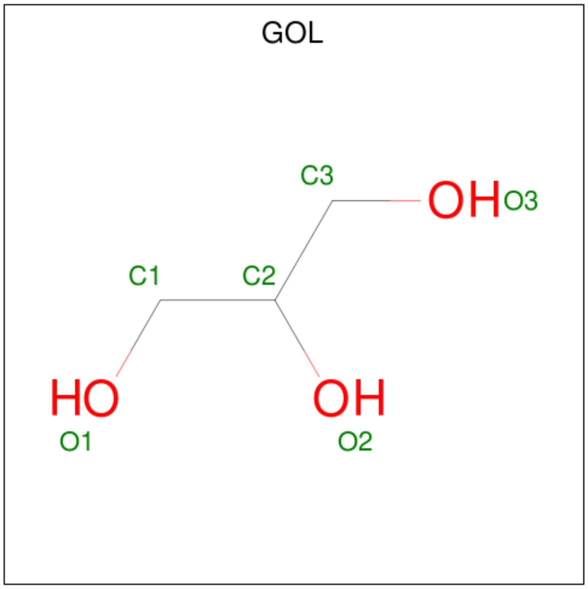


For Manuscript Review

| Mol | Chain | Residues | Atoms | ZeroOcc | AltConf |
|---|---|---|---|---|---|
| 4 | A | 1 | Total C O<br>6 3 3 | 0 | 0 |
| 4 | A | 1 | Total C O<br>6 3 3 | 0 | 0 |

- Molecule 5 is water.

| Mol | Chain | Residues | Atoms | ZeroOcc | AltConf |
|---|---|---|---|---|---|
| 5 | A | 53 | Total O<br>53 53 | 0 | 0 |

For Manuscript Review

### 3 Residue-property plots (i)

These plots are drawn for all protein, RNA, DNA and oligosaccharide chains in the entry. The first graphic for a chain summarises the proportions of the various outlier classes displayed in the second graphic. The second graphic shows the sequence view annotated by issues in geometry and electron density. Residues are color-coded according to the number of geometric quality criteria for which they contain at least one outlier: green = 0, yellow = 1, orange = 2 and red = 3 or more. A red dot above a residue indicates a poor fit to the electron density (RSRZ > 2). Stretches of 2 or more consecutive residues without any outlier are shown as a green connector. Residues present in the sample, but not in the model, are shown in grey.

• Molecule 1: Lysozyme C

Chain A:

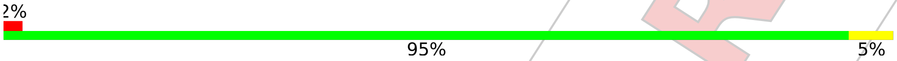


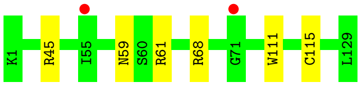


For Manuscript Review

### 4 Data and refinement statistics (i)

| Property | Value | Source |
|---|---|---|
| Space group | P 43 21 2 | Depositor |
| Cell constants<br>a, b, c, $\alpha$, $\beta$, $\gamma$ | 79.79Å 79.79Å 37.78Å<br>90.00° 90.00° 90.00° | Depositor |
| Resolution (Å) | 27.45 – 1.50<br>27.43 – 1.48 | Depositor<br>EDS |
| % Data completeness<br>(in resolution range) | 97.1 (27.45-1.50)<br>97.1 (27.43-1.48) | Depositor<br>EDS |
| $R_{merge}$ | (Not available) | Depositor |
| $R_{sym}$ | (Not available) | Depositor |
| $< I/\sigma(I) >$ [1] | 1.14 (at 1.48Å) | Xtriage |
| Refinement program | REFMAC 5.8.0267 | Depositor |
| R, $R_{free}$ | 0.212 , 0.234<br>0.219 , 0.238 | Depositor<br>DCC |
| $R_{free}$ test set | 1056 reflections (5.17%) | wwPDB-VP |
| Wilson B-factor (Å$^2$) | 13.4 | Xtriage |
| Anisotropy | 0.144 | Xtriage |
| Bulk solvent $k_{sol}$(e/Å$^3$), $B_{sol}$(Å$^2$) | 0.41 , 43.5 | EDS |
| L-test for twinning[2] | $< \|L\| >= 0.47$, $< L^2 >= 0.31$ | Xtriage |
| Estimated twinning fraction | No twinning to report. | Xtriage |
| $F_o$,$F_c$ correlation | 0.95 | EDS |
| Total number of atoms | 1074 | wwPDB-VP |
| Average B, all atoms (Å$^2$) | 20.0 | wwPDB-VP |

Xtriage's analysis on translational NCS is as follows: *The largest off-origin peak in the Patterson function is 11.60% of the height of the origin peak. No significant pseudotranslation is detected.*

[1] Intensities estimated from amplitudes.

[2] Theoretical values of $< |L| >$, $< L^2 >$ for acentric reflections are 0.5, 0.333 respectively for untwinned datasets, and 0.375, 0.2 for perfectly twinned datasets.

For Manuscript Review

### 5 Model quality (i)

#### 5.1 Standard geometry (i)

Bond lengths and bond angles in the following residue types are not validated in this section: GOL, CL, NA

The Z score for a bond length (or angle) is the number of standard deviations the observed value is removed from the expected value. A bond length (or angle) with $|Z| > 5$ is considered an outlier worth inspection. RMSZ is the root-mean-square of all Z scores of the bond lengths (or angles).

| Mol | Chain | Bond lengths | | Bond angles | |
|---|---|---|---|---|---|
| | | RMSZ | #$\|Z\|$ >5 | RMSZ | #$\|Z\|$ >5 |
| 1 | A | 0.80 | 0/1021 | 0.90 | 0/1379 |

There are no bond length outliers.

There are no bond angle outliers.

There are no chirality outliers.

There are no planarity outliers.

#### 5.2 Too-close contacts (i)

In the following table, the Non-H and H(model) columns list the number of non-hydrogen atoms and hydrogen atoms in the chain respectively. The H(added) column lists the number of hydrogen atoms added and optimized by MolProbity. The Clashes column lists the number of clashes within the asymmetric unit, whereas Symm-Clashes lists symmetry-related clashes.

| Mol | Chain | Non-H | H(model) | H(added) | Clashes | Symm-Clashes |
|---|---|---|---|---|---|---|
| 1 | A | 1001 | 0 | 958 | 7 | 0 |
| 2 | A | 1 | 0 | 0 | 0 | 0 |
| 3 | A | 7 | 0 | 0 | 0 | 0 |
| 4 | A | 12 | 0 | 16 | 0 | 0 |
| 5 | A | 53 | 0 | 0 | 0 | 0 |
| All | All | 1074 | 0 | 974 | 7 | 0 |

The all-atom clashscore is defined as the number of clashes found per 1000 atoms (including hydrogen atoms). The all-atom clashscore for this structure is 4.

All (7) close contacts within the same asymmetric unit are listed below, sorted by their clash magnitude.

For Manuscript Review

| Atom-1 | Atom-2 | Interatomic distance (Å) | Clash overlap (Å) |
|---|---|---|---|
| 1:A:59:ASN:ND2 | 1:A:61:ARG:HB3 | 1.95 | 0.80 |
| 1:A:59:ASN:HD21 | 1:A:61:ARG:HB3 | 1.63 | 0.60 |
| 1:A:59:ASN:HD22 | 1:A:61:ARG:HB3 | 1.79 | 0.47 |
| 1:A:45:ARG:NE | 1:A:68:ARG:NH2 | 2.66 | 0.44 |
| 1:A:111:TRP:CD1 | 1:A:115:CYS:HB2 | 2.54 | 0.43 |
| 1:A:59:ASN:ND2 | 1:A:61:ARG:H | 2.18 | 0.41 |
| 1:A:59:ASN:HD21 | 1:A:61:ARG:CB | 2.29 | 0.41 |

There are no symmetry-related clashes.

#### 5.3 Torsion angles (i)

##### 5.3.1 Protein backbone (i)

In the following table, the Percentiles column shows the percent Ramachandran outliers of the chain as a percentile score with respect to all X-ray entries followed by that with respect to entries of similar resolution.

The Analysed column shows the number of residues for which the backbone conformation was analysed, and the total number of residues.

| Mol | Chain | Analysed | Favoured | Allowed | Outliers | Percentiles |
|---|---|---|---|---|---|---|
| 1 | A | 127/129 (98%) | 125 (98%) | 2 (2%) | 0 | 100 100 |

There are no Ramachandran outliers to report.

##### 5.3.2 Protein sidechains (i)

In the following table, the Percentiles column shows the percent sidechain outliers of the chain as a percentile score with respect to all X-ray entries followed by that with respect to entries of similar resolution.

The Analysed column shows the number of residues for which the sidechain conformation was analysed, and the total number of residues.

| Mol | Chain | Analysed | Rotameric | Outliers | Percentiles |
|---|---|---|---|---|---|
| 1 | A | 105/105 (100%) | 105 (100%) | 0 | 100 100 |

There are no protein residues with a non-rotameric sidechain to report.

Sometimes sidechains can be flipped to improve hydrogen bonding and reduce clashes. All (3) such sidechains are listed below:

For Manuscript Review

| Mol | Chain | Res | Type |
|---|---|---|---|
| 1 | A | 15 | HIS |
| 1 | A | 46 | ASN |
| 1 | A | 59 | ASN |

##### 5.3.3 RNA (i)

There are no RNA molecules in this entry.

#### 5.4 Non-standard residues in protein, DNA, RNA chains (i)

There are no non-standard protein/DNA/RNA residues in this entry.

#### 5.5 Carbohydrates (i)

There are no monosaccharides in this entry.

#### 5.6 Ligand geometry (i)

Of 10 ligands modelled in this entry, 8 are monoatomic - leaving 2 for Mogul analysis.

In the following table, the Counts columns list the number of bonds (or angles) for which Mogul statistics could be retrieved, the number of bonds (or angles) that are observed in the model and the number of bonds (or angles) that are defined in the Chemical Component Dictionary. The Link column lists molecule types, if any, to which the group is linked. The Z score for a bond length (or angle) is the number of standard deviations the observed value is removed from the expected value. A bond length (or angle) with $|Z| > 2$ is considered an outlier worth inspection. RMSZ is the root-mean-square of all Z scores of the bond lengths (or angles).

| Mol | Type | Chain | Res | Link | Bond lengths | | | Bond angles | | |
|---|---|---|---|---|---|---|---|---|---|---|
| | | | | | Counts | RMSZ | $\#\|Z\| > 2$ | Counts | RMSZ | $\#\|Z\| > 2$ |
| 4 | GOL | A | 210 | - | 5,5,5 | 0.11 | 0 | 5,5,5 | 0.33 | 0 |
| 4 | GOL | A | 209 | - | 5,5,5 | 0.10 | 0 | 5,5,5 | 0.38 | 0 |

In the following table, the Chirals column lists the number of chiral outliers, the number of chiral centers analysed, the number of these observed in the model and the number defined in the Chemical Component Dictionary. Similar counts are reported in the Torsion and Rings columns. '-' means no outliers of that kind were identified.

| Mol | Type | Chain | Res | Link | Chirals | Torsions | Rings |
|---|---|---|---|---|---|---|---|
| 4 | GOL | A | 210 | - | - | 2/4/4/4 | - |
| 4 | GOL | A | 209 | - | - | 0/4/4/4 | - |

For Manuscript Review

There are no bond length outliers.

There are no bond angle outliers.

There are no chirality outliers.

All (2) torsion outliers are listed below:

| Mol | Chain | Res | Type | Atoms |
|---|---|---|---|---|
| 4 | A | 210 | GOL | C1-C2-C3-O3 |
| 4 | A | 210 | GOL | O2-C2-C3-O3 |

There are no ring outliers.

No monomer is involved in short contacts.

#### 5.7 Other polymers (i)

There are no such residues in this entry.

#### 5.8 Polymer linkage issues (i)

There are no chain breaks in this entry.

For Manuscript Review

### 6 Fit of model and data (i)

#### 6.1 Protein, DNA and RNA chains (i)

In the following table, the column labelled '#RSRZ> 2' contains the number (and percentage) of RSRZ outliers, followed by percent RSRZ outliers for the chain as percentile scores relative to all X-ray entries and entries of similar resolution. The OWAB column contains the minimum, median, $95^{th}$ percentile and maximum values of the occupancy-weighted average B-factor per residue. The column labelled 'Q< 0.9' lists the number of (and percentage) of residues with an average occupancy less than 0.9.

| Mol | Chain | Analysed | <RSRZ> | #RSRZ>2 | | | OWAB(Å$^2$) | Q<0.9 |
|---|---|---|---|---|---|---|---|---|
| 1 | A | 129/129 (100%) | 0.07 | 2 (1%) | 72 | 77 | 10, 17, 31, 44 | 0 |

All (2) RSRZ outliers are listed below:

| Mol | Chain | Res | Type | RSRZ |
|---|---|---|---|---|
| 1 | A | 55 | ILE | 2.4 |
| 1 | A | 71 | GLY | 2.0 |

#### 6.2 Non-standard residues in protein, DNA, RNA chains (i)

There are no non-standard protein/DNA/RNA residues in this entry.

#### 6.3 Carbohydrates (i)

There are no monosaccharides in this entry.

#### 6.4 Ligands (i)

In the following table, the Atoms column lists the number of modelled atoms in the group and the number defined in the chemical component dictionary. The B-factors column lists the minimum, median, $95^{th}$ percentile and maximum values of B factors of atoms in the group. The column labelled 'Q< 0.9' lists the number of atoms with occupancy less than 0.9.

| Mol | Type | Chain | Res | Atoms | RSCC | RSR | B-factors(Å$^2$) | Q<0.9 |
|---|---|---|---|---|---|---|---|---|
| 4 | GOL | A | 210 | 6/6 | 0.56 | 0.24 | 56,58,60,68 | 0 |
| 4 | GOL | A | 209 | 6/6 | 0.64 | 0.27 | 45,48,52,55 | 0 |
| 3 | CL | A | 206 | 1/1 | 0.77 | 0.25 | 54,54,54,54 | 0 |
| 3 | CL | A | 208 | 1/1 | 0.88 | 0.12 | 48,48,48,48 | 0 |
| 3 | CL | A | 205 | 1/1 | 0.89 | 0.13 | 53,53,53,53 | 0 |

*Continued on next page...*

For Manuscript Review

*Continued from previous page...*

| Mol | Type | Chain | Res | Atoms | RSCC | RSR | B-factors($Å^2$) | Q<0.9 |
|---|---|---|---|---|---|---|---|---|
| 3 | CL | A | 207 | 1/1 | 0.91 | 0.10 | 46,46,46,46 | 0 |
| 3 | CL | A | 203 | 1/1 | 0.99 | 0.04 | 24,24,24,24 | 0 |
| 3 | CL | A | 204 | 1/1 | 0.99 | 0.03 | 27,27,27,27 | 0 |
| 2 | NA | A | 201 | 1/1 | 0.99 | 0.04 | 16,16,16,16 | 0 |
| 3 | CL | A | 202 | 1/1 | 0.99 | 0.04 | 18,18,18,18 | 0 |

#### 6.5 Other polymers (i)

There are no such residues in this entry.

For Manuscript Review

### Full wwPDB X-ray Structure Validation Report (i)

Jun 8, 2022 – 04:09 PM EDT

| | | |
|---|---|---|
| PDB ID | : | 8D8A |
| Title | : | Crystal structure of hen egg white lysozyme at 275 Kelvin (Duplicate) |
| Deposited on | : | 2022-06-08 |
| Resolution | : | 1.50 Å(reported) |

**This wwPDB validation report is for manuscript review**

This is a Full wwPDB X-ray Structure Validation Report.

This report is produced by the wwPDB biocuration pipeline after annotation of the structure.

We welcome your comments at **
A user guide is available at
https://www.wwpdb.org/validation/2017/XrayValidationReportHelp
with specific help available everywhere you see the (i) symbol.

The following versions of software and data (see references (i)) were used in the production of this report:

| | | |
|---|---|---|
| MolProbity | : | 4.02b-467 |
| Mogul | : | 1.8.5 (274361), CSD as541be (2020) |
| Xtriage (Phenix) | : | 1.13 |
| EDS | : | 2.28 |
| Percentile statistics | : | 20191225.v01 (using entries in the PDB archive December 25th 2019) |
| Refmac | : | 5.8.0158 |
| CCP4 | : | 7.0.044 (Gargrove) |
| Ideal geometry (proteins) | : | Engh & Huber (2001) |
| Ideal geometry (DNA, RNA) | : | Parkinson et al. (1996) |
| Validation Pipeline (wwPDB-VP) | : | 2.28 |

For Manuscript Review

### 1 Overall quality at a glance (i)

The following experimental techniques were used to determine the structure:
*X-RAY DIFFRACTION*

The reported resolution of this entry is 1.50 Å.

Percentile scores (ranging between 0-100) for global validation metrics of the entry are shown in the following graphic. The table shows the number of entries on which the scores are based.

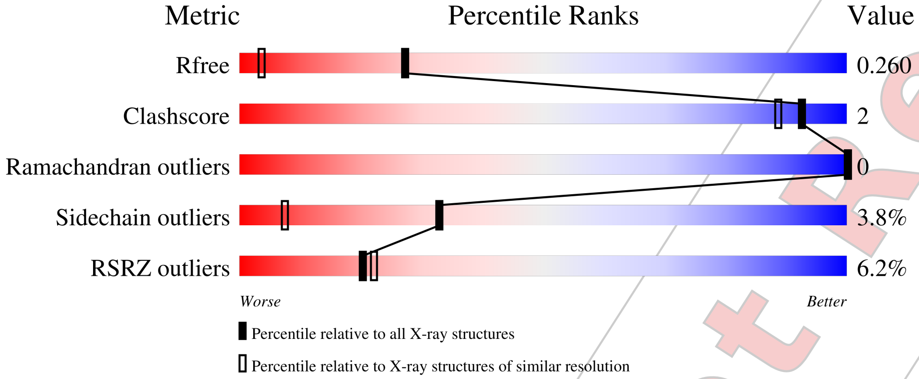


| Metric | Whole archive (#Entries) | Similar resolution (#Entries, resolution range(Å)) |
|---|---|---|
| $R_{free}$ | 130704 | 2936 (1.50-1.50) |
| Clashscore | 141614 | 3144 (1.50-1.50) |
| Ramachandran outliers | 138981 | 3066 (1.50-1.50) |
| Sidechain outliers | 138945 | 3064 (1.50-1.50) |
| RSRZ outliers | 127900 | 2884 (1.50-1.50) |

The table below summarises the geometric issues observed across the polymeric chains and their fit to the electron density. The red, orange, yellow and green segments of the lower bar indicate the fraction of residues that contain outliers for >=3, 2, 1 and 0 types of geometric quality criteria respectively. A grey segment represents the fraction of residues that are not modelled. The numeric value for each fraction is indicated below the corresponding segment, with a dot representing fractions <=5% The upper red bar (where present) indicates the fraction of residues that have poor fit to the electron density. The numeric value is given above the bar.

| Mol | Chain | Length | Quality of chain |
|---|---|---|---|
| 1 | A | 129 | 6% / 93% 7% |

For Manuscript Review

### 2 Entry composition ⓘ

There are 5 unique types of molecules in this entry. The entry contains 1064 atoms, of which 0 are hydrogens and 0 are deuteriums.

In the tables below, the ZeroOcc column contains the number of atoms modelled with zero occupancy, the AltConf column contains the number of residues with at least one atom in alternate conformation and the Trace column contains the number of residues modelled with at most 2 atoms.

- Molecule 1 is a protein called Lysozyme C.

| Mol | Chain | Residues | Atoms | | | | | ZeroOcc | AltConf | Trace |
|---|---|---|---|---|---|---|---|---|---|---|
| | | | Total | C | N | O | S | | | |
| 1 | A | 129 | 1001 | 613 | 193 | 185 | 10 | 0 | 0 | 0 |

- Molecule 2 is SODIUM ION (three-letter code: NA) (formula: Na).

| Mol | Chain | Residues | Atoms | | ZeroOcc | AltConf |
|---|---|---|---|---|---|---|
| | | | Total | Na | | |
| 2 | A | 1 | 1 | 1 | 0 | 0 |

- Molecule 3 is CHLORIDE ION (three-letter code: CL) (formula: Cl).

| Mol | Chain | Residues | Atoms | | ZeroOcc | AltConf |
|---|---|---|---|---|---|---|
| | | | Total | Cl | | |
| 3 | A | 7 | 7 | 7 | 0 | 0 |

- Molecule 4 is GLYCEROL (three-letter code: GOL) (formula: $C_3H_8O_3$).

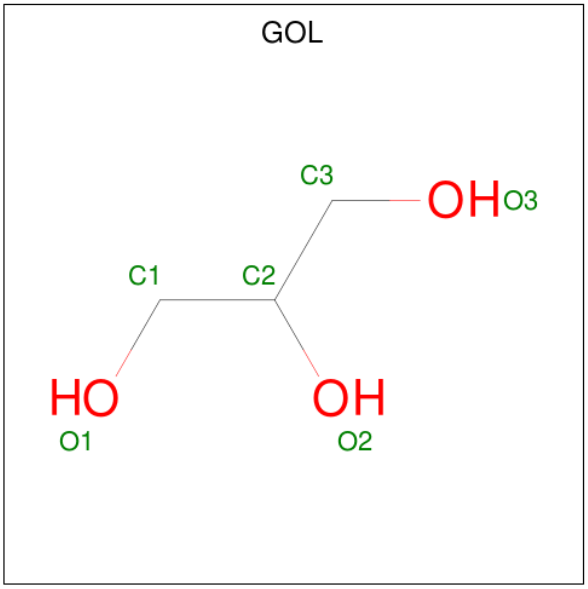


For Manuscript Review

| Mol | Chain | Residues | Atoms | ZeroOcc | AltConf |
|---|---|---|---|---|---|
| 4 | A | 1 | Total C O<br>6 3 3 | 0 | 0 |
| 4 | A | 1 | Total C O<br>6 3 3 | 0 | 0 |

- Molecule 5 is water.

| Mol | Chain | Residues | Atoms | ZeroOcc | AltConf |
|---|---|---|---|---|---|
| 5 | A | 43 | Total O<br>43 43 | 0 | 0 |

For Manuscript Review

### 3 Residue-property plots (i)

These plots are drawn for all protein, RNA, DNA and oligosaccharide chains in the entry. The first graphic for a chain summarises the proportions of the various outlier classes displayed in the second graphic. The second graphic shows the sequence view annotated by issues in geometry and electron density. Residues are color-coded according to the number of geometric quality criteria for which they contain at least one outlier: green = 0, yellow = 1, orange = 2 and red = 3 or more. A red dot above a residue indicates a poor fit to the electron density (RSRZ > 2). Stretches of 2 or more consecutive residues without any outlier are shown as a green connector. Residues present in the sample, but not in the model, are shown in grey.

• Molecule 1: Lysozyme C

Chain A:

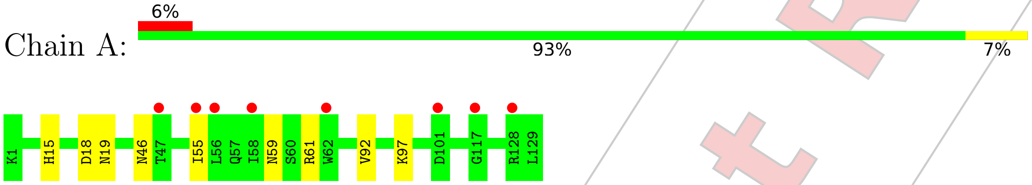


For Manuscript Review

### 4 Data and refinement statistics (i)

| Property | Value | Source |
|---|---|---|
| Space group | P 43 21 2 | Depositor |
| Cell constants<br>a, b, c, $\alpha$, $\beta$, $\gamma$ | 79.72Å 79.72Å 37.43Å<br>90.00° 90.00° 90.00° | Depositor |
| Resolution (Å) | 22.53 – 1.50<br>22.52 – 1.49 | Depositor<br>EDS |
| % Data completeness<br>(in resolution range) | 93.9 (22.53-1.50)<br>93.9 (22.52-1.49) | Depositor<br>EDS |
| $R_{merge}$ | (Not available) | Depositor |
| $R_{sym}$ | (Not available) | Depositor |
| $< I/\sigma(I) >$ [1] | 1.20 (at 1.49Å) | Xtriage |
| Refinement program | REFMAC 5.8.0267 | Depositor |
| R, $R_{free}$ | 0.230 , 0.260<br>0.231 , 0.260 | Depositor<br>DCC |
| $R_{free}$ test set | 973 reflections (5.08%) | wwPDB-VP |
| Wilson B-factor (Å$^2$) | 18.1 | Xtriage |
| Anisotropy | 0.035 | Xtriage |
| Bulk solvent $k_{sol}$(e/Å$^3$), $B_{sol}$(Å$^2$) | 0.43 , 44.5 | EDS |
| L-test for twinning[2] | $< \|L\| > = 0.43$, $< L^2 > = 0.25$ | Xtriage |
| Estimated twinning fraction | No twinning to report. | Xtriage |
| $F_o$,$F_c$ correlation | 0.94 | EDS |
| Total number of atoms | 1064 | wwPDB-VP |
| Average B, all atoms (Å$^2$) | 22.0 | wwPDB-VP |

Xtriage's analysis on translational NCS is as follows: *The largest off-origin peak in the Patterson function is 8.24% of the height of the origin peak. No significant pseudotranslation is detected.*

[1] Intensities estimated from amplitudes.

[2] Theoretical values of $< |L| >$, $< L^2 >$ for acentric reflections are 0.5, 0.333 respectively for untwinned datasets, and 0.375, 0.2 for perfectly twinned datasets.

For Manuscript Review

### 5 Model quality (i)

#### 5.1 Standard geometry (i)

Bond lengths and bond angles in the following residue types are not validated in this section: NA, GOL, CL

The Z score for a bond length (or angle) is the number of standard deviations the observed value is removed from the expected value. A bond length (or angle) with $|Z| > 5$ is considered an outlier worth inspection. RMSZ is the root-mean-square of all Z scores of the bond lengths (or angles).

| Mol | Chain | Bond lengths | | Bond angles | |
|---|---|---|---|---|---|
| | | RMSZ | $\#\|Z\| >5$ | RMSZ | $\#\|Z\| >5$ |
| 1 | A | 0.83 | 0/1021 | 0.97 | 0/1379 |

There are no bond length outliers.

There are no bond angle outliers.

There are no chirality outliers.

There are no planarity outliers.

#### 5.2 Too-close contacts (i)

In the following table, the Non-H and H(model) columns list the number of non-hydrogen atoms and hydrogen atoms in the chain respectively. The H(added) column lists the number of hydrogen atoms added and optimized by MolProbity. The Clashes column lists the number of clashes within the asymmetric unit, whereas Symm-Clashes lists symmetry-related clashes.

| Mol | Chain | Non-H | H(model) | H(added) | Clashes | Symm-Clashes |
|---|---|---|---|---|---|---|
| 1 | A | 1001 | 0 | 958 | 3 | 0 |
| 2 | A | 1 | 0 | 0 | 0 | 0 |
| 3 | A | 7 | 0 | 0 | 0 | 0 |
| 4 | A | 12 | 0 | 16 | 0 | 0 |
| 5 | A | 43 | 0 | 0 | 0 | 1 |
| All | All | 1064 | 0 | 974 | 3 | 1 |

The all-atom clashscore is defined as the number of clashes found per 1000 atoms (including hydrogen atoms). The all-atom clashscore for this structure is 2.

All (3) close contacts within the same asymmetric unit are listed below, sorted by their clash magnitude.

For Manuscript Review

| Atom-1 | Atom-2 | Interatomic distance (Å) | Clash overlap (Å) |
|---|---|---|---|
| 1:A:59:ASN:ND2 | 1:A:61:ARG:HB3 | 2.22 | 0.54 |
| 1:A:15:HIS:HB3 | 1:A:92:VAL:HG11 | 1.98 | 0.45 |
| 1:A:46:ASN:ND2 | 1:A:59:ASN:OD1 | 2.54 | 0.41 |

All (1) symmetry-related close contacts are listed below. The label for Atom-2 includes the symmetry operator and encoded unit-cell translations to be applied.

| Atom-1 | Atom-2 | Interatomic distance (Å) | Clash overlap (Å) |
|---|---|---|---|
| 5:A:303:HOH:O | 5:A:303:HOH:O[7_554] | 1.22 | 0.98 |

#### 5.3 Torsion angles (i)

##### 5.3.1 Protein backbone (i)

In the following table, the Percentiles column shows the percent Ramachandran outliers of the chain as a percentile score with respect to all X-ray entries followed by that with respect to entries of similar resolution.

The Analysed column shows the number of residues for which the backbone conformation was analysed, and the total number of residues.

| Mol | Chain | Analysed | Favoured | Allowed | Outliers | Percentiles |
|---|---|---|---|---|---|---|
| 1 | A | 127/129 (98%) | 124 (98%) | 3 (2%) | 0 | 100 100 |

There are no Ramachandran outliers to report.

##### 5.3.2 Protein sidechains (i)

In the following table, the Percentiles column shows the percent sidechain outliers of the chain as a percentile score with respect to all X-ray entries followed by that with respect to entries of similar resolution.

The Analysed column shows the number of residues for which the sidechain conformation was analysed, and the total number of residues.

| Mol | Chain | Analysed | Rotameric | Outliers | Percentiles |
|---|---|---|---|---|---|
| 1 | A | 105/105 (100%) | 101 (96%) | 4 (4%) | 33 7 |

All (4) residues with a non-rotameric sidechain are listed below:

For Manuscript Review

| Mol | Chain | Res | Type |
|---|---|---|---|
| 1 | A | 18 | ASP |
| 1 | A | 19 | ASN |
| 1 | A | 55 | ILE |
| 1 | A | 97 | LYS |

Sometimes sidechains can be flipped to improve hydrogen bonding and reduce clashes. All (6) such sidechains are listed below:

| Mol | Chain | Res | Type |
|---|---|---|---|
| 1 | A | 15 | HIS |
| 1 | A | 19 | ASN |
| 1 | A | 44 | ASN |
| 1 | A | 46 | ASN |
| 1 | A | 59 | ASN |
| 1 | A | 65 | ASN |

###### 5.3.3 RNA (i)

There are no RNA molecules in this entry.

##### 5.4 Non-standard residues in protein, DNA, RNA chains (i)

There are no non-standard protein/DNA/RNA residues in this entry.

##### 5.5 Carbohydrates (i)

There are no monosaccharides in this entry.

##### 5.6 Ligand geometry (i)

Of 10 ligands modelled in this entry, 8 are monoatomic - leaving 2 for Mogul analysis.

In the following table, the Counts columns list the number of bonds (or angles) for which Mogul statistics could be retrieved, the number of bonds (or angles) that are observed in the model and the number of bonds (or angles) that are defined in the Chemical Component Dictionary. The Link column lists molecule types, if any, to which the group is linked. The Z score for a bond length (or angle) is the number of standard deviations the observed value is removed from the expected value. A bond length (or angle) with $|Z| > 2$ is considered an outlier worth inspection. RMSZ is the root-mean-square of all Z scores of the bond lengths (or angles).

For Manuscript Review

| Mol | Type | Chain | Res | Link | Bond lengths | | | Bond angles | | |
|---|---|---|---|---|---|---|---|---|---|---|
| | | | | | Counts | RMSZ | $\#\|Z\| > 2$ | Counts | RMSZ | $\#\|Z\| > 2$ |
| 4 | GOL | A | 210 | - | 5,5,5 | 0.11 | 0 | 5,5,5 | 0.26 | 0 |
| 4 | GOL | A | 209 | - | 5,5,5 | 0.12 | 0 | 5,5,5 | 0.25 | 0 |

In the following table, the Chirals column lists the number of chiral outliers, the number of chiral centers analysed, the number of these observed in the model and the number defined in the Chemical Component Dictionary. Similar counts are reported in the Torsion and Rings columns. '-' means no outliers of that kind were identified.

| Mol | Type | Chain | Res | Link | Chirals | Torsions | Rings |
|---|---|---|---|---|---|---|---|
| 4 | GOL | A | 210 | - | - | 0/4/4/4 | - |
| 4 | GOL | A | 209 | - | - | 1/4/4/4 | - |

There are no bond length outliers.

There are no bond angle outliers.

There are no chirality outliers.

All (1) torsion outliers are listed below:

| Mol | Chain | Res | Type | Atoms |
|---|---|---|---|---|
| 4 | A | 209 | GOL | O1-C1-C2-O2 |

There are no ring outliers.

No monomer is involved in short contacts.

#### 5.7 Other polymers (i)

There are no such residues in this entry.

#### 5.8 Polymer linkage issues (i)

There are no chain breaks in this entry.

For Manuscript Review

### 6 Fit of model and data (i)

#### 6.1 Protein, DNA and RNA chains (i)

In the following table, the column labelled '#RSRZ> 2' contains the number (and percentage) of RSRZ outliers, followed by percent RSRZ outliers for the chain as percentile scores relative to all X-ray entries and entries of similar resolution. The OWAB column contains the minimum, median, $95^{th}$ percentile and maximum values of the occupancy-weighted average B-factor per residue. The column labelled 'Q< 0.9' lists the number of (and percentage) of residues with an average occupancy less than 0.9.

| Mol | Chain | Analysed | <RSRZ> | #RSRZ>2 | | | OWAB(Å$^2$) | Q<0.9 |
|---|---|---|---|---|---|---|---|---|
| 1 | A | 129/129 (100%) | 0.57 | 8 (6%) | 20 | 22 | 13, 20, 33, 39 | 0 |

All (8) RSRZ outliers are listed below:

| Mol | Chain | Res | Type | RSRZ |
|---|---|---|---|---|
| 1 | A | 47 | THR | 4.0 |
| 1 | A | 56 | LEU | 3.2 |
| 1 | A | 101 | ASP | 3.1 |
| 1 | A | 62 | TRP | 2.6 |
| 1 | A | 58 | ILE | 2.3 |
| 1 | A | 55 | ILE | 2.1 |
| 1 | A | 117 | GLY | 2.1 |
| 1 | A | 128 | ARG | 2.1 |

#### 6.2 Non-standard residues in protein, DNA, RNA chains (i)

There are no non-standard protein/DNA/RNA residues in this entry.

#### 6.3 Carbohydrates (i)

There are no monosaccharides in this entry.

#### 6.4 Ligands (i)

In the following table, the Atoms column lists the number of modelled atoms in the group and the number defined in the chemical component dictionary. The B-factors column lists the minimum, median, $95^{th}$ percentile and maximum values of B factors of atoms in the group. The column labelled 'Q< 0.9' lists the number of atoms with occupancy less than 0.9.

For Manuscript Review

| Mol | Type | Chain | Res | Atoms | RSCC | RSR | B-factors($Å^2$) | Q<0.9 |
|---|---|---|---|---|---|---|---|---|
| 4 | GOL | A | 210 | 6/6 | 0.39 | 0.30 | 49,54,56,57 | 0 |
| 3 | CL | A | 207 | 1/1 | 0.60 | 0.12 | 52,52,52,52 | 0 |
| 4 | GOL | A | 209 | 6/6 | 0.74 | 0.27 | 45,53,57,60 | 0 |
| 3 | CL | A | 206 | 1/1 | 0.86 | 0.20 | 56,56,56,56 | 0 |
| 3 | CL | A | 205 | 1/1 | 0.86 | 0.09 | 54,54,54,54 | 0 |
| 3 | CL | A | 208 | 1/1 | 0.94 | 0.19 | 42,42,42,42 | 0 |
| 3 | CL | A | 204 | 1/1 | 0.96 | 0.05 | 31,31,31,31 | 0 |
| 3 | CL | A | 203 | 1/1 | 0.98 | 0.04 | 28,28,28,28 | 0 |
| 2 | NA | A | 201 | 1/1 | 0.99 | 0.07 | 20,20,20,20 | 0 |
| 3 | CL | A | 202 | 1/1 | 0.99 | 0.05 | 20,20,20,20 | 0 |

#### 6.5 Other polymers ⓘ

There are no such residues in this entry.

For Manuscript Review

### Full wwPDB X-ray Structure Validation Report (i)

Jun 9, 2022 – 11:26 AM EDT

| | | |
|---|---|---|
| PDB ID | : | 8D8B |
| Title | : | Crystal structure of hen egg white lysozyme at 275 Kelvin (Triplicate) |
| Deposited on | : | 2022-06-08 |
| Resolution | : | 1.50 Å(reported) |

**This wwPDB validation report is for manuscript review**

This is a Full wwPDB X-ray Structure Validation Report.

This report is produced by the wwPDB biocuration pipeline after annotation of the structure.

We welcome your comments at **
A user guide is available at
https://www.wwpdb.org/validation/2017/XrayValidationReportHelp
with specific help available everywhere you see the (i) symbol.

The following versions of software and data (see references (i)) were used in the production of this report:

| | | |
|---|---|---|
| MolProbity | : | 4.02b-467 |
| Mogul | : | 1.8.5 (274361), CSD as541be (2020) |
| Xtriage (Phenix) | : | 1.13 |
| EDS | : | 2.28 |
| Percentile statistics | : | 20191225.v01 (using entries in the PDB archive December 25th 2019) |
| Refmac | : | 5.8.0158 |
| CCP4 | : | 7.0.044 (Gargrove) |
| Ideal geometry (proteins) | : | Engh & Huber (2001) |
| Ideal geometry (DNA, RNA) | : | Parkinson et al. (1996) |
| Validation Pipeline (wwPDB-VP) | : | 2.28 |

For Manuscript Review

### 1 Overall quality at a glance

The following experimental techniques were used to determine the structure:
*X-RAY DIFFRACTION*

The reported resolution of this entry is 1.50 Å.

Percentile scores (ranging between 0-100) for global validation metrics of the entry are shown in the following graphic. The table shows the number of entries on which the scores are based.

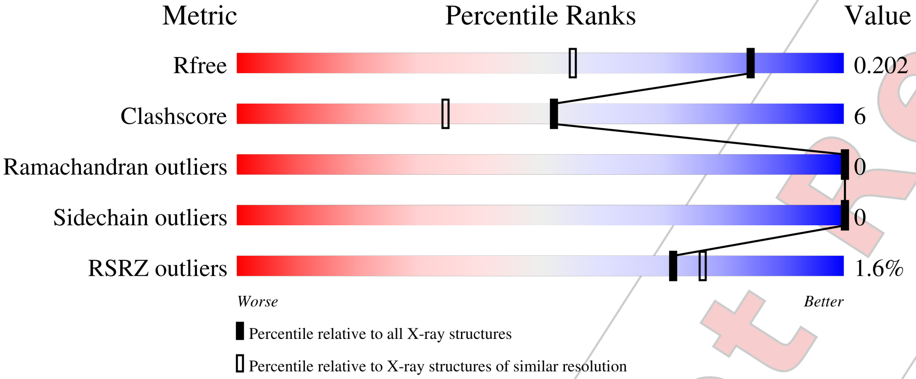


| Metric | Whole archive (#Entries) | Similar resolution (#Entries, resolution range(Å)) |
|---|---|---|
| $R_{free}$ | 130704 | 2936 (1.50-1.50) |
| Clashscore | 141614 | 3144 (1.50-1.50) |
| Ramachandran outliers | 138981 | 3066 (1.50-1.50) |
| Sidechain outliers | 138945 | 3064 (1.50-1.50) |
| RSRZ outliers | 127900 | 2884 (1.50-1.50) |

The table below summarises the geometric issues observed across the polymeric chains and their fit to the electron density. The red, orange, yellow and green segments of the lower bar indicate the fraction of residues that contain outliers for >=3, 2, 1 and 0 types of geometric quality criteria respectively. A grey segment represents the fraction of residues that are not modelled. The numeric value for each fraction is indicated below the corresponding segment, with a dot representing fractions <=5% The upper red bar (where present) indicates the fraction of residues that have poor fit to the electron density. The numeric value is given above the bar.

| Mol | Chain | Length | Quality of chain |
|---|---|---|---|
| 1 | A | 129 | 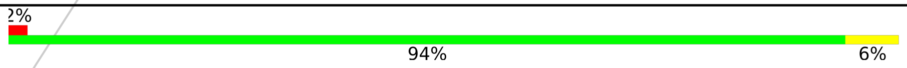  |

For Manuscript Review

### 2 Entry composition (i)

There are 5 unique types of molecules in this entry. The entry contains 1072 atoms, of which 0 are hydrogens and 0 are deuteriums.

In the tables below, the ZeroOcc column contains the number of atoms modelled with zero occupancy, the AltConf column contains the number of residues with at least one atom in alternate conformation and the Trace column contains the number of residues modelled with at most 2 atoms.

- Molecule 1 is a protein called Lysozyme C.

| Mol | Chain | Residues | Atoms | | | | | ZeroOcc | AltConf | Trace |
|---|---|---|---|---|---|---|---|---|---|---|
| | | | Total | C | N | O | S | | | |
| 1 | A | 129 | 1001 | 613 | 193 | 185 | 10 | 0 | 0 | 0 |

- Molecule 2 is SODIUM ION (three-letter code: NA) (formula: Na).

| Mol | Chain | Residues | Atoms | | ZeroOcc | AltConf |
|---|---|---|---|---|---|---|
| | | | Total | Na | | |
| 2 | A | 1 | 1 | 1 | 0 | 0 |

- Molecule 3 is CHLORIDE ION (three-letter code: CL) (formula: Cl).

| Mol | Chain | Residues | Atoms | | ZeroOcc | AltConf |
|---|---|---|---|---|---|---|
| | | | Total | Cl | | |
| 3 | A | 7 | 7 | 7 | 0 | 0 |

- Molecule 4 is GLYCEROL (three-letter code: GOL) (formula: $C_3H_8O_3$).

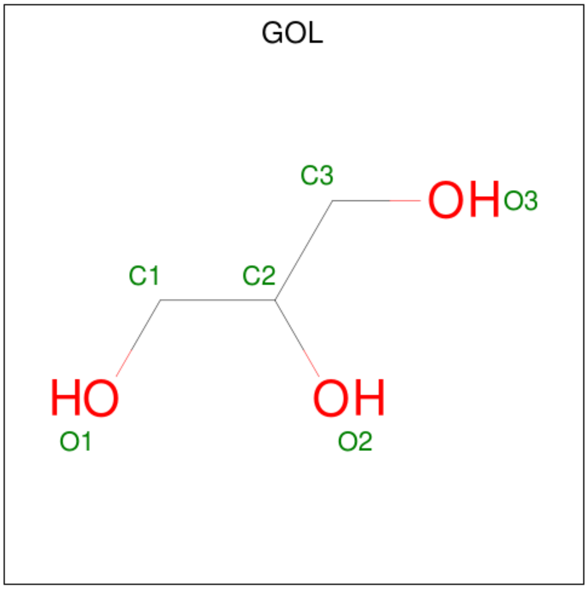


For Manuscript Review

| Mol | Chain | Residues | Atoms | ZeroOcc | AltConf |
|---|---|---|---|---|---|
| 4 | A | 1 | Total C O<br>6 3 3 | 0 | 0 |
| 4 | A | 1 | Total C O<br>6 3 3 | 0 | 0 |

- Molecule 5 is water.

| Mol | Chain | Residues | Atoms | ZeroOcc | AltConf |
|---|---|---|---|---|---|
| 5 | A | 51 | Total O<br>51 51 | 0 | 0 |

For Manuscript Review

### 3 Residue-property plots (i)

These plots are drawn for all protein, RNA, DNA and oligosaccharide chains in the entry. The first graphic for a chain summarises the proportions of the various outlier classes displayed in the second graphic. The second graphic shows the sequence view annotated by issues in geometry and electron density. Residues are color-coded according to the number of geometric quality criteria for which they contain at least one outlier: green = 0, yellow = 1, orange = 2 and red = 3 or more. A red dot above a residue indicates a poor fit to the electron density (RSRZ > 2). Stretches of 2 or more consecutive residues without any outlier are shown as a green connector. Residues present in the sample, but not in the model, are shown in grey.

• Molecule 1: Lysozyme C

Chain A:

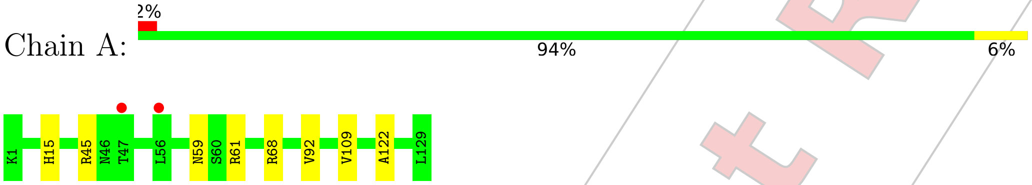


For Manuscript Review

### 4 Data and refinement statistics (i)

| Property | Value | Source |
|---|---|---|
| Space group | P 43 21 2 | Depositor |
| Cell constants<br>a, b, c, $\alpha$, $\beta$, $\gamma$ | 79.74Å 79.74Å 37.70Å<br>90.00° 90.00° 90.00° | Depositor |
| Resolution (Å) | 22.59 – 1.50<br>22.58 – 1.50 | Depositor<br>EDS |
| % Data completeness<br>(in resolution range) | 99.8 (22.59-1.50)<br>99.8 (22.58-1.50) | Depositor<br>EDS |
| $R_{merge}$ | (Not available) | Depositor |
| $R_{sym}$ | (Not available) | Depositor |
| $< I/\sigma(I) >$ [1] | 4.33 (at 1.50Å) | Xtriage |
| Refinement program | REFMAC 5.8.0267 | Depositor |
| R, $R_{free}$ | 0.171 , 0.190<br>0.180 , 0.202 | Depositor<br>DCC |
| $R_{free}$ test set | 1032 reflections (5.14%) | wwPDB-VP |
| Wilson B-factor (Å$^2$) | 12.6 | Xtriage |
| Anisotropy | 0.017 | Xtriage |
| Bulk solvent $k_{sol}$(e/Å$^3$), $B_{sol}$(Å$^2$) | 0.40 , 44.3 | EDS |
| L-test for twinning[2] | $< \|L\| > = 0.49$, $< L^2 > = 0.32$ | Xtriage |
| Estimated twinning fraction | No twinning to report. | Xtriage |
| $F_o$,$F_c$ correlation | 0.96 | EDS |
| Total number of atoms | 1072 | wwPDB-VP |
| Average B, all atoms (Å$^2$) | 18.0 | wwPDB-VP |

Xtriage's analysis on translational NCS is as follows: *The largest off-origin peak in the Patterson function is 10.99% of the height of the origin peak. No significant pseudotranslation is detected.*

[1] Intensities estimated from amplitudes.

[2] Theoretical values of $< |L| >$, $< L^2 >$ for acentric reflections are 0.5, 0.333 respectively for untwinned datasets, and 0.375, 0.2 for perfectly twinned datasets.

For Manuscript Review

### 5 Model quality (i)

#### 5.1 Standard geometry (i)

Bond lengths and bond angles in the following residue types are not validated in this section: GOL, CL, NA

The Z score for a bond length (or angle) is the number of standard deviations the observed value is removed from the expected value. A bond length (or angle) with $|Z| > 5$ is considered an outlier worth inspection. RMSZ is the root-mean-square of all Z scores of the bond lengths (or angles).

| Mol | Chain | Bond lengths | | Bond angles | |
|---|---|---|---|---|---|
| | | RMSZ | #\|Z\| >5 | RMSZ | #\|Z\| >5 |
| 1 | A | 0.76 | 0/1021 | 0.88 | 0/1379 |

There are no bond length outliers.

There are no bond angle outliers.

There are no chirality outliers.

There are no planarity outliers.

#### 5.2 Too-close contacts (i)

In the following table, the Non-H and H(model) columns list the number of non-hydrogen atoms and hydrogen atoms in the chain respectively. The H(added) column lists the number of hydrogen atoms added and optimized by MolProbity. The Clashes column lists the number of clashes within the asymmetric unit, whereas Symm-Clashes lists symmetry-related clashes.

| Mol | Chain | Non-H | H(model) | H(added) | Clashes | Symm-Clashes |
|---|---|---|---|---|---|---|
| 1 | A | 1001 | 0 | 958 | 11 | 0 |
| 2 | A | 1 | 0 | 0 | 0 | 0 |
| 3 | A | 7 | 0 | 0 | 0 | 0 |
| 4 | A | 12 | 0 | 16 | 1 | 0 |
| 5 | A | 51 | 0 | 0 | 4 | 0 |
| All | All | 1072 | 0 | 974 | 11 | 0 |

The all-atom clashscore is defined as the number of clashes found per 1000 atoms (including hydrogen atoms). The all-atom clashscore for this structure is 6.

All (11) close contacts within the same asymmetric unit are listed below, sorted by their clash magnitude.

For Manuscript Review

| Atom-1 | Atom-2 | Interatomic distance (Å) | Clash overlap (Å) |
|---|---|---|---|
| 1:A:59:ASN:ND2 | 1:A:61:ARG:HB3 | 1.99 | 0.77 |
| 1:A:59:ASN:ND2 | 1:A:61:ARG:H | 1.88 | 0.71 |
| 1:A:68:ARG:CD | 5:A:301:HOH:O | 2.45 | 0.65 |
| 1:A:59:ASN:HD22 | 1:A:61:ARG:HB3 | 1.65 | 0.62 |
| 1:A:59:ASN:ND2 | 1:A:61:ARG:CB | 2.70 | 0.54 |
| 1:A:45:ARG:NH1 | 5:A:301:HOH:O | 2.48 | 0.47 |
| 1:A:59:ASN:HD21 | 1:A:61:ARG:CB | 2.31 | 0.44 |
| 1:A:122:ALA:HB1 | 4:A:209:GOL:H32 | 1.98 | 0.44 |
| 1:A:68:ARG:HD3 | 5:A:301:HOH:O | 2.14 | 0.41 |
| 1:A:109:VAL:HB | 5:A:309:HOH:O | 2.20 | 0.41 |
| 1:A:15:HIS:HB3 | 1:A:92:VAL:HG11 | 2.04 | 0.40 |

There are no symmetry-related clashes.

#### 5.3 Torsion angles (i)

##### 5.3.1 Protein backbone (i)

In the following table, the Percentiles column shows the percent Ramachandran outliers of the chain as a percentile score with respect to all X-ray entries followed by that with respect to entries of similar resolution.

The Analysed column shows the number of residues for which the backbone conformation was analysed, and the total number of residues.

| Mol | Chain | Analysed | Favoured | Allowed | Outliers | Percentiles | |
|---|---|---|---|---|---|---|---|
| 1 | A | 127/129 (98%) | 125 (98%) | 2 (2%) | 0 | 100 | 100 |

There are no Ramachandran outliers to report.

##### 5.3.2 Protein sidechains (i)

In the following table, the Percentiles column shows the percent sidechain outliers of the chain as a percentile score with respect to all X-ray entries followed by that with respect to entries of similar resolution.

The Analysed column shows the number of residues for which the sidechain conformation was analysed, and the total number of residues.

| Mol | Chain | Analysed | Rotameric | Outliers | Percentiles | |
|---|---|---|---|---|---|---|
| 1 | A | 105/105 (100%) | 105 (100%) | 0 | 100 | 100 |

For Manuscript Review

There are no protein residues with a non-rotameric sidechain to report.

Sometimes sidechains can be flipped to improve hydrogen bonding and reduce clashes. All (4) such sidechains are listed below:

| Mol | Chain | Res | Type |
|---|---|---|---|
| 1 | A | 15 | HIS |
| 1 | A | 46 | ASN |
| 1 | A | 59 | ASN |
| 1 | A | 65 | ASN |

##### 5.3.3 RNA (i)

There are no RNA molecules in this entry.

#### 5.4 Non-standard residues in protein, DNA, RNA chains (i)

There are no non-standard protein/DNA/RNA residues in this entry.

#### 5.5 Carbohydrates (i)

There are no monosaccharides in this entry.

#### 5.6 Ligand geometry (i)

Of 10 ligands modelled in this entry, 8 are monoatomic - leaving 2 for Mogul analysis.

In the following table, the Counts columns list the number of bonds (or angles) for which Mogul statistics could be retrieved, the number of bonds (or angles) that are observed in the model and the number of bonds (or angles) that are defined in the Chemical Component Dictionary. The Link column lists molecule types, if any, to which the group is linked. The Z score for a bond length (or angle) is the number of standard deviations the observed value is removed from the expected value. A bond length (or angle) with $|Z| > 2$ is considered an outlier worth inspection. RMSZ is the root-mean-square of all Z scores of the bond lengths (or angles).

| Mol | Type | Chain | Res | Link | Bond lengths | | | Bond angles | | |
|---|---|---|---|---|---|---|---|---|---|---|
| | | | | | Counts | RMSZ | $\#\|Z\| > 2$ | Counts | RMSZ | $\#\|Z\| > 2$ |
| 4 | GOL | A | 210 | - | 5,5,5 | 0.10 | 0 | 5,5,5 | 0.33 | 0 |
| 4 | GOL | A | 209 | - | 5,5,5 | 0.12 | 0 | 5,5,5 | 0.30 | 0 |

In the following table, the Chirals column lists the number of chiral outliers, the number of chiral centers analysed, the number of these observed in the model and the number defined in the Chemical Component Dictionary. Similar counts are reported in the Torsion and Rings columns. '-' means no outliers of that kind were identified.

For Manuscript Review

| Mol | Type | Chain | Res | Link | Chirals | Torsions | Rings |
|---|---|---|---|---|---|---|---|
| 4 | GOL | A | 210 | - | - | 2/4/4/4 | - |
| 4 | GOL | A | 209 | - | - | 2/4/4/4 | - |

There are no bond length outliers.

There are no bond angle outliers.

There are no chirality outliers.

All (4) torsion outliers are listed below:

| Mol | Chain | Res | Type | Atoms |
|---|---|---|---|---|
| 4 | A | 209 | GOL | C1-C2-C3-O3 |
| 4 | A | 210 | GOL | C1-C2-C3-O3 |
| 4 | A | 210 | GOL | O2-C2-C3-O3 |
| 4 | A | 209 | GOL | O2-C2-C3-O3 |

There are no ring outliers.

1 monomer is involved in 1 short contact:

| Mol | Chain | Res | Type | Clashes | Symm-Clashes |
|---|---|---|---|---|---|
| 4 | A | 209 | GOL | 1 | 0 |

#### 5.7 Other polymers (i)

There are no such residues in this entry.

#### 5.8 Polymer linkage issues (i)

There are no chain breaks in this entry.

For Manuscript Review

### 6 Fit of model and data (i)

#### 6.1 Protein, DNA and RNA chains (i)

In the following table, the column labelled '#RSRZ> 2' contains the number (and percentage) of RSRZ outliers, followed by percent RSRZ outliers for the chain as percentile scores relative to all X-ray entries and entries of similar resolution. The OWAB column contains the minimum, median, $95^{th}$ percentile and maximum values of the occupancy-weighted average B-factor per residue. The column labelled 'Q< 0.9' lists the number of (and percentage) of residues with an average occupancy less than 0.9.

| Mol | Chain | Analysed | <RSRZ> | #RSRZ>2 | | | OWAB(Å$^2$) | Q<0.9 |
|---|---|---|---|---|---|---|---|---|
| 1 | A | 129/129 (100%) | -0.05 | 2 (1%) | 72 | 77 | 8, 15, 30, 49 | 0 |

All (2) RSRZ outliers are listed below:

| Mol | Chain | Res | Type | RSRZ |
|---|---|---|---|---|
| 1 | A | 56 | LEU | 2.2 |
| 1 | A | 47 | THR | 2.1 |

#### 6.2 Non-standard residues in protein, DNA, RNA chains (i)

There are no non-standard protein/DNA/RNA residues in this entry.

#### 6.3 Carbohydrates (i)

There are no monosaccharides in this entry.

#### 6.4 Ligands (i)

In the following table, the Atoms column lists the number of modelled atoms in the group and the number defined in the chemical component dictionary. The B-factors column lists the minimum, median, $95^{th}$ percentile and maximum values of B factors of atoms in the group. The column labelled 'Q< 0.9' lists the number of atoms with occupancy less than 0.9.

| Mol | Type | Chain | Res | Atoms | RSCC | RSR | B-factors(Å$^2$) | Q<0.9 |
|---|---|---|---|---|---|---|---|---|
| 4 | GOL | A | 209 | 6/6 | 0.68 | 0.32 | 60,61,64,64 | 0 |
| 4 | GOL | A | 210 | 6/6 | 0.69 | 0.29 | 59,61,61,67 | 0 |
| 3 | CL | A | 208 | 1/1 | 0.81 | 0.21 | 48,48,48,48 | 0 |
| 3 | CL | A | 206 | 1/1 | 0.86 | 0.29 | 53,53,53,53 | 0 |
| 3 | CL | A | 205 | 1/1 | 0.87 | 0.22 | 49,49,49,49 | 0 |

*Continued on next page...*

For Manuscript Review

*Continued from previous page...*

| Mol | Type | Chain | Res | Atoms | RSCC | RSR | B-factors($Å^2$) | Q<0.9 |
|---|---|---|---|---|---|---|---|---|
| 3 | CL | A | 207 | 1/1 | 0.96 | 0.15 | 38,38,38,38 | 0 |
| 3 | CL | A | 203 | 1/1 | 0.99 | 0.04 | 22,22,22,22 | 0 |
| 3 | CL | A | 204 | 1/1 | 0.99 | 0.08 | 24,24,24,24 | 0 |
| 2 | NA | A | 201 | 1/1 | 0.99 | 0.04 | 12,12,12,12 | 0 |
| 3 | CL | A | 202 | 1/1 | 0.99 | 0.04 | 17,17,17,17 | 0 |

#### 6.5 Other polymers (i)

There are no such residues in this entry.

For Manuscript Review

### Full wwPDB X-ray Structure Validation Report (i)

Jun 9, 2022 – 11:27 AM EDT

| | | |
|---|---|---|
| PDB ID | : | 8D8C |
| Title | : | Crystal structure of hen egg white lysozyme at 300 Kelvin |
| Deposited on | : | 2022-06-08 |
| Resolution | : | 1.50 Å(reported) |

**This wwPDB validation report is for manuscript review**

This is a Full wwPDB X-ray Structure Validation Report.

This report is produced by the wwPDB biocuration pipeline after annotation of the structure.

We welcome your comments at **
A user guide is available at
https://www.wwpdb.org/validation/2017/XrayValidationReportHelp
with specific help available everywhere you see the (i) symbol.

The following versions of software and data (see references (i)) were used in the production of this report:

| | | |
|---|---|---|
| MolProbity | : | 4.02b-467 |
| Mogul | : | 1.8.5 (274361), CSD as541be (2020) |
| Xtriage (Phenix) | : | 1.13 |
| EDS | : | 2.28 |
| Percentile statistics | : | 20191225.v01 (using entries in the PDB archive December 25th 2019) |
| Refmac | : | 5.8.0158 |
| CCP4 | : | 7.0.044 (Gargrove) |
| Ideal geometry (proteins) | : | Engh & Huber (2001) |
| Ideal geometry (DNA, RNA) | : | Parkinson et al. (1996) |
| Validation Pipeline (wwPDB-VP) | : | 2.28 |

For Manuscript Review

### 1 Overall quality at a glance (i)

The following experimental techniques were used to determine the structure:
*X-RAY DIFFRACTION*

The reported resolution of this entry is 1.50 Å.

Percentile scores (ranging between 0-100) for global validation metrics of the entry are shown in the following graphic. The table shows the number of entries on which the scores are based.

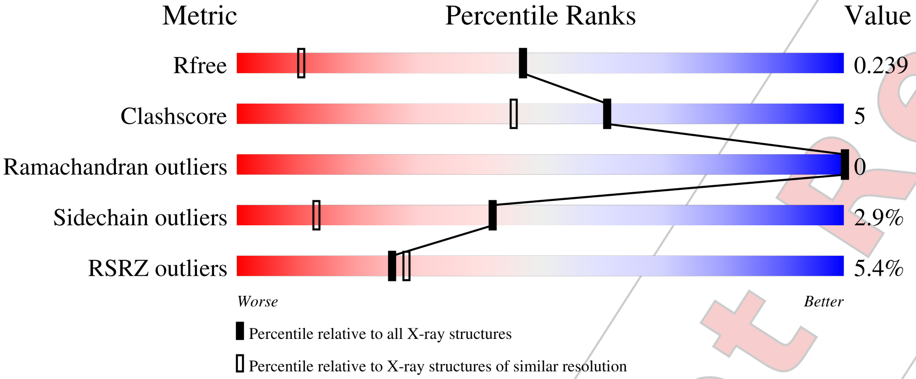


| Metric | Whole archive (#Entries) | Similar resolution (#Entries, resolution range(Å)) |
|---|---|---|
| $R_{free}$ | 130704 | 2936 (1.50-1.50) |
| Clashscore | 141614 | 3144 (1.50-1.50) |
| Ramachandran outliers | 138981 | 3066 (1.50-1.50) |
| Sidechain outliers | 138945 | 3064 (1.50-1.50) |
| RSRZ outliers | 127900 | 2884 (1.50-1.50) |

The table below summarises the geometric issues observed across the polymeric chains and their fit to the electron density. The red, orange, yellow and green segments of the lower bar indicate the fraction of residues that contain outliers for >=3, 2, 1 and 0 types of geometric quality criteria respectively. A grey segment represents the fraction of residues that are not modelled. The numeric value for each fraction is indicated below the corresponding segment, with a dot representing fractions <=5% The upper red bar (where present) indicates the fraction of residues that have poor fit to the electron density. The numeric value is given above the bar.

| Mol | Chain | Length | Quality of chain |
|---|---|---|---|
| 1 | A | 129 | 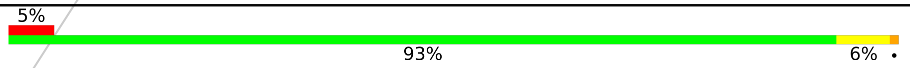 |

The following table lists non-polymeric compounds, carbohydrate monomers and non-standard residues in protein, DNA, RNA chains that are outliers for geometric or electron-density-fit criteria:

| Mol | Type | Chain | Res | Chirality | Geometry | Clashes | Electron density |
|---|---|---|---|---|---|---|---|
| 4 | GOL | A | 209 | - | - | - | X |

*Continued on next page...*

For Manuscript Review

*Continued from previous page...*

| Mol | Type | Chain | Res | Chirality | Geometry | Clashes | Electron density |
|---|---|---|---|---|---|---|---|
| 4 | GOL | A | 210 | - | - | - | X |

For Manuscript Review

### 2 Entry composition (i)

There are 5 unique types of molecules in this entry. The entry contains 1062 atoms, of which 0 are hydrogens and 0 are deuteriums.

In the tables below, the ZeroOcc column contains the number of atoms modelled with zero occupancy, the AltConf column contains the number of residues with at least one atom in alternate conformation and the Trace column contains the number of residues modelled with at most 2 atoms.

- Molecule 1 is a protein called Lysozyme C.

| Mol | Chain | Residues | Atoms | | | | | ZeroOcc | AltConf | Trace |
|---|---|---|---|---|---|---|---|---|---|---|
| | | | Total | C | N | O | S | | | |
| 1 | A | 129 | 1001 | 613 | 193 | 185 | 10 | 0 | 0 | 0 |

- Molecule 2 is SODIUM ION (three-letter code: NA) (formula: Na).

| Mol | Chain | Residues | Atoms | | ZeroOcc | AltConf |
|---|---|---|---|---|---|---|
| | | | Total | Na | | |
| 2 | A | 1 | 1 | 1 | 0 | 0 |

- Molecule 3 is CHLORIDE ION (three-letter code: CL) (formula: Cl).

| Mol | Chain | Residues | Atoms | | ZeroOcc | AltConf |
|---|---|---|---|---|---|---|
| | | | Total | Cl | | |
| 3 | A | 7 | 7 | 7 | 0 | 0 |

- Molecule 4 is GLYCEROL (three-letter code: GOL) (formula: $C_3H_8O_3$).

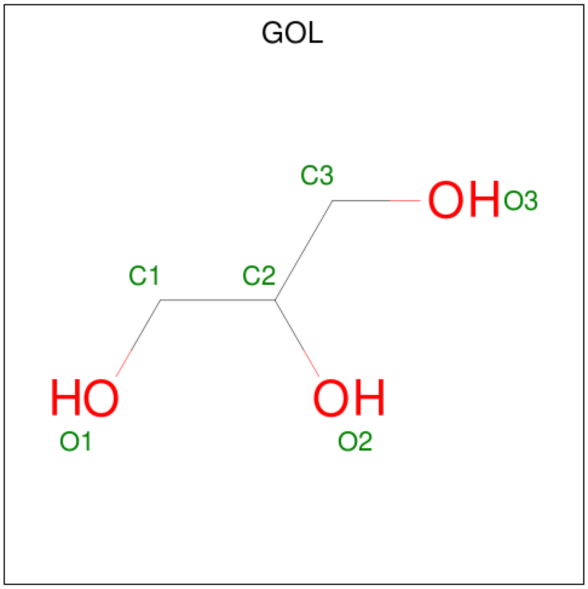


For Manuscript Review

| Mol | Chain | Residues | Atoms | ZeroOcc | AltConf |
|---|---|---|---|---|---|
| 4 | A | 1 | Total C O<br>6 3 3 | 0 | 0 |
| 4 | A | 1 | Total C O<br>6 3 3 | 0 | 0 |

- Molecule 5 is water.

| Mol | Chain | Residues | Atoms | ZeroOcc | AltConf |
|---|---|---|---|---|---|
| 5 | A | 41 | Total O<br>41 41 | 0 | 0 |

For Manuscript Review

### 3 Residue-property plots (i)

These plots are drawn for all protein, RNA, DNA and oligosaccharide chains in the entry. The first graphic for a chain summarises the proportions of the various outlier classes displayed in the second graphic. The second graphic shows the sequence view annotated by issues in geometry and electron density. Residues are color-coded according to the number of geometric quality criteria for which they contain at least one outlier: green = 0, yellow = 1, orange = 2 and red = 3 or more. A red dot above a residue indicates a poor fit to the electron density (RSRZ > 2). Stretches of 2 or more consecutive residues without any outlier are shown as a green connector. Residues present in the sample, but not in the model, are shown in grey.

- Molecule 1: Lysozyme C

Chain A:

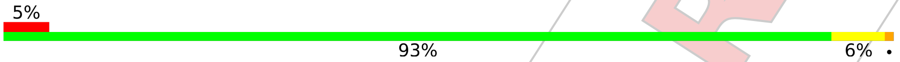


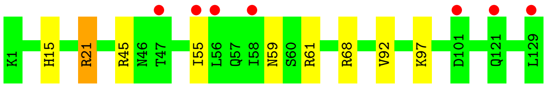


For Manuscript Review

### 4 Data and refinement statistics (i)

| Property | Value | Source |
|---|---|---|
| Space group | P 43 21 2 | Depositor |
| Cell constants<br>a, b, c, $\alpha$, $\beta$, $\gamma$ | 80.42Å 80.42Å 38.00Å<br>90.00° 90.00° 90.00° | Depositor |
| Resolution (Å) | 22.78 – 1.50<br>22.77 – 1.49 | Depositor<br>EDS |
| % Data completeness<br>(in resolution range) | 98.1 (22.78-1.50)<br>98.1 (22.77-1.49) | Depositor<br>EDS |
| $R_{merge}$ | (Not available) | Depositor |
| $R_{sym}$ | (Not available) | Depositor |
| $< I/\sigma(I) >$ [1] | 1.09 (at 1.49Å) | Xtriage |
| Refinement program | REFMAC 5.8.0267 | Depositor |
| R, $R_{free}$ | 0.199 , 0.228<br>0.209 , 0.239 | Depositor<br>DCC |
| $R_{free}$ test set | 1073 reflections (5.19%) | wwPDB-VP |
| Wilson B-factor (Å$^2$) | 18.1 | Xtriage |
| Anisotropy | 0.041 | Xtriage |
| Bulk solvent $k_{sol}$(e/Å$^3$), $B_{sol}$(Å$^2$) | 0.36 , 41.6 | EDS |
| L-test for twinning[2] | $< \|L\| > = 0.46$, $< L^2 > = 0.29$ | Xtriage |
| Estimated twinning fraction | No twinning to report. | Xtriage |
| $F_o$,$F_c$ correlation | 0.96 | EDS |
| Total number of atoms | 1062 | wwPDB-VP |
| Average B, all atoms (Å$^2$) | 23.0 | wwPDB-VP |

Xtriage's analysis on translational NCS is as follows: *The largest off-origin peak in the Patterson function is 11.47% of the height of the origin peak. No significant pseudotranslation is detected.*

[1] Intensities estimated from amplitudes.

[2] Theoretical values of $< |L| >$, $< L^2 >$ for acentric reflections are 0.5, 0.333 respectively for untwinned datasets, and 0.375, 0.2 for perfectly twinned datasets.

For Manuscript Review

### 5 Model quality (i)

#### 5.1 Standard geometry (i)

Bond lengths and bond angles in the following residue types are not validated in this section: CL, NA, GOL

The Z score for a bond length (or angle) is the number of standard deviations the observed value is removed from the expected value. A bond length (or angle) with $|Z| > 5$ is considered an outlier worth inspection. RMSZ is the root-mean-square of all Z scores of the bond lengths (or angles).

| Mol | Chain | Bond lengths | | Bond angles | |
|---|---|---|---|---|---|
| | | RMSZ | #$\|Z\|$ >5 | RMSZ | #$\|Z\|$ >5 |
| 1 | A | 0.84 | 0/1021 | 0.92 | 0/1379 |

There are no bond length outliers.

There are no bond angle outliers.

There are no chirality outliers.

There are no planarity outliers.

#### 5.2 Too-close contacts (i)

In the following table, the Non-H and H(model) columns list the number of non-hydrogen atoms and hydrogen atoms in the chain respectively. The H(added) column lists the number of hydrogen atoms added and optimized by MolProbity. The Clashes column lists the number of clashes within the asymmetric unit, whereas Symm-Clashes lists symmetry-related clashes.

| Mol | Chain | Non-H | H(model) | H(added) | Clashes | Symm-Clashes |
|---|---|---|---|---|---|---|
| 1 | A | 1001 | 0 | 958 | 9 | 0 |
| 2 | A | 1 | 0 | 0 | 0 | 0 |
| 3 | A | 7 | 0 | 0 | 0 | 0 |
| 4 | A | 12 | 0 | 16 | 0 | 0 |
| 5 | A | 41 | 0 | 0 | 0 | 0 |
| All | All | 1062 | 0 | 974 | 9 | 0 |

The all-atom clashscore is defined as the number of clashes found per 1000 atoms (including hydrogen atoms). The all-atom clashscore for this structure is 5.

All (9) close contacts within the same asymmetric unit are listed below, sorted by their clash magnitude.

For Manuscript Review

| Atom-1 | Atom-2 | Interatomic distance (Å) | Clash overlap (Å) |
|---|---|---|---|
| 1:A:21:ARG:HH21 | 1:A:21:ARG:HG3 | 1.59 | 0.68 |
| 1:A:59:ASN:ND2 | 1:A:61:ARG:H | 1.92 | 0.66 |
| 1:A:59:ASN:ND2 | 1:A:61:ARG:HB3 | 2.18 | 0.59 |
| 1:A:21:ARG:HH21 | 1:A:21:ARG:CG | 2.24 | 0.50 |
| 1:A:15:HIS:HB3 | 1:A:92:VAL:HG11 | 1.98 | 0.46 |
| 1:A:59:ASN:HD22 | 1:A:61:ARG:HB3 | 1.81 | 0.46 |
| 1:A:59:ASN:ND2 | 1:A:61:ARG:CB | 2.79 | 0.45 |
| 1:A:45:ARG:NE | 1:A:68:ARG:NH2 | 2.65 | 0.45 |
| 1:A:21:ARG:CG | 1:A:21:ARG:NH2 | 2.82 | 0.43 |

There are no symmetry-related clashes.

#### 5.3 Torsion angles (i)

##### 5.3.1 Protein backbone (i)

In the following table, the Percentiles column shows the percent Ramachandran outliers of the chain as a percentile score with respect to all X-ray entries followed by that with respect to entries of similar resolution.

The Analysed column shows the number of residues for which the backbone conformation was analysed, and the total number of residues.

| Mol | Chain | Analysed | Favoured | Allowed | Outliers | Percentiles | |
|---|---|---|---|---|---|---|---|
| 1 | A | 127/129 (98%) | 125 (98%) | 2 (2%) | 0 | 100 | 100 |

There are no Ramachandran outliers to report.

##### 5.3.2 Protein sidechains (i)

In the following table, the Percentiles column shows the percent sidechain outliers of the chain as a percentile score with respect to all X-ray entries followed by that with respect to entries of similar resolution.

The Analysed column shows the number of residues for which the sidechain conformation was analysed, and the total number of residues.

| Mol | Chain | Analysed | Rotameric | Outliers | Percentiles | |
|---|---|---|---|---|---|---|
| 1 | A | 105/105 (100%) | 102 (97%) | 3 (3%) | 42 | 13 |

All (3) residues with a non-rotameric sidechain are listed below:

For Manuscript Review

| Mol | Chain | Res | Type |
|---|---|---|---|
| 1 | A | 21 | ARG |
| 1 | A | 55 | ILE |
| 1 | A | 97 | LYS |

Sometimes sidechains can be flipped to improve hydrogen bonding and reduce clashes. All (3) such sidechains are listed below:

| Mol | Chain | Res | Type |
|---|---|---|---|
| 1 | A | 15 | HIS |
| 1 | A | 46 | ASN |
| 1 | A | 59 | ASN |

##### 5.3.3 RNA (i)

There are no RNA molecules in this entry.

#### 5.4 Non-standard residues in protein, DNA, RNA chains (i)

There are no non-standard protein/DNA/RNA residues in this entry.

#### 5.5 Carbohydrates (i)

There are no monosaccharides in this entry.

#### 5.6 Ligand geometry (i)

Of 10 ligands modelled in this entry, 8 are monoatomic - leaving 2 for Mogul analysis.

In the following table, the Counts columns list the number of bonds (or angles) for which Mogul statistics could be retrieved, the number of bonds (or angles) that are observed in the model and the number of bonds (or angles) that are defined in the Chemical Component Dictionary. The Link column lists molecule types, if any, to which the group is linked. The Z score for a bond length (or angle) is the number of standard deviations the observed value is removed from the expected value. A bond length (or angle) with $|Z| > 2$ is considered an outlier worth inspection. RMSZ is the root-mean-square of all Z scores of the bond lengths (or angles).

| Mol | Type | Chain | Res | Link | Bond lengths | | | Bond angles | | |
|---|---|---|---|---|---|---|---|---|---|---|
| | | | | | Counts | RMSZ | $\#\|Z\| > 2$ | Counts | RMSZ | $\#\|Z\| > 2$ |
| 4 | GOL | A | 209 | - | 5,5,5 | 0.10 | 0 | 5,5,5 | 0.31 | 0 |
| 4 | GOL | A | 210 | - | 5,5,5 | 0.08 | 0 | 5,5,5 | 0.29 | 0 |

For Manuscript Review

In the following table, the Chirals column lists the number of chiral outliers, the number of chiral centers analysed, the number of these observed in the model and the number defined in the Chemical Component Dictionary. Similar counts are reported in the Torsion and Rings columns. '-' means no outliers of that kind were identified.

| Mol | Type | Chain | Res | Link | Chirals | Torsions | Rings |
|---|---|---|---|---|---|---|---|
| 4 | GOL | A | 209 | - | - | 2/4/4/4 | - |
| 4 | GOL | A | 210 | - | - | 4/4/4/4 | - |

There are no bond length outliers.

There are no bond angle outliers.

There are no chirality outliers.

All (6) torsion outliers are listed below:

| Mol | Chain | Res | Type | Atoms |
|---|---|---|---|---|
| 4 | A | 209 | GOL | C1-C2-C3-O3 |
| 4 | A | 210 | GOL | O1-C1-C2-C3 |
| 4 | A | 209 | GOL | O2-C2-C3-O3 |
| 4 | A | 210 | GOL | O1-C1-C2-O2 |
| 4 | A | 210 | GOL | O2-C2-C3-O3 |
| 4 | A | 210 | GOL | C1-C2-C3-O3 |

There are no ring outliers.

No monomer is involved in short contacts.

#### 5.7 Other polymers (i)

There are no such residues in this entry.

#### 5.8 Polymer linkage issues (i)

There are no chain breaks in this entry.

For Manuscript Review

### 6 Fit of model and data (i)

#### 6.1 Protein, DNA and RNA chains (i)

In the following table, the column labelled '#RSRZ> 2' contains the number (and percentage) of RSRZ outliers, followed by percent RSRZ outliers for the chain as percentile scores relative to all X-ray entries and entries of similar resolution. The OWAB column contains the minimum, median, $95^{th}$ percentile and maximum values of the occupancy-weighted average B-factor per residue. The column labelled 'Q< 0.9' lists the number of (and percentage) of residues with an average occupancy less than 0.9.

| Mol | Chain | Analysed | <RSRZ> | #RSRZ>2 | | | OWAB($Å^2$) | Q<0.9 |
|---|---|---|---|---|---|---|---|---|
| 1 | A | 129/129 (100%) | 0.51 | 7 (5%) | 25 | 28 | 13, 20, 38, 46 | 0 |

All (7) RSRZ outliers are listed below:

| Mol | Chain | Res | Type | RSRZ |
|---|---|---|---|---|
| 1 | A | 47 | THR | 3.5 |
| 1 | A | 55 | ILE | 3.2 |
| 1 | A | 101 | ASP | 2.9 |
| 1 | A | 56 | LEU | 2.6 |
| 1 | A | 58 | ILE | 2.2 |
| 1 | A | 129 | LEU | 2.1 |
| 1 | A | 121 | GLN | 2.1 |

#### 6.2 Non-standard residues in protein, DNA, RNA chains (i)

There are no non-standard protein/DNA/RNA residues in this entry.

#### 6.3 Carbohydrates (i)

There are no monosaccharides in this entry.

#### 6.4 Ligands (i)

In the following table, the Atoms column lists the number of modelled atoms in the group and the number defined in the chemical component dictionary. The B-factors column lists the minimum, median, $95^{th}$ percentile and maximum values of B factors of atoms in the group. The column labelled 'Q< 0.9' lists the number of atoms with occupancy less than 0.9.

| Mol | Type | Chain | Res | Atoms | RSCC | RSR | B-factors($Å^2$) | Q<0.9 |
|---|---|---|---|---|---|---|---|---|
| 4 | GOL | A | 210 | 6/6 | 0.35 | 0.43 | 71,75,76,79 | 0 |

*Continued on next page...*

For Manuscript Review

*Continued from previous page...*

| Mol | Type | Chain | Res | Atoms | RSCC | RSR | B-factors($Å^2$) | Q<0.9 |
|---|---|---|---|---|---|---|---|---|
| 4 | GOL | A | 209 | 6/6 | 0.48 | 0.48 | 60,64,68,71 | 0 |
| 3 | CL | A | 206 | 1/1 | 0.71 | 0.26 | 57,57,57,57 | 0 |
| 3 | CL | A | 208 | 1/1 | 0.86 | 0.33 | 61,61,61,61 | 0 |
| 3 | CL | A | 207 | 1/1 | 0.87 | 0.14 | 55,55,55,55 | 0 |
| 3 | CL | A | 205 | 1/1 | 0.88 | 0.26 | 62,62,62,62 | 0 |
| 3 | CL | A | 203 | 1/1 | 0.97 | 0.04 | 34,34,34,34 | 0 |
| 3 | CL | A | 202 | 1/1 | 0.98 | 0.06 | 24,24,24,24 | 0 |
| 3 | CL | A | 204 | 1/1 | 0.98 | 0.07 | 32,32,32,32 | 0 |
| 2 | NA | A | 201 | 1/1 | 0.99 | 0.09 | 21,21,21,21 | 0 |

#### 6.5 Other polymers (i)

There are no such residues in this entry.

For Manuscript Review

### Full wwPDB X-ray Structure Validation Report (i)

Jun 9, 2022 – 11:17 AM EDT

| | | |
|---|---|---|
| PDB ID | : | 8D8D |
| Title | : | Crystal structure of hen egg white lysozyme at 300 Kelvin (Duplicate) |
| Deposited on | : | 2022-06-08 |
| Resolution | : | 1.50 Å(reported) |

**This wwPDB validation report is for manuscript review**

This is a Full wwPDB X-ray Structure Validation Report.

This report is produced by the wwPDB biocuration pipeline after annotation of the structure.

We welcome your comments at **
A user guide is available at
https://www.wwpdb.org/validation/2017/XrayValidationReportHelp
with specific help available everywhere you see the (i) symbol.

The following versions of software and data (see references (i)) were used in the production of this report:

| | | |
|---|---|---|
| MolProbity | : | 4.02b-467 |
| Mogul | : | 1.8.5 (274361), CSD as541be (2020) |
| Xtriage (Phenix) | : | 1.13 |
| EDS | : | 2.28 |
| Percentile statistics | : | 20191225.v01 (using entries in the PDB archive December 25th 2019) |
| Refmac | : | 5.8.0158 |
| CCP4 | : | 7.0.044 (Gargrove) |
| Ideal geometry (proteins) | : | Engh & Huber (2001) |
| Ideal geometry (DNA, RNA) | : | Parkinson et al. (1996) |
| Validation Pipeline (wwPDB-VP) | : | 2.28 |

For Manuscript Review

### 1 Overall quality at a glance (i)

The following experimental techniques were used to determine the structure:
*X-RAY DIFFRACTION*

The reported resolution of this entry is 1.50 Å.

Percentile scores (ranging between 0-100) for global validation metrics of the entry are shown in the following graphic. The table shows the number of entries on which the scores are based.

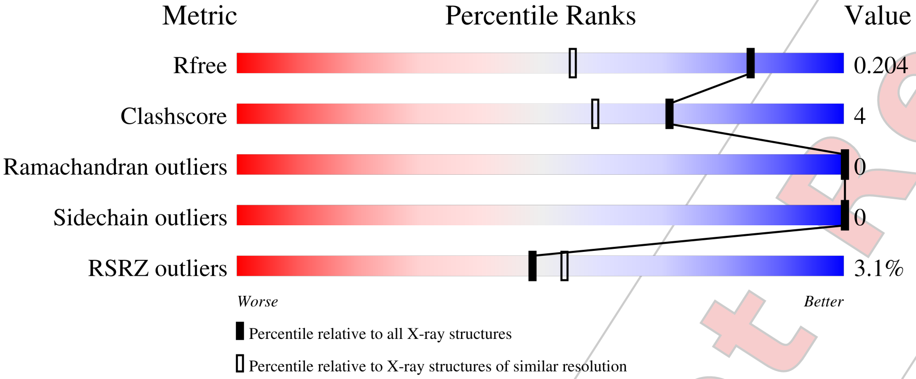


| Metric | Whole archive (#Entries) | Similar resolution (#Entries, resolution range(Å)) |
|---|---|---|
| $R_{free}$ | 130704 | 2936 (1.50-1.50) |
| Clashscore | 141614 | 3144 (1.50-1.50) |
| Ramachandran outliers | 138981 | 3066 (1.50-1.50) |
| Sidechain outliers | 138945 | 3064 (1.50-1.50) |
| RSRZ outliers | 127900 | 2884 (1.50-1.50) |

The table below summarises the geometric issues observed across the polymeric chains and their fit to the electron density. The red, orange, yellow and green segments of the lower bar indicate the fraction of residues that contain outliers for >=3, 2, 1 and 0 types of geometric quality criteria respectively. A grey segment represents the fraction of residues that are not modelled. The numeric value for each fraction is indicated below the corresponding segment, with a dot representing fractions <=5% The upper red bar (where present) indicates the fraction of residues that have poor fit to the electron density. The numeric value is given above the bar.

| Mol | Chain | Length | Quality of chain |
|---|---|---|---|
| 1 | A | 129 | 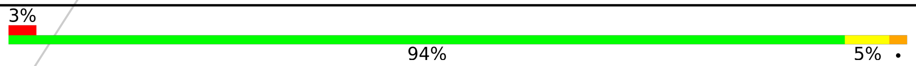  |

The following table lists non-polymeric compounds, carbohydrate monomers and non-standard residues in protein, DNA, RNA chains that are outliers for geometric or electron-density-fit criteria:

| Mol | Type | Chain | Res | Chirality | Geometry | Clashes | Electron density |
|---|---|---|---|---|---|---|---|
| 4 | GOL | A | 209 | - | - | - | X |

For Manuscript Review

### 2 Entry composition (i)

There are 5 unique types of molecules in this entry. The entry contains 1067 atoms, of which 0 are hydrogens and 0 are deuteriums.

In the tables below, the ZeroOcc column contains the number of atoms modelled with zero occupancy, the AltConf column contains the number of residues with at least one atom in alternate conformation and the Trace column contains the number of residues modelled with at most 2 atoms.

- Molecule 1 is a protein called Lysozyme C.

| Mol | Chain | Residues | Atoms | | | | | ZeroOcc | AltConf | Trace |
|---|---|---|---|---|---|---|---|---|---|---|
| | | | Total | C | N | O | S | | | |
| 1 | A | 129 | 1001 | 613 | 193 | 185 | 10 | 0 | 0 | 0 |

- Molecule 2 is SODIUM ION (three-letter code: NA) (formula: Na).

| Mol | Chain | Residues | Atoms | | ZeroOcc | AltConf |
|---|---|---|---|---|---|---|
| | | | Total | Na | | |
| 2 | A | 1 | 1 | 1 | 0 | 0 |

- Molecule 3 is CHLORIDE ION (three-letter code: CL) (formula: Cl).

| Mol | Chain | Residues | Atoms | | ZeroOcc | AltConf |
|---|---|---|---|---|---|---|
| | | | Total | Cl | | |
| 3 | A | 7 | 7 | 7 | 0 | 0 |

- Molecule 4 is GLYCEROL (three-letter code: GOL) (formula: $C_3H_8O_3$).

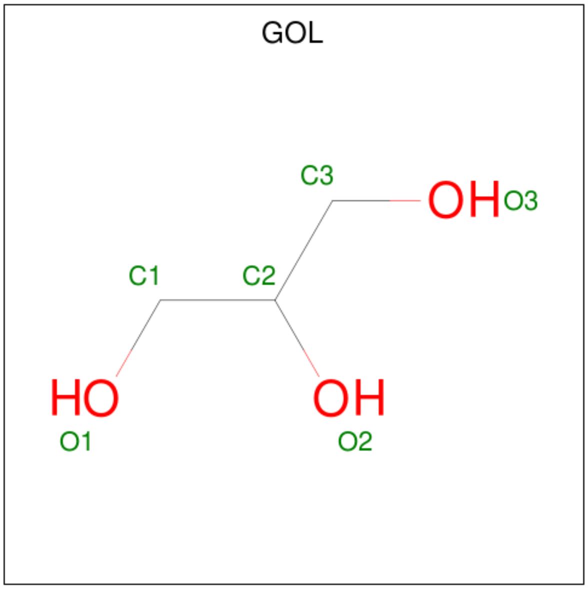


For Manuscript Review

| Mol | Chain | Residues | Atoms | ZeroOcc | AltConf |
|---|---|---|---|---|---|
| 4 | A | 1 | Total C O<br>6 3 3 | 0 | 0 |
| 4 | A | 1 | Total C O<br>6 3 3 | 0 | 0 |

- Molecule 5 is water.

| Mol | Chain | Residues | Atoms | ZeroOcc | AltConf |
|---|---|---|---|---|---|
| 5 | A | 46 | Total O<br>46 46 | 0 | 0 |

For Manuscript Review

### 3 Residue-property plots (i)

These plots are drawn for all protein, RNA, DNA and oligosaccharide chains in the entry. The first graphic for a chain summarises the proportions of the various outlier classes displayed in the second graphic. The second graphic shows the sequence view annotated by issues in geometry and electron density. Residues are color-coded according to the number of geometric quality criteria for which they contain at least one outlier: green = 0, yellow = 1, orange = 2 and red = 3 or more. A red dot above a residue indicates a poor fit to the electron density (RSRZ > 2). Stretches of 2 or more consecutive residues without any outlier are shown as a green connector. Residues present in the sample, but not in the model, are shown in grey.

• Molecule 1: Lysozyme C

Chain A:
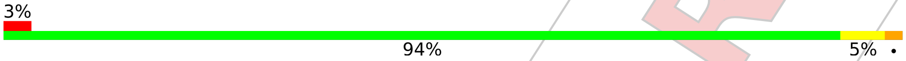


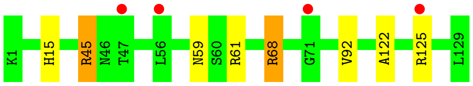


For Manuscript Review

### 4 Data and refinement statistics (i)

| Property | Value | Source |
|---|---|---|
| Space group | P 43 21 2 | Depositor |
| Cell constants<br>a, b, c, $\alpha$, $\beta$, $\gamma$ | 79.48Å 79.48Å 37.86Å<br>90.00° 90.00° 90.00° | Depositor |
| Resolution (Å) | 22.57 – 1.50<br>22.57 – 1.50 | Depositor<br>EDS |
| % Data completeness<br>(in resolution range) | 99.5 (22.57-1.50)<br>99.6 (22.57-1.50) | Depositor<br>EDS |
| $R_{merge}$ | (Not available) | Depositor |
| $R_{sym}$ | (Not available) | Depositor |
| $< I/\sigma(I) >$ [1] | 2.47 (at 1.50Å) | Xtriage |
| Refinement program | REFMAC 5.8.0267 | Depositor |
| R, $R_{free}$ | 0.176 , 0.203<br>0.177 , 0.204 | Depositor<br>DCC |
| $R_{free}$ test set | 1026 reflections (5.15%) | wwPDB-VP |
| Wilson B-factor (Å$^2$) | 16.6 | Xtriage |
| Anisotropy | 0.073 | Xtriage |
| Bulk solvent $k_{sol}$(e/Å$^3$), $B_{sol}$(Å$^2$) | 0.36 , 41.1 | EDS |
| L-test for twinning[2] | $< \|L\| > = 0.52$, $< L^2 > = 0.36$ | Xtriage |
| Estimated twinning fraction | No twinning to report. | Xtriage |
| $F_o$,$F_c$ correlation | 0.97 | EDS |
| Total number of atoms | 1067 | wwPDB-VP |
| Average B, all atoms (Å$^2$) | 21.0 | wwPDB-VP |

Xtriage's analysis on translational NCS is as follows: *The largest off-origin peak in the Patterson function is 10.76% of the height of the origin peak. No significant pseudotranslation is detected.*

[1] Intensities estimated from amplitudes.

[2] Theoretical values of $< |L| >$, $< L^2 >$ for acentric reflections are 0.5, 0.333 respectively for untwinned datasets, and 0.375, 0.2 for perfectly twinned datasets.

For Manuscript Review

### 5 Model quality (i)

#### 5.1 Standard geometry (i)

Bond lengths and bond angles in the following residue types are not validated in this section: NA, CL, GOL

The Z score for a bond length (or angle) is the number of standard deviations the observed value is removed from the expected value. A bond length (or angle) with $|Z| > 5$ is considered an outlier worth inspection. RMSZ is the root-mean-square of all Z scores of the bond lengths (or angles).

| Mol | Chain | Bond lengths | | Bond angles | |
|---|---|---|---|---|---|
| | | RMSZ | #$\|Z\|$ >5 | RMSZ | #$\|Z\|$ >5 |
| 1 | A | 0.76 | 0/1021 | 0.94 | 2/1379 (0.1%) |

There are no bond length outliers.

All (2) bond angle outliers are listed below:

| Mol | Chain | Res | Type | Atoms | Z | Observed($^o$) | Ideal($^o$) |
|---|---|---|---|---|---|---|---|
| 1 | A | 68 | ARG | NE-CZ-NH1 | -7.01 | 116.79 | 120.30 |
| 1 | A | 45 | ARG | CG-CD-NE | -5.04 | 101.22 | 111.80 |

There are no chirality outliers.

There are no planarity outliers.

#### 5.2 Too-close contacts (i)

In the following table, the Non-H and H(model) columns list the number of non-hydrogen atoms and hydrogen atoms in the chain respectively. The H(added) column lists the number of hydrogen atoms added and optimized by MolProbity. The Clashes column lists the number of clashes within the asymmetric unit, whereas Symm-Clashes lists symmetry-related clashes.

| Mol | Chain | Non-H | H(model) | H(added) | Clashes | Symm-Clashes |
|---|---|---|---|---|---|---|
| 1 | A | 1001 | 0 | 958 | 7 | 0 |
| 2 | A | 1 | 0 | 0 | 0 | 0 |
| 3 | A | 7 | 0 | 0 | 0 | 0 |
| 4 | A | 12 | 0 | 16 | 0 | 0 |
| 5 | A | 46 | 0 | 0 | 0 | 0 |
| All | All | 1067 | 0 | 974 | 7 | 0 |

The all-atom clashscore is defined as the number of clashes found per 1000 atoms (including hydrogen atoms). The all-atom clashscore for this structure is 4.

For Manuscript Review

All (7) close contacts within the same asymmetric unit are listed below, sorted by their clash magnitude.

| Atom-1 | Atom-2 | Interatomic distance (Å) | Clash overlap (Å) |
|---|---|---|---|
| 1:A:59:ASN:ND2 | 1:A:61:ARG:HB3 | 2.08 | 0.69 |
| 1:A:59:ASN:ND2 | 1:A:61:ARG:H | 1.95 | 0.63 |
| 1:A:45:ARG:NE | 1:A:68:ARG:NH2 | 2.62 | 0.48 |
| 1:A:15:HIS:HB3 | 1:A:92:VAL:HG11 | 1.96 | 0.47 |
| 1:A:122:ALA:HA | 1:A:125:ARG:HG3 | 2.01 | 0.43 |
| 1:A:59:ASN:HD22 | 1:A:61:ARG:HB3 | 1.81 | 0.42 |
| 1:A:59:ASN:ND2 | 1:A:61:ARG:CB | 2.81 | 0.41 |

There are no symmetry-related clashes.

#### 5.3 Torsion angles (i)

##### 5.3.1 Protein backbone (i)

In the following table, the Percentiles column shows the percent Ramachandran outliers of the chain as a percentile score with respect to all X-ray entries followed by that with respect to entries of similar resolution.

The Analysed column shows the number of residues for which the backbone conformation was analysed, and the total number of residues.

| Mol | Chain | Analysed | Favoured | Allowed | Outliers | Percentiles | |
|---|---|---|---|---|---|---|---|
| 1 | A | 127/129 (98%) | 126 (99%) | 1 (1%) | 0 | 100 | 100 |

There are no Ramachandran outliers to report.

##### 5.3.2 Protein sidechains (i)

In the following table, the Percentiles column shows the percent sidechain outliers of the chain as a percentile score with respect to all X-ray entries followed by that with respect to entries of similar resolution.

The Analysed column shows the number of residues for which the sidechain conformation was analysed, and the total number of residues.

| Mol | Chain | Analysed | Rotameric | Outliers | Percentiles | |
|---|---|---|---|---|---|---|
| 1 | A | 105/105 (100%) | 105 (100%) | 0 | 100 | 100 |

There are no protein residues with a non-rotameric sidechain to report.

For Manuscript Review

Sometimes sidechains can be flipped to improve hydrogen bonding and reduce clashes. All (3) such sidechains are listed below:

| Mol | Chain | Res | Type |
|---|---|---|---|
| 1 | A | 15 | HIS |
| 1 | A | 46 | ASN |
| 1 | A | 59 | ASN |

##### 5.3.3 RNA (i)

There are no RNA molecules in this entry.

#### 5.4 Non-standard residues in protein, DNA, RNA chains (i)

There are no non-standard protein/DNA/RNA residues in this entry.

#### 5.5 Carbohydrates (i)

There are no monosaccharides in this entry.

#### 5.6 Ligand geometry (i)

Of 10 ligands modelled in this entry, 8 are monoatomic - leaving 2 for Mogul analysis.

In the following table, the Counts columns list the number of bonds (or angles) for which Mogul statistics could be retrieved, the number of bonds (or angles) that are observed in the model and the number of bonds (or angles) that are defined in the Chemical Component Dictionary. The Link column lists molecule types, if any, to which the group is linked. The Z score for a bond length (or angle) is the number of standard deviations the observed value is removed from the expected value. A bond length (or angle) with $|Z| > 2$ is considered an outlier worth inspection. RMSZ is the root-mean-square of all Z scores of the bond lengths (or angles).

| Mol | Type | Chain | Res | Link | Bond lengths | | | Bond angles | | |
|---|---|---|---|---|---|---|---|---|---|---|
| | | | | | Counts | RMSZ | $\#\|Z\| > 2$ | Counts | RMSZ | $\#\|Z\| > 2$ |
| 4 | GOL | A | 209 | - | 5,5,5 | 0.12 | 0 | 5,5,5 | 0.29 | 0 |
| 4 | GOL | A | 210 | - | 5,5,5 | 0.08 | 0 | 5,5,5 | 0.31 | 0 |

In the following table, the Chirals column lists the number of chiral outliers, the number of chiral centers analysed, the number of these observed in the model and the number defined in the Chemical Component Dictionary. Similar counts are reported in the Torsion and Rings columns. '-' means no outliers of that kind were identified.

For Manuscript Review

| Mol | Type | Chain | Res | Link | Chirals | Torsions | Rings |
|---|---|---|---|---|---|---|---|
| 4 | GOL | A | 209 | - | - | 2/4/4/4 | - |
| 4 | GOL | A | 210 | - | - | 2/4/4/4 | - |

There are no bond length outliers.

There are no bond angle outliers.

There are no chirality outliers.

All (4) torsion outliers are listed below:

| Mol | Chain | Res | Type | Atoms |
|---|---|---|---|---|
| 4 | A | 210 | GOL | O1-C1-C2-C3 |
| 4 | A | 209 | GOL | O2-C2-C3-O3 |
| 4 | A | 210 | GOL | O1-C1-C2-O2 |
| 4 | A | 209 | GOL | C1-C2-C3-O3 |

There are no ring outliers.

No monomer is involved in short contacts.

#### 5.7 Other polymers (i)

There are no such residues in this entry.

#### 5.8 Polymer linkage issues (i)

There are no chain breaks in this entry.

For Manuscript Review

For Manuscript Review

### 6 Fit of model and data (i)

#### 6.1 Protein, DNA and RNA chains (i)

In the following table, the column labelled '#RSRZ> 2' contains the number (and percentage) of RSRZ outliers, followed by percent RSRZ outliers for the chain as percentile scores relative to all X-ray entries and entries of similar resolution. The OWAB column contains the minimum, median, $95^{th}$ percentile and maximum values of the occupancy-weighted average B-factor per residue. The column labelled 'Q< 0.9' lists the number of (and percentage) of residues with an average occupancy less than 0.9.

| Mol | Chain | Analysed | <RSRZ> | #RSRZ>2 | | | OWAB(Å$^2$) | Q<0.9 |
|---|---|---|---|---|---|---|---|---|
| 1 | A | 129/129 (100%) | 0.07 | 4 (3%) | 49 | 54 | 11, 18, 33, 44 | 0 |

All (4) RSRZ outliers are listed below:

| Mol | Chain | Res | Type | RSRZ |
|---|---|---|---|---|
| 1 | A | 71 | GLY | 2.9 |
| 1 | A | 56 | LEU | 2.3 |
| 1 | A | 47 | THR | 2.2 |
| 1 | A | 125 | ARG | 2.1 |

#### 6.2 Non-standard residues in protein, DNA, RNA chains (i)

There are no non-standard protein/DNA/RNA residues in this entry.

#### 6.3 Carbohydrates (i)

There are no monosaccharides in this entry.

#### 6.4 Ligands (i)

In the following table, the Atoms column lists the number of modelled atoms in the group and the number defined in the chemical component dictionary. The B-factors column lists the minimum, median, $95^{th}$ percentile and maximum values of B factors of atoms in the group. The column labelled 'Q< 0.9' lists the number of atoms with occupancy less than 0.9.

| Mol | Type | Chain | Res | Atoms | RSCC | RSR | B-factors(Å$^2$) | Q<0.9 |
|---|---|---|---|---|---|---|---|---|
| 4 | GOL | A | 209 | 6/6 | 0.45 | 0.48 | 70,79,80,82 | 0 |
| 4 | GOL | A | 210 | 6/6 | 0.63 | 0.36 | 84,86,87,88 | 0 |
| 3 | CL | A | 206 | 1/1 | 0.68 | 0.33 | 65,65,65,65 | 0 |

*Continued on next page...*

*Continued from previous page...*

| Mol | Type | Chain | Res | Atoms | RSCC | RSR | B-factors(Å$^2$) | Q<0.9 |
|---|---|---|---|---|---|---|---|---|
| 3 | CL | A | 205 | 1/1 | 0.78 | 0.24 | 61,61,61,61 | 0 |
| 3 | CL | A | 208 | 1/1 | 0.83 | 0.29 | 62,62,62,62 | 0 |
| 3 | CL | A | 207 | 1/1 | 0.91 | 0.18 | 50,50,50,50 | 0 |
| 3 | CL | A | 202 | 1/1 | 0.98 | 0.04 | 22,22,22,22 | 0 |
| 3 | CL | A | 204 | 1/1 | 0.98 | 0.09 | 31,31,31,31 | 0 |
| 3 | CL | A | 203 | 1/1 | 0.99 | 0.12 | 32,32,32,32 | 0 |
| 2 | NA | A | 201 | 1/1 | 0.99 | 0.07 | 19,19,19,19 | 0 |

#### 6.5 Other polymers (i)

There are no such residues in this entry.

For Manuscript Review

### Full wwPDB X-ray Structure Validation Report (i)

Jun 9, 2022 – 11:19 AM EDT

| | | |
|---|---|---|
| PDB ID | : | 8D8E |
| Title | : | Crystal structure of hen egg white lysozyme at 300 Kelvin (Triplicate) |
| Deposited on | : | 2022-06-08 |
| Resolution | : | 1.50 Å(reported) |

**This wwPDB validation report is for manuscript review**

This is a Full wwPDB X-ray Structure Validation Report.

This report is produced by the wwPDB biocuration pipeline after annotation of the structure.

We welcome your comments at **
A user guide is available at
https://www.wwpdb.org/validation/2017/XrayValidationReportHelp
with specific help available everywhere you see the (i) symbol.

The following versions of software and data (see references (i)) were used in the production of this report:

| | | |
|---|---|---|
| MolProbity | : | 4.02b-467 |
| Mogul | : | 1.8.5 (274361), CSD as541be (2020) |
| Xtriage (Phenix) | : | 1.13 |
| EDS | : | 2.28 |
| Percentile statistics | : | 20191225.v01 (using entries in the PDB archive December 25th 2019) |
| Refmac | : | 5.8.0158 |
| CCP4 | : | 7.0.044 (Gargrove) |
| Ideal geometry (proteins) | : | Engh & Huber (2001) |
| Ideal geometry (DNA, RNA) | : | Parkinson et al. (1996) |
| Validation Pipeline (wwPDB-VP) | : | 2.28 |

For Manuscript Review

### 1 Overall quality at a glance (i)

The following experimental techniques were used to determine the structure:
*X-RAY DIFFRACTION*

The reported resolution of this entry is 1.50 Å.

Percentile scores (ranging between 0-100) for global validation metrics of the entry are shown in the following graphic. The table shows the number of entries on which the scores are based.

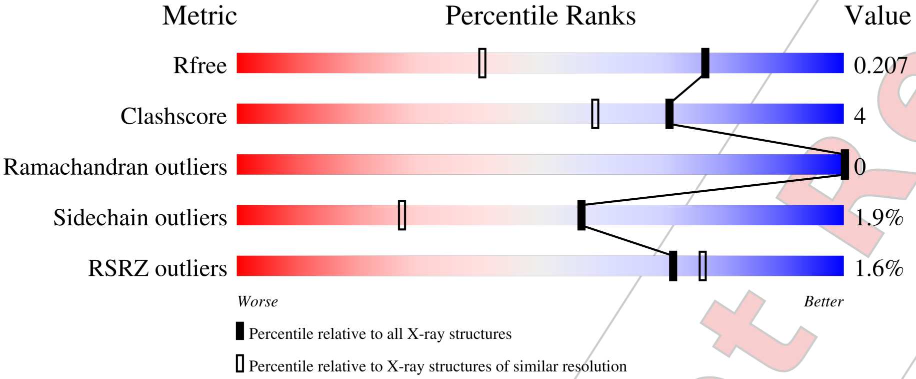


| Metric | Whole archive (#Entries) | Similar resolution (#Entries, resolution range(Å)) |
|---|---|---|
| $R_{free}$ | 130704 | 2936 (1.50-1.50) |
| Clashscore | 141614 | 3144 (1.50-1.50) |
| Ramachandran outliers | 138981 | 3066 (1.50-1.50) |
| Sidechain outliers | 138945 | 3064 (1.50-1.50) |
| RSRZ outliers | 127900 | 2884 (1.50-1.50) |

The table below summarises the geometric issues observed across the polymeric chains and their fit to the electron density. The red, orange, yellow and green segments of the lower bar indicate the fraction of residues that contain outliers for >=3, 2, 1 and 0 types of geometric quality criteria respectively. A grey segment represents the fraction of residues that are not modelled. The numeric value for each fraction is indicated below the corresponding segment, with a dot representing fractions <=5% The upper red bar (where present) indicates the fraction of residues that have poor fit to the electron density. The numeric value is given above the bar.

| Mol | Chain | Length | Quality of chain |
|---|---|---|---|
| 1 | A | 129 | 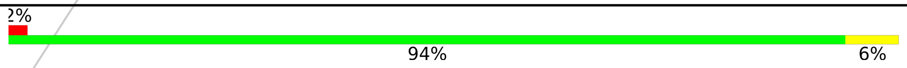 |



The following table lists non-polymeric compounds, carbohydrate monomers and non-standard residues in protein, DNA, RNA chains that are outliers for geometric or electron-density-fit criteria:

| Mol | Type | Chain | Res | Chirality | Geometry | Clashes | Electron density |
|---|---|---|---|---|---|---|---|
| 4 | GOL | A | 209 | - | - | - | X |

For Manuscript Review

#### 2 Entry composition (i)

There are 5 unique types of molecules in this entry. The entry contains 1063 atoms, of which 0 are hydrogens and 0 are deuteriums.

In the tables below, the ZeroOcc column contains the number of atoms modelled with zero occupancy, the AltConf column contains the number of residues with at least one atom in alternate conformation and the Trace column contains the number of residues modelled with at most 2 atoms.

- Molecule 1 is a protein called Lysozyme C.

| Mol | Chain | Residues | Atoms | | | | | ZeroOcc | AltConf | Trace |
|---|---|---|---|---|---|---|---|---|---|---|
| | | | Total | C | N | O | S | | | |
| 1 | A | 129 | 1001 | 613 | 193 | 185 | 10 | 0 | 0 | 0 |

- Molecule 2 is SODIUM ION (three-letter code: NA) (formula: Na).

| Mol | Chain | Residues | Atoms | | ZeroOcc | AltConf |
|---|---|---|---|---|---|---|
| | | | Total | Na | | |
| 2 | A | 1 | 1 | 1 | 0 | 0 |

- Molecule 3 is CHLORIDE ION (three-letter code: CL) (formula: Cl).

| Mol | Chain | Residues | Atoms | | ZeroOcc | AltConf |
|---|---|---|---|---|---|---|
| | | | Total | Cl | | |
| 3 | A | 7 | 7 | 7 | 0 | 0 |

- Molecule 4 is GLYCEROL (three-letter code: GOL) (formula: $C_3H_8O_3$).

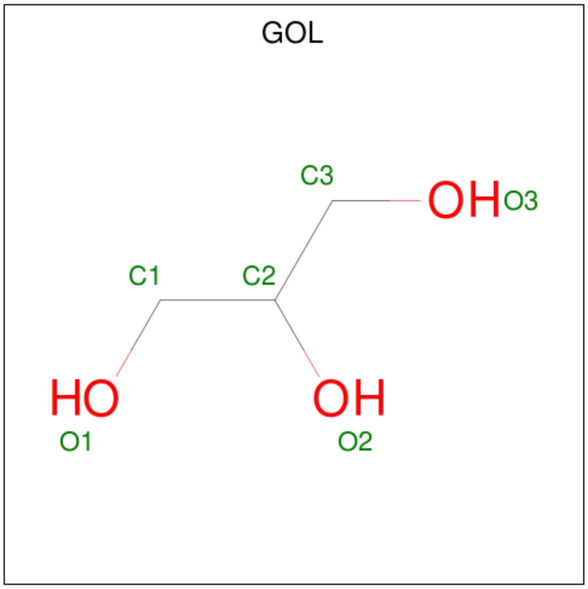


For Manuscript Review

| Mol | Chain | Residues | Atoms | ZeroOcc | AltConf |
|---|---|---|---|---|---|
| 4 | A | 1 | Total C O<br>6 3 3 | 0 | 0 |
| 4 | A | 1 | Total C O<br>6 3 3 | 0 | 0 |

- Molecule 5 is water.

| Mol | Chain | Residues | Atoms | ZeroOcc | AltConf |
|---|---|---|---|---|---|
| 5 | A | 42 | Total O<br>42 42 | 0 | 0 |

For Manuscript Review

### 3 Residue-property plots (i)

These plots are drawn for all protein, RNA, DNA and oligosaccharide chains in the entry. The first graphic for a chain summarises the proportions of the various outlier classes displayed in the second graphic. The second graphic shows the sequence view annotated by issues in geometry and electron density. Residues are color-coded according to the number of geometric quality criteria for which they contain at least one outlier: green = 0, yellow = 1, orange = 2 and red = 3 or more. A red dot above a residue indicates a poor fit to the electron density (RSRZ > 2). Stretches of 2 or more consecutive residues without any outlier are shown as a green connector. Residues present in the sample, but not in the model, are shown in grey.

- Molecule 1: Lysozyme C

Chain A:

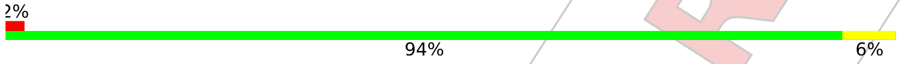


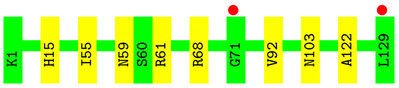


For Manuscript Review

### 4 Data and refinement statistics (i)

| Property | Value | Source |
|---|---|---|
| Space group | P 43 21 2 | Depositor |
| Cell constants<br>a, b, c, $\alpha$, $\beta$, $\gamma$ | 79.50Å 79.50Å 37.77Å<br>90.00° 90.00° 90.00° | Depositor |
| Resolution (Å) | 56.21 – 1.50<br>56.21 – 1.50 | Depositor<br>EDS |
| % Data completeness<br>(in resolution range) | 99.6 (56.21-1.50)<br>99.6 (56.21-1.50) | Depositor<br>EDS |
| $R_{merge}$ | (Not available) | Depositor |
| $R_{sym}$ | (Not available) | Depositor |
| $< I/\sigma(I) >$ [1] | 1.53 (at 1.50Å) | Xtriage |
| Refinement program | REFMAC 5.8.0267 | Depositor |
| R, $R_{free}$ | 0.175 , 0.196<br>0.183 , 0.207 | Depositor<br>DCC |
| $R_{free}$ test set | 1032 reflections (5.16%) | wwPDB-VP |
| Wilson B-factor (Å$^2$) | 18.3 | Xtriage |
| Anisotropy | 0.068 | Xtriage |
| Bulk solvent $k_{sol}$(e/Å$^3$), $B_{sol}$(Å$^2$) | 0.38 , 41.6 | EDS |
| L-test for twinning[2] | $< \lvert L \rvert >$ = 0.49, $< L^2 >$ = 0.33 | Xtriage |
| Estimated twinning fraction | No twinning to report. | Xtriage |
| $F_o$,$F_c$ correlation | 0.97 | EDS |
| Total number of atoms | 1063 | wwPDB-VP |
| Average B, all atoms (Å$^2$) | 23.0 | wwPDB-VP |

Xtriage's analysis on translational NCS is as follows: *The largest off-origin peak in the Patterson function is 10.19% of the height of the origin peak. No significant pseudotranslation is detected.*

[1] Intensities estimated from amplitudes.

[2] Theoretical values of $< |L| >$, $< L^2 >$ for acentric reflections are 0.5, 0.333 respectively for untwinned datasets, and 0.375, 0.2 for perfectly twinned datasets.

For Manuscript Review

### 5 Model quality (i)

#### 5.1 Standard geometry (i)

Bond lengths and bond angles in the following residue types are not validated in this section: GOL, CL, NA

The Z score for a bond length (or angle) is the number of standard deviations the observed value is removed from the expected value. A bond length (or angle) with $|Z| > 5$ is considered an outlier worth inspection. RMSZ is the root-mean-square of all Z scores of the bond lengths (or angles).

| Mol | Chain | Bond lengths | | Bond angles | |
|---|---|---|---|---|---|
| | | RMSZ | #$|Z|$ >5 | RMSZ | #$|Z|$ >5 |
| 1 | A | 0.79 | 0/1021 | 0.91 | 1/1379 (0.1%) |

There are no bond length outliers.

All (1) bond angle outliers are listed below:

| Mol | Chain | Res | Type | Atoms | Z | Observed($^o$) | Ideal($^o$) |
|---|---|---|---|---|---|---|---|
| 1 | A | 68 | ARG | NE-CZ-NH1 | -5.15 | 117.72 | 120.30 |

There are no chirality outliers.

There are no planarity outliers.

#### 5.2 Too-close contacts (i)

In the following table, the Non-H and H(model) columns list the number of non-hydrogen atoms and hydrogen atoms in the chain respectively. The H(added) column lists the number of hydrogen atoms added and optimized by MolProbity. The Clashes column lists the number of clashes within the asymmetric unit, whereas Symm-Clashes lists symmetry-related clashes.

| Mol | Chain | Non-H | H(model) | H(added) | Clashes | Symm-Clashes |
|---|---|---|---|---|---|---|
| 1 | A | 1001 | 0 | 958 | 7 | 0 |
| 2 | A | 1 | 0 | 0 | 0 | 0 |
| 3 | A | 7 | 0 | 0 | 0 | 0 |
| 4 | A | 12 | 0 | 16 | 1 | 0 |
| 5 | A | 42 | 0 | 0 | 0 | 0 |
| All | All | 1063 | 0 | 974 | 7 | 0 |

The all-atom clashscore is defined as the number of clashes found per 1000 atoms (including hydrogen atoms). The all-atom clashscore for this structure is 4.

For Manuscript Review

All (7) close contacts within the same asymmetric unit are listed below, sorted by their clash magnitude.

| Atom-1 | Atom-2 | Interatomic distance (Å) | Clash overlap (Å) |
|---|---|---|---|
| 1:A:59:ASN:ND2 | 1:A:61:ARG:HB3 | 2.05 | 0.71 |
| 1:A:59:ASN:ND2 | 1:A:61:ARG:H | 1.99 | 0.61 |
| 1:A:59:ASN:HD22 | 1:A:61:ARG:HB3 | 1.73 | 0.53 |
| 1:A:59:ASN:ND2 | 1:A:61:ARG:CB | 2.73 | 0.49 |
| 1:A:15:HIS:HB3 | 1:A:92:VAL:HG11 | 1.96 | 0.47 |
| 1:A:59:ASN:HD21 | 1:A:61:ARG:CB | 2.29 | 0.45 |
| 1:A:122:ALA:HB1 | 4:A:209:GOL:H32 | 2.01 | 0.42 |

There are no symmetry-related clashes.

#### 5.3 Torsion angles (i)

##### 5.3.1 Protein backbone (i)

In the following table, the Percentiles column shows the percent Ramachandran outliers of the chain as a percentile score with respect to all X-ray entries followed by that with respect to entries of similar resolution.

The Analysed column shows the number of residues for which the backbone conformation was analysed, and the total number of residues.

| Mol | Chain | Analysed | Favoured | Allowed | Outliers | Percentiles | |
|---|---|---|---|---|---|---|---|
| 1 | A | 127/129 (98%) | 126 (99%) | 1 (1%) | 0 | 100 | 100 |

There are no Ramachandran outliers to report.

##### 5.3.2 Protein sidechains (i)

In the following table, the Percentiles column shows the percent sidechain outliers of the chain as a percentile score with respect to all X-ray entries followed by that with respect to entries of similar resolution.

The Analysed column shows the number of residues for which the sidechain conformation was analysed, and the total number of residues.

| Mol | Chain | Analysed | Rotameric | Outliers | Percentiles | |
|---|---|---|---|---|---|---|
| 1 | A | 105/105 (100%) | 103 (98%) | 2 (2%) | 57 | 27 |

All (2) residues with a non-rotameric sidechain are listed below:

For Manuscript Review

| Mol | Chain | Res | Type |
|---|---|---|---|
| 1 | A | 55 | ILE |
| 1 | A | 103 | ASN |

Sometimes sidechains can be flipped to improve hydrogen bonding and reduce clashes. All (4) such sidechains are listed below:

| Mol | Chain | Res | Type |
|---|---|---|---|
| 1 | A | 15 | HIS |
| 1 | A | 46 | ASN |
| 1 | A | 59 | ASN |
| 1 | A | 103 | ASN |

##### 5.3.3 RNA (i)

There are no RNA molecules in this entry.

#### 5.4 Non-standard residues in protein, DNA, RNA chains (i)

There are no non-standard protein/DNA/RNA residues in this entry.

#### 5.5 Carbohydrates (i)

There are no monosaccharides in this entry.

#### 5.6 Ligand geometry (i)

Of 10 ligands modelled in this entry, 8 are monoatomic - leaving 2 for Mogul analysis.

In the following table, the Counts columns list the number of bonds (or angles) for which Mogul statistics could be retrieved, the number of bonds (or angles) that are observed in the model and the number of bonds (or angles) that are defined in the Chemical Component Dictionary. The Link column lists molecule types, if any, to which the group is linked. The Z score for a bond length (or angle) is the number of standard deviations the observed value is removed from the expected value. A bond length (or angle) with $|Z| > 2$ is considered an outlier worth inspection. RMSZ is the root-mean-square of all Z scores of the bond lengths (or angles).

| Mol | Type | Chain | Res | Link | Bond lengths | | | Bond angles | | |
|---|---|---|---|---|---|---|---|---|---|---|
| | | | | | Counts | RMSZ | $\#\|Z\| > 2$ | Counts | RMSZ | $\#\|Z\| > 2$ |
| 4 | GOL | A | 210 | - | 5,5,5 | 0.10 | 0 | 5,5,5 | 0.31 | 0 |
| 4 | GOL | A | 209 | - | 5,5,5 | 0.11 | 0 | 5,5,5 | 0.30 | 0 |

For Manuscript Review

In the following table, the Chirals column lists the number of chiral outliers, the number of chiral centers analysed, the number of these observed in the model and the number defined in the Chemical Component Dictionary. Similar counts are reported in the Torsion and Rings columns. '-' means no outliers of that kind were identified.

| Mol | Type | Chain | Res | Link | Chirals | Torsions | Rings |
|---|---|---|---|---|---|---|---|
| 4 | GOL | A | 210 | - | - | 2/4/4/4 | - |
| 4 | GOL | A | 209 | - | - | 2/4/4/4 | - |

There are no bond length outliers.

There are no bond angle outliers.

There are no chirality outliers.

All (4) torsion outliers are listed below:

| Mol | Chain | Res | Type | Atoms |
|---|---|---|---|---|
| 4 | A | 209 | GOL | O2-C2-C3-O3 |
| 4 | A | 210 | GOL | O1-C1-C2-O2 |
| 4 | A | 209 | GOL | C1-C2-C3-O3 |
| 4 | A | 210 | GOL | O1-C1-C2-C3 |

There are no ring outliers.

1 monomer is involved in 1 short contact:

| Mol | Chain | Res | Type | Clashes | Symm-Clashes |
|---|---|---|---|---|---|
| 4 | A | 209 | GOL | 1 | 0 |

#### 5.7 Other polymers (i)

There are no such residues in this entry.

#### 5.8 Polymer linkage issues (i)

There are no chain breaks in this entry.

For Manuscript Review

### 6 Fit of model and data (i)

#### 6.1 Protein, DNA and RNA chains (i)

In the following table, the column labelled '#RSRZ> 2' contains the number (and percentage) of RSRZ outliers, followed by percent RSRZ outliers for the chain as percentile scores relative to all X-ray entries and entries of similar resolution. The OWAB column contains the minimum, median, $95^{th}$ percentile and maximum values of the occupancy-weighted average B-factor per residue. The column labelled 'Q< 0.9' lists the number of (and percentage) of residues with an average occupancy less than 0.9.

| Mol | Chain | Analysed | <RSRZ> | #RSRZ>2 | | | OWAB(Å$^2$) | Q<0.9 |
|---|---|---|---|---|---|---|---|---|
| 1 | A | 129/129 (100%) | -0.24 | 2 (1%) | 72 | 77 | 12, 19, 37, 50 | 0 |

All (2) RSRZ outliers are listed below:

| Mol | Chain | Res | Type | RSRZ |
|---|---|---|---|---|
| 1 | A | 71 | GLY | 3.0 |
| 1 | A | 129 | LEU | 2.4 |

#### 6.2 Non-standard residues in protein, DNA, RNA chains (i)

There are no non-standard protein/DNA/RNA residues in this entry.

#### 6.3 Carbohydrates (i)

There are no monosaccharides in this entry.

#### 6.4 Ligands (i)

In the following table, the Atoms column lists the number of modelled atoms in the group and the number defined in the chemical component dictionary. The B-factors column lists the minimum, median, $95^{th}$ percentile and maximum values of B factors of atoms in the group. The column labelled 'Q< 0.9' lists the number of atoms with occupancy less than 0.9.

| Mol | Type | Chain | Res | Atoms | RSCC | RSR | B-factors(Å$^2$) | Q<0.9 |
|---|---|---|---|---|---|---|---|---|
| 4 | GOL | A | 209 | 6/6 | 0.47 | 0.46 | 75,79,81,82 | 0 |
| 4 | GOL | A | 210 | 6/6 | 0.51 | 0.28 | 77,79,81,81 | 0 |
| 3 | CL | A | 208 | 1/1 | 0.70 | 0.27 | 76,76,76,76 | 0 |
| 3 | CL | A | 205 | 1/1 | 0.71 | 0.21 | 65,65,65,65 | 0 |
| 3 | CL | A | 206 | 1/1 | 0.77 | 0.21 | 63,63,63,63 | 0 |

*Continued on next page...*

For Manuscript Review

*Continued from previous page...*

| Mol | Type | Chain | Res | Atoms | RSCC | RSR | B-factors($Å^2$) | Q<0.9 |
|---|---|---|---|---|---|---|---|---|
| 3 | CL | A | 207 | 1/1 | 0.90 | 0.16 | 52,52,52,52 | 0 |
| 3 | CL | A | 202 | 1/1 | 0.98 | 0.05 | 24,24,24,24 | 0 |
| 3 | CL | A | 204 | 1/1 | 0.98 | 0.09 | 31,31,31,31 | 0 |
| 3 | CL | A | 203 | 1/1 | 0.99 | 0.08 | 36,36,36,36 | 0 |
| 2 | NA | A | 201 | 1/1 | 0.99 | 0.09 | 21,21,21,21 | 0 |

#### 6.5 Other polymers ⓘ

There are no such residues in this entry.

For Manuscript Review

### Full wwPDB X-ray Structure Validation Report (i)

Jun 9, 2022 – 11:20 AM EDT

| | | |
|---|---|---|
| PDB ID | : | 8D8F |
| Title | : | Crystal structure of hen egg white lysozyme at 325 Kelvin |
| Deposited on | : | 2022-06-08 |
| Resolution | : | 1.50 Å(reported) |

**This wwPDB validation report is for manuscript review**

This is a Full wwPDB X-ray Structure Validation Report.

This report is produced by the wwPDB biocuration pipeline after annotation of the structure.

We welcome your comments at **
A user guide is available at
https://www.wwpdb.org/validation/2017/XrayValidationReportHelp
with specific help available everywhere you see the (i) symbol.

For Manuscript Review

The following versions of software and data (see references (i)) were used in the production of this report:

| | | |
|---|---|---|
| MolProbity | : | 4.02b-467 |
| Mogul | : | 1.8.5 (274361), CSD as541be (2020) |
| Xtriage (Phenix) | : | 1.13 |
| EDS | : | 2.28 |
| Percentile statistics | : | 20191225.v01 (using entries in the PDB archive December 25th 2019) |
| Refmac | : | 5.8.0158 |
| CCP4 | : | 7.0.044 (Gargrove) |
| Ideal geometry (proteins) | : | Engh & Huber (2001) |
| Ideal geometry (DNA, RNA) | : | Parkinson et al. (1996) |
| Validation Pipeline (wwPDB-VP) | : | 2.28 |

### 1 Overall quality at a glance (i)

The following experimental techniques were used to determine the structure:
*X-RAY DIFFRACTION*

The reported resolution of this entry is 1.50 Å.

Percentile scores (ranging between 0-100) for global validation metrics of the entry are shown in the following graphic. The table shows the number of entries on which the scores are based.

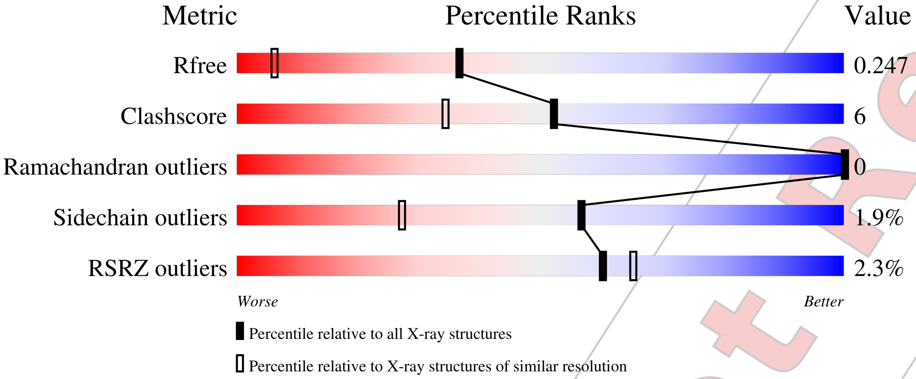


| Metric | Whole archive (#Entries) | Similar resolution (#Entries, resolution range(Å)) |
|---|---|---|
| $R_{free}$ | 130704 | 2936 (1.50-1.50) |
| Clashscore | 141614 | 3144 (1.50-1.50) |
| Ramachandran outliers | 138981 | 3066 (1.50-1.50) |
| Sidechain outliers | 138945 | 3064 (1.50-1.50) |
| RSRZ outliers | 127900 | 2884 (1.50-1.50) |

The table below summarises the geometric issues observed across the polymeric chains and their fit to the electron density. The red, orange, yellow and green segments of the lower bar indicate the fraction of residues that contain outliers for >=3, 2, 1 and 0 types of geometric quality criteria respectively. A grey segment represents the fraction of residues that are not modelled. The numeric value for each fraction is indicated below the corresponding segment, with a dot representing fractions <=5% The upper red bar (where present) indicates the fraction of residues that have poor fit to the electron density. The numeric value is given above the bar.

| Mol | Chain | Length | Quality of chain |
|---|---|---|---|
| 1 | A | 129 | 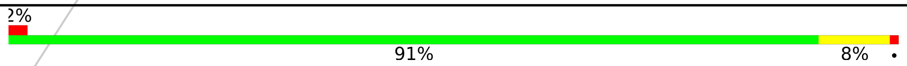  |

The following table lists non-polymeric compounds, carbohydrate monomers and non-standard residues in protein, DNA, RNA chains that are outliers for geometric or electron-density-fit criteria:

| Mol | Type | Chain | Res | Chirality | Geometry | Clashes | Electron density |
|---|---|---|---|---|---|---|---|
| 4 | GOL | A | 209 | - | - | - | X |

*Continued on next page...*



For Manuscript Review

*Continued from previous page...*

| Mol | Type | Chain | Res | Chirality | Geometry | Clashes | Electron density |
|---|---|---|---|---|---|---|---|
| 4 | GOL | A | 210 | - | - | - | X |

For Manuscript Review

#### 2 Entry composition (i)

There are 5 unique types of molecules in this entry. The entry contains 1051 atoms, of which 0 are hydrogens and 0 are deuteriums.

In the tables below, the ZeroOcc column contains the number of atoms modelled with zero occupancy, the AltConf column contains the number of residues with at least one atom in alternate conformation and the Trace column contains the number of residues modelled with at most 2 atoms.

- Molecule 1 is a protein called Lysozyme C.

| Mol | Chain | Residues | Atoms | | | | | ZeroOcc | AltConf | Trace |
|---|---|---|---|---|---|---|---|---|---|---|
| | | | Total | C | N | O | S | | | |
| 1 | A | 129 | 1001 | 613 | 193 | 185 | 10 | 0 | 0 | 0 |

- Molecule 2 is SODIUM ION (three-letter code: NA) (formula: Na).

| Mol | Chain | Residues | Atoms | | ZeroOcc | AltConf |
|---|---|---|---|---|---|---|
| | | | Total | Na | | |
| 2 | A | 1 | 1 | 1 | 0 | 0 |

- Molecule 3 is CHLORIDE ION (three-letter code: CL) (formula: Cl).

| Mol | Chain | Residues | Atoms | | ZeroOcc | AltConf |
|---|---|---|---|---|---|---|
| | | | Total | Cl | | |
| 3 | A | 7 | 7 | 7 | 0 | 0 |

- Molecule 4 is GLYCEROL (three-letter code: GOL) (formula: $C_3H_8O_3$).

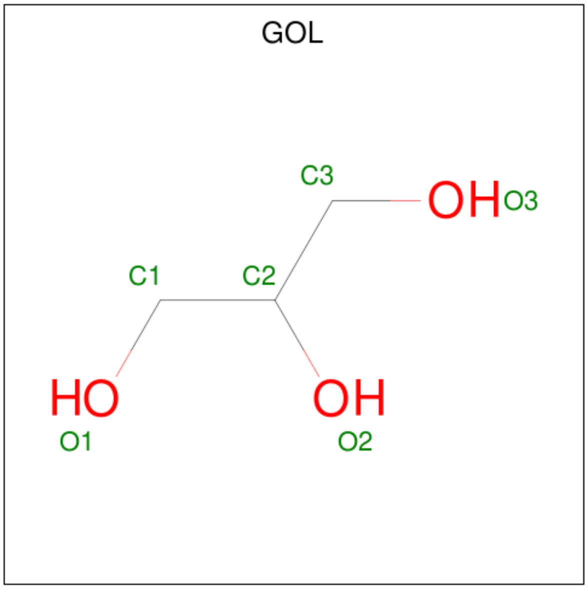


For Manuscript Review

| Mol | Chain | Residues | Atoms | ZeroOcc | AltConf |
|---|---|---|---|---|---|
| 4 | A | 1 | Total C O<br>6 3 3 | 0 | 0 |
| 4 | A | 1 | Total C O<br>6 3 3 | 0 | 0 |

- Molecule 5 is water.

| Mol | Chain | Residues | Atoms | ZeroOcc | AltConf |
|---|---|---|---|---|---|
| 5 | A | 30 | Total O<br>30 30 | 0 | 0 |

For Manuscript Review

### 3 Residue-property plots (i)

These plots are drawn for all protein, RNA, DNA and oligosaccharide chains in the entry. The first graphic for a chain summarises the proportions of the various outlier classes displayed in the second graphic. The second graphic shows the sequence view annotated by issues in geometry and electron density. Residues are color-coded according to the number of geometric quality criteria for which they contain at least one outlier: green = 0, yellow = 1, orange = 2 and red = 3 or more. A red dot above a residue indicates a poor fit to the electron density (RSRZ > 2). Stretches of 2 or more consecutive residues without any outlier are shown as a green connector. Residues present in the sample, but not in the model, are shown in grey.

- Molecule 1: Lysozyme C

Chain A:

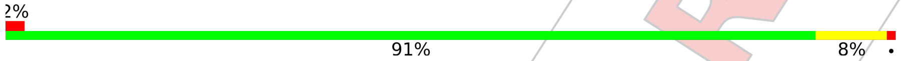


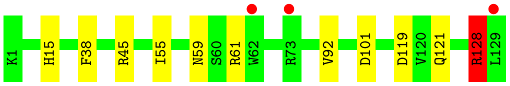


For Manuscript Review

### 4 Data and refinement statistics (i)

| Property | Value | Source |
|---|---|---|
| Space group | P 43 21 2 | Depositor |
| Cell constants<br>a, b, c, $\alpha$, $\beta$, $\gamma$ | 79.61Å 79.61Å 37.72Å<br>90.00° 90.00° 90.00° | Depositor |
| Resolution (Å) | 35.63 – 1.50<br>35.60 – 1.49 | Depositor<br>EDS |
| % Data completeness<br>(in resolution range) | 98.1 (35.63-1.50)<br>98.1 (35.60-1.49) | Depositor<br>EDS |
| $R_{merge}$ | (Not available) | Depositor |
| $R_{sym}$ | (Not available) | Depositor |
| $< I/\sigma(I) >$ [1] | 1.07 (at 1.50Å) | Xtriage |
| Refinement program | REFMAC 5.8.0267 | Depositor |
| R, $R_{free}$ | 0.202 , 0.243<br>0.210 , 0.247 | Depositor<br>DCC |
| $R_{free}$ test set | 1027 reflections (5.17%) | wwPDB-VP |
| Wilson B-factor (Å$^2$) | 19.7 | Xtriage |
| Anisotropy | 0.094 | Xtriage |
| Bulk solvent $k_{sol}$(e/Å$^3$), $B_{sol}$(Å$^2$) | 0.35 , 38.4 | EDS |
| L-test for twinning[2] | $< \|L\| > = 0.51$, $< L^2 > = 0.34$ | Xtriage |
| Estimated twinning fraction | No twinning to report. | Xtriage |
| $F_o$,$F_c$ correlation | 0.96 | EDS |
| Total number of atoms | 1051 | wwPDB-VP |
| Average B, all atoms (Å$^2$) | 27.0 | wwPDB-VP |

Xtriage's analysis on translational NCS is as follows: *The largest off-origin peak in the Patterson function is 12.12% of the height of the origin peak. No significant pseudotranslation is detected.*

[1] Intensities estimated from amplitudes.
[2] Theoretical values of $< |L| >$, $< L^2 >$ for acentric reflections are 0.5, 0.333 respectively for untwinned datasets, and 0.375, 0.2 for perfectly twinned datasets.

For Manuscript Review

### 5 Model quality (i)

#### 5.1 Standard geometry (i)

Bond lengths and bond angles in the following residue types are not validated in this section: CL, NA, GOL

The Z score for a bond length (or angle) is the number of standard deviations the observed value is removed from the expected value. A bond length (or angle) with $|Z| > 5$ is considered an outlier worth inspection. RMSZ is the root-mean-square of all Z scores of the bond lengths (or angles).

| Mol | Chain | Bond lengths | | Bond angles | |
|---|---|---|---|---|---|
| | | RMSZ | #$\|Z\|$ >5 | RMSZ | #$\|Z\|$ >5 |
| 1 | A | 0.73 | 0/1021 | 0.86 | 1/1379 (0.1%) |

There are no bond length outliers.

All (1) bond angle outliers are listed below:

| Mol | Chain | Res | Type | Atoms | Z | Observed($^o$) | Ideal($^o$) |
|---|---|---|---|---|---|---|---|
| 1 | A | 128 | ARG | NE-CZ-NH1 | 6.29 | 123.44 | 120.30 |

There are no chirality outliers.

There are no planarity outliers.

#### 5.2 Too-close contacts (i)

In the following table, the Non-H and H(model) columns list the number of non-hydrogen atoms and hydrogen atoms in the chain respectively. The H(added) column lists the number of hydrogen atoms added and optimized by MolProbity. The Clashes column lists the number of clashes within the asymmetric unit, whereas Symm-Clashes lists symmetry-related clashes.

| Mol | Chain | Non-H | H(model) | H(added) | Clashes | Symm-Clashes |
|---|---|---|---|---|---|---|
| 1 | A | 1001 | 0 | 958 | 11 | 0 |
| 2 | A | 1 | 0 | 0 | 0 | 0 |
| 3 | A | 7 | 0 | 0 | 0 | 0 |
| 4 | A | 12 | 0 | 16 | 1 | 0 |
| 5 | A | 30 | 0 | 0 | 2 | 0 |
| All | All | 1051 | 0 | 974 | 11 | 0 |

The all-atom clashscore is defined as the number of clashes found per 1000 atoms (including hydrogen atoms). The all-atom clashscore for this structure is 6.

For Manuscript Review

All (11) close contacts within the same asymmetric unit are listed below, sorted by their clash magnitude.

| Atom-1 | Atom-2 | Interatomic distance (Å) | Clash overlap (Å) |
|---|---|---|---|
| 1:A:119:ASP:OD1 | 1:A:121:GLN:NE2 | 2.05 | 0.89 |
| 1:A:128:ARG:HD2 | 5:A:330:HOH:O | 1.94 | 0.66 |
| 1:A:38:PHE:HA | 1:A:55:ILE:HD11 | 1.81 | 0.61 |
| 1:A:59:ASN:ND2 | 1:A:61:ARG:HB3 | 2.24 | 0.53 |
| 1:A:101:ASP:OD2 | 4:A:210:GOL:H11 | 2.09 | 0.53 |
| 1:A:15:HIS:HB3 | 1:A:92:VAL:HG11 | 1.91 | 0.52 |
| 1:A:121:GLN:H | 1:A:121:GLN:CD | 2.13 | 0.52 |
| 1:A:128:ARG:HH11 | 1:A:128:ARG:CG | 2.23 | 0.52 |
| 1:A:128:ARG:HH11 | 1:A:128:ARG:HG2 | 1.75 | 0.51 |
| 1:A:59:ASN:ND2 | 1:A:61:ARG:H | 2.10 | 0.49 |
| 1:A:128:ARG:CD | 5:A:330:HOH:O | 2.63 | 0.42 |

There are no symmetry-related clashes.

#### 5.3 Torsion angles (i)

##### 5.3.1 Protein backbone (i)

In the following table, the Percentiles column shows the percent Ramachandran outliers of the chain as a percentile score with respect to all X-ray entries followed by that with respect to entries of similar resolution.

The Analysed column shows the number of residues for which the backbone conformation was analysed, and the total number of residues.

| Mol | Chain | Analysed | Favoured | Allowed | Outliers | Percentiles | |
|---|---|---|---|---|---|---|---|
| 1 | A | 127/129 (98%) | 126 (99%) | 1 (1%) | 0 | 100 | 100 |

There are no Ramachandran outliers to report.

##### 5.3.2 Protein sidechains (i)

In the following table, the Percentiles column shows the percent sidechain outliers of the chain as a percentile score with respect to all X-ray entries followed by that with respect to entries of similar resolution.

The Analysed column shows the number of residues for which the sidechain conformation was analysed, and the total number of residues.

For Manuscript Review

| Mol | Chain | Analysed | Rotameric | Outliers | Percentiles | |
|---|---|---|---|---|---|---|
| 1 | A | 105/105 (100%) | 103 (98%) | 2 (2%) | 57 | 27 |

All (2) residues with a non-rotameric sidechain are listed below:

| Mol | Chain | Res | Type |
|---|---|---|---|
| 1 | A | 45 | ARG |
| 1 | A | 128 | ARG |

Sometimes sidechains can be flipped to improve hydrogen bonding and reduce clashes. All (4) such sidechains are listed below:

| Mol | Chain | Res | Type |
|---|---|---|---|
| 1 | A | 15 | HIS |
| 1 | A | 46 | ASN |
| 1 | A | 59 | ASN |
| 1 | A | 65 | ASN |

##### 5.3.3 RNA (i)

There are no RNA molecules in this entry.

#### 5.4 Non-standard residues in protein, DNA, RNA chains (i)

There are no non-standard protein/DNA/RNA residues in this entry.

#### 5.5 Carbohydrates (i)

There are no monosaccharides in this entry.

#### 5.6 Ligand geometry (i)

Of 10 ligands modelled in this entry, 8 are monoatomic - leaving 2 for Mogul analysis.

In the following table, the Counts columns list the number of bonds (or angles) for which Mogul statistics could be retrieved, the number of bonds (or angles) that are observed in the model and the number of bonds (or angles) that are defined in the Chemical Component Dictionary. The Link column lists molecule types, if any, to which the group is linked. The Z score for a bond length (or angle) is the number of standard deviations the observed value is removed from the expected value. A bond length (or angle) with $|Z| > 2$ is considered an outlier worth inspection. RMSZ is the root-mean-square of all Z scores of the bond lengths (or angles).

For Manuscript Review

| Mol | Type | Chain | Res | Link | Bond lengths | | | Bond angles | | |
|---|---|---|---|---|---|---|---|---|---|---|
| | | | | | Counts | RMSZ | #\|Z\| > 2 | Counts | RMSZ | #\|Z\| > 2 |
| 4 | GOL | A | 210 | - | 5,5,5 | 0.08 | 0 | 5,5,5 | 0.28 | 0 |
| 4 | GOL | A | 209 | - | 5,5,5 | 0.12 | 0 | 5,5,5 | 0.31 | 0 |

In the following table, the Chirals column lists the number of chiral outliers, the number of chiral centers analysed, the number of these observed in the model and the number defined in the Chemical Component Dictionary. Similar counts are reported in the Torsion and Rings columns. '-' means no outliers of that kind were identified.

| Mol | Type | Chain | Res | Link | Chirals | Torsions | Rings |
|---|---|---|---|---|---|---|---|
| 4 | GOL | A | 210 | - | - | 2/4/4/4 | - |
| 4 | GOL | A | 209 | - | - | 2/4/4/4 | - |

There are no bond length outliers.

There are no bond angle outliers.

There are no chirality outliers.

All (4) torsion outliers are listed below:

| Mol | Chain | Res | Type | Atoms |
|---|---|---|---|---|
| 4 | A | 209 | GOL | C1-C2-C3-O3 |
| 4 | A | 209 | GOL | O2-C2-C3-O3 |
| 4 | A | 210 | GOL | O1-C1-C2-O2 |
| 4 | A | 210 | GOL | O1-C1-C2-C3 |

There are no ring outliers.

1 monomer is involved in 1 short contact:

| Mol | Chain | Res | Type | Clashes | Symm-Clashes |
|---|---|---|---|---|---|
| 4 | A | 210 | GOL | 1 | 0 |

#### 5.7 Other polymers (i)

There are no such residues in this entry.

#### 5.8 Polymer linkage issues (i)

There are no chain breaks in this entry.

For Manuscript Review

### 6 Fit of model and data (i)

#### 6.1 Protein, DNA and RNA chains (i)

In the following table, the column labelled '#RSRZ> 2' contains the number (and percentage) of RSRZ outliers, followed by percent RSRZ outliers for the chain as percentile scores relative to all X-ray entries and entries of similar resolution. The OWAB column contains the minimum, median, $95^{th}$ percentile and maximum values of the occupancy-weighted average B-factor per residue. The column labelled 'Q< 0.9' lists the number of (and percentage) of residues with an average occupancy less than 0.9.

| Mol | Chain | Analysed | <RSRZ> | #RSRZ>2 | | | OWAB(Å$^2$) | Q<0.9 |
|---|---|---|---|---|---|---|---|---|
| 1 | A | 129/129 (100%) | 0.09 | 3 (2%) | 60 | 65 | 15, 24, 43, 51 | 0 |

All (3) RSRZ outliers are listed below:

| Mol | Chain | Res | Type | RSRZ |
|---|---|---|---|---|
| 1 | A | 62 | TRP | 2.6 |
| 1 | A | 73 | ARG | 2.6 |
| 1 | A | 129 | LEU | 2.0 |

#### 6.2 Non-standard residues in protein, DNA, RNA chains (i)

There are no non-standard protein/DNA/RNA residues in this entry.

#### 6.3 Carbohydrates (i)

There are no monosaccharides in this entry.

#### 6.4 Ligands (i)

In the following table, the Atoms column lists the number of modelled atoms in the group and the number defined in the chemical component dictionary. The B-factors column lists the minimum, median, $95^{th}$ percentile and maximum values of B factors of atoms in the group. The column labelled 'Q< 0.9' lists the number of atoms with occupancy less than 0.9.

| Mol | Type | Chain | Res | Atoms | RSCC | RSR | B-factors(Å$^2$) | Q<0.9 |
|---|---|---|---|---|---|---|---|---|
| 4 | GOL | A | 209 | 6/6 | 0.37 | 0.52 | 83,89,91,92 | 0 |
| 4 | GOL | A | 210 | 6/6 | 0.40 | 0.64 | 91,94,94,95 | 0 |
| 3 | CL | A | 206 | 1/1 | 0.56 | 0.31 | 68,68,68,68 | 0 |
| 3 | CL | A | 208 | 1/1 | 0.63 | 0.36 | 79,79,79,79 | 0 |

*Continued on next page...*

For Manuscript Review

*Continued from previous page...*

| Mol | Type | Chain | Res | Atoms | RSCC | RSR | B-factors($Å^2$) | Q<0.9 |
|---|---|---|---|---|---|---|---|---|
| 3 | CL | A | 205 | 1/1 | 0.74 | 0.30 | 76,76,76,76 | 0 |
| 3 | CL | A | 207 | 1/1 | 0.91 | 0.12 | 58,58,58,58 | 0 |
| 3 | CL | A | 203 | 1/1 | 0.96 | 0.18 | 48,48,48,48 | 0 |
| 2 | NA | A | 201 | 1/1 | 0.97 | 0.16 | 32,32,32,32 | 0 |
| 3 | CL | A | 204 | 1/1 | 0.97 | 0.06 | 39,39,39,39 | 0 |
| 3 | CL | A | 202 | 1/1 | 0.99 | 0.04 | 29,29,29,29 | 0 |

#### 6.5 Other polymers (i)

There are no such residues in this entry.

For Manuscript Review

### Full wwPDB X-ray Structure Validation Report (i)

Jun 9, 2022 – 11:22 AM EDT

| | | |
|---|---|---|
| PDB ID | : | 8D8G |
| Title | : | Crystal structure of hen egg white lysozyme at 325 Kelvin (Duplicate) |
| Deposited on | : | 2022-06-08 |
| Resolution | : | 1.50 Å(reported) |

**This wwPDB validation report is for manuscript review**

This is a Full wwPDB X-ray Structure Validation Report.

This report is produced by the wwPDB biocuration pipeline after annotation of the structure.

We welcome your comments at **
A user guide is available at
https://www.wwpdb.org/validation/2017/XrayValidationReportHelp
with specific help available everywhere you see the (i) symbol.

The following versions of software and data (see references (i)) were used in the production of this report:

| | | |
|---|---|---|
| MolProbity | : | 4.02b-467 |
| Mogul | : | 1.8.5 (274361), CSD as541be (2020) |
| Xtriage (Phenix) | : | 1.13 |
| EDS | : | 2.28 |
| Percentile statistics | : | 20191225.v01 (using entries in the PDB archive December 25th 2019) |
| Refmac | : | 5.8.0158 |
| CCP4 | : | 7.0.044 (Gargrove) |
| Ideal geometry (proteins) | : | Engh & Huber (2001) |
| Ideal geometry (DNA, RNA) | : | Parkinson et al. (1996) |
| Validation Pipeline (wwPDB-VP) | : | 2.28 |

For Manuscript Review

### 1 Overall quality at a glance (i)

The following experimental techniques were used to determine the structure:
*X-RAY DIFFRACTION*

The reported resolution of this entry is 1.50 Å.

Percentile scores (ranging between 0-100) for global validation metrics of the entry are shown in the following graphic. The table shows the number of entries on which the scores are based.

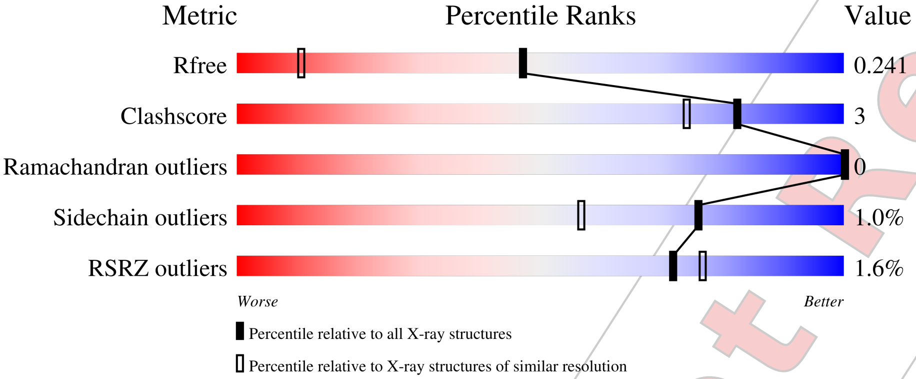


| Metric | Whole archive (#Entries) | Similar resolution (#Entries, resolution range(Å)) |
|---|---|---|
| $R_{free}$ | 130704 | 2936 (1.50-1.50) |
| Clashscore | 141614 | 3144 (1.50-1.50) |
| Ramachandran outliers | 138981 | 3066 (1.50-1.50) |
| Sidechain outliers | 138945 | 3064 (1.50-1.50) |
| RSRZ outliers | 127900 | 2884 (1.50-1.50) |

The table below summarises the geometric issues observed across the polymeric chains and their fit to the electron density. The red, orange, yellow and green segments of the lower bar indicate the fraction of residues that contain outliers for >=3, 2, 1 and 0 types of geometric quality criteria respectively. A grey segment represents the fraction of residues that are not modelled. The numeric value for each fraction is indicated below the corresponding segment, with a dot representing fractions <=5% The upper red bar (where present) indicates the fraction of residues that have poor fit to the electron density. The numeric value is given above the bar.

| Mol | Chain | Length | Quality of chain |
|---|---|---|---|
| 1 | A | 129 | 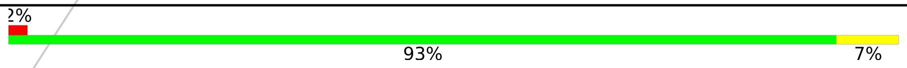  |

The following table lists non-polymeric compounds, carbohydrate monomers and non-standard residues in protein, DNA, RNA chains that are outliers for geometric or electron-density-fit criteria:

| Mol | Type | Chain | Res | Chirality | Geometry | Clashes | Electron density |
|---|---|---|---|---|---|---|---|
| 3 | CL | A | 208 | - | - | - | X |

*Continued on next page...*

For Manuscript Review

*Continued from previous page...*

| Mol | Type | Chain | Res | Chirality | Geometry | Clashes | Electron density |
|---|---|---|---|---|---|---|---|
| 4 | GOL | A | 209 | - | - | - | X |
| 4 | GOL | A | 210 | - | - | - | X |

For Manuscript Review

#### 2 Entry composition (i)

There are 5 unique types of molecules in this entry. The entry contains 1052 atoms, of which 0 are hydrogens and 0 are deuteriums.

In the tables below, the ZeroOcc column contains the number of atoms modelled with zero occupancy, the AltConf column contains the number of residues with at least one atom in alternate conformation and the Trace column contains the number of residues modelled with at most 2 atoms.

- Molecule 1 is a protein called Lysozyme C.

| Mol | Chain | Residues | Atoms | | | | | ZeroOcc | AltConf | Trace |
|---|---|---|---|---|---|---|---|---|---|---|
| | | | Total | C | N | O | S | | | |
| 1 | A | 129 | 1001 | 613 | 193 | 185 | 10 | 0 | 0 | 0 |

- Molecule 2 is SODIUM ION (three-letter code: NA) (formula: Na).

| Mol | Chain | Residues | Atoms | | ZeroOcc | AltConf |
|---|---|---|---|---|---|---|
| | | | Total | Na | | |
| 2 | A | 1 | 1 | 1 | 0 | 0 |

- Molecule 3 is CHLORIDE ION (three-letter code: CL) (formula: Cl).

| Mol | Chain | Residues | Atoms | | ZeroOcc | AltConf |
|---|---|---|---|---|---|---|
| | | | Total | Cl | | |
| 3 | A | 7 | 7 | 7 | 0 | 0 |

- Molecule 4 is GLYCEROL (three-letter code: GOL) (formula: $C_3H_8O_3$).

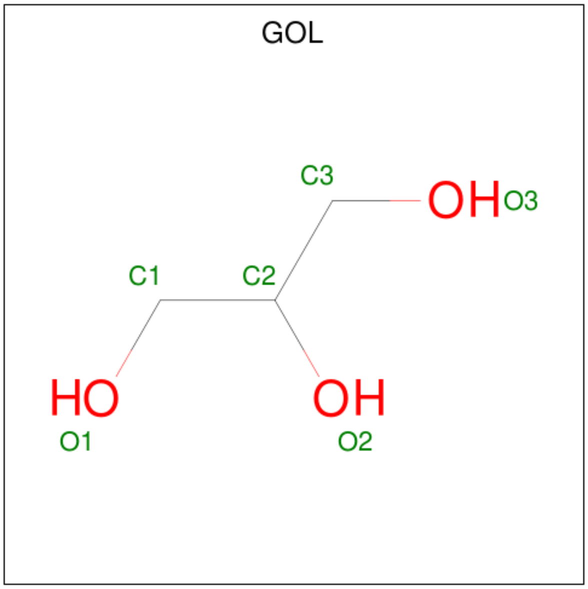


For Manuscript Review

| Mol | Chain | Residues | Atoms | ZeroOcc | AltConf |
|---|---|---|---|---|---|
| 4 | A | 1 | Total C O<br>6 3 3 | 0 | 0 |
| 4 | A | 1 | Total C O<br>6 3 3 | 0 | 0 |

- Molecule 5 is water.

| Mol | Chain | Residues | Atoms | ZeroOcc | AltConf |
|---|---|---|---|---|---|
| 5 | A | 31 | Total O<br>31 31 | 0 | 0 |

For Manuscript Review

### 3 Residue-property plots (i)

These plots are drawn for all protein, RNA, DNA and oligosaccharide chains in the entry. The first graphic for a chain summarises the proportions of the various outlier classes displayed in the second graphic. The second graphic shows the sequence view annotated by issues in geometry and electron density. Residues are color-coded according to the number of geometric quality criteria for which they contain at least one outlier: green = 0, yellow = 1, orange = 2 and red = 3 or more. A red dot above a residue indicates a poor fit to the electron density (RSRZ > 2). Stretches of 2 or more consecutive residues without any outlier are shown as a green connector. Residues present in the sample, but not in the model, are shown in grey.

• Molecule 1: Lysozyme C

Chain A:
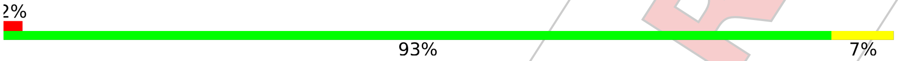


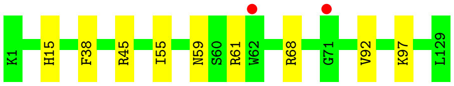


For Manuscript Review

### 4 Data and refinement statistics (i)

| Property | Value | Source |
|---|---|---|
| Space group | P 43 21 2 | Depositor |
| Cell constants<br>a, b, c, $\alpha$, $\beta$, $\gamma$ | 79.81Å 79.81Å 37.71Å<br>90.00° 90.00° 90.00° | Depositor |
| Resolution (Å) | 35.72 – 1.50<br>35.69 – 1.49 | Depositor<br>EDS |
| % Data completeness<br>(in resolution range) | 97.5 (35.72-1.50)<br>97.6 (35.69-1.49) | Depositor<br>EDS |
| $R_{merge}$ | (Not available) | Depositor |
| $R_{sym}$ | (Not available) | Depositor |
| $< I/\sigma(I) >$ [1] | 1.08 (at 1.50Å) | Xtriage |
| Refinement program | REFMAC 5.8.0267 | Depositor |
| R, $R_{free}$ | 0.200 , 0.236<br>0.208 , 0.241 | Depositor<br>DCC |
| $R_{free}$ test set | 1077 reflections (5.42%) | wwPDB-VP |
| Wilson B-factor (Å$^2$) | 19.8 | Xtriage |
| Anisotropy | 0.102 | Xtriage |
| Bulk solvent $k_{sol}$(e/Å$^3$), $B_{sol}$(Å$^2$) | 0.35 , 38.3 | EDS |
| L-test for twinning[2] | $< \|L\| >$ = 0.50, $< L^2 >$ = 0.33 | Xtriage |
| Estimated twinning fraction | No twinning to report. | Xtriage |
| $F_o$,$F_c$ correlation | 0.96 | EDS |
| Total number of atoms | 1052 | wwPDB-VP |
| Average B, all atoms (Å$^2$) | 27.0 | wwPDB-VP |

Xtriage's analysis on translational NCS is as follows: *The largest off-origin peak in the Patterson function is 12.73% of the height of the origin peak. No significant pseudotranslation is detected.*

[1]Intensities estimated from amplitudes.

[2]Theoretical values of $< |L| >$, $< L^2 >$ for acentric reflections are 0.5, 0.333 respectively for untwinned datasets, and 0.375, 0.2 for perfectly twinned datasets.

For Manuscript Review

### 5 Model quality (i)

#### 5.1 Standard geometry (i)

Bond lengths and bond angles in the following residue types are not validated in this section: GOL, NA, CL

The Z score for a bond length (or angle) is the number of standard deviations the observed value is removed from the expected value. A bond length (or angle) with $|Z| > 5$ is considered an outlier worth inspection. RMSZ is the root-mean-square of all Z scores of the bond lengths (or angles).

| Mol | Chain | Bond lengths | | Bond angles | |
|---|---|---|---|---|---|
| | | RMSZ | #$\|Z\|$ >5 | RMSZ | #$\|Z\|$ >5 |
| 1 | A | 0.75 | 0/1021 | 0.86 | 0/1379 |

There are no bond length outliers.

There are no bond angle outliers.

There are no chirality outliers.

There are no planarity outliers.

#### 5.2 Too-close contacts (i)

In the following table, the Non-H and H(model) columns list the number of non-hydrogen atoms and hydrogen atoms in the chain respectively. The H(added) column lists the number of hydrogen atoms added and optimized by MolProbity. The Clashes column lists the number of clashes within the asymmetric unit, whereas Symm-Clashes lists symmetry-related clashes.

| Mol | Chain | Non-H | H(model) | H(added) | Clashes | Symm-Clashes |
|---|---|---|---|---|---|---|
| 1 | A | 1001 | 0 | 958 | 5 | 0 |
| 2 | A | 1 | 0 | 0 | 0 | 0 |
| 3 | A | 7 | 0 | 0 | 0 | 0 |
| 4 | A | 12 | 0 | 16 | 0 | 0 |
| 5 | A | 31 | 0 | 0 | 0 | 0 |
| All | All | 1052 | 0 | 974 | 5 | 0 |

The all-atom clashscore is defined as the number of clashes found per 1000 atoms (including hydrogen atoms). The all-atom clashscore for this structure is 3.

All (5) close contacts within the same asymmetric unit are listed below, sorted by their clash magnitude.

For Manuscript Review

| Atom-1 | Atom-2 | Interatomic distance (Å) | Clash overlap (Å) |
|---|---|---|---|
| 1:A:59:ASN:ND2 | 1:A:61:ARG:HB3 | 2.18 | 0.59 |
| 1:A:15:HIS:HB3 | 1:A:92:VAL:HG11 | 1.93 | 0.51 |
| 1:A:38:PHE:HA | 1:A:55:ILE:HD11 | 1.92 | 0.50 |
| 1:A:59:ASN:ND2 | 1:A:61:ARG:H | 2.11 | 0.48 |
| 1:A:45:ARG:NH2 | 1:A:68:ARG:NE | 2.69 | 0.40 |

There are no symmetry-related clashes.

#### 5.3 Torsion angles (i)

##### 5.3.1 Protein backbone (i)

In the following table, the Percentiles column shows the percent Ramachandran outliers of the chain as a percentile score with respect to all X-ray entries followed by that with respect to entries of similar resolution.

The Analysed column shows the number of residues for which the backbone conformation was analysed, and the total number of residues.

| Mol | Chain | Analysed | Favoured | Allowed | Outliers | Percentiles | |
|---|---|---|---|---|---|---|---|
| 1 | A | 127/129 (98%) | 126 (99%) | 1 (1%) | 0 | 100 | 100 |

There are no Ramachandran outliers to report.

##### 5.3.2 Protein sidechains (i)

In the following table, the Percentiles column shows the percent sidechain outliers of the chain as a percentile score with respect to all X-ray entries followed by that with respect to entries of similar resolution.

The Analysed column shows the number of residues for which the sidechain conformation was analysed, and the total number of residues.

| Mol | Chain | Analysed | Rotameric | Outliers | Percentiles | |
|---|---|---|---|---|---|---|
| 1 | A | 105/105 (100%) | 104 (99%) | 1 (1%) | 76 | 57 |

All (1) residues with a non-rotameric sidechain are listed below:

| Mol | Chain | Res | Type |
|---|---|---|---|
| 1 | A | 97 | LYS |

For Manuscript Review

Sometimes sidechains can be flipped to improve hydrogen bonding and reduce clashes. All (4) such sidechains are listed below:

| Mol | Chain | Res | Type |
|---|---|---|---|
| 1 | A | 15 | HIS |
| 1 | A | 46 | ASN |
| 1 | A | 59 | ASN |
| 1 | A | 65 | ASN |

###### 5.3.3 RNA (i)

There are no RNA molecules in this entry.

##### 5.4 Non-standard residues in protein, DNA, RNA chains (i)

There are no non-standard protein/DNA/RNA residues in this entry.

##### 5.5 Carbohydrates (i)

There are no monosaccharides in this entry.

##### 5.6 Ligand geometry (i)

Of 10 ligands modelled in this entry, 8 are monoatomic - leaving 2 for Mogul analysis.

In the following table, the Counts columns list the number of bonds (or angles) for which Mogul statistics could be retrieved, the number of bonds (or angles) that are observed in the model and the number of bonds (or angles) that are defined in the Chemical Component Dictionary. The Link column lists molecule types, if any, to which the group is linked. The Z score for a bond length (or angle) is the number of standard deviations the observed value is removed from the expected value. A bond length (or angle) with $|Z| > 2$ is considered an outlier worth inspection. RMSZ is the root-mean-square of all Z scores of the bond lengths (or angles).

| Mol | Type | Chain | Res | Link | Bond lengths | | | Bond angles | | |
|---|---|---|---|---|---|---|---|---|---|---|
| | | | | | Counts | RMSZ | $\#\|Z\| > 2$ | Counts | RMSZ | $\#\|Z\| > 2$ |
| 4 | GOL | A | 209 | - | 5,5,5 | 0.12 | 0 | 5,5,5 | 0.33 | 0 |
| 4 | GOL | A | 210 | - | 5,5,5 | 0.09 | 0 | 5,5,5 | 0.28 | 0 |

In the following table, the Chirals column lists the number of chiral outliers, the number of chiral centers analysed, the number of these observed in the model and the number defined in the Chemical Component Dictionary. Similar counts are reported in the Torsion and Rings columns. '-' means no outliers of that kind were identified.

For Manuscript Review

| Mol | Type | Chain | Res | Link | Chirals | Torsions | Rings |
|---|---|---|---|---|---|---|---|
| 4 | GOL | A | 209 | - | - | 2/4/4/4 | - |
| 4 | GOL | A | 210 | - | - | 0/4/4/4 | - |

There are no bond length outliers.

There are no bond angle outliers.

There are no chirality outliers.

All (2) torsion outliers are listed below:

| Mol | Chain | Res | Type | Atoms |
|---|---|---|---|---|
| 4 | A | 209 | GOL | C1-C2-C3-O3 |
| 4 | A | 209 | GOL | O2-C2-C3-O3 |

There are no ring outliers.

No monomer is involved in short contacts.

#### 5.7 Other polymers (i)

There are no such residues in this entry.

#### 5.8 Polymer linkage issues (i)

There are no chain breaks in this entry.

For Manuscript Review

### 6 Fit of model and data (i)

#### 6.1 Protein, DNA and RNA chains (i)

In the following table, the column labelled '#RSRZ> 2' contains the number (and percentage) of RSRZ outliers, followed by percent RSRZ outliers for the chain as percentile scores relative to all X-ray entries and entries of similar resolution. The OWAB column contains the minimum, median, $95^{th}$ percentile and maximum values of the occupancy-weighted average B-factor per residue. The column labelled 'Q< 0.9' lists the number of (and percentage) of residues with an average occupancy less than 0.9.

| Mol | Chain | Analysed | <RSRZ> | #RSRZ>2 | | | OWAB(Å$^2$) | Q<0.9 |
|---|---|---|---|---|---|---|---|---|
| 1 | A | 129/129 (100%) | -0.04 | 2 (1%) | 72 | 77 | 15, 24, 44, 51 | 0 |

All (2) RSRZ outliers are listed below:

| Mol | Chain | Res | Type | RSRZ |
|---|---|---|---|---|
| 1 | A | 71 | GLY | 2.5 |
| 1 | A | 62 | TRP | 2.3 |

#### 6.2 Non-standard residues in protein, DNA, RNA chains (i)

There are no non-standard protein/DNA/RNA residues in this entry.

#### 6.3 Carbohydrates (i)

There are no monosaccharides in this entry.

#### 6.4 Ligands (i)

In the following table, the Atoms column lists the number of modelled atoms in the group and the number defined in the chemical component dictionary. The B-factors column lists the minimum, median, $95^{th}$ percentile and maximum values of B factors of atoms in the group. The column labelled 'Q< 0.9' lists the number of atoms with occupancy less than 0.9.

| Mol | Type | Chain | Res | Atoms | RSCC | RSR | B-factors(Å$^2$) | Q<0.9 |
|---|---|---|---|---|---|---|---|---|
| 4 | GOL | A | 210 | 6/6 | 0.39 | 0.56 | 86,89,90,90 | 0 |
| 3 | CL | A | 206 | 1/1 | 0.51 | 0.27 | 68,68,68,68 | 0 |
| 4 | GOL | A | 209 | 6/6 | 0.53 | 0.51 | 89,92,95,96 | 0 |
| 3 | CL | A | 205 | 1/1 | 0.69 | 0.25 | 74,74,74,74 | 0 |
| 3 | CL | A | 208 | 1/1 | 0.76 | 0.44 | 82,82,82,82 | 0 |

*Continued on next page...*

For Manuscript Review

*Continued from previous page...*

| Mol | Type | Chain | Res | Atoms | RSCC | RSR | B-factors($Å^2$) | Q<0.9 |
|---|---|---|---|---|---|---|---|---|
| 3 | CL | A | 207 | 1/1 | 0.93 | 0.15 | 59,59,59,59 | 0 |
| 2 | NA | A | 201 | 1/1 | 0.96 | 0.14 | 31,31,31,31 | 0 |
| 3 | CL | A | 204 | 1/1 | 0.97 | 0.07 | 39,39,39,39 | 0 |
| 3 | CL | A | 203 | 1/1 | 0.98 | 0.19 | 48,48,48,48 | 0 |
| 3 | CL | A | 202 | 1/1 | 0.98 | 0.05 | 29,29,29,29 | 0 |

#### 6.5 Other polymers (i)

There are no such residues in this entry.

For Manuscript Review

For Manuscript Review

### Full wwPDB X-ray Structure Validation Report (i)

Jun 9, 2022 – 11:24 AM EDT

| | | |
|---|---|---|
| PDB ID | : | 8D8H |
| Title | : | Crystal structure of hen egg white lysozyme at 325 Kelvin (Triplicate) |
| Deposited on | : | 2022-06-08 |
| Resolution | : | 1.50 Å(reported) |

**This wwPDB validation report is for manuscript review**

This is a Full wwPDB X-ray Structure Validation Report.

This report is produced by the wwPDB biocuration pipeline after annotation of the structure.

We welcome your comments at **
A user guide is available at
https://www.wwpdb.org/validation/2017/XrayValidationReportHelp
with specific help available everywhere you see the (i) symbol.

The following versions of software and data (see references (i)) were used in the production of this report:

| | | |
|---|---|---|
| MolProbity | : | 4.02b-467 |
| Mogul | : | 1.8.5 (274361), CSD as541be (2020) |
| Xtriage (Phenix) | : | 1.13 |
| EDS | : | 2.28 |
| Percentile statistics | : | 20191225.v01 (using entries in the PDB archive December 25th 2019) |
| Refmac | : | 5.8.0158 |
| CCP4 | : | 7.0.044 (Gargrove) |
| Ideal geometry (proteins) | : | Engh & Huber (2001) |
| Ideal geometry (DNA, RNA) | : | Parkinson et al. (1996) |
| Validation Pipeline (wwPDB-VP) | : | 2.28 |

### 1 Overall quality at a glance (i)

The following experimental techniques were used to determine the structure:
*X-RAY DIFFRACTION*

The reported resolution of this entry is 1.50 Å.

Percentile scores (ranging between 0-100) for global validation metrics of the entry are shown in the following graphic. The table shows the number of entries on which the scores are based.

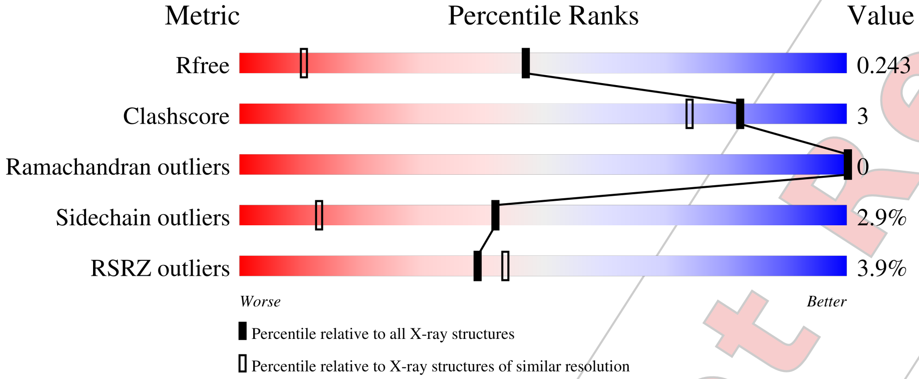


| Metric | Whole archive (#Entries) | Similar resolution (#Entries, resolution range(Å)) |
|---|---|---|
| $R_{free}$ | 130704 | 2936 (1.50-1.50) |
| Clashscore | 141614 | 3144 (1.50-1.50) |
| Ramachandran outliers | 138981 | 3066 (1.50-1.50) |
| Sidechain outliers | 138945 | 3064 (1.50-1.50) |
| RSRZ outliers | 127900 | 2884 (1.50-1.50) |

The table below summarises the geometric issues observed across the polymeric chains and their fit to the electron density. The red, orange, yellow and green segments of the lower bar indicate the fraction of residues that contain outliers for >=3, 2, 1 and 0 types of geometric quality criteria respectively. A grey segment represents the fraction of residues that are not modelled. The numeric value for each fraction is indicated below the corresponding segment, with a dot representing fractions <=5% The upper red bar (where present) indicates the fraction of residues that have poor fit to the electron density. The numeric value is given above the bar.

| Mol | Chain | Length | Quality of chain |
|---|---|---|---|
| 1 | A | 129 | 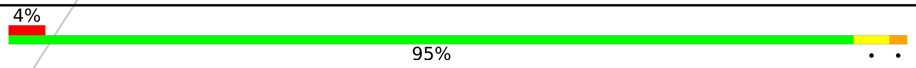 |

The following table lists non-polymeric compounds, carbohydrate monomers and non-standard residues in protein, DNA, RNA chains that are outliers for geometric or electron-density-fit criteria:

| Mol | Type | Chain | Res | Chirality | Geometry | Clashes | Electron density |
|---|---|---|---|---|---|---|---|
| 4 | GOL | A | 209 | - | - | - | X |

For Manuscript Review

### 2 Entry composition (i)

There are 5 unique types of molecules in this entry. The entry contains 1048 atoms, of which 0 are hydrogens and 0 are deuteriums.

In the tables below, the ZeroOcc column contains the number of atoms modelled with zero occupancy, the AltConf column contains the number of residues with at least one atom in alternate conformation and the Trace column contains the number of residues modelled with at most 2 atoms.

- Molecule 1 is a protein called Lysozyme C.

| Mol | Chain | Residues | Atoms | | | | | ZeroOcc | AltConf | Trace |
|---|---|---|---|---|---|---|---|---|---|---|
| | | | Total | C | N | O | S | | | |
| 1 | A | 129 | 1001 | 613 | 193 | 185 | 10 | 0 | 0 | 0 |

- Molecule 2 is SODIUM ION (three-letter code: NA) (formula: Na).

| Mol | Chain | Residues | Atoms | | ZeroOcc | AltConf |
|---|---|---|---|---|---|---|
| | | | Total | Na | | |
| 2 | A | 1 | 1 | 1 | 0 | 0 |

- Molecule 3 is CHLORIDE ION (three-letter code: CL) (formula: Cl).

| Mol | Chain | Residues | Atoms | | ZeroOcc | AltConf |
|---|---|---|---|---|---|---|
| | | | Total | Cl | | |
| 3 | A | 7 | 7 | 7 | 0 | 0 |

- Molecule 4 is GLYCEROL (three-letter code: GOL) (formula: $C_3H_8O_3$).

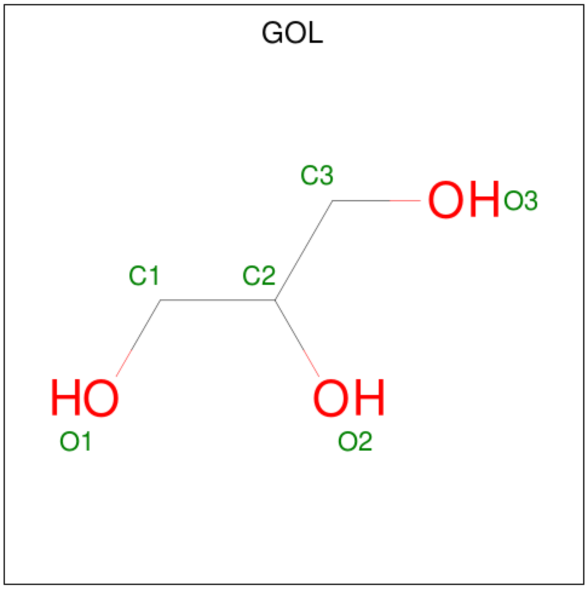


For Manuscript Review

| Mol | Chain | Residues | Atoms | ZeroOcc | AltConf |
|---|---|---|---|---|---|
| 4 | A | 1 | Total C O<br>6 3 3 | 0 | 0 |
| 4 | A | 1 | Total C O<br>6 3 3 | 0 | 0 |

- Molecule 5 is water.

| Mol | Chain | Residues | Atoms | ZeroOcc | AltConf |
|---|---|---|---|---|---|
| 5 | A | 27 | Total O<br>27 27 | 0 | 0 |

For Manuscript Review

### 3 Residue-property plots (i)

These plots are drawn for all protein, RNA, DNA and oligosaccharide chains in the entry. The first graphic for a chain summarises the proportions of the various outlier classes displayed in the second graphic. The second graphic shows the sequence view annotated by issues in geometry and electron density. Residues are color-coded according to the number of geometric quality criteria for which they contain at least one outlier: green = 0, yellow = 1, orange = 2 and red = 3 or more. A red dot above a residue indicates a poor fit to the electron density (RSRZ > 2). Stretches of 2 or more consecutive residues without any outlier are shown as a green connector. Residues present in the sample, but not in the model, are shown in grey.

- Molecule 1: Lysozyme C

Chain A:

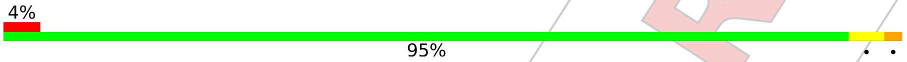


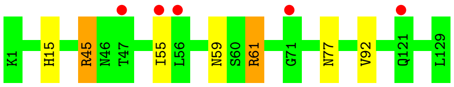


For Manuscript Review

### 4 Data and refinement statistics (i)

| Property | Value | Source |
|---|---|---|
| Space group | P 43 21 2 | Depositor |
| Cell constants<br>a, b, c, $\alpha$, $\beta$, $\gamma$ | 79.89Å 79.89Å 37.80Å<br>90.00° 90.00° 90.00° | Depositor |
| Resolution (Å) | 22.64 – 1.50<br>22.63 – 1.50 | Depositor<br>EDS |
| % Data completeness<br>(in resolution range) | 98.3 (22.64-1.50)<br>98.3 (22.63-1.50) | Depositor<br>EDS |
| $R_{merge}$ | (Not available) | Depositor |
| $R_{sym}$ | (Not available) | Depositor |
| $< I/\sigma(I) >$ [1] | 1.45 (at 1.50Å) | Xtriage |
| Refinement program | REFMAC 5.8.0267 | Depositor |
| R, $R_{free}$ | 0.213 , 0.241<br>0.221 , 0.243 | Depositor<br>DCC |
| $R_{free}$ test set | 1013 reflections (5.11%) | wwPDB-VP |
| Wilson B-factor (Å$^2$) | 17.4 | Xtriage |
| Anisotropy | 0.012 | Xtriage |
| Bulk solvent $k_{sol}$(e/Å$^3$), $B_{sol}$(Å$^2$) | 0.39 , 45.1 | EDS |
| L-test for twinning[2] | $< \|L\| >$ = 0.50, $< L^2 >$ = 0.34 | Xtriage |
| Estimated twinning fraction | No twinning to report. | Xtriage |
| $F_o$,$F_c$ correlation | 0.97 | EDS |
| Total number of atoms | 1048 | wwPDB-VP |
| Average B, all atoms (Å$^2$) | 27.0 | wwPDB-VP |

Xtriage's analysis on translational NCS is as follows: *The largest off-origin peak in the Patterson function is 11.51% of the height of the origin peak. No significant pseudotranslation is detected.*

[1] Intensities estimated from amplitudes.

[2] Theoretical values of $< |L| >$, $< L^2 >$ for acentric reflections are 0.5, 0.333 respectively for untwinned datasets, and 0.375, 0.2 for perfectly twinned datasets.

For Manuscript Review

### 5 Model quality (i)

#### 5.1 Standard geometry (i)

Bond lengths and bond angles in the following residue types are not validated in this section: CL, GOL, NA

The Z score for a bond length (or angle) is the number of standard deviations the observed value is removed from the expected value. A bond length (or angle) with $|Z| > 5$ is considered an outlier worth inspection. RMSZ is the root-mean-square of all Z scores of the bond lengths (or angles).

| Mol | Chain | Bond lengths | | Bond angles | |
|---|---|---|---|---|---|
| | | RMSZ | #\|Z\| >5 | RMSZ | #\|Z\| >5 |
| 1 | A | 0.76 | 0/1021 | 0.91 | 1/1379 (0.1%) |

There are no bond length outliers.

All (1) bond angle outliers are listed below:

| Mol | Chain | Res | Type | Atoms | Z | Observed($^o$) | Ideal($^o$) |
|---|---|---|---|---|---|---|---|
| 1 | A | 45 | ARG | CG-CD-NE | -5.17 | 100.94 | 111.80 |

There are no chirality outliers.

There are no planarity outliers.

#### 5.2 Too-close contacts (i)

In the following table, the Non-H and H(model) columns list the number of non-hydrogen atoms and hydrogen atoms in the chain respectively. The H(added) column lists the number of hydrogen atoms added and optimized by MolProbity. The Clashes column lists the number of clashes within the asymmetric unit, whereas Symm-Clashes lists symmetry-related clashes.

| Mol | Chain | Non-H | H(model) | H(added) | Clashes | Symm-Clashes |
|---|---|---|---|---|---|---|
| 1 | A | 1001 | 0 | 958 | 5 | 0 |
| 2 | A | 1 | 0 | 0 | 0 | 0 |
| 3 | A | 7 | 0 | 0 | 0 | 0 |
| 4 | A | 12 | 0 | 16 | 0 | 0 |
| 5 | A | 27 | 0 | 0 | 1 | 0 |
| All | All | 1048 | 0 | 974 | 5 | 0 |

The all-atom clashscore is defined as the number of clashes found per 1000 atoms (including hydrogen atoms). The all-atom clashscore for this structure is 3.

For Manuscript Review

All (5) close contacts within the same asymmetric unit are listed below, sorted by their clash magnitude.

| Atom-1 | Atom-2 | Interatomic distance (Å) | Clash overlap (Å) |
|---|---|---|---|
| 1:A:59:ASN:ND2 | 1:A:61:ARG:H | 1.99 | 0.60 |
| 1:A:45:ARG:NH2 | 5:A:301:HOH:O | 2.37 | 0.58 |
| 1:A:59:ASN:ND2 | 1:A:61:ARG:HB3 | 2.19 | 0.57 |
| 1:A:15:HIS:HB3 | 1:A:92:VAL:HG11 | 1.93 | 0.49 |
| 1:A:59:ASN:ND2 | 1:A:61:ARG:CB | 2.83 | 0.41 |

There are no symmetry-related clashes.

#### 5.3 Torsion angles (i)

##### 5.3.1 Protein backbone (i)

In the following table, the Percentiles column shows the percent Ramachandran outliers of the chain as a percentile score with respect to all X-ray entries followed by that with respect to entries of similar resolution.

The Analysed column shows the number of residues for which the backbone conformation was analysed, and the total number of residues.

| Mol | Chain | Analysed | Favoured | Allowed | Outliers | Percentiles | |
|---|---|---|---|---|---|---|---|
| 1 | A | 127/129 (98%) | 125 (98%) | 2 (2%) | 0 | 100 | 100 |

There are no Ramachandran outliers to report.

##### 5.3.2 Protein sidechains (i)

In the following table, the Percentiles column shows the percent sidechain outliers of the chain as a percentile score with respect to all X-ray entries followed by that with respect to entries of similar resolution.

The Analysed column shows the number of residues for which the sidechain conformation was analysed, and the total number of residues.

| Mol | Chain | Analysed | Rotameric | Outliers | Percentiles | |
|---|---|---|---|---|---|---|
| 1 | A | 105/105 (100%) | 102 (97%) | 3 (3%) | 42 | 13 |

All (3) residues with a non-rotameric sidechain are listed below:

For Manuscript Review

| Mol | Chain | Res | Type |
|---|---|---|---|
| 1 | A | 55 | ILE |
| 1 | A | 61 | ARG |
| 1 | A | 77 | ASN |

Sometimes sidechains can be flipped to improve hydrogen bonding and reduce clashes. All (3) such sidechains are listed below:

| Mol | Chain | Res | Type |
|---|---|---|---|
| 1 | A | 15 | HIS |
| 1 | A | 46 | ASN |
| 1 | A | 59 | ASN |

##### 5.3.3 RNA (i)

There are no RNA molecules in this entry.

#### 5.4 Non-standard residues in protein, DNA, RNA chains (i)

There are no non-standard protein/DNA/RNA residues in this entry.

#### 5.5 Carbohydrates (i)

There are no monosaccharides in this entry.

#### 5.6 Ligand geometry (i)

Of 10 ligands modelled in this entry, 8 are monoatomic - leaving 2 for Mogul analysis.

In the following table, the Counts columns list the number of bonds (or angles) for which Mogul statistics could be retrieved, the number of bonds (or angles) that are observed in the model and the number of bonds (or angles) that are defined in the Chemical Component Dictionary. The Link column lists molecule types, if any, to which the group is linked. The Z score for a bond length (or angle) is the number of standard deviations the observed value is removed from the expected value. A bond length (or angle) with $|Z| > 2$ is considered an outlier worth inspection. RMSZ is the root-mean-square of all Z scores of the bond lengths (or angles).

| Mol | Type | Chain | Res | Link | Bond lengths | | | Bond angles | | |
|---|---|---|---|---|---|---|---|---|---|---|
| | | | | | Counts | RMSZ | $\#\|Z\| > 2$ | Counts | RMSZ | $\#\|Z\| > 2$ |
| 4 | GOL | A | 209 | - | 5,5,5 | 0.09 | 0 | 5,5,5 | 0.25 | 0 |
| 4 | GOL | A | 210 | - | 5,5,5 | 0.07 | 0 | 5,5,5 | 0.24 | 0 |

For Manuscript Review

In the following table, the Chirals column lists the number of chiral outliers, the number of chiral centers analysed, the number of these observed in the model and the number defined in the Chemical Component Dictionary. Similar counts are reported in the Torsion and Rings columns. '-' means no outliers of that kind were identified.

| Mol | Type | Chain | Res | Link | Chirals | Torsions | Rings |
|---|---|---|---|---|---|---|---|
| 4 | GOL | A | 209 | - | - | 2/4/4/4 | - |
| 4 | GOL | A | 210 | - | - | 2/4/4/4 | - |

There are no bond length outliers.

There are no bond angle outliers.

There are no chirality outliers.

All (4) torsion outliers are listed below:

| Mol | Chain | Res | Type | Atoms |
|---|---|---|---|---|
| 4 | A | 209 | GOL | C1-C2-C3-O3 |
| 4 | A | 209 | GOL | O2-C2-C3-O3 |
| 4 | A | 210 | GOL | O1-C1-C2-O2 |
| 4 | A | 210 | GOL | O1-C1-C2-C3 |

There are no ring outliers.

No monomer is involved in short contacts.

#### 5.7 Other polymers (i)

There are no such residues in this entry.

#### 5.8 Polymer linkage issues (i)

There are no chain breaks in this entry.

For Manuscript Review

### 6 Fit of model and data (i)

#### 6.1 Protein, DNA and RNA chains (i)

In the following table, the column labelled '#RSRZ> 2' contains the number (and percentage) of RSRZ outliers, followed by percent RSRZ outliers for the chain as percentile scores relative to all X-ray entries and entries of similar resolution. The OWAB column contains the minimum, median, $95^{th}$ percentile and maximum values of the occupancy-weighted average B-factor per residue. The column labelled 'Q< 0.9' lists the number of (and percentage) of residues with an average occupancy less than 0.9.

| Mol | Chain | Analysed | <RSRZ> | #RSRZ>2 | | | OWAB(Å$^2$) | Q<0.9 |
|---|---|---|---|---|---|---|---|---|
| 1 | A | 129/129 (100%) | 0.15 | 5 (3%) | 39 | 44 | 15, 24, 45, 53 | 0 |

All (5) RSRZ outliers are listed below:

| Mol | Chain | Res | Type | RSRZ |
|---|---|---|---|---|
| 1 | A | 71 | GLY | 2.7 |
| 1 | A | 47 | THR | 2.6 |
| 1 | A | 56 | LEU | 2.3 |
| 1 | A | 55 | ILE | 2.1 |
| 1 | A | 121 | GLN | 2.0 |

#### 6.2 Non-standard residues in protein, DNA, RNA chains (i)

There are no non-standard protein/DNA/RNA residues in this entry.

#### 6.3 Carbohydrates (i)

There are no monosaccharides in this entry.

#### 6.4 Ligands (i)

In the following table, the Atoms column lists the number of modelled atoms in the group and the number defined in the chemical component dictionary. The B-factors column lists the minimum, median, $95^{th}$ percentile and maximum values of B factors of atoms in the group. The column labelled 'Q< 0.9' lists the number of atoms with occupancy less than 0.9.

| Mol | Type | Chain | Res | Atoms | RSCC | RSR | B-factors(Å$^2$) | Q<0.9 |
|---|---|---|---|---|---|---|---|---|
| 4 | GOL | A | 209 | 6/6 | 0.40 | 0.62 | 89,93,94,95 | 0 |
| 3 | CL | A | 205 | 1/1 | 0.43 | 0.28 | 76,76,76,76 | 0 |

*Continued on next page...*

For Manuscript Review

*Continued from previous page...*

| Mol | Type | Chain | Res | Atoms | RSCC | RSR | B-factors($Å^2$) | Q<0.9 |
|---|---|---|---|---|---|---|---|---|
| 4 | GOL | A | 210 | 6/6 | 0.47 | 0.38 | 82,83,86,86 | 0 |
| 3 | CL | A | 206 | 1/1 | 0.54 | 0.34 | 70,70,70,70 | 0 |
| 3 | CL | A | 207 | 1/1 | 0.72 | 0.20 | 60,60,60,60 | 0 |
| 3 | CL | A | 208 | 1/1 | 0.85 | 0.26 | 66,66,66,66 | 0 |
| 3 | CL | A | 203 | 1/1 | 0.96 | 0.09 | 43,43,43,43 | 0 |
| 3 | CL | A | 202 | 1/1 | 0.98 | 0.04 | 28,28,28,28 | 0 |
| 2 | NA | A | 201 | 1/1 | 0.98 | 0.08 | 26,26,26,26 | 0 |
| 3 | CL | A | 204 | 1/1 | 0.98 | 0.11 | 42,42,42,42 | 0 |

#### 6.5 Other polymers (i)

There are no such residues in this entry.

For Manuscript Review
